# A Thousand Meters Deep: Vertical Profiling of the Subterranean Microbes of Gourgouthakas Cave

**DOI:** 10.64898/2026.03.19.712943

**Authors:** Savvas Paragkamian, Christos A. Christakis, Vassiliki A. Michalopoulou, Maria Plakogiannaki, Stefanos Soultatos, Nikolaos P. Arapitsas, Markos Vaxevanopoulos, Yorgos Sotiriadis, Christos Pennos, Emmanouil A. Markakis, Panagiotis F. Sarris

**Affiliations:** Department of Biology, University of Crete, 714 09 Heraklion, Crete, Greece; Hellenic Institute of Speleological Research, Irakleio, Crete, Greece; Institute of Molecular Biology and Biotechnology, Foundation for Research and Technology-Hellas, 714 09 Heraklion, Crete, Greece; Department of Agriculture, School of Agricultural Sciences, Hellenic Mediterranean University, Estavromenos, 71410, Heraklion, Greece; Department Of History, Archaeology and Social Anthropology, University of Thessaly, Greece; Proteas caving club, 54636 Thessaloniki, Greece; School of Geology, Department of Physical Geography, Aristotle University of Thessaloniki, Thessaloniki, Greece; Department of Biosciences, College of Life and Environmental Sciences, University of Exeter, Exeter, UK

**Keywords:** Subterranean microbiology, Gourgouthakas cave, Bioprospecting, Bacterial diversity, Antimicrobial activity, Phytopathogens, *Streptomyces*, Biocontrol.

## Abstract

**Introduction:** Caves represent unique, nutrient-limited windows into the deep biosphere, yet the microbiology of the deep terrestrial subsurface remains remarkably under-explored. In this work, we conducted a rare expedition into Gourgouthakas Cave (Crete, Greece), one of the world’s deepest vertical systems, which had remained untouched by humans for 19 years.

**Methods:** We performed a high-resolution vertical profiling of the cavès microbes by sampling rock surfaces across nine different depths down to 1,100 meters. Through extensive cultivation on various media and at different temperatures, we established a biobank of 820 bacterial isolates.

**Results:** Taxonomic identification of a 374-isolate subset revealed a diverse community spanning 35 genera and 4 phyla, dominated by *Pseudomonas*, *Aquipseudomonas*, *Bacillus*, and *Stenotrophomonas*.

Beyond characterizing this taxonomic diversity, we explored the biotechnological potential of these subterranean microbes against major agricultural threats. Screening 70 representative isolates against six key pathogens, including *Ralstonia solanacearum*, *Verticillium dahliae*, and *Phytophthora nicotianae*, uncovered a notable group of strains with potent antagonistic activity, particularly within the *Pseudomonas* and *Brevibacillus* groups. Genomic sequencing of cave-derived *Actinobacteria* (*Streptomyces* and *Nocardiopsis* isolates) further highlighted this potential, revealing 142 biosynthetic gene clusters (BGCs), over half of which showed little to no similarity to known clusters, suggesting a hidden reservoir of novel secondary metabolites. Pangenomic analysis of *Streptomyces* revealed 1,497 unique gene clusters. Finally, *ex vivo* trials showed that the *Aquipseudomonas paracarnis* (formerly *Pseudomonas sp.*) isolate SRL917 significantly reduced *Botrytis cinerea* infections on tomato leaves, even surpassing the performance of a commercial biocontrol agent.

**Discussion:** Collectively, our results demonstrate that deep karstic systems are not merely geological wonders but vital hotspots for microbial innovation with tangible applications for sustainable agriculture.

## 1 Introduction

Caves are lightless subterranean ecosystems that are large enough for humans to explore (Culver and Pipan, 2019). Cave environments have been studied for centuries, and cave life in particular has offered novel insights into dispersal, adaptation, and the evolution of novel taxa ranging from bacteria to invertebrates (arthropods, molluscs, etc.) to chordates (fish, mammals, etc.) (Mammola et al., 2020). Their isolation, coupled with their distinctive abiotic conditions, makes caves ideal as natural laboratories (Poulson and White, 1969) and promising settings for exploring possible extraterrestrial life signatures (Boston et al., 2001). Unique ecosystems exist in caves such as the chemoautotrophic Movile Cave in Romania, where microbes produce the organic carbon used by the rest of the biotic community (Sarbu et al., 1996). Bacterial diversity has also been explored in two of the world’s deepest caves, in the Caucasus: in the Veryovkina cave (deepest sample at 2204m; Hodzhev et al., 2026), and in Krubera-Voronja Cave (deepest sample at 1640m; Kieraite-Aleksandrova et al., 2015), more than 20 bacterial phyla were present in total, yet owing to low nutrient availability the recovered DNA was relatively scarce compared with other systems such as soil. In general, even in the isolated subterranean systems, biomass distribution projections estimate that bacteria are the most abundant organisms, followed by arthropods (Bar-On et al., 2018).

There are two complementary approaches to studying microbes in extreme habitats such as caves (Tighe et al., 2017): culturing the microbial isolates (Mikell et al., 1996), and applying DNA sequencing technologies (Pace, 1997; Barton et al., 2004). In terms of biodiversity, the isolates capture only a fraction of the microbial community present, whereas shotgun metagenomics can recover both its functional and taxonomic diversity (Wu et al., 2025). The benefits of isolating microbial strains from extreme environments are multifaceted, ranging from characterizing Earth’s biodiversity, to investigating the origins of life, to bioprospecting and human-health advances, including the supply of beneficial microbes for agrifood (Schultz et al., 2023). Still, the majority of microbial lineages and functions remain uncultured (Lloyd et al., 2018).

Isolating microbial strains from extreme environments such as caves is a laborious, multi-step process: transferring the samples to the laboratory, handling them across multiple conditions (temperature, nutrients, pH, etc.), colony picking, culturing the isolates, identification, and cryopreservation. Sample processing is crucial for recovering as many strains as possible, and it is ecosystem-specific, as researchers aim to emulate natural conditions (Kaeberlein et al., 2002). For example, because limestone caves and karstic aquifers are resource-limited environments, nutrient-poor media are preferred for cultivation (Bender et al., 2020).

Microbes are the cornerstone of cave ecosystem processes (Engel, 2015), for example nutrient cycling, speleothem formation (secondary mineral deposition), and carbonate-rock precipitation (Northup and Lavoie, 2001). Oligotrophic, isolated caves are known as microbial reservoirs with antimicrobial functionality (Cheeptham, 2013). A notable example is the microbial collection from Lechuguilla Cave (New Mexico, USA), in which 93 isolates were tested against 26 different antimicrobial agents, yielding three strains resistant to 14 antibiotics (Bhullar et al., 2012). This collection also holds the *Paenibacillus* sp. LC231, a strain resistant to 26 antibiotics (Pawlowski et al., 2016). Lechuguilla is more than 200 km long and is closed to visitors, so human influence on its resistome is unlikely. In the Krubera-Voronja Cave isolates (the deepest cave in the world as of this article), cultured bacteria showed depth-dependent stratification of antimicrobial activity, with higher antagonistic activity reported in the deeper parts of the cave (Klusaite et al., 2016).

Here, we developed a biobank of bacterial isolates from Gourgouthakas Cave, a 1,129-meter-deep cave in Crete, Greece, to explore their diversity and functional properties. We profiled the cave vertically across its full depth and its variable rock substrates to sample its microbial communities. By isolating these microbes, we recovered several hundred strains spanning 35 genera and 4 phyla, all cryopreserved in a local biobank.

We further demonstrate the antimicrobial potential of Gourgouthakas microbes against six major crop pathogens, including pathogenic bacteria, fungi and oomycetes. Together, this work provides strain-level insights and showcases their potential applications using isolates from one of the deepest microbial cave samplings worldwide.

## 2 Material and Methods

### 2.1 Study site

Sampling was performed in Gourgouthakas Cave, a 1,129 m vertical cave with a 2,384 m horizontal extent, located in the Atzinolakas area of the Lefka Ori (White Mountains) massif, Crete, Greece (Figure 1). The massif is composed predominantly of carbonate rocks (limestones and dolomites), highly fractured and intersected by prominent tectonic faults. The carbonate bedrock is steeply inclined, and its rapid tectonic uplift favours the development of deep vertical cave systems (Moraetis et al., 2025). The entrance of Gourgouthakas lies at 1,550 m elevation in an area of high speleological interest (Adamopoulos, 2005). The cave is part of an extensive karst aquifer in which groundwater circulates through open vadose canyons and phreatic corridors; as part of the Koiliaris drainage basin, the system discharges at the Stylos spring, 50 km away of the cave entrance (Lilli et al., 2020). The site’s biodiversity is protected by the European Natura 2000 network (GR4340008; LEFKA ORI KAI PARAKTIA ZONI).

**Figure 1.**
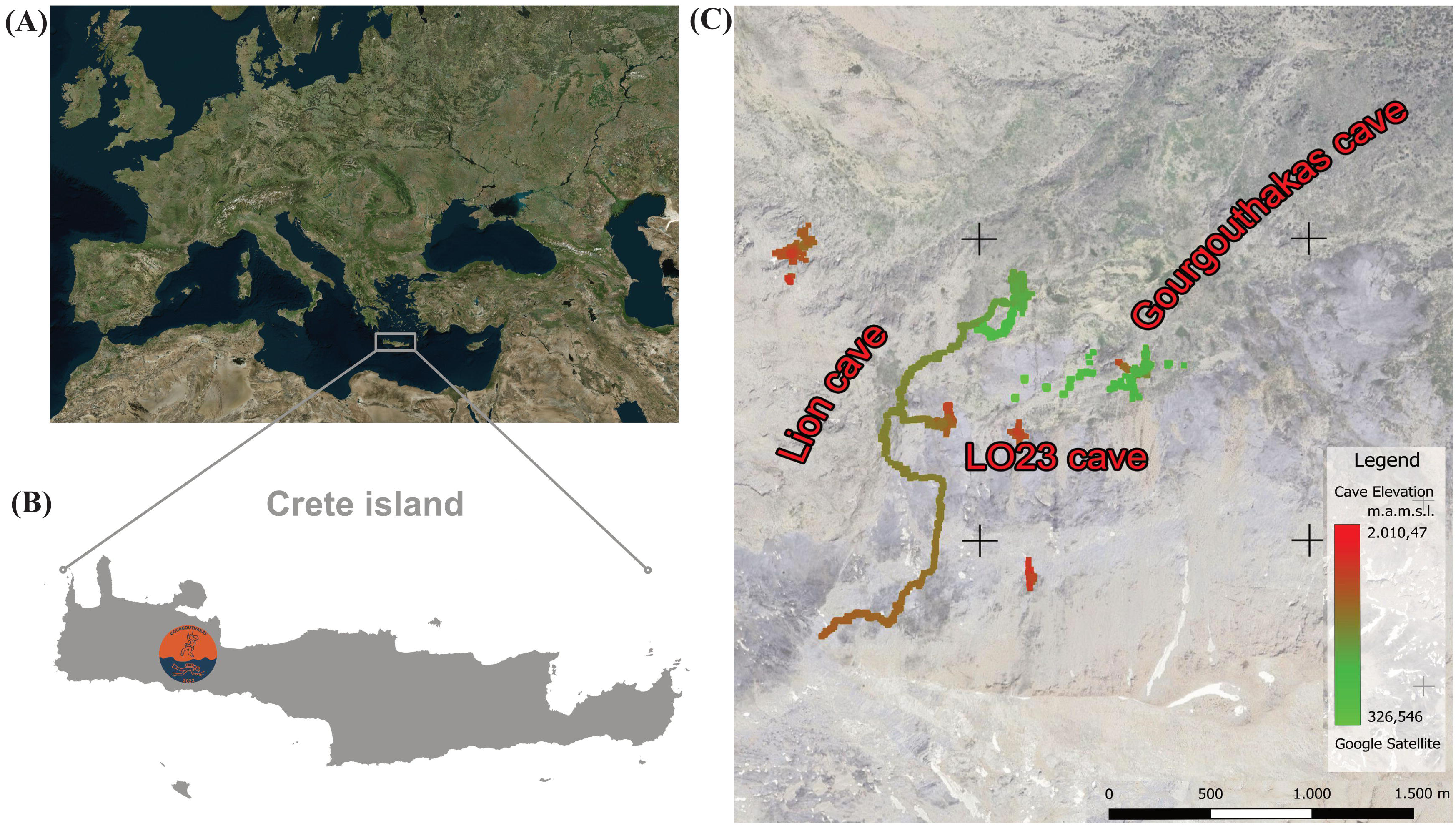
Maps of the location of Gourgouthakas cave. **(A)** Location of the cave in the eastern Mediterranean region. **(B)** Crete island and the area of interest (symbol of the expedition) at Lefka Ori (White Mountains). **(C)** Satellite image of the Atzinolakas area in the north-eastern part of Lefka Ori (White Mountains). Projected are the vertical profiles of the cave systems in the area, including Gourgouthakas cave on the right (map modified from Pennos et al. (2025) with cave data shared by Adamopoulos (2005)). The color of the cave passage corresponds to altitude, where greener hues indicate deeper points within the cave system.

### 2.2 Sampling

In the summer of 2022 the caving expedition “Gourgouthakas 2022” was organized, marking 19 years since the last human visit to the cave (Sotiriadis et al., 2023). This provided a unique opportunity for microbial sampling in the deepest cave in Greece. Sampling and photography during the expedition were authorized by the Ministry of Environment (ID: ΑΔΑ: Ψ50Η4653Π8-ΤΦΥ, available at https://diavgeia.gov.gr/). Expedition coordinators synchronized the sampling along with the rigging of the cave to minimize possible contamination. During sampling, we first descended to the bottom of the cave while identifying sampling sites. Sites were selected across depths, favouring locations a few meters away from the cavers’ route and with different rock substrate types. Sampling was initiated from the bottom of the cave at -1,100 meters.

At each site, disposable gloves, a sterile razor to scrape the rock surface, and sterile falcon tubes of 50 mL were used to collect and store the material. Two falcon tubes were used per site: one with material for culturing isolates and one for metagenomics. At each site, air temperature and relative humidity were also measured with a HANNA Instruments Portable Low Range EC/TDS Meter - HI99300 (accuracy: EC and TDS ±2% FS, temperature ±0.5°C). Μetagenomics samples were stored at -20 °C until their processing.

Sampling was carried out overnight at nine different depths (Figure 2), from the bottom to the entrance of the cave while ascending. Once the entrance was reached, the samples for the culturing of isolates were placed in a freezer box with ice and those for the metagenomics on dry ice, then immediately transferred to the nearest village and sent to our lab facilities in Heraklion, Crete. The transfer from the cave entrance to the laboratory took approximately 7 hours, the same period over which sample handling had began.

**Figure 2.**
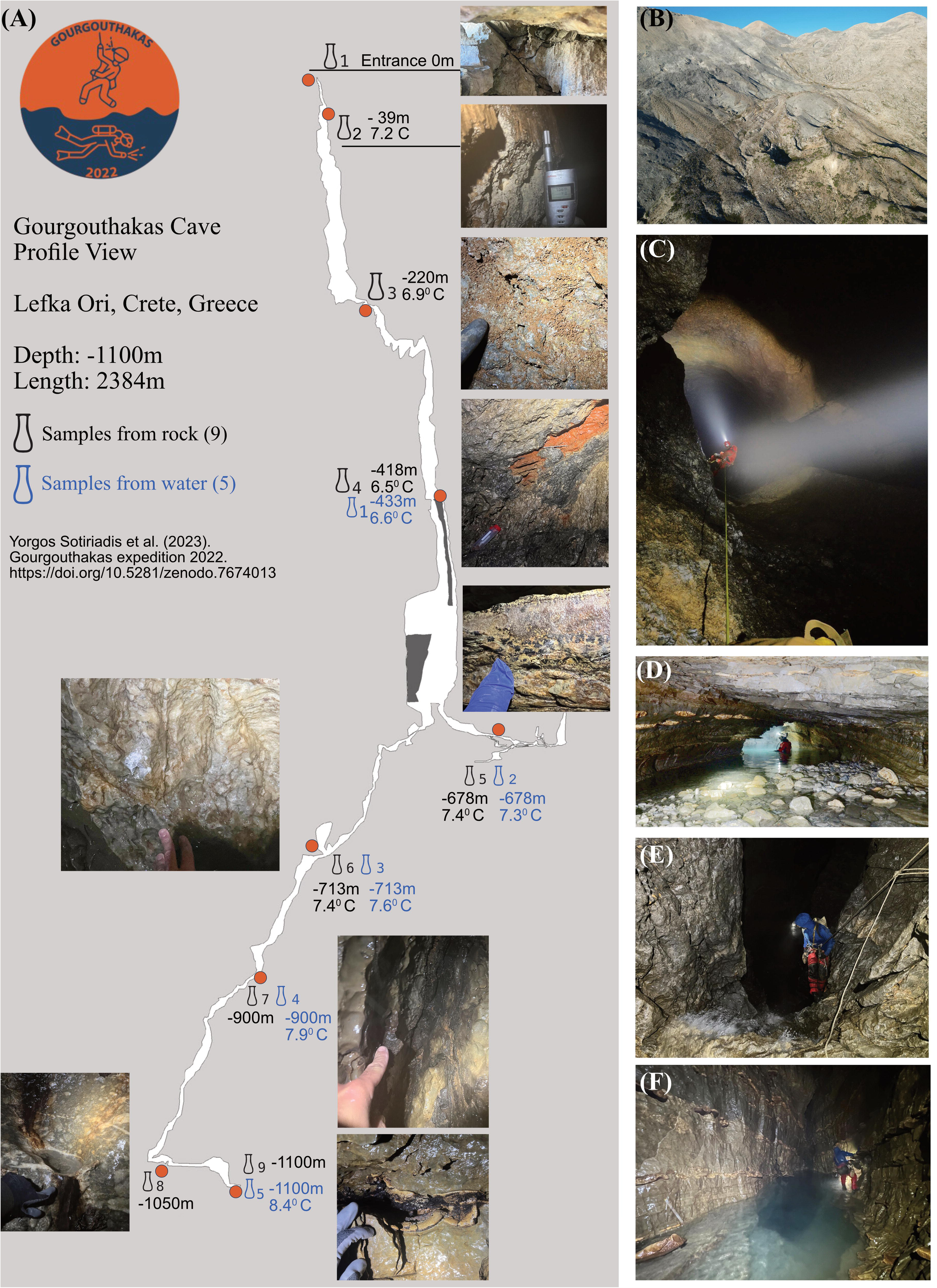
Gourgouthakas cave microbiome sampling and vertical profile. **(A)** Sampling sites along the cave with the respective picture of the substrate material (Photos S. Paragkamian). **(B)** The area Atzinolakas with the complex karst and faults (Photo M. Vaxevanopoulos). **(C)** One of the main pits of the cave, named Guillaume, at -450 m (Photo Y. Sotiriadis). **(D)** First siphon of the karstic aquifer at -678m (Photo Y. Sotiriadis). **(E)** Pit “Samuel” at -950m with constant flow of water (Photo M. Vaxevanopoulos). **(F)** Last siphon at -1100 m depth (Photo M. Vaxevanopoulos)

### 2.3 Isolation, culture and cryopreservation

The cultivation strategy was built on three axes: fast and minimal sample processing; multiple growth media; and different temperature conditions. First, environmental samples were processed shortly (<12 hours) after collection, under sterile conditions and with minimal dilution. 200 uL of sterile ultrapure water was used to hydrate each sample and allow plating on solid media. As a control, the same water aliquot was also plated to confirm the absence of microbial growth. Fast, minimal sample processing has been documented to increase colony-forming units during cultivation (Bender et al., 2020). Second, all samples and controls were plated on four different growth media, ranging from nutrient-rich to nutrient-poor:

1. Nutrient Agar (N: in g L⁻¹: 5 bacto-peptone, 3 yeast extract, 5 NaCl, 15 agar, pH 7.4);
2. Half-strength Nutrient Agar (½ NA; contained in g L⁻¹: 2.5 bacto-peptone, 1.5 yeast extract, 2.5 NaCl, 15 agar, pH 7.4);
3. Reasoner’s 2A agar (R2A; in g L⁻¹: 0.5 yeast extract, 0.5 proteose peptone, 0.5 casamino acids, 0.5 glucose, 0.5 soluble starch, 0.3 sodium-pyruvate, 0.3 K2PO4, 0.05 MgSO4·7H2O, 15 agar, pH 7.2); and
4. Mannitol Nitrogen Free Mineral Medium (MNFMM medium contained in g L⁻¹: CaCl₂ 1, NaCl 0.5, K₂HPO₄ 1, D-mannitol 7, Na₂MoO₄·2H₂O 0.01, MgSO₄·7H₂O 0.25, FeSO₄·7H₂O 0.01, MnSO₄ 0.01, agar 20, pH 6.8–7.2).

This range of nutrient conditions was used because extremophiles from oligotrophic biomes have been can be stressed by rich media, owing to their nutrient-poor native environment (Bender et al., 2020). In particular, R2A has been applied in many large-scale cultivations of cave microbes (Zhu et al., 2021; Reasoner and Geldreich, 1985). MNFMM medium was selected, despite its limited prior use on cave samples, for its ability to enable the isolation of nitrogen-fixing bacteria from the environment (Aquilanti et al., 2004).

Third, plates were incubated at 5 °C and 28 °C to allow the growth and isolation of both cold-adapted, slow-growing and mesophilic microorganisms (Schultz et al., 2023). Developing colonies were monitored periodically and repeatedly streaked to obtain pure isolates. Pure isolates were preserved in their respective liquid media supplemented with 50% (v/v) glycerol and stored at −80 °C.

### 2.4 Taxonomic identification using the 16S rRNA gene

For taxonomic identification, the full-length 16S rRNA gene (∼1,400bp) of the isolates was amplified. Cells were lysed by heating the bacterial cells at 98 °C for 10 min, followed by centrifugation of the lysate at 10,000 × g for 10 min. Primers 27F/1492R (27F: AGAGTTTGATCCTGGCTCAG, 1492R: GGTTACCTTGTTACGACTT) and DF Taq DNA polymerase (Enzyquest, PD010S) were used according to the manufacturer’s instructions.

The samples were shipped to Eurofins Scientific for Sanger sequencing using the SeqPlate&Clean service in 96-well plates. For samples that yielded with low quality sequences, the procedure was repeated and the samples were then sent for sequencing to a different commercial vendor (Genewiz).

For all samples, sequences were trimmed according to the quality of the resulting .ab1 chromatograms and subsequently processed for taxonomic identification. Based on isolateR workflow (Daisley et al., 2024) and extra analysis in Biopython (v1.87) (Cock et al., 2009), reads were quality-trimmed (isoQC step), using the “auto” cutoff, with a fixed sliding-window alternative (mean Phred ≥ 20 over a 15-bp window); reads shorter than 200 bp after trimming were discarded (Daisley et al., 2024). Taxonomic assignment (isoTAX) was performed with vsearch (v2.31.0) (Rognes et al., 2016) (--usearch_global) against three reference databases - the NCBI 16S RefSeq Targeted Loci type-strain database (Schoch et al., 2020), the SILVA SSU Ref NR99 release 138.2 (Chuvochina et al., 2026) and the GTDB SSU representative database (Parks et al., 2025)(bac120_ssu_reps + ar53_ssu_reps). Full lineages were retrieved with taxonkit (v0.20.0) from the NCBI taxonomy, and each query was assigned to the most specific rank passing the isolateR percent-identity cutoffs (phylum 75.0, class 78.5, order 82.0, family 86.5, genus 96.5, and species 98.7 %). The three taxonomic databases were used for completeness and complementarity of the results (Vinje et al., 2025).

For phylogenetic context, the GTDB bac120 master tree was pruned to the best-hit reference genomes of the isolates with the keep.tip function of the R package ape (Paradis and Schliep, 2019). The resulting tree was visualized in R (v4.5.2) (R Core Team, 2025) with tidytree (Yu, 2023) and treeio (Wang et al., 2020), collapsed to one representative tip per genus and annotated with per-isolate abundances across the nine cave sampling depths (0 to −1,100 m) drawn from the joined isolate metadata.

### 2.5 *In vitro* assays

The *in vitro* inhibitory activity of selected bacterial isolates was evaluated against several plant pathogens at 28 °C. Representative strains were selected for each identified genus and screened against *Paracidovorax citrulli*, *Clavibacter michiganensis*, *Ralstonia solanacearum* and *Xanthomonas campestris* pv. *campestris* using an overlay assay on NA plates. Briefly, each pathogen was grown to exponential phase in liquid NB, and then plated on NA. Concurrently, each bacterial strain was grown on fresh NA, and then a single colony was then spotted onto the NA plates for the overlay assay. Antagonistic activity was recorded by monitoring the growth inhibition zones (in cm) around the bacterial colonies up to 72 hours post-inoculation (hpi).

Antifungal and anti-oomycete activities were also evaluated against *Verticillium dahliae* and *Phytophthora nicotianae*, respectively, using dual-culture assays on PDA plates (Potato (infusion form) 200 g L^-1^, Dextrose 20 g L^-1^, Agar 15 g L^-1^) plates. Briefly, the pathogens were grown on PDA, and metabolically active plugs were transferred to fresh PDA plates. Concurrently, each bacterial isolate was grown on fresh NA, and a single colony was then streaked vertically at a distance of 4 cm from each pathogen. Growth reduction was classified follows: 1 = inhibition, 0.5 = partial inhibition, 0 = no inhibition. Cases of space limitation due to the motility of bacterial isolates were also noted.

The haemolytic activity of the *Pseudomonas aeruginosa* isolates was assessed by plating isolates on Columbia agar supplemented with 5% sheep blood (COS; bioMérieux) and incubating the plates at 28 °C for 24–48 h. of Haemolysis was scored by inspecting the plates for clear zones surrounding the bacterial colonies.

### 2.6 Whole genome sequencing and analysis

Following the taxonomic identification and the *in vitro* assays, three *Streptomyces* isolates, one *Nocardiopsis* isolate and one *Aquipseudomonas* isolate were selected for whole-genome sequencing . High-quality genomic DNA was extracted using a phenol-based protocol optimized for Gram-positive and Gram-negative bacteria (Kiledal and Maresca, 2023). After high-molecular-weight genomic DNA extraction using the phenol-based protocol, DNA concentration was quantified with a Qubit 4.0 Fluorometer (Qubit dsDNA HS Assay Kit, Thermo Fisher Scientific), and purity was assessed with a NanoDrop spectrophotometer; only samples meeting strict quality thresholds (A260/280 ratio of 1.8–2.0 and A260/230 ratio of 2.0–2.2) were accepted.

Hybrid DNA sequencing was performed at BGI Genomics (*Streptomyces, Nocardiopsis*) and at the Laboratory of Clinical Microbiology and Microbial Pathogenesis, School of Medicine, University of Crete (*Aquipseudomonas*), combining short reads (DNBSEQ, 150 bp paired-end for *Streptomyces* and *Nocardiopsis*, and Illumina NextSeq2000, 150 bp paired-end for *Aquipseudomonas*) and long reads (Oxford Nanopore Technologies with PromethION and MinION Mk1B). The sequences were trimmed using fastp 1.0 and fastplong (Chen, 2025) with default parameters. Short reads were filtered based on quality and removal of N-contenting reads (if 0.1% of a read is N it was discarded). Long reads were filtered to remove reads shorter than 2,000 bp or with mean quality below 9.

The genomes were assembled with Unicycler, following a hybrid assembly pipeline (Wick et al., 2025). The assembled genomes were annotated with Bakta v1.11.2 using the local reference database v6.0 (Beyvers et al., 2025). Taxonomic placement was performed with GTDB-Tk 2.5.2 using the r226 reference data (Parks et al., 2025), and the biosynthetic gene clusters were identified with antiSMASH v8.0 (Blin et al., 2025).

Publicly available *Streptomyces* genomes were retrieved from NCBI using ncbi-genome-download bacteria (Blin, 2023; O’Leary et al., 2024) and ere selected at the contig, chromosome, or complete-genome assembly levels. Pangenomic analyses were performed with anvi’o v9 (Eren et al., 2021). Genome storage databases were generated with anvi-gen-genomes-storage, genome annotations were performed with anvi-run-hmms using the -I Bacteria_71, along with --also-scan-trnas options (Eddy, 2011), anvi-run-ncbi-cogs (Galperin et al., 2021), anvi-run-kegg-kofams (Kanehisa et al., 2020), anvi-run-pfams (Mistry et al., 2021), and anvi-run-scg-taxonomy (Parks et al., 2025). The pangenome was then constructed with anvi-pan-genome using --minbit 0.5, --mcl-inflation 10, and --use-ncbi-blast (Eren et al., 2021). Gene cluster presence/absence data were extracted from the resulting PAN.db files and processed with sqlite3 v3.45.3 and gawk v5.3.0 to count gene cluster occurrences per genome. Visualizations were generated in R v4.5.2 (R Core Team, 2025) using tidyverse v2.0.0 (Wickham et al., 2019) and UpSetR v1.4.0 (Conway et al., 2017).

To identify gene clusters specific to our newly sequenced cave isolates, the gene cluster membership table from the anvi’o PAN.db output was exported with anvi-export-gene-clusters, and functional annotations from the corresponding contigs databases with anvi-export-functions. We then identified the SRL1060-specific gene clusters, the gene clusters uniquely shared by SRL1060, SRL740, and SRL742, and those uniquely shared by SRL740 and SRL742. Genes belonging to these cluster categories were then extracted, matched to their functional annotations, and summarized accordingly.

### 2.7 *Ex vivo* assay

The bacterial isolate SRL917 was cultivated in 200 mL tryptic soy broth (TSB) in Erlenmeyer flasks at 28 °C with shaking (180 rpm) for 48 h in the dark. Cells were harvested by centrifugation (3,000 × g, 10 min) and resuspended in sterile water to a final concentration of 10⁸ CFU x mL⁻¹, determined by dilution plating. Detached tomato leaflets (hybrid Belladonna F1) were immersed in the bacterial suspension for 2 min. A commercially registered product (X) was also included, which was applied according to the manufacturer’s instructions to allow comparison with the bacterial treatment.

The leaflets were inoculated with *Botrytis cinerea* (isolate B.14.9) two days after the bacterial treatment. The conidial suspension was prepared from a 10-day-old PDA culture and the concentration was adjusted to 1 × 10⁶ conidia x mL⁻¹ using a haemocytometer. For infection, two small wounds were made on each leaflet with a sterile dissecting needle, and 10 μL of the conidial suspension was applied to each wound; control leaflets received 10 μL of sterile distilled water.

The leaflets were placed on plastic racks inside transparent boxes (50 × 38 × 36 cm) lined with moist tissue paper to maintain ∼100% relative humidity and incubated in a growth chamber at 20 ± 2 °C under fluorescent light (12 h photoperiod) for 6 days. Disease symptoms were recorded by measuring the diameter of the necrotic lesion around each wound. Disease severity was plotted over time to construct disease progress curves and the area under the disease progress curve (AUDPC) was calculated using the trapezoidal integration method (Campbell and Madden, 1990). Statistical tests were performed with R programming language v4.5.2 (R Core Team, 2025) using the aov() function for ANOVA and TukeyHSD() for post-hoc Tukey HSD tests.

## 3 Results and Discussion

### 3.1 Vertical microbial isolates profiling

The “Gourgouthakas 2022” expedition (Sotiriadis et al., 2023) provided a unique opportunity for microbial sampling along a 1,100-meter vertical extent of a complex and important karst aquifer (Lilli et al., 2020) on the western part of Crete. The cave had been untouched by human visitors for 19 years, reducing the risk of contamination; cavers are known to alter native microbial communities, particularly in vertical, narrow caves where they increase diversity, as shown in the world’s deepest caves in the Caucasus (Kieraite-Aleksandrova et al., 2015; Hodzhev et al., 2026). In addition, having a microbiology laboratory close to the cave allowed rapid sample shipment and immediate isolation; transfer time from caves and other extreme habitats to the laboratory, is often a limiting step in the isolation of microbes (Bender et al., 2020).

A total of 820 isolates were obtained from nine rock-surface samples (Figure 2), each from a different depth, ranging from −1,100 m to the surface (0 m) (Supplementary Presentation 1). Humidity and temperature were constant along the cave, at 100% and 7.2-8.4 °C, respectively (Figure 2A). The highest number of isolates was obtained at −900 m (177 isolates), followed by −1,100 m (143), −713 m (125), −1,050 m (118), −678 m (74), −418 m (50), −220 m (48), −39 m (47), and 0 m (38 isolates). Surface samples yielded fewer isolates due to fungal overgrowth (Figure 3B). Among the growth media, the highest number of isolates grew on R2A (363 isolates), followed by ½ NA (218), NA (137), and MNFMM (101). Most isolates grew at 28 °C (574 isolates), while 236 isolates grew at 5 °C, although not all isolates were tested for psychrophilic/psychrotolerant growth (Supplementary Table 1). For example, 10 isolates that grew at 5 °C, but also grew at 28 °C. The resulting biobank comprised phenotypically diverse isolates, varying in texture and colour (e.g. orange, iridescent, purple) (Figure 3C).

**Figure 3.**
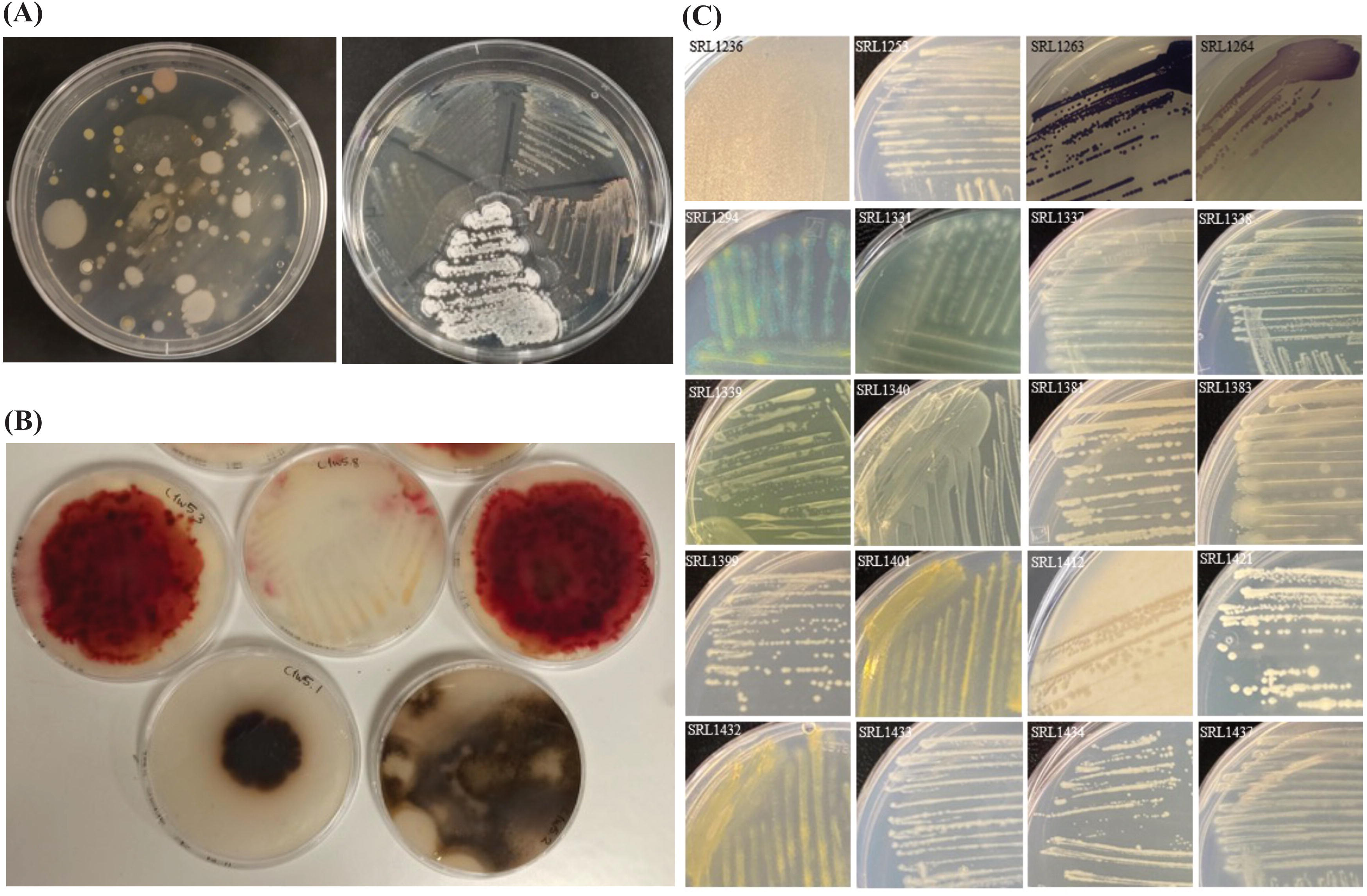
Selected pictures of microbial diversity of Gourgouthakas isolated bacteria. **(A-B)** Initial screenings of sample material prior to pure cultures. **(C)** A “color palette” of isolated microbes from the collection of Gourgouthakas cave. Variety of colors from yellow, purple and bioiridescent colonies.

A total of 374 isolates were identified to genus or species level across taxonomies using the 16S rRNA gene (Figure 4). The taxon identification varied with the reference database: 28 genera from SILVA, 27 genera from NCBI, and 35 genera from GTDB (Supplementary Table 2). Under GTDB, the most abundant genus was *Aquipseudomonas* (146 isolates), followed by *Bacillus_A* (44 isolates) and *Stenotrophomonas* (34 isolates). Other relatively abundant genera included *Paenibacillus* (Paenibacillus_AQ: 8; *Paenibacillus*: 5; *Paenibacillus_B*: 4 isolates), *Brevibacillus_B* (16 isolates), *Peribacillus* (15 isolates), *Flavobacterium* (11 isolates), *Variovorax* (8 isolates), *Brevundimonas* (7 isolates), and *Janthinobacterium* (7 isolates). This profiling of isolated microbes showcased the culturable microbial diversity of the cave, although culture-independent methods (i.g. metagenomics) are needed to fully grasp its microbiome (Hershey and Barton, 2018). However, even with state-of-the-art methods recover only a small percentage of the microbial diversity is culturable (Wu et al., 2025).

**Figure 4.**
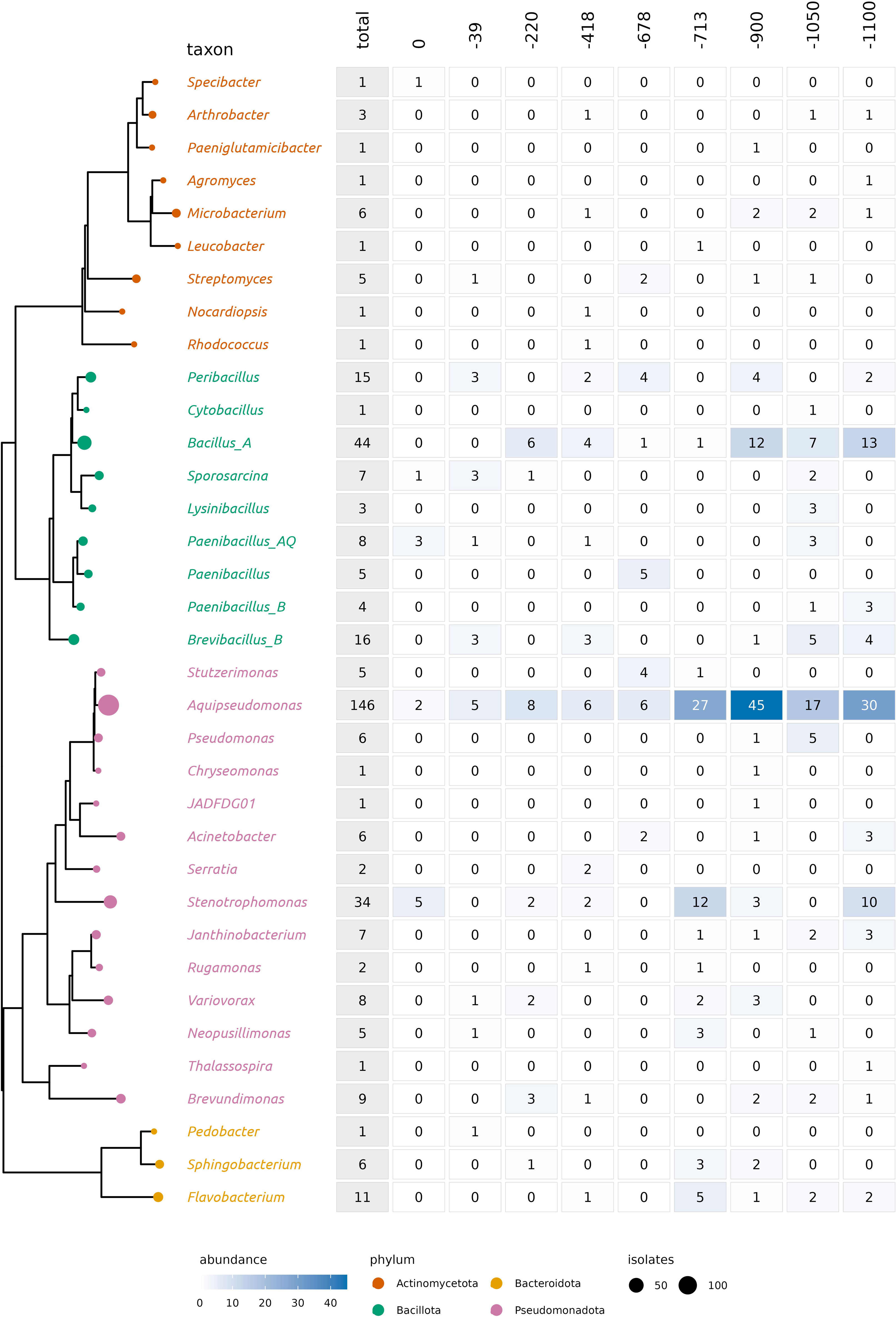
Diversity of genera of Gourgouthakas isolated bacteria. On the left is the pruned GTDB reference tree with the taxa from the biobank at the genus level. The size of the leaf node is relative to the number of isolates in that specific genus. The heatmap on the right breaks down the isolates based on the sample depth in the cave.

### 3.2 Pathogen growth-inhibition screening

Beneficial microbes have been studied for their role in inhibiting phytopathogen growth, with different mechanisms of action underlying the effect in each case. Biobanks such as the one established here are valuable resources for testing and detecting beneficial microbes. Direct inhibition often involves the secretion of antibiotics, lipopeptides, bacteriocins, biosurfactants, and/or volatile organic compounds, which interfere with the pathogen metabolism, membrane integrity, or cellular development (Caulier et al., 2019; Villavicencio-Vásquez et al., 2025). Alternatively, beneficial microbes may outcompete the pathogens for physical space and nutrients or bolster plant immunity via induced systemic resistance (ISR), enhancing the plant’s capacity to respond rapidly and effectively to subsequent pathogen attacks (Licciardello et al., 2018, Chepsergon and Moleleki, 2023, Pieterse et al., 2014).

To evaluate the phytopathogen inhibition capabilities of the isolates, 70 bacterial isolates from different taxonomic groups, but from the same cave origin, were used for direct inhibition studies. The phytopathogens tested are all known to infect economically important crops. Specifically, *Paracidovorax citrulli* (a Cucurbitaceae pathogen), *Clavibacter michiganensis* and *Ralstonia solanacearum* all have host ranges that include the Solanaceae, while *Xanthomonas campestris* pv. *campestris* is a major pathogen of Brassicaceae. Beyond bacterial pathogens, the fungus *Verticillium dahliae* (the agent of verticillium wilt in many plant species) and the oomycete *Phytophthora nicotianae* (a damaging Solanaceae pathogen) were also included. These antagonistic activities were demonstrated under *in vitro* conditions (Supplementary Presentations 2 and 3). Among the 70 isolates that were tested, 15 showed an inhibitory potential (Figure 5E), while several isolates of *Aquipseudomonas* sp. and *Brevibacillus* sp. outcompeted all of the pathogens tested (Figure 5A, B, C; Supplementary Table 3).

**Figure 5.**
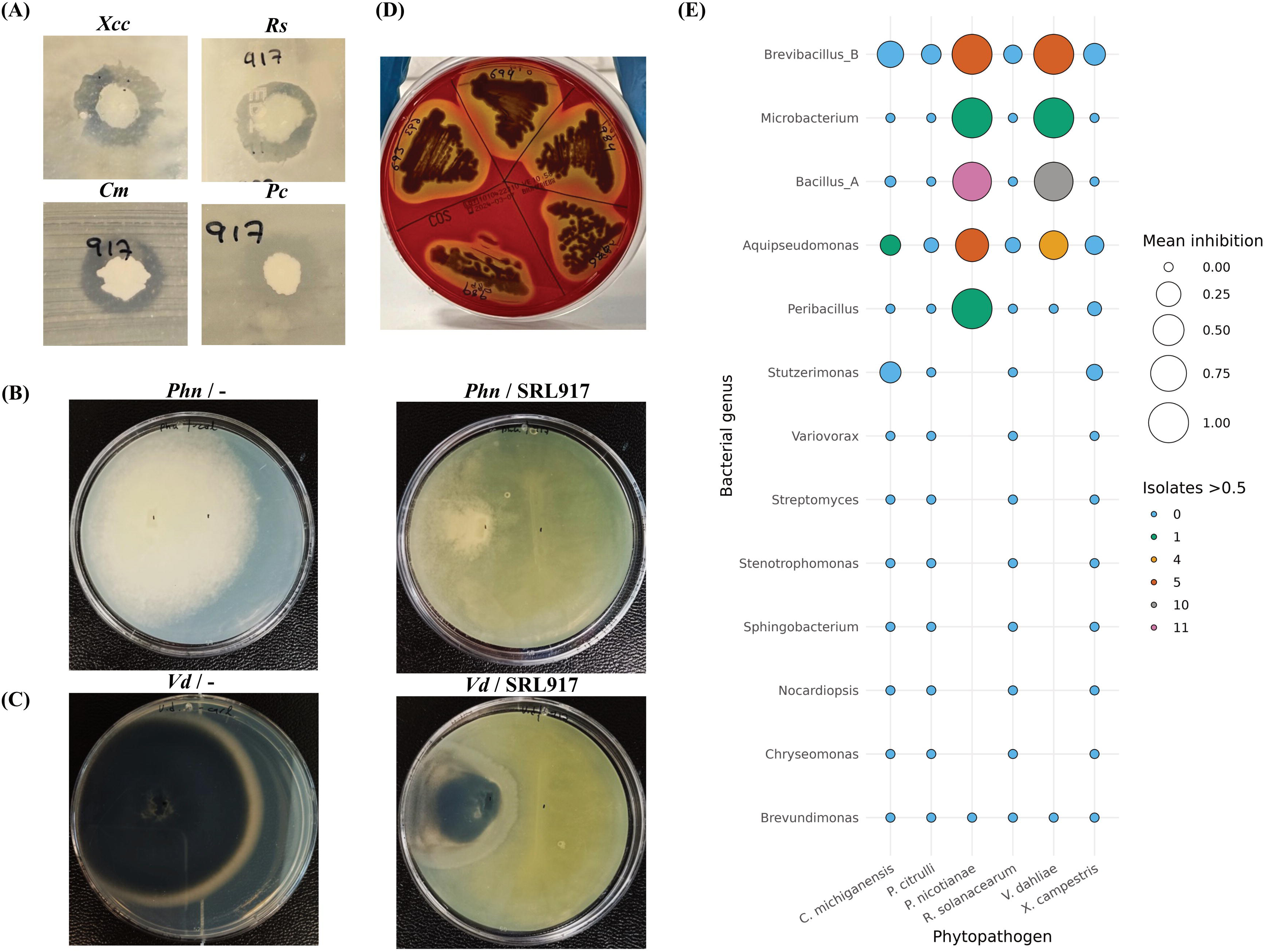
*In vitro* examination of phytopathogens growth by the beneficial *Aquipseudomonas paracarnis* SRL917. **(A)** Overlay inhibition assays of *Xanthomonas campestris* pv. *campestris* (*Xcc*), *Ralstonia solanacearum* (*Rs*), *Clavibacter michiganensis* (*Cm*) and *Paracidovorax citrulli* (*Pc*) with SRL917. The inhibition is shown by the clear absence of phytopathogens growth around the beneficial bacteria, at 72 h. **(B, C).** Dual inhibition assays between the SRL917 and the phytopathogens *Verticillium dahliae* (**B**) and *Phytophthora nicotianae* (**C**). At 17 days, the SRL917 shows space-limiting activity in PDA when grown together with the phytopathogens (right side) compared to the control plates (left side). (**D**) Various *Pseudomonas aeruginosa* isolates, from different depths, reveal strong hemolytic activity. **(E)** Antagonistic activity of selected isolates against plant pathogens.

Such *in vitro* assays provide a controlled framework for identifying promising microbial candidates and for elucidating the molecular and biochemical traits underlying microbial antagonism before further validation *in planta* /*ex vivo* or under field conditions (Christakis et al., 2021; Villavicencio-Vásquez et al., 2025).

### 3.3 Genomic analysis of selected *Actinobacteria* isolates

#### 3.3.1 *Streptomyces sp.* genomes

*Streptomyces* is one of the most widely studied bacterial genera for antibiotic and antimicrobial compound discovery (Quinn et al., 2020). Regarded as a cornerstone of industrial and medical biotechnology, *Streptomyces* strains produce over two-thirds of all clinically used, naturally derived antibiotics, alongside vital anticancer, immunosuppressive, and antiviral drugs (Alam et al., 2022). Recently, Kaur et al. (2026) found that *Streptomyces rimosus* produces a cyclic depsipeptide antibiotic, the manikomycin by improving the fractionation of previously overlooked minor products. Historically, the discovery of compounds such as streptomycin, the tetracyclines, and the immunosuppressant tacrolimus relied on culture-dependent, phenotypic screening (Alam et al., 2022). However, this traditional pipeline has reached a bottleneck of frequent compound rediscovery and an inability to access the true genetic capacity of these bacteria under standard laboratory conditions (Lee et al., 2020). Whole-genome sequencing has revealed that a single *Streptomyces* chromosome encodes far more biosynthetic gene clusters (BGCs), typically around 30 per species, than the fraction of secondary metabolites detectable in axenic culture (Lee et al., 2020). Moreover, cave-derived *Streptomyces* strains are promising sources of novel antibiotics, as subterranean environments harbor diverse actinobacteria with unique biosynthetic potential (Rangseekaew and Pathom-aree., 2019). For these reasons, we first investigated the whole genomes of several *Streptomyces* isolates from our biobank.

The genomes of three cave-derived *Streptomyces* isolates, SRL740, SRL742 and SRL1060 were sequenced and analyzed using a hybrid short- and long-read approach for each isolate. These isolates were recovered from distinct thin biofilms on the cave walls and all belong to *Streptomyces microflavus* species detected in samples across a broad depth range (678–1,050 m).

The resulting high-quality obtained genomes were examined *in silico* examined for their encoded biosynthetic gene clusters (BGCs), to estimate their potential to produce bioactive secondary metabolites, including antimicrobial compounds. A total of 116 BGCs were detected across the three genomes, approximately, 57% of which show low or no similarity to known clusters, suggesting the presence of biosynthetic features worthy of further investigation. SRL740 and SRL742 each contain 37 BGCs, while SRL1060 harbours 42 BGCs (Table 1). Most of the predicted clusters are associated with terpenes, followed by ribosomally synthesized and post-translationally modified peptides (RiPPs) (Table 1). BGCs associated with other diverse metabolites (e.g., ectoine, betalactone, butyrolactone and prodigiosin), non-ribosomal peptide synthetases (NRPS), polyketide synthases (PKS) and siderophores are also present (Table 1). Cave-derived *Streptomyces* can harbour diverse and novel BGCs within these classes.

**Table 1.**
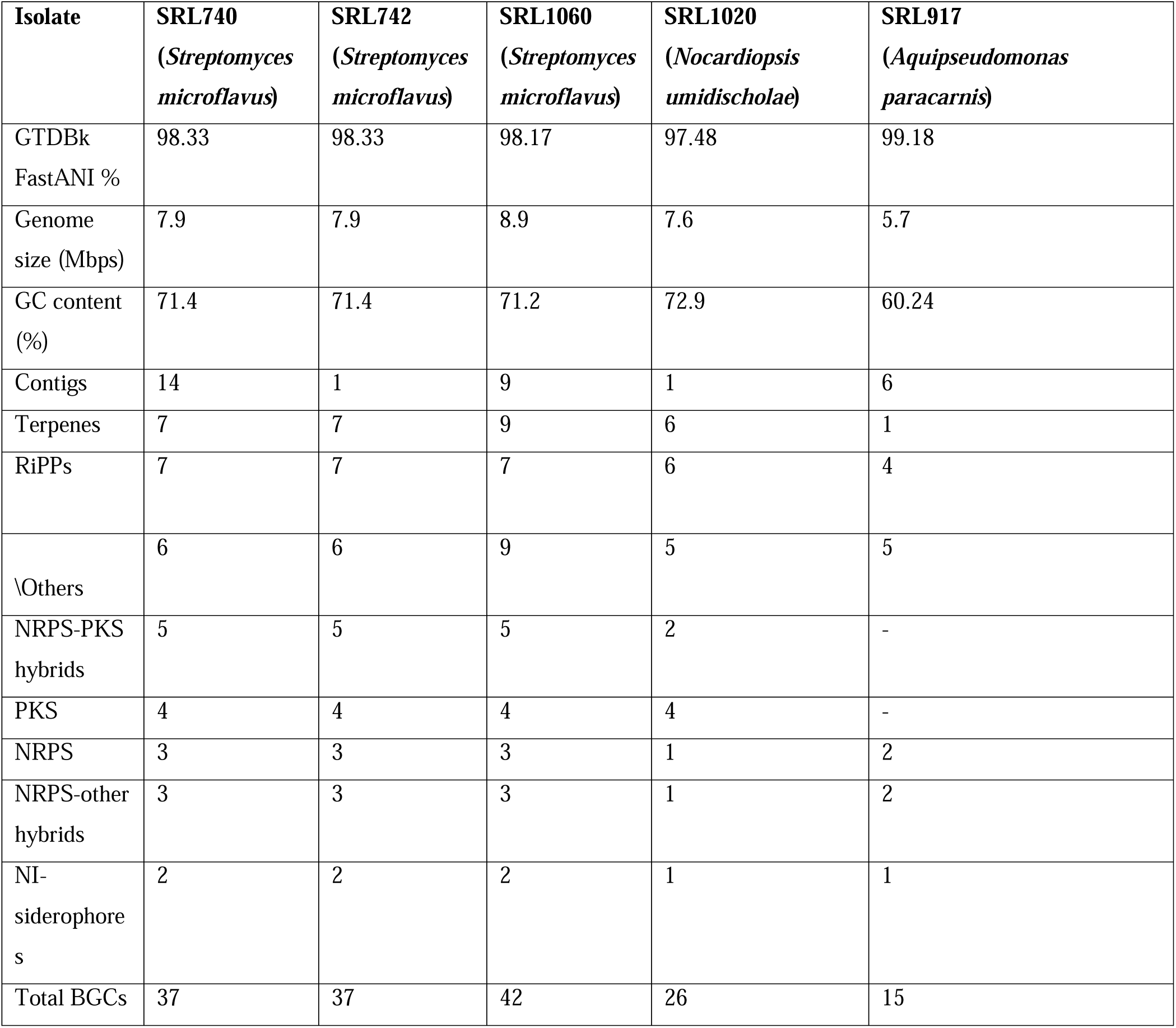
Summaries of the whole genome assemblies and their Biosynthetic Gene Cluster annotations.

Genome mining with antiSMASH (Blin et al., 2025) revealed several potentially “silent” or cryptic BGCs (Gosse et al., 2019; Lee et al., 2020; Hoskisson and Seipke, 2020) (Supplementary Table 4). Such BGCs could be exploited through modern synthetic biology approaches, including genetic refactoring, promoter engineering, and heterologous expression, to activate dormant pathways and harvest the structurally unique chemical scaffolds needed to combat global antimicrobial resistance (Lee et al., 2020). The abundance of antibiotic-associated BGCs in our isolates points to their potential use against pathogens. Future work involving metabolite isolation, purification, and biochemical characterization will be necessary to directly link this inhibitory directly activity to specific known or novel compounds.

Comparative analysis of the three newly sequenced cave-derived *Streptomyces* isolates provided insight into their genomic relatedness and differentiation. Most gene clusters were shared among all three isolates, representing 81.7% of the three-isolate pangenome (Figure 6A). SRL740 and SRL742 showed nearly identical gene cluster repertoires, with only one genome-specific gene cluster detected in each isolate (Figure 6A; Supplementary Figure 1); given this small number of unique clusters, the differences may reflect minor assembly, annotation, or gene-fragmentation effects rather than biological divergence. SRL740 and SRL742 also uniquely shared 269 gene clusters that were absent from SRL1060, indicating a distinct shared genomic fraction between these two isolates (Supplementary Figure 1). Consistent with their high genomic similarity, SRL740 and SRL742 were isolated from samples collected at the same cave depth, suggesting that they may represent closely related isolates. In contrast, SRL1060 contained 1,181 gene clusters absent from the other two isolates, indicating a more distinct genomic profile (Figure 6A; Supplementary Figure 1).

**Figure 6.**
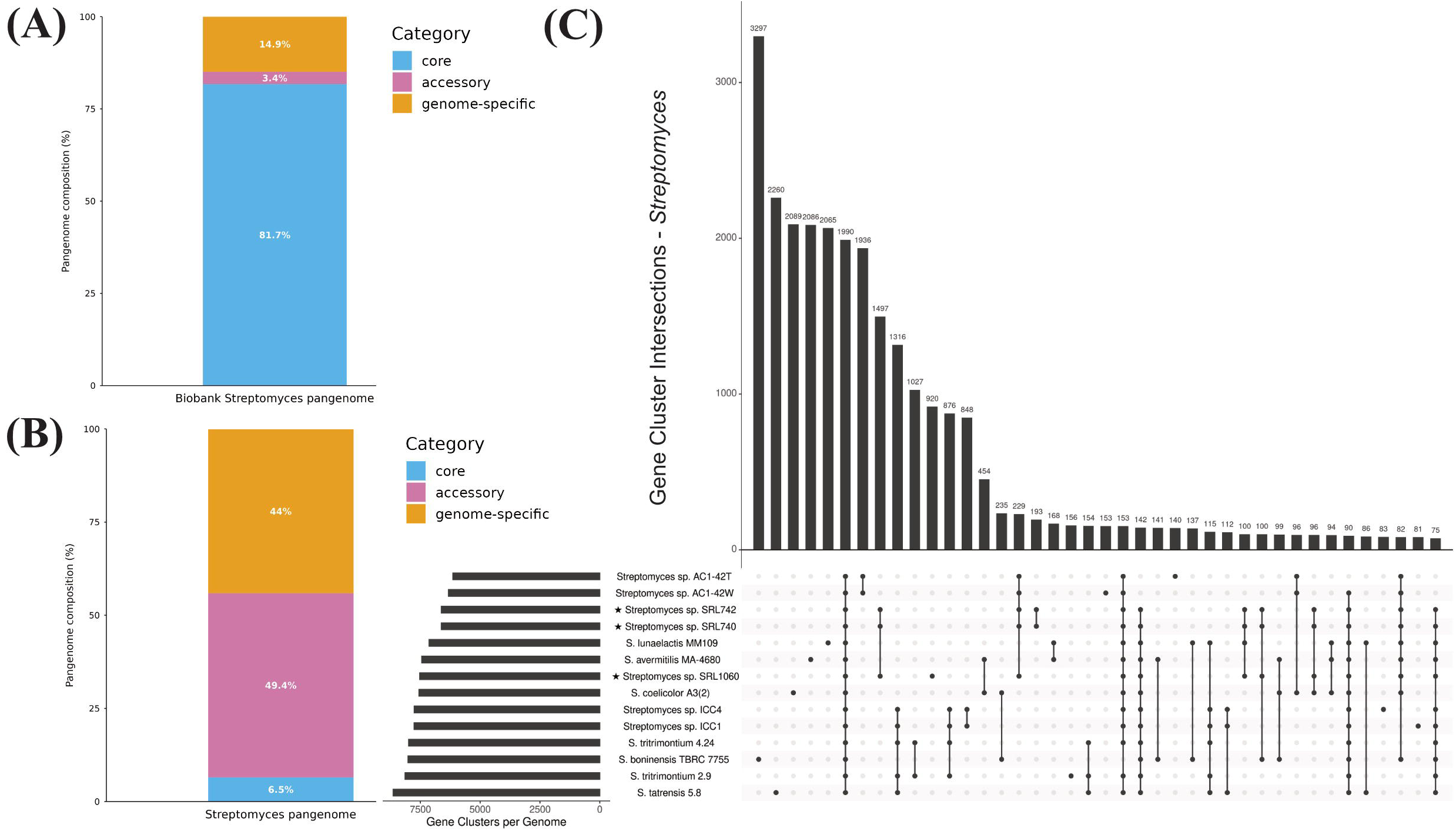
Pangenome analysis of *Streptomyces* genomes. **(A)** Distribution of core, accessory, and genome-specific genes (%) in the three-isolate pangenome of the newly sequenced cave-derived *Streptomyces* isolates. **(B)** Distribution of core, accessory, and genome-specific genes (%) in the pangenome analysis including the three cave-derived *Streptomyces* isolates and 11 publicly available *Streptomyces* genomes. **(C)** UpSet plot of the pangenome analysis of the three cave-derived *Streptomyces* isolates and 11 publicly available *Streptomyces* genomes. Horizontal bars (left) indicate the total number of gene clusters per genome, while vertical bars (top) indicate the number of gene clusters shared by each genome combination, as indicated by the connected dots below.

To place our three *Streptomyces* isolates in a broader context, we compared them with 11 publicly available *Streptomyces* genomes, emphasizing cave-derived or cave-associated strains. These included the Iron Curtain Cave isolates *Streptomyces* sp. ICC1 and ICC4 (Gosse et al., 2019), the moonmilk derived *Streptomyces lunaelactis* MM109^T^ (Naômé et al., 2018), the limestone cave derived *Streptomyces boninensis* TBRC 7755 (Yasawong et al., 2025), the moonmilk derived *Streptomyces tritrimontium* strains 2.9^T^ and 4.24 (Bielańska et al., 2025), the moonmilk derived *Streptomyces tatrensis* 5.8^T^ (Bielańska et al., 2026), and the guano/bat-associated *Streptomyces* sp. AC1-42T and AC1-42W (Leon et al., 2018). In addition, *Streptomyces avermitilis* MA-4680^T^(= NBRC 14893^T^) (Ikeda et al., 2003) and *Streptomyces coelicolor* A3(2) (Bentley et al., 2002) were included as a non-cave, commercial antibiotic production strain and a well-characterized model *Streptomyces* genome, respectively.

The pangenome analysis revealed extensive genomic diversity among the 14 *Streptomyces* genomes. Of a total of 30,423 gene clusters, only 1,990 (6.5%) belong to the core genome, while the remainder are either shared by subsets of genomes (accessory gene clusters, 49.4%) or are genome-specific (44%) (Figure 6B, Figure 6C). Several genomes display elevated numbers of genome-specific gene clusters, with *Streptomyces boninensis* TBRC 7755 showing the most (3,297), followed by *Streptomyces tatrensis* 5.8^T^ (2,260) (Figure 6C). Among the Gourgouthakas cave isolates, SRL1060 contains 920 genome-specific gene clusters, while 193 clusters are uniquely shared between SRL740 and SRL742 (Figure 6C). Notably, the three Gourgouthakas isolates SRL740, SRL742, and SRL1060 uniquely share 1,497 gene clusters that are absent from all the other cave-associated and non-cave reference genomes included in the analysis (Figure 6C). These Gourgouthakas-shared clusters may represent a distinctive genomic fraction and provide useful targets for future investigation of genetic features potentially associated with adaptation to the specific environmental conditions of Gourgouthakas Cave.

Functional annotation of the genes in Gourgouthakas-isolate-specific or shared gene clusters revealed a considerable fraction of poorly characterized genes. Specifically, genes lacking functional annotation account for 28.1% of SRL1060-specific genes, 18.8% of genes in clusters uniquely shared by SRL740 and SRL742, and 8.8% of genes in clusters uniquely shared by all three Gourgouthakas isolates (Supplementary Table 5). Among the annotated genes, recurrent functional categories include transcription, signal transduction, lipid and carbohydrate transport and metabolism, and secondary metabolite biosynthesis, transport, and catabolism (Supplementary Table 5). Several frequently detected domains, including tetratricopeptide repeats, methyltransferase domains, helix-turn-helix domains, and other regulatory or protein-interaction-associated domains, are also prominent. Tetratricopeptide repeat-containing proteins can contribute to the control of antibiotic biosynthesis and morphological differentiation in *Streptomyces* (Shi, et al., 2024), and methyltransferases are likewise important for antibiotic biosynthesis and activity in *Streptomyces* (Sokolova et al., 2023). Together, these features suggest that the unique genomic fraction of the Gourgouthakas isolates contains both poorly characterized genes and genes potentially associated with regulation, environmental adaptation, specialized metabolism and biosynthetic potential.

#### 3.3.2 *Nocardiopsis* sp. (SRL1020) genome

The genus *Nocardiopsis* sits at the intersection of ecological adaptability and industrial innovation (Bennur et al., 2015), and cave isolates of this genus have been reported to produce promising antimicrobial compounds (Bennur et al., 2015). While *Streptomyces* has been thoroughly sampled, *Nocardiopsis* species, mainly isolated from extreme marine habitats, deep-sea sediments, and hypersaline zones, represent an untapped reservoir of novel chemistry, with more than 67% of their natural products reported as entirely new chemical skeletons (Shi et al., 2022). Comparative genomic analyses of *Nocardiopsis* strains have revealed expansive pangenomes rich in unique gene content, encoding a diverse portfolio of polyketide synthases (PKS), non-ribosomal peptide synthetases (NRPS), and alkaloid biosynthetic pathways (Shi et al., 2022).

Isolate SRL1020 belongs to *Nocardiopsis umidischolae* and was recovered from clay found at a depth of 418 m (Figure 2A). The genome of *Nocardiopsis* SRL1020 contains 26 biosynthetic gene clusters (BGCs), of which 17 showed no similarity to known clusters and three showed low similarity confidence, indicating the presence of potentially underexplored secondary metabolites and antimicrobial compounds. Although SRL1020 harbours fewer BGCs than the *Streptomyces* isolates analyzed here, its clusters are likewise associated with terpenes, RiPPs, NRPS, and PKS, suggesting that this strain may also possess antimicrobial and antibiotic potential. Recent studies further highlight the value of *Nocardiopsis* and its metabolites for plant protection and biocontrol applications (Grigoryan et al., 2021).

### 3.4 *Aquipseudomonas paracarnis* (SRL917) genome

To gain insight into the genomic repertoire of SRL917, which showed inhibitory activity against all pathogens tested (Figure 3), a whole-genome hybrid sequencing was performed as described above for the *Streptomyces* and *Nocardiopsis* isolates. According to GTDB classification, the isolate belongs to *Aquipseudomonas paracarnis*. A secondary-metabolite analysis was then carried out to assess its genomic potential and identify any putative genes linked to antimicrobial capacity. A total of 15 BGC regions were detected (Table 1), comprising one terpene, four NRPS/NRPS-other hybrids, four RiPPs, one NI-siderophore and five other regions (specifically NAGGN, betalactone, arylpolyene, redox-cofactor and phosphonate). Only one NRPS-hybrid region showed high similarity to a known cluster, specifically the pseudomonine BGC from *Pseudomonas fluorescens*. Conversely, low similarity was observed for three other regions: the azole-containing-RiPP showed low similarity to the asplenin BGC from *Pseudomonas fuscovaginae* UPB0736; the arylpolyene region matched the APE Vf BGC from *Aliivibrio fischeri* ES114; and one NRPS region matched ambactin from *Xenorhabdus miraniensis* (Table 1; Supplementary Table 4). Ambactin has already been shown to act as an antibacterial agent and has been successfully cloned by the ExRec method (Schimming et al., 2014). Conversely, asplenin is a cyclic lipopeptide shown to be responsible for the bacterial motility in strain UPB0736, without activity against *Rhizoctonia solani* (Ferrarini et al., 2022). Nonetheless, because the SRL917 BGC showed only low similarity to the asplenin, a putative antimicrobial role cannot be excluded without further testing. Overall, the novel BGCs found in the SRL917 isolate may act as antipathogenic molecules.

### 3.5 *Botrytis cinerea ex vivo* inhibition

*Botrytis cinerea* is a globally distributed phytopathogenic fungus responsible for gray mould, an economically important disease affecting numerous crops, particularly those grown under greenhouse conditions. As a necrotrophic pathogen, it can infect various plant organs and typically initiates infection through wounds created during harvesting or handling (Elad et al. 2007). In many cases, infections remain latent and symptoms emerge only at the post-harvest stage, significantly reducing the market quality and shelf life of agricultural products (Hu et al., 2016).

On the basis of its strong *in vitro* performance against multiple phytopathogens and the putative antimicrobial agents that it may release, *Aquipseudomonas paracarnis* SRL917 was selected for further *ex vivo* experiments. Application of the bacteria to tomato leaves (Figure 7D) led to a marked reduction in disease symptoms caused by *Botrytis cinerea*, compared with both the positive control (Figure 7B) and the commercial product X (Figure 7C; Supplementary Table 6). Overall, the *in vitro* and *ex vivo* results demonstrate the antimicrobial potential of the *Aquipseudomonas paracarnis* isolate and further investigation regarding its secondary metabolites will be of crucial importance.

**Figure 7.**
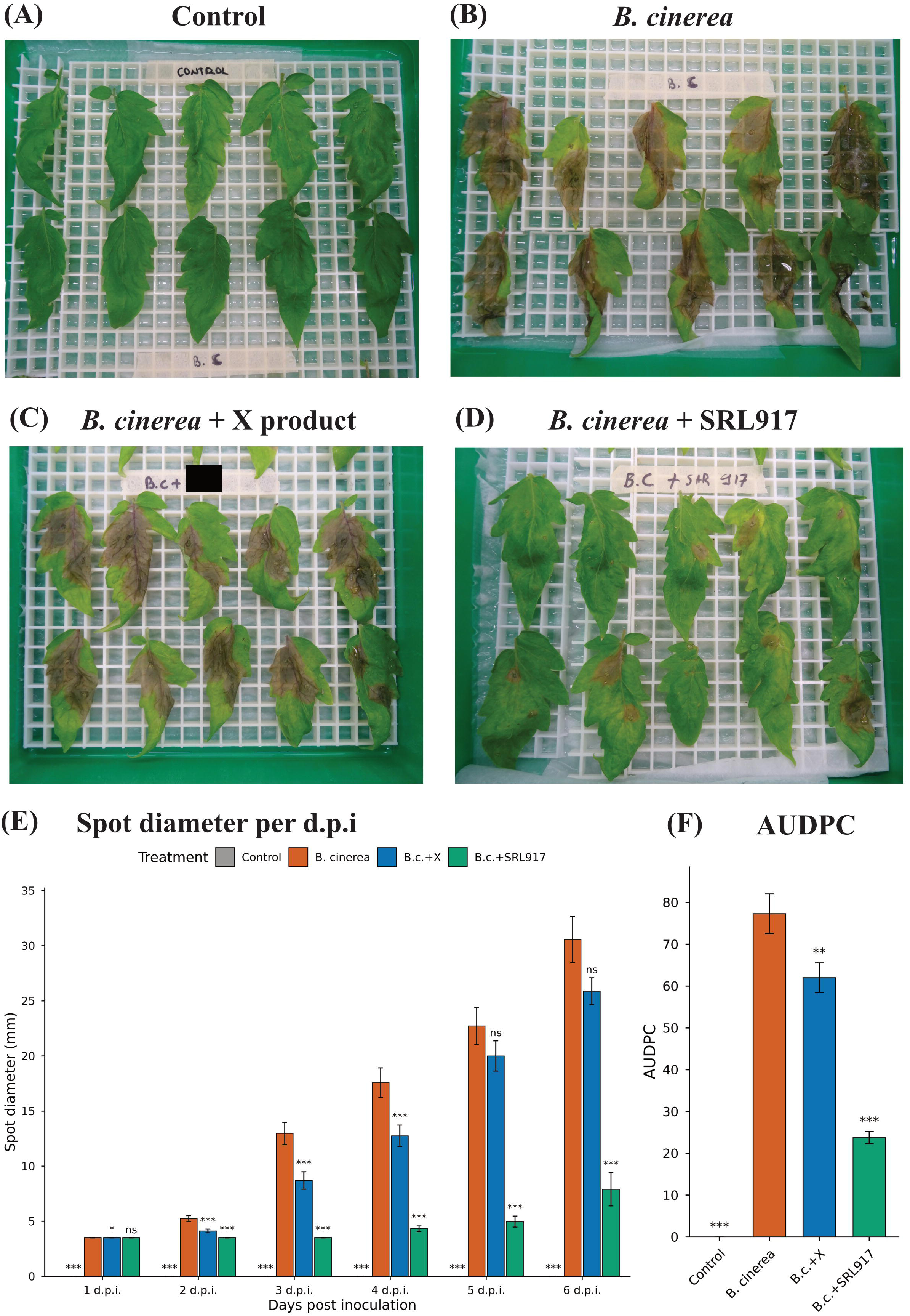
*In planta* assay in tomato leaves (Belladona F1) examining the disease severity of *B. cinerea* and the inhibition of a commercial product (X) and bacterial isolate SRL917. Photos A-D are on the 6^th^ DPI. **(A)** Healthy control leaves. **(B)** Diseased leaves without any treatment. **(C)** Diseased leaves treated with a commercial product (X). **(D)** Diseased leaves treated with inoculation of SRL917. **(E)** Bar plot of the spot diameter of leaves in the assay with error bars across the DPI. **(F)** Area Under the Disease Progress Curve of the assay with error bars and statistical significance. Stars correspond to statistically significant difference between groups, i.e. *p-value* ***=0.001, **=0.01, *=0.05 and non-significant=ns, as derived from ANOVA followed by post-hoc Tukey tests. Error bars represent the standard error of the mean.

## 4 Conclusion

The importance of caves as reservoirs of unique microbial life and functionality, has long been recognized (Engel, 2015; Ghosh et al., 2017). The terrestrial deep subsurface remains comparatively unexplored (Beaver and Neufeld, 2024), and deep caves – being humanly accessible – offer a valuable window onto it.

Cave microbes have provided templates for various drugs that help manage both human and plant health, and the discovery of novel microbes in extreme environments such as deep caves can benefit public health, climate resilience, and sustainable agriculture (Liu et al., 2025). To date, we have likely exploited only a fraction of the chemical space of microbial natural products: compared with the molecules catalogued in NPAtlas (Poynton et al., 2024), only around 3-4% of biosynthetic diversity has been experimentally accessed (Gavriilidou et al., 2022). Actinobacteria, such as *Streptomyces* and *Nocardiopsis* were until recently considered “exhausted” as a source of new drugs; the unique biosynthetic pathways in the genomes of the strains we studied suggest otherwise. Indeed, the recent finding of Kaur et al. (2026) shows that new antibiotics may still be discovered from genera such as *Streptomyces,* despite decades of intensive study.

Greece’s remarkable and unique cave systems remain poorly studied in this respect. Our microbial collection from the deep cave of Gourgouthakas showcases promising biotechnological applications across agrifood and medical research, and is, to our knowledge, the first large microbial collection from a single oligotrophic karstic system in Greece. Future metagenomic sequencing of these samples will aim to reveal the full microbial diversity of the cave.

## 5 Data Availability Statement

The 16s rRNA and whole genome sequences of the microbial collection of Gourgouthakas are deposited in ENA under the accession number PRJEB109961. The metadata of the samples of the ENA project are provided at Supplementary Table 7. All code generated for this work is also available in this repository https://github.com/savvas-paragkamian/gourgouthakas_isolates.git

## 6 Supplementary Material

All supplementary material is deposited on Zenodo under the identifier: https://zenodo.org/records/20694818

More specifically the contents of the repository are:

● Supplementary Presentation 1: Sampling photos of sampling in Gourgouthakas cave
● Supplementary Presentation 2: Photos of inhibitions *in vitro* of bacterial phytopathogens
● Supplementary Presentation 3: Photos of inhibitions *in vitro* of fungal phytopathogens
● Supplementary Table 1: Growth information of stabs of the isolates of SRL Gourgouthakas cave biobank
● Supplementary Table 2: Taxonomic information of 16s rRNA of the SRL biobank isolates
● Supplementary Table 3: Measurements of the phytopathogen *in vitro* inhibitions
● Supplementary Table 4: Data on the Biosynthetic Gene Clusters of the assembled genomes
● Supplementary Table 5: Data on the comparative genomics and Pangenomes of *Streptomyces* genomes
● Supplementary Table 6: Data of the *ex vivo* assay of SRL917 against *Botrytis cinerea*
● Supplementary Table 7: GSC MixS (ERC000021) sample metadata submitted to project PRJEB109961
● Supplementary Figure 1: UpSet plot of the pangenome analysis of the three newly sequenced cave-derived *Streptomyces* spp. isolates

## 7 Conflict of Interest

*The authors declare that the research was conducted in the absence of any commercial or financial relationships that could be construed as a potential conflict of interest*.

## 8 Author Contributions

Conceptualization: PFS, SP, CAC

Data curation: SP, CAC, VAM, NPA

Formal analysis: SP, CAC, VAM, NPA

Funding acquisition: PFS, MV

Investigation: CAC, VAM, MP, SS

Methodology: CAC, VAM

Project administration: PFS, CAC, SP

Resources: SP, PFS, EAM

Software: SP, CAC, MP, NPA

Supervision: PFS, EAM, CAC, MV, YS

Validation: PFS, CAC, VAM, EAM, CP, MV

Visualization: SP, NPA, VAM, YS, SS, CAC, MP, MV

Writing – original draft: SP, VAM, SS, NPA, MP, YS, CP, PFS

Writing – review & editing: All authors reviewed and accepted the manuscript

## 9 Funding

This research received NO funding.

## Supporting information

Supplementary Presentation 1

Supplementary Presentation 2

Supplementary Presentation 3

## 10 Acknowledgments

The authors would like to thank all the members of the expedition “Gourgouthakas 2022” that made this sampling possible (in alphabetical order): Adamopoulos Kostas, Alexandrou Stratis, Aggelopoulos Giorgos, Antonopoulos Dimitris, Argyris Giorgos, Benezes Makis, Bourdas Dimitris, Chalampardaki Katerina – Ahaka, Charitakis Giannis, Chatziapostolou Makis, Digenis Markos, Drakopoulou Katerina, Georgopoulou Xenia, Kadas Pavlos-Apostolos, Kaplantzis Takis, Kelaidi Maria, Kostidis Kostis, Kouzmina Evgenia, Ladakis Petros, Lambrinos Stelios, Loukas Charisis, Manolas Argyris, Margiolis Alexandros, Maroulis Stavros, Mitsakis Nikos, Nikolakaki Sofia, Papaeliou Chara, Pantelios Vasilis, Papoulias Christos, Partsios Ilias, Smarianaki Katerina, Stamoulou Maria, Tsichlakis Nikiforos, Tsopelas Michalis, Vasilopoulou Katerina, Vlachos Giorgos The authors thank also Prof. Dimitris Goumas for his kind donation of *Paracidovorax citrulli*, Dr. Emilia Markellou for her kind donation of *Phytopthora nicotianae*, Dr Ilias Lagkouvardos and Ms Maria Malliarou for their kind contribution with the extraction and sequencing of the SRL917 isolate and the reviewers for their valuable input and suggestions which improved our manuscript.

## Notes

https://zenodo.org/records/20694818

### Competing Interest Statement

The authors have declared no competing interest.

### Summary of Updates

We now have extensively revised our manuscript. In this version we decided to include one more microbial genome. We have now added a revised pipeline of the taxonomy identification of full length 16S rDNA, as was suggested by our respected reviewers.

