## Supplementary Presentation 1 for "A Thousand Meters Deep: Vertical Profiling of the Subterranean Microbes of Gourgouthakas Cave"

### gourgouthakas_-0000_entrance_1

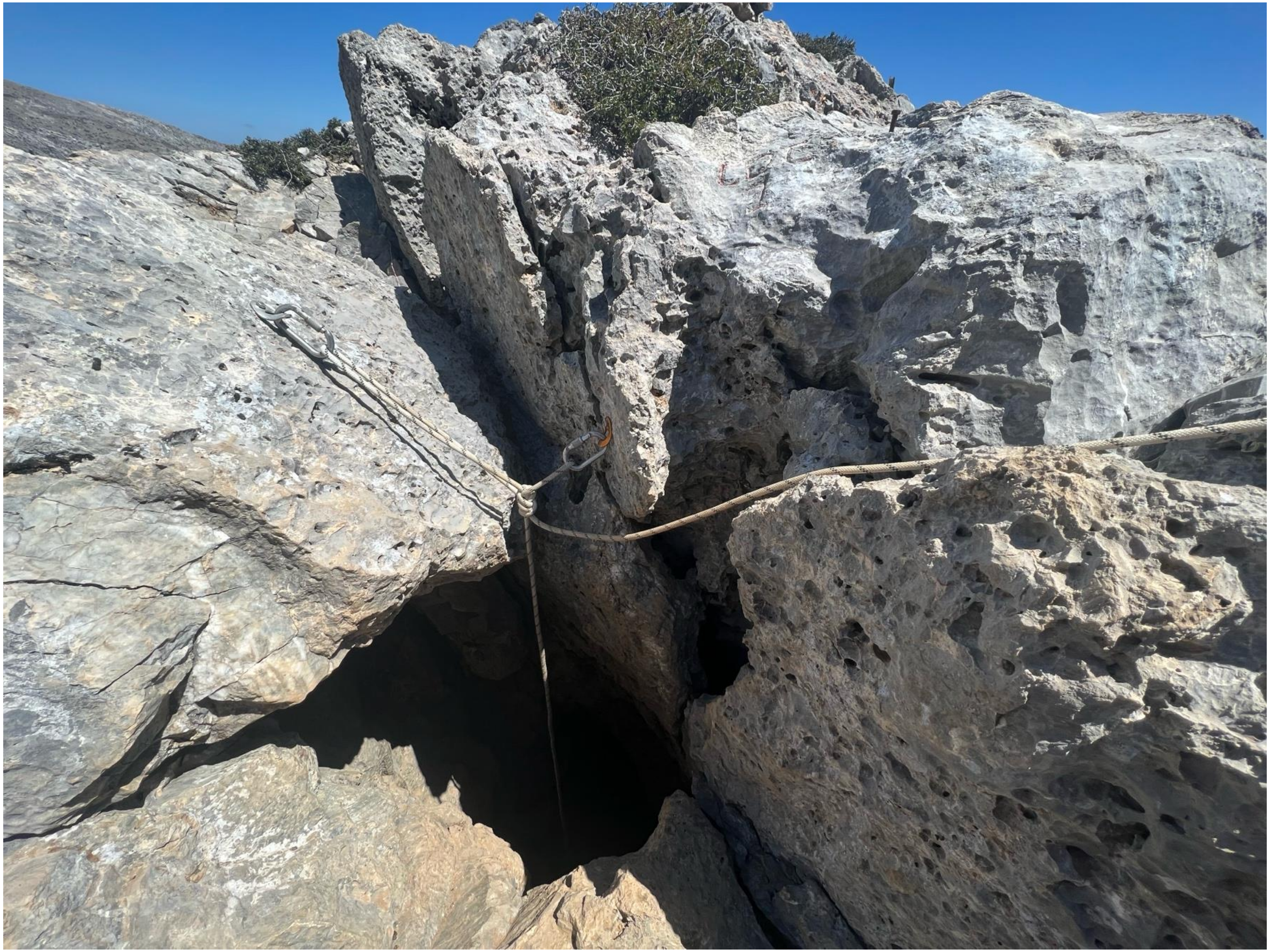

### gourgouthakas_-0000_entrance_2

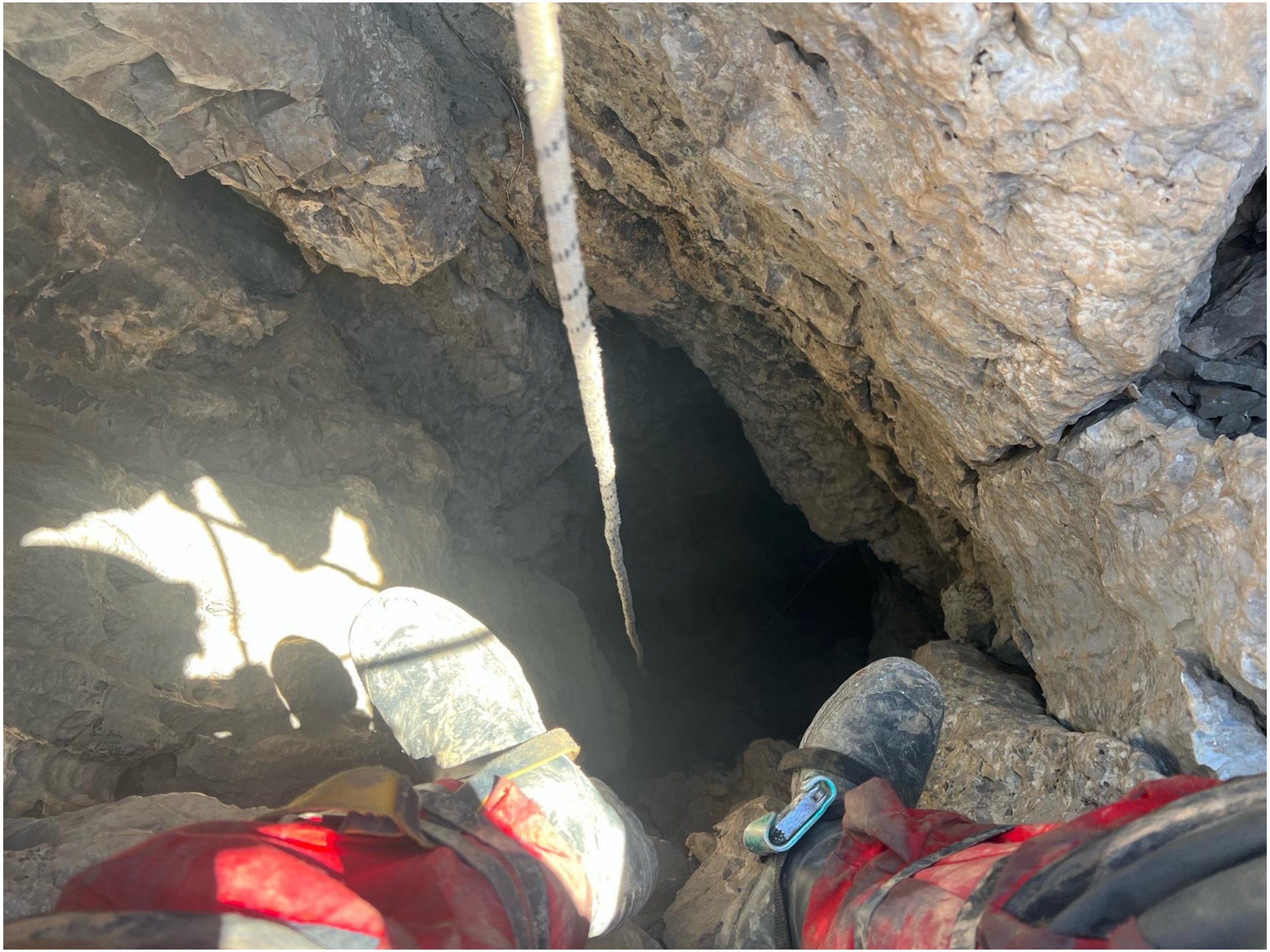

### gourgouthakas_-0000_entrance_3

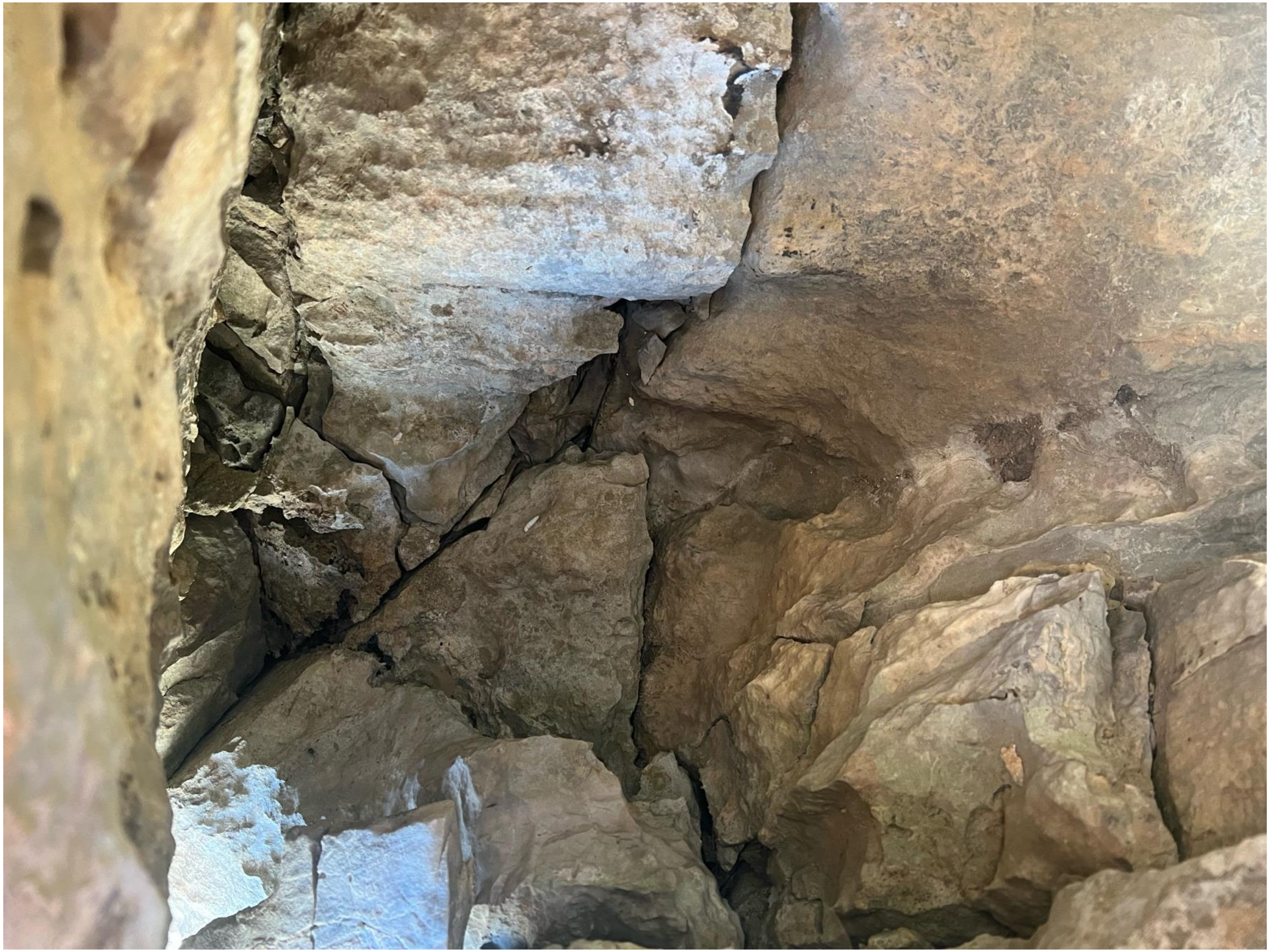

### gourgouthakas_-0000_entrance_4

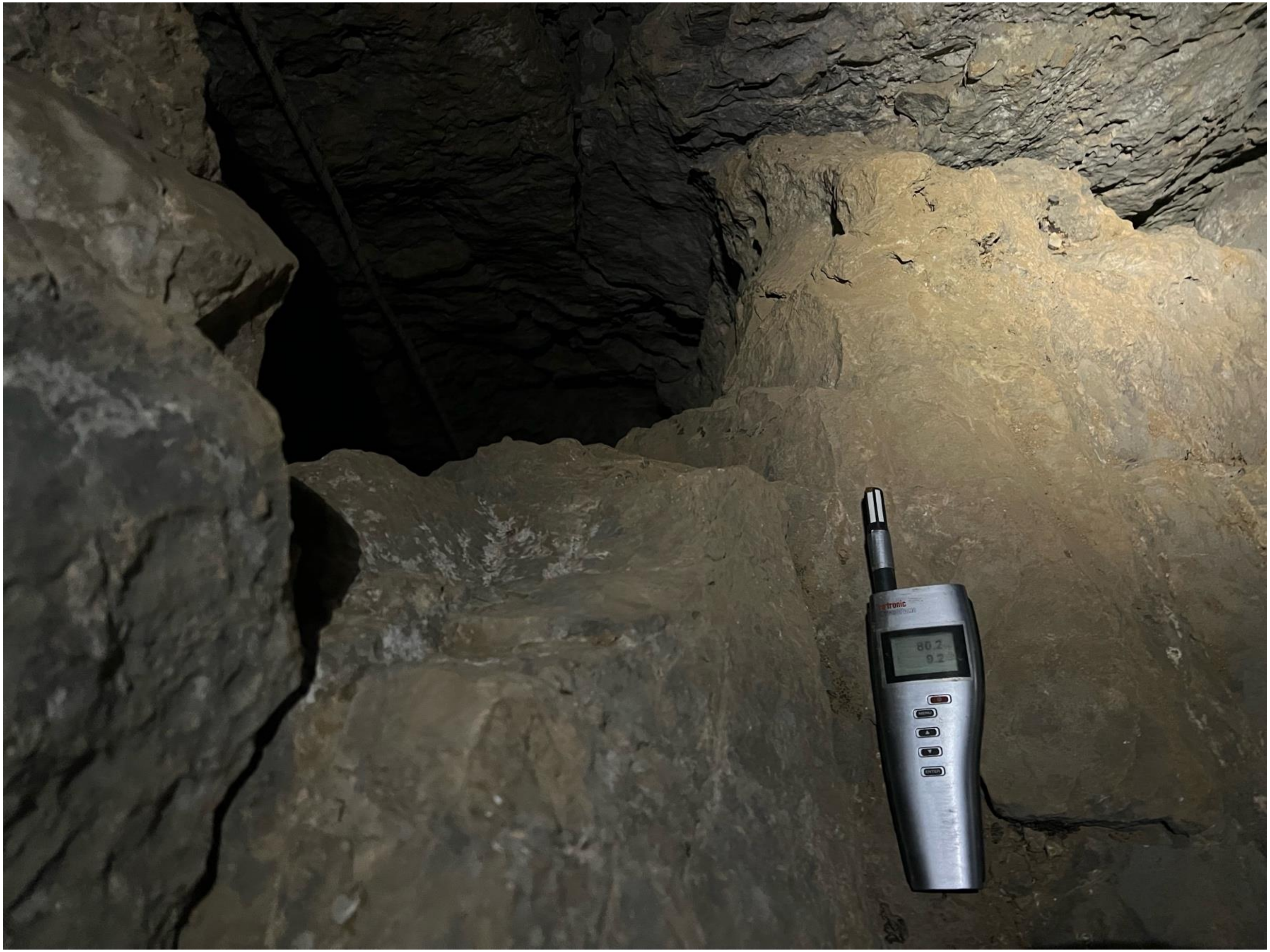

### gourgouthakas_-0000_entrance_5

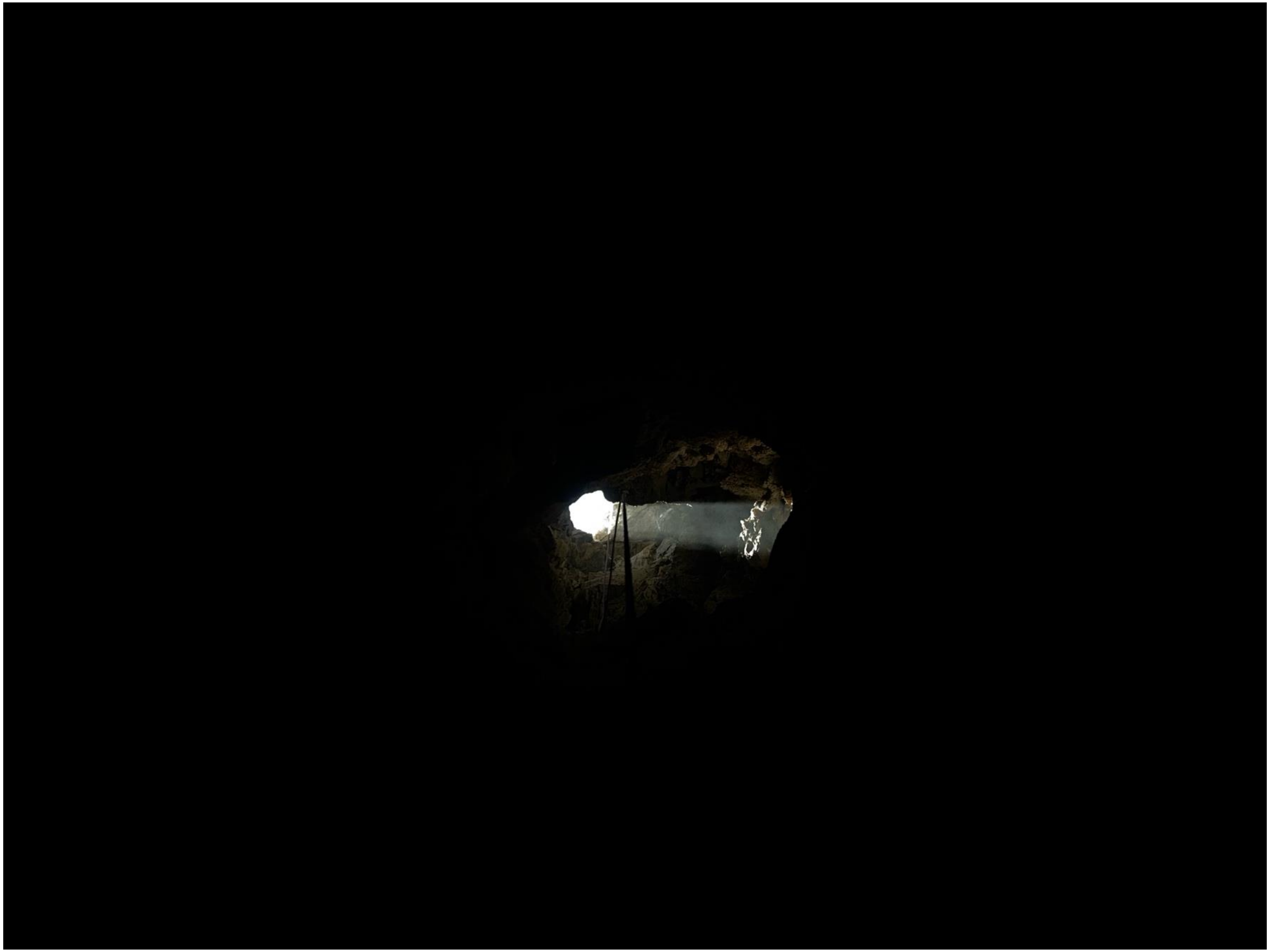

gourgouthakas_-0039_2

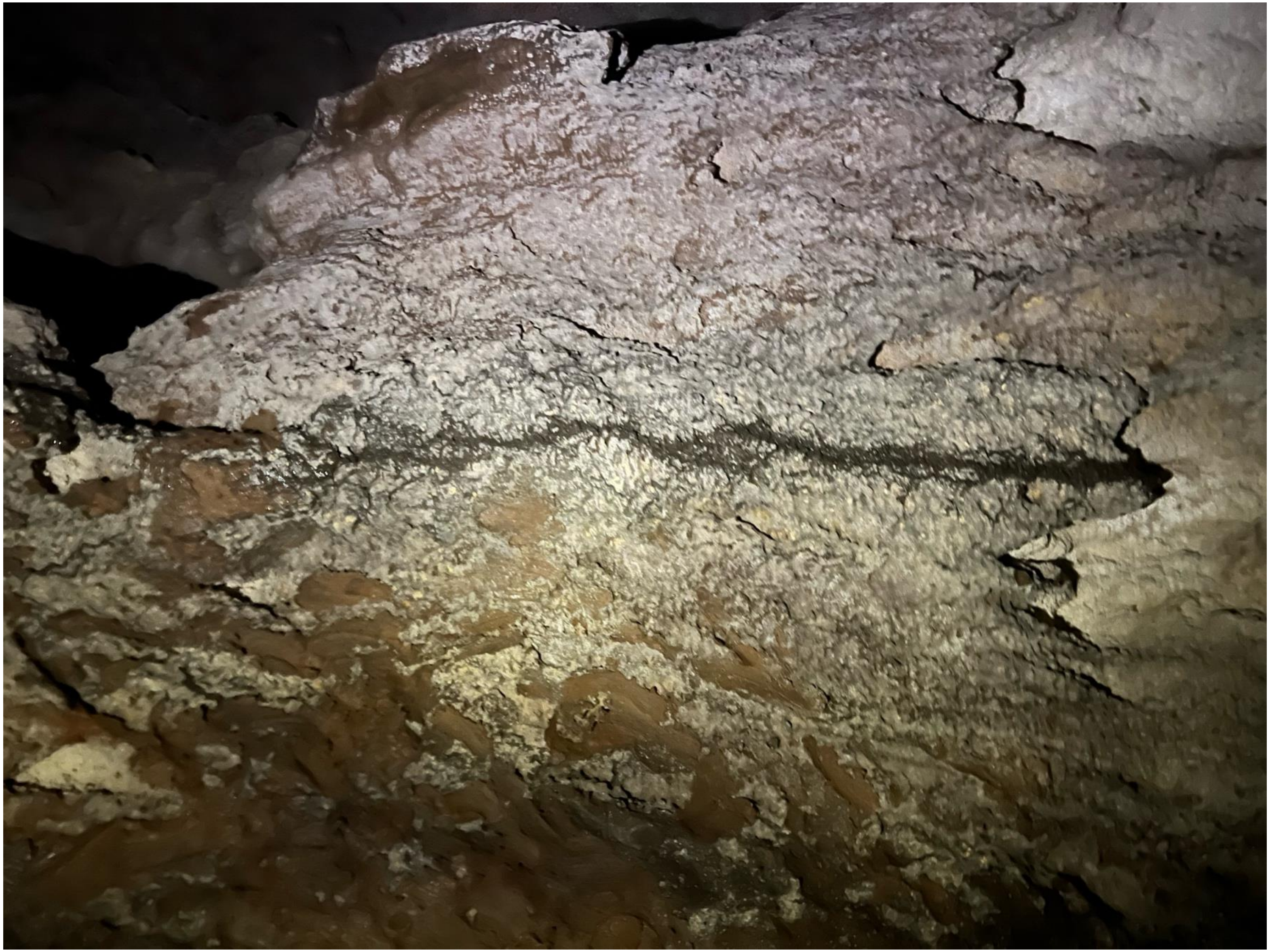

### gourgouthakas_-0039_3

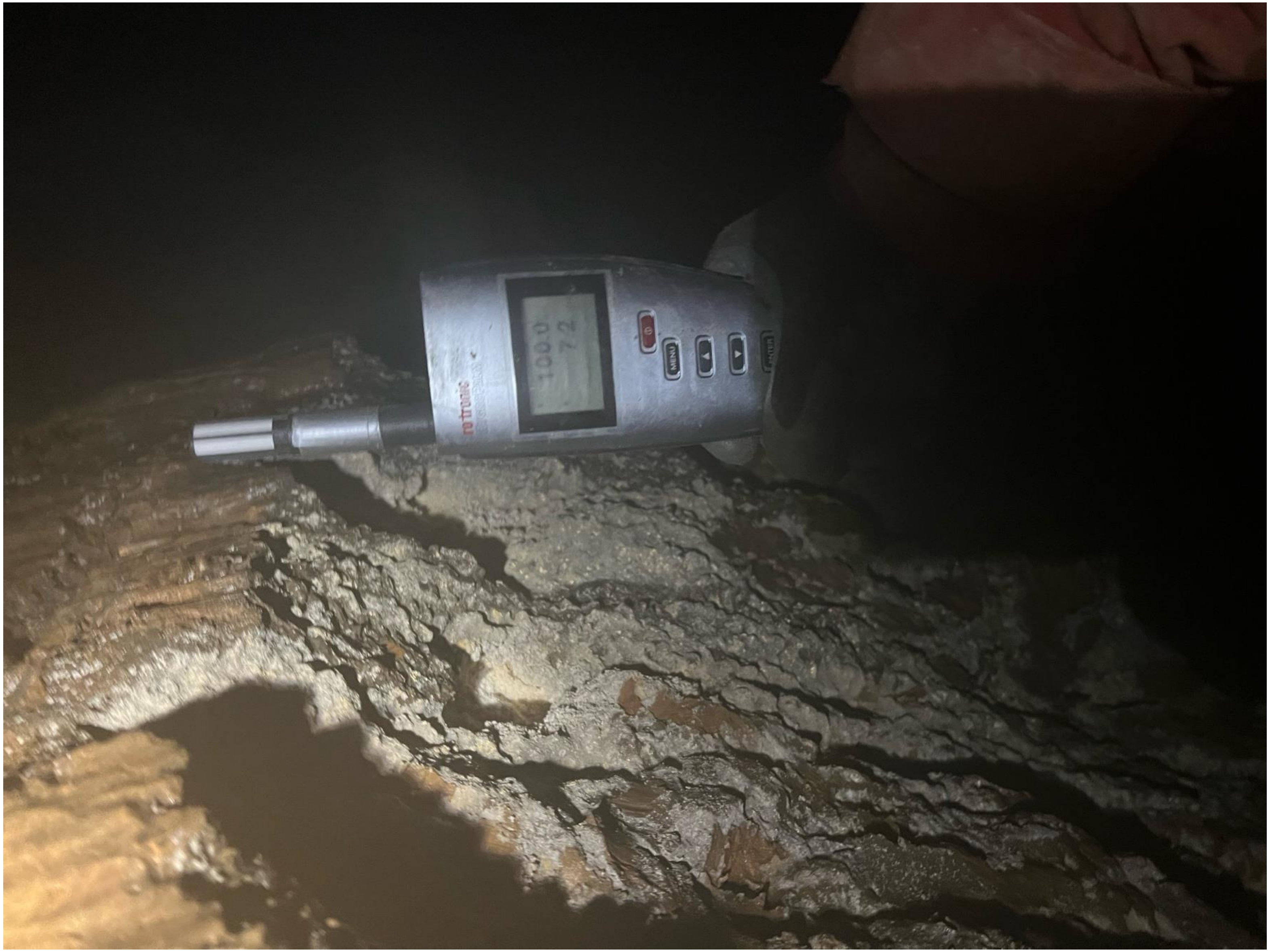

### gourgouthakas_-0039_4

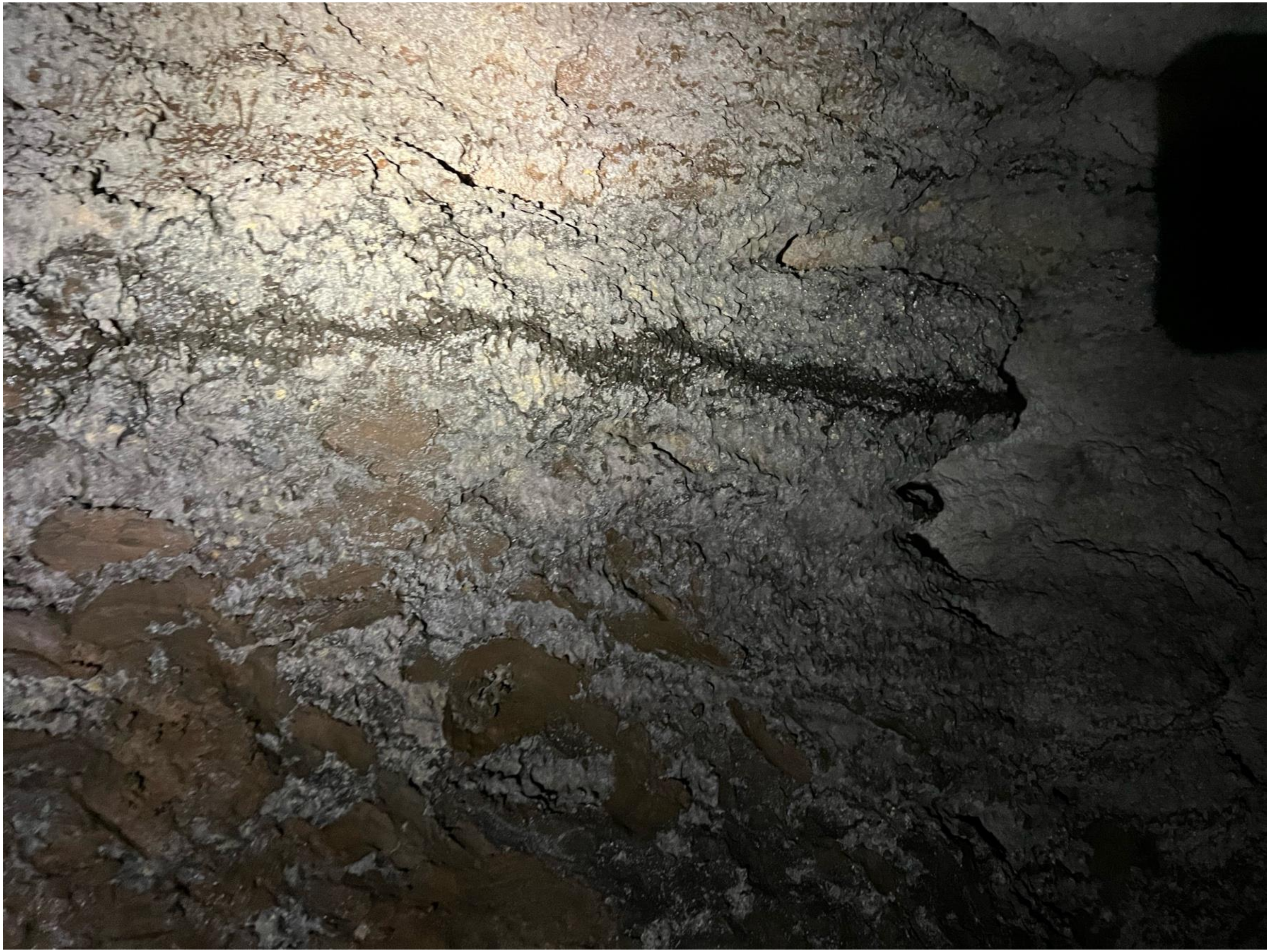

### gourgouthakas_-0220_meander_1

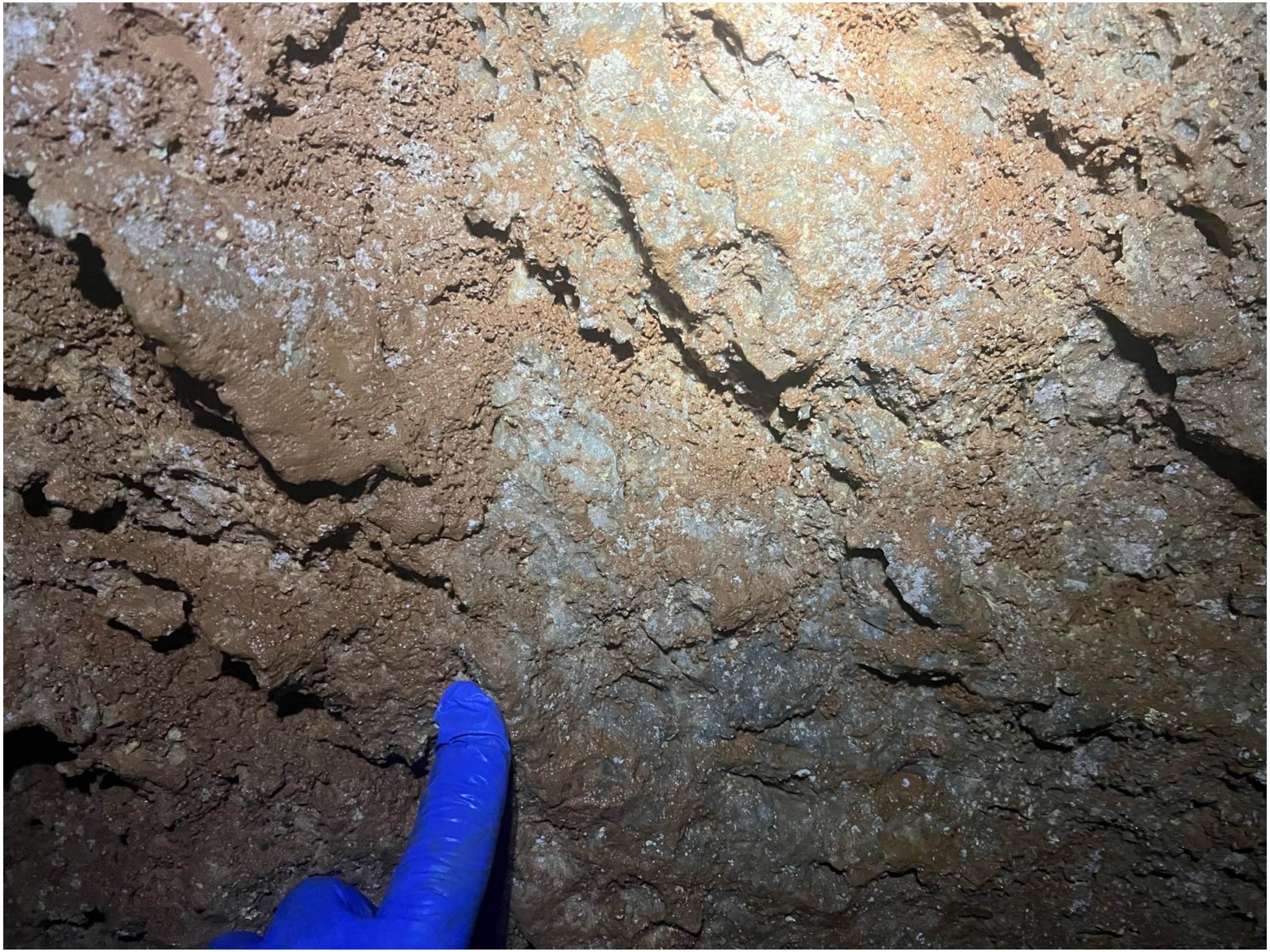

### gourgouthakas_-0220_meander_2

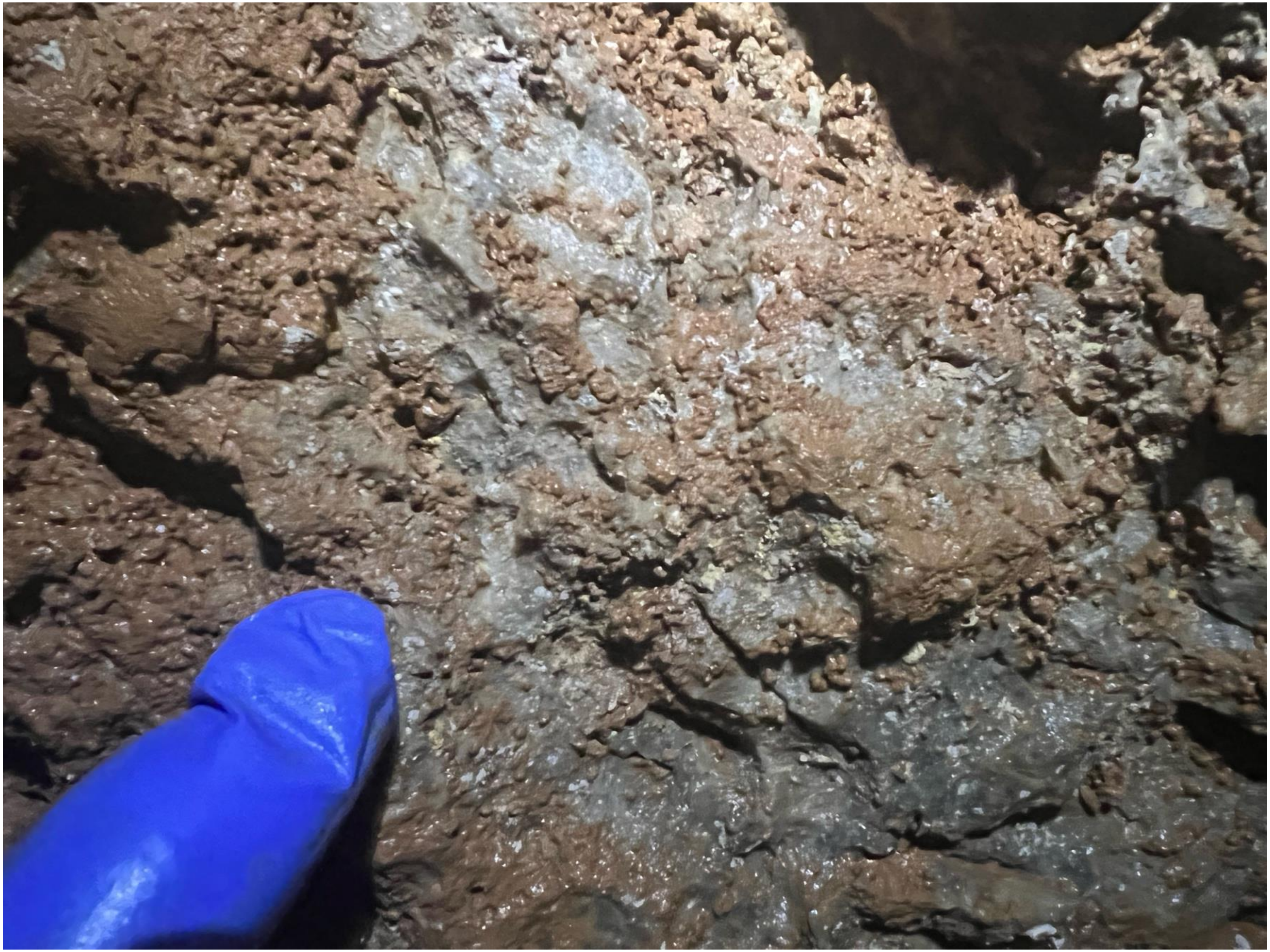

### gourgouthakas_-0220_meander_3

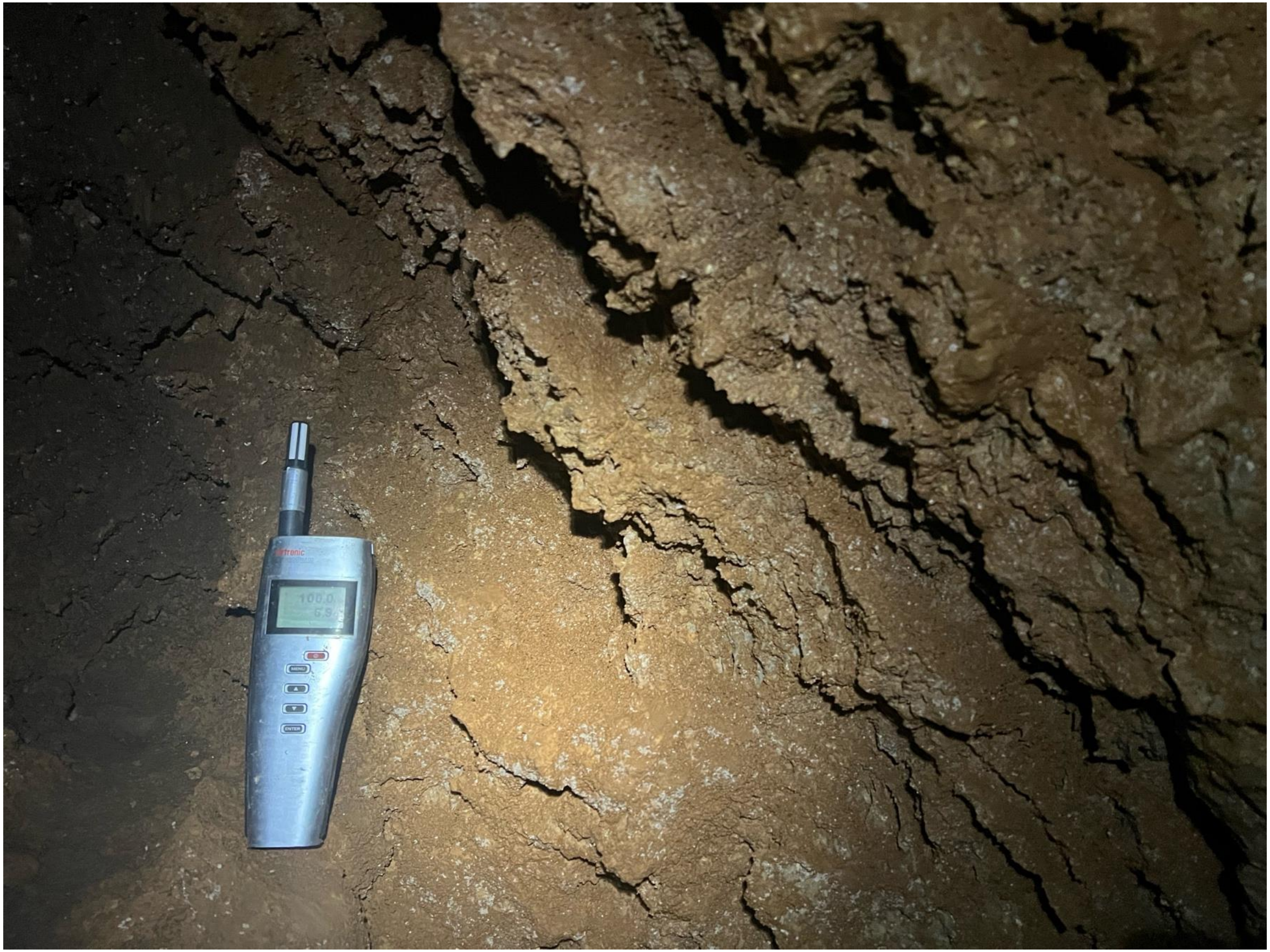

### gourgouthakas_-0220_meander_4

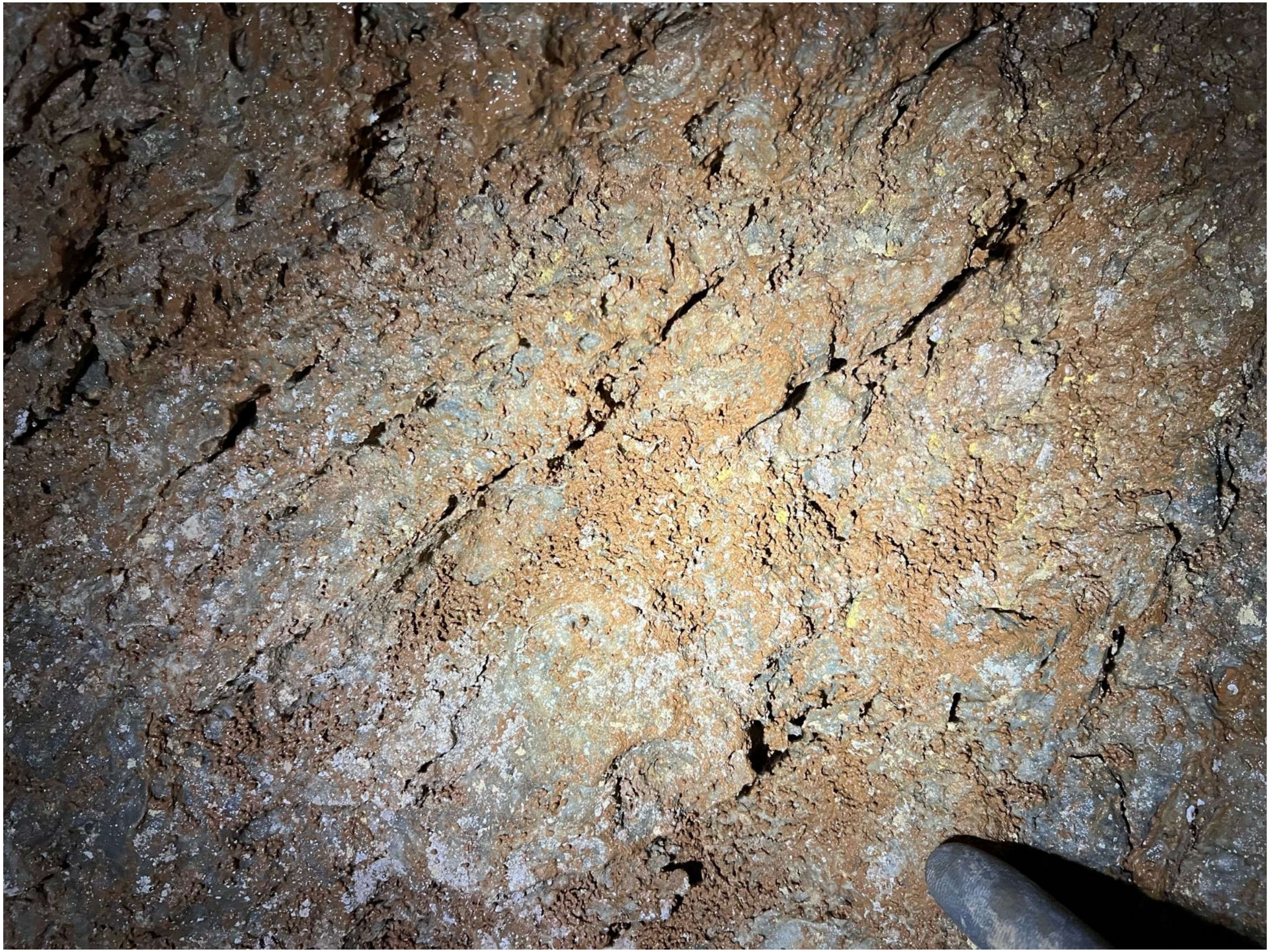

### gourgouthakas_-0418_leon_1

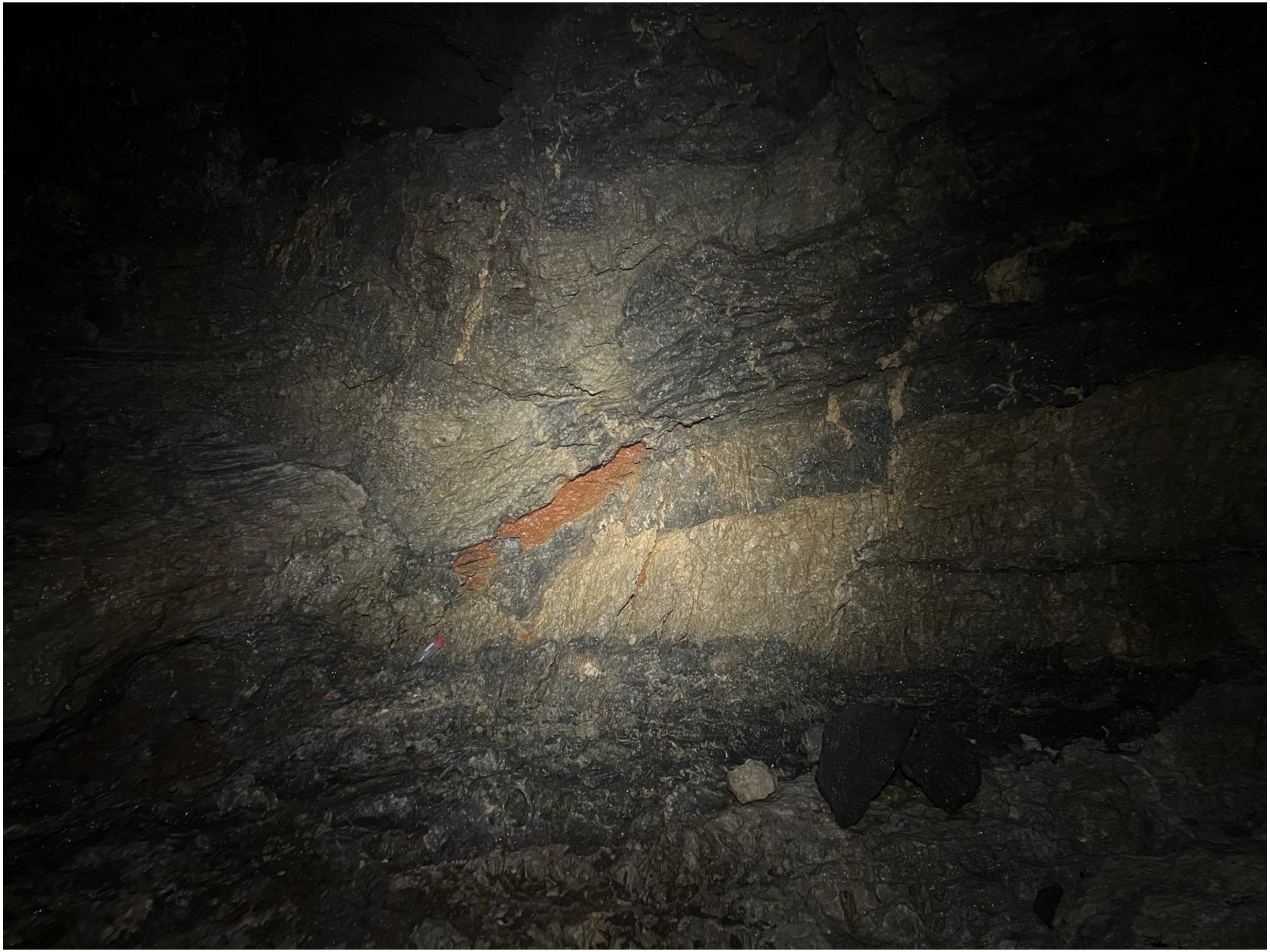

### gourgouthakas_-0418_leon_2

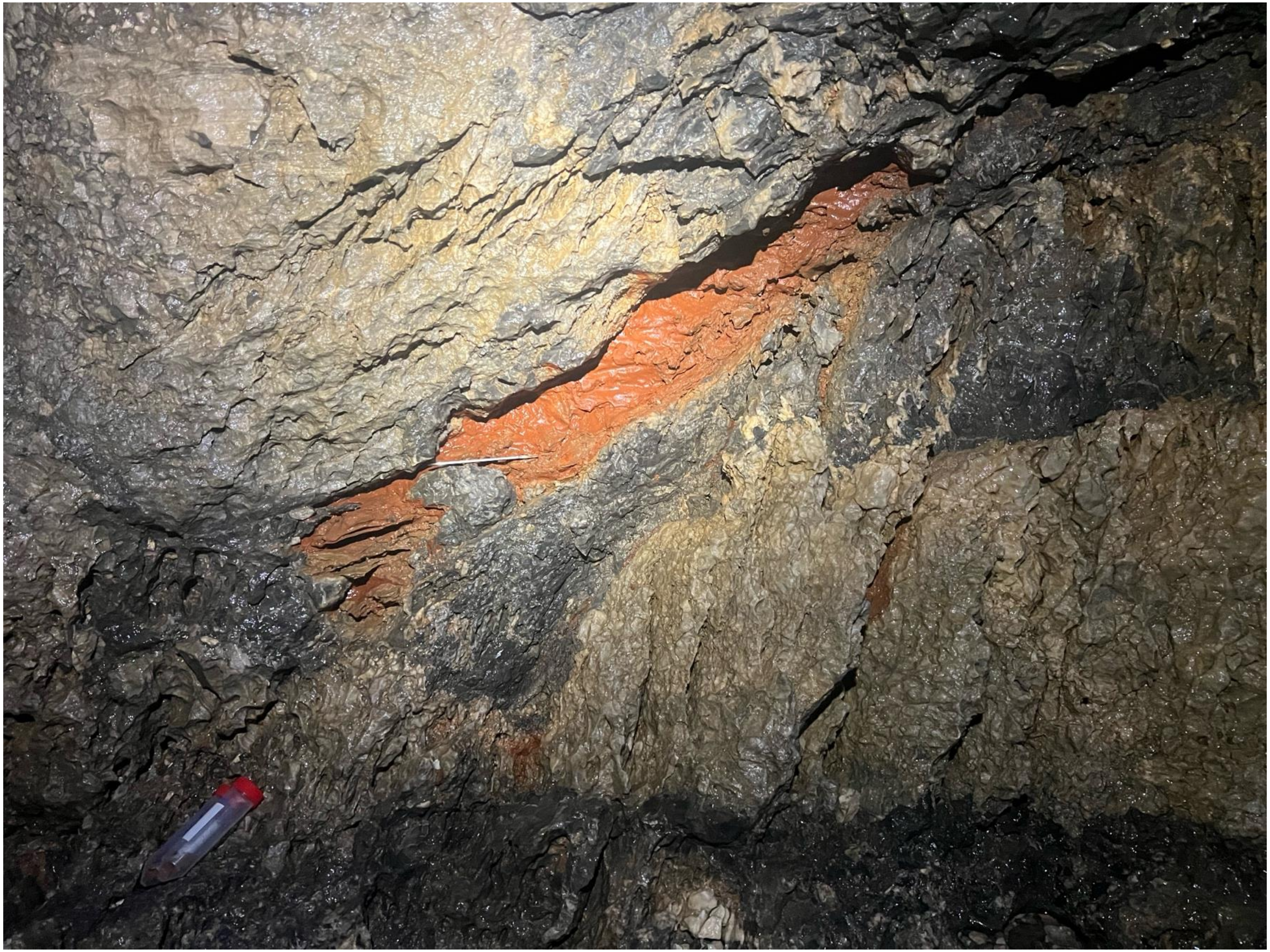

### gourgouthakas_-0418_leon_3

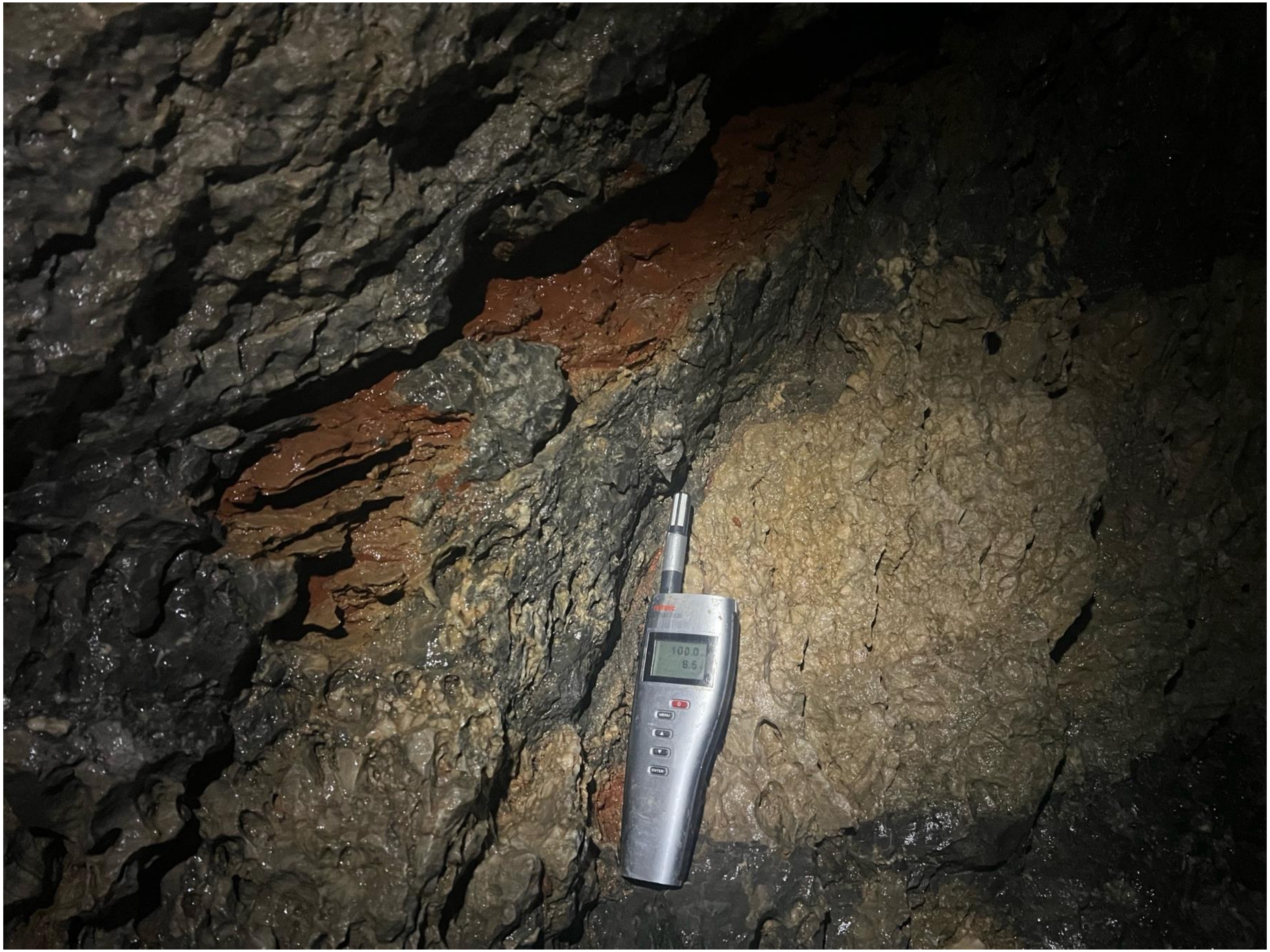

### gourgouthakas_-0418_leon_4

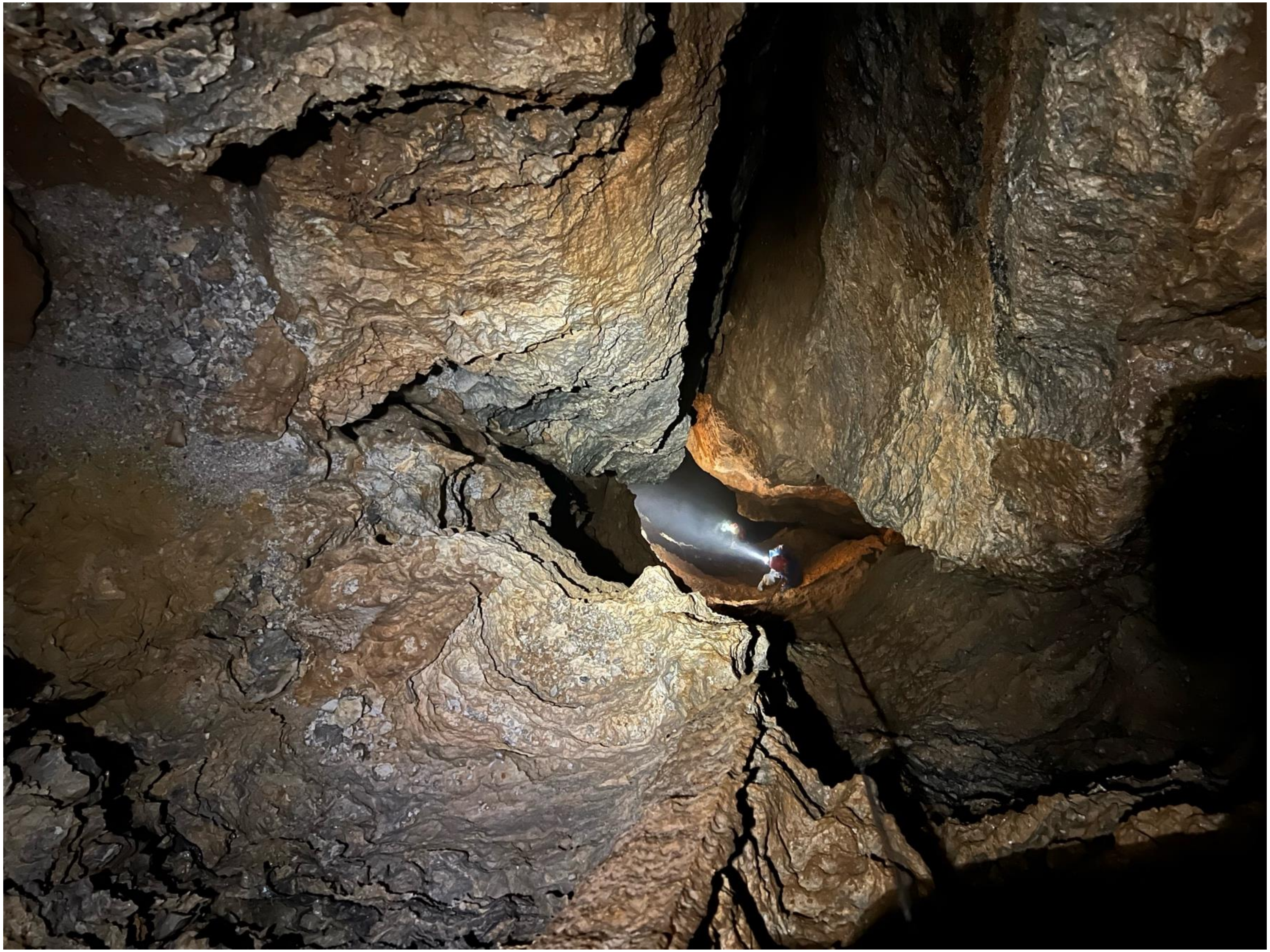

### gourgouthakas_-0433_gour_1

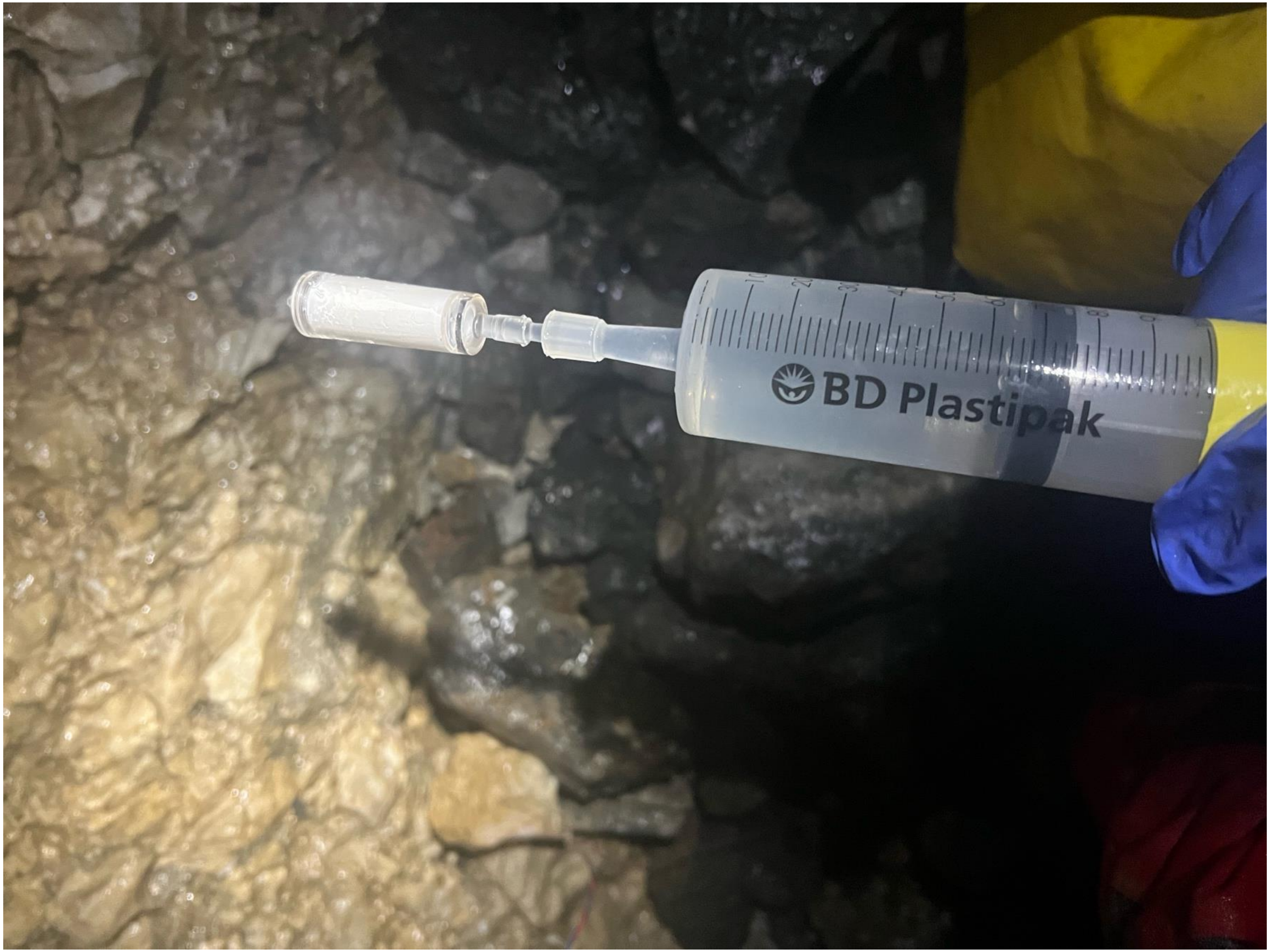

### gourgouthakas_-0433_gour_2

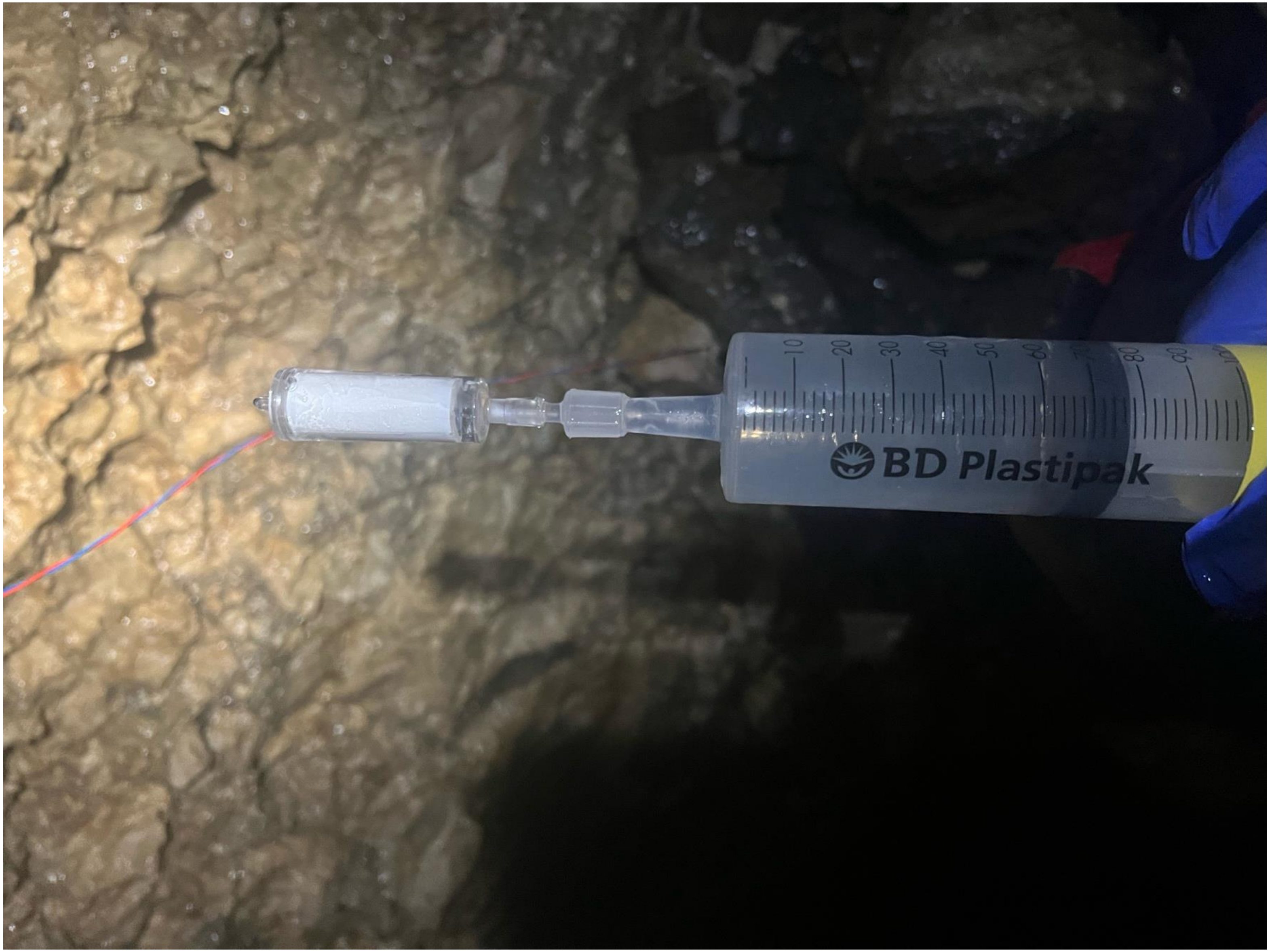

### gourgouthakas_-0433_gour_3

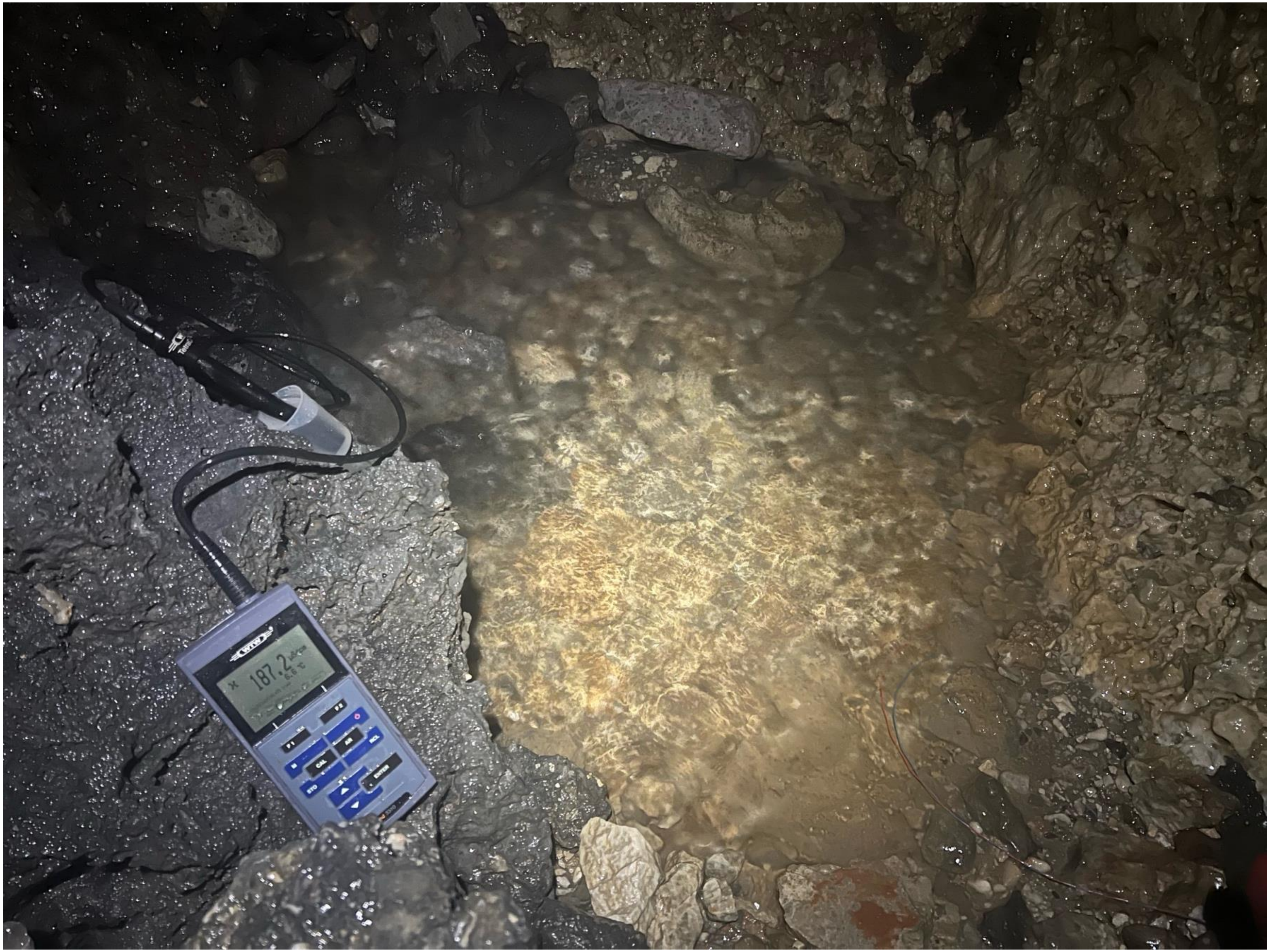

### gourgouthakas_-0433_gour_4

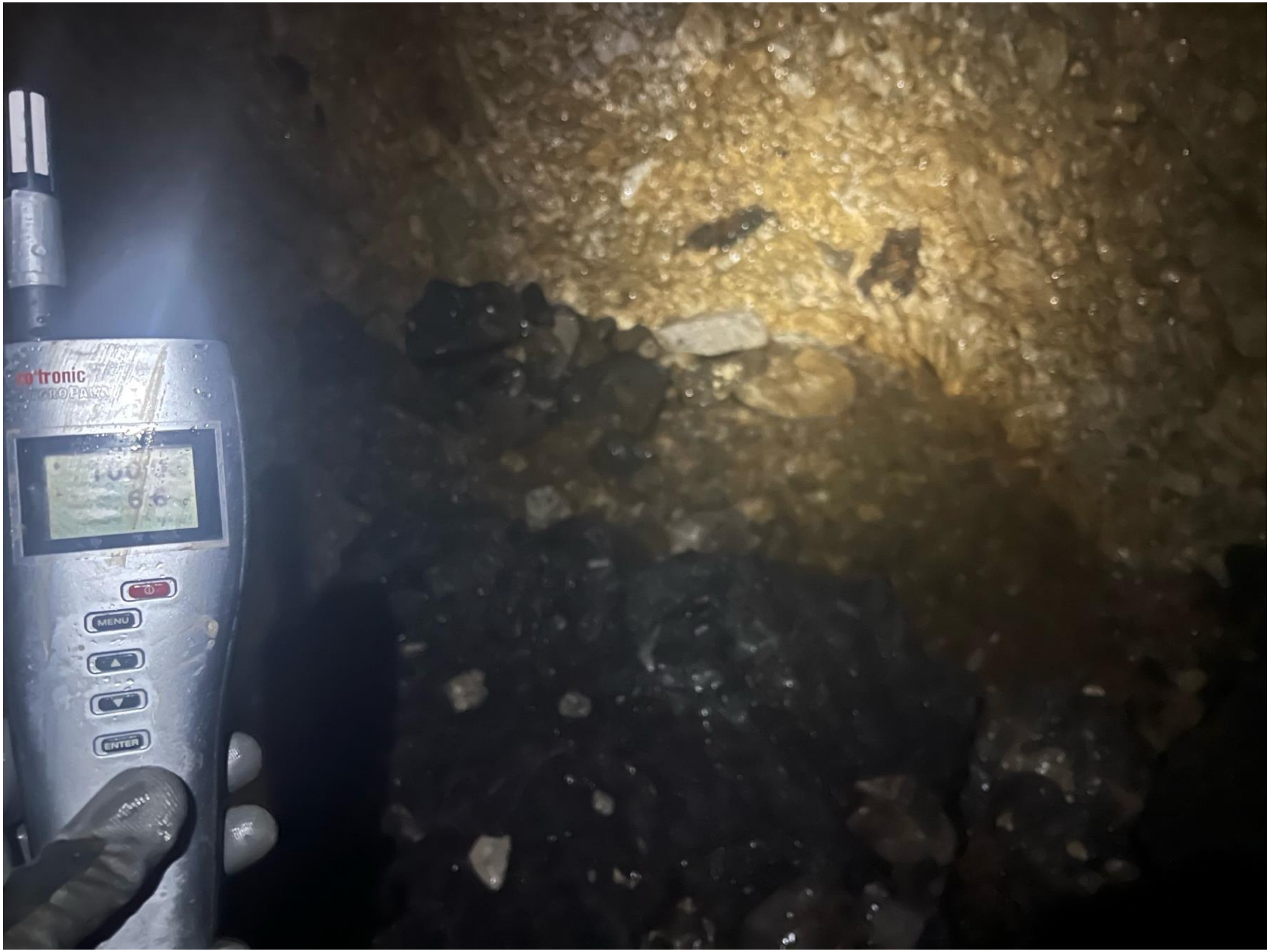

### gourgouthakas_-0678_siphon_1

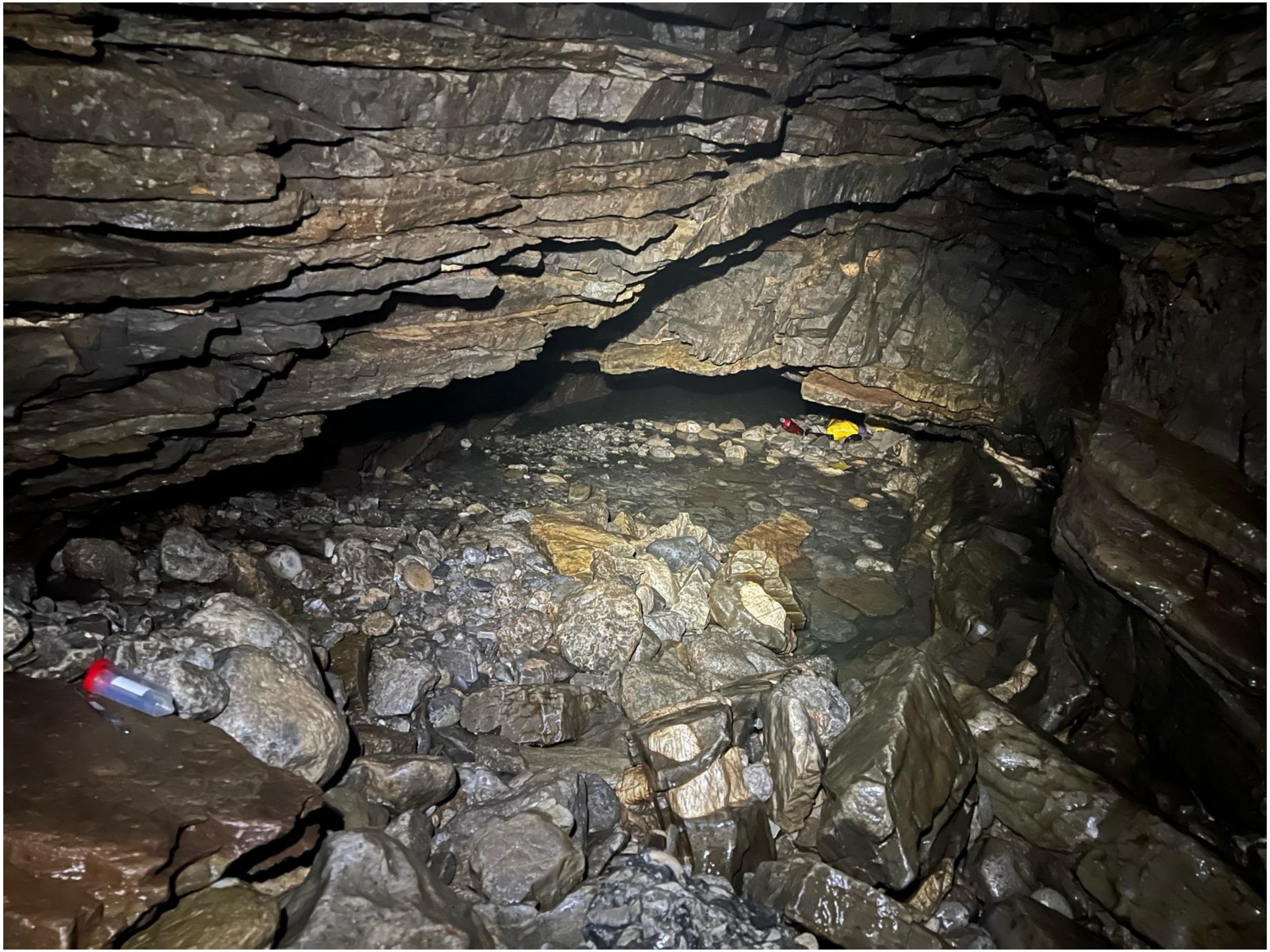

### gourgouthakas_-0678_siphon_2

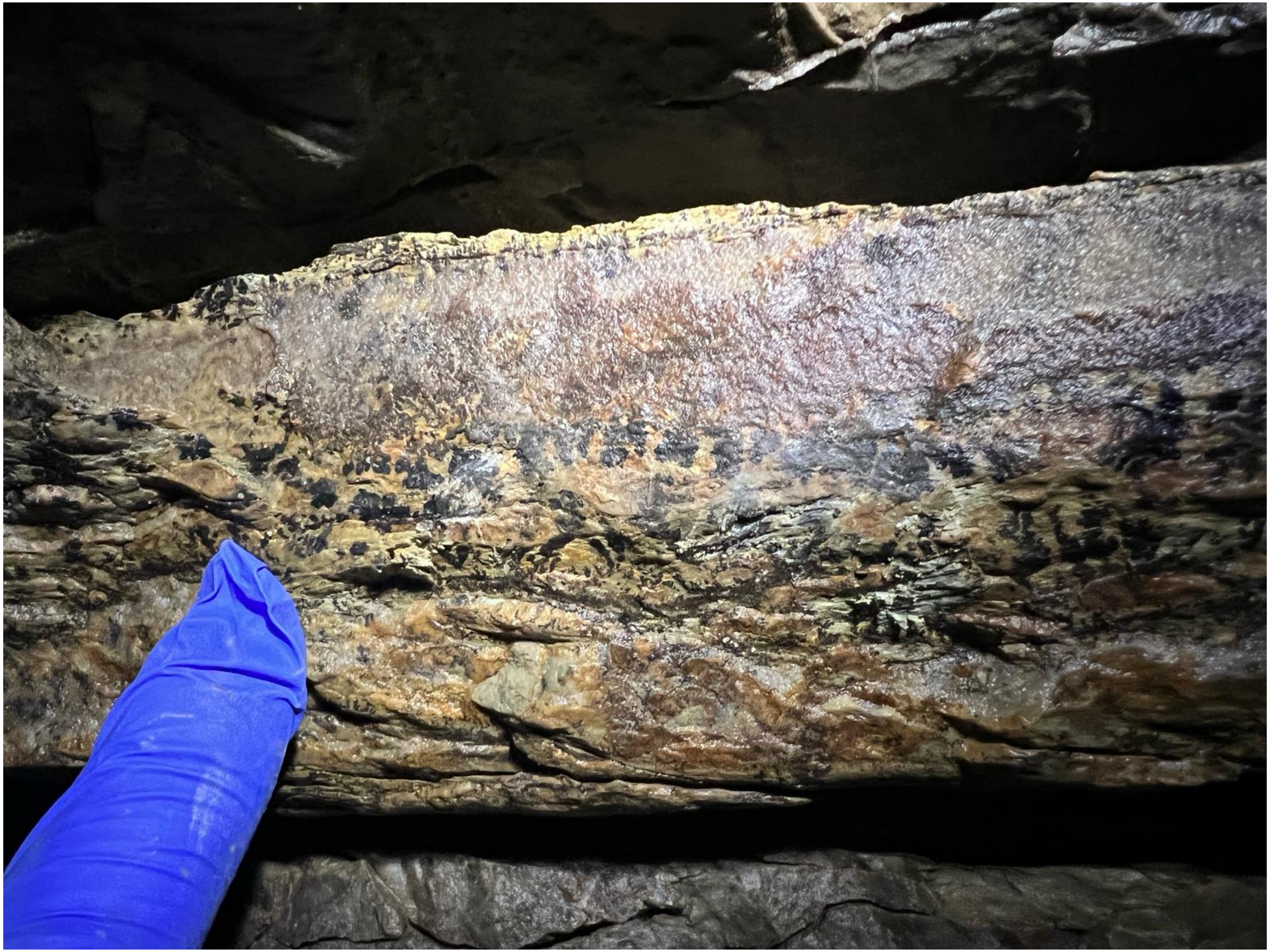

### gourgouthakas_-0678_siphon_3

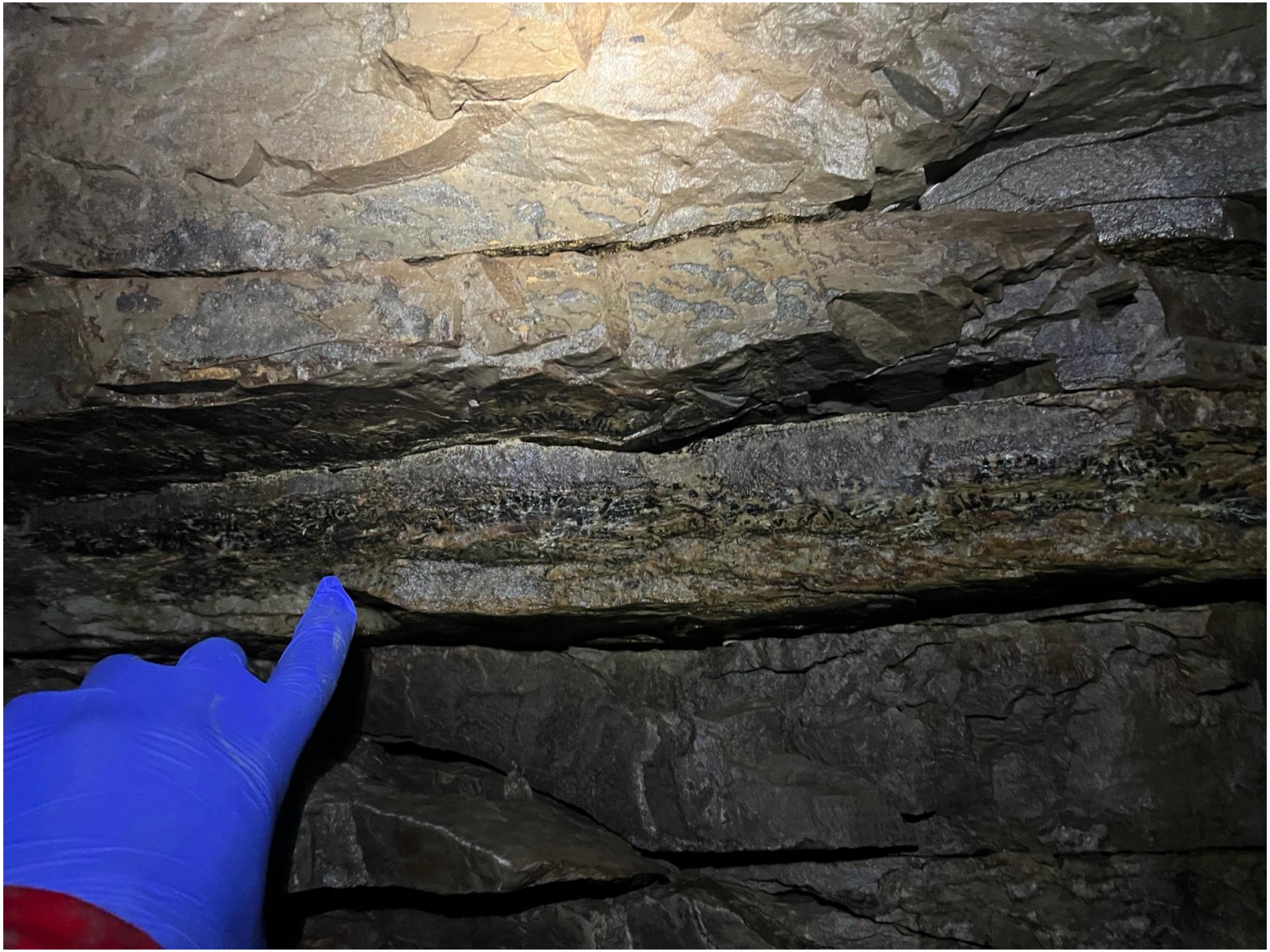

### gourgouthakas_-0678_siphon_4

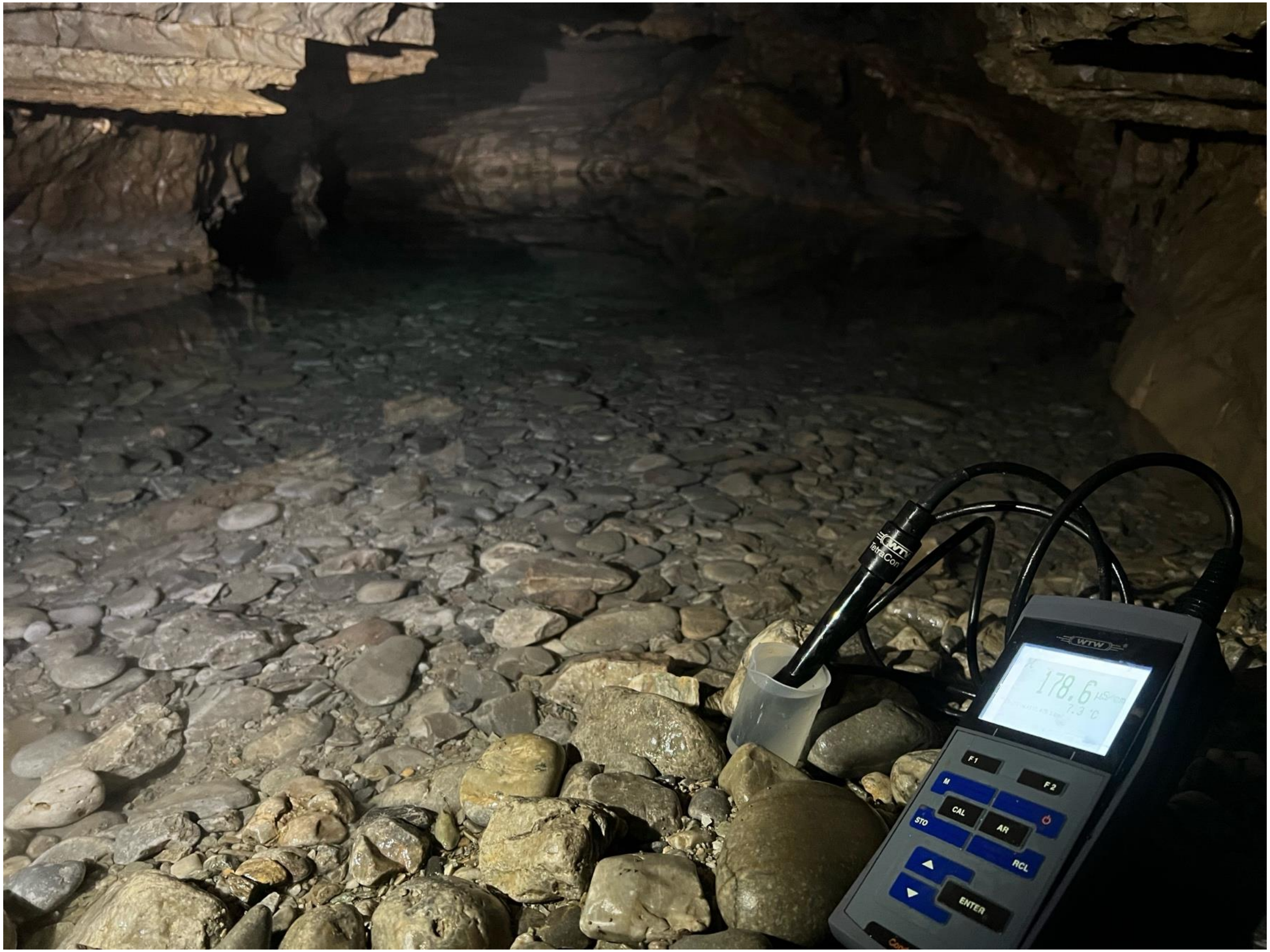

### gourgouthakas_-0678_siphon_5

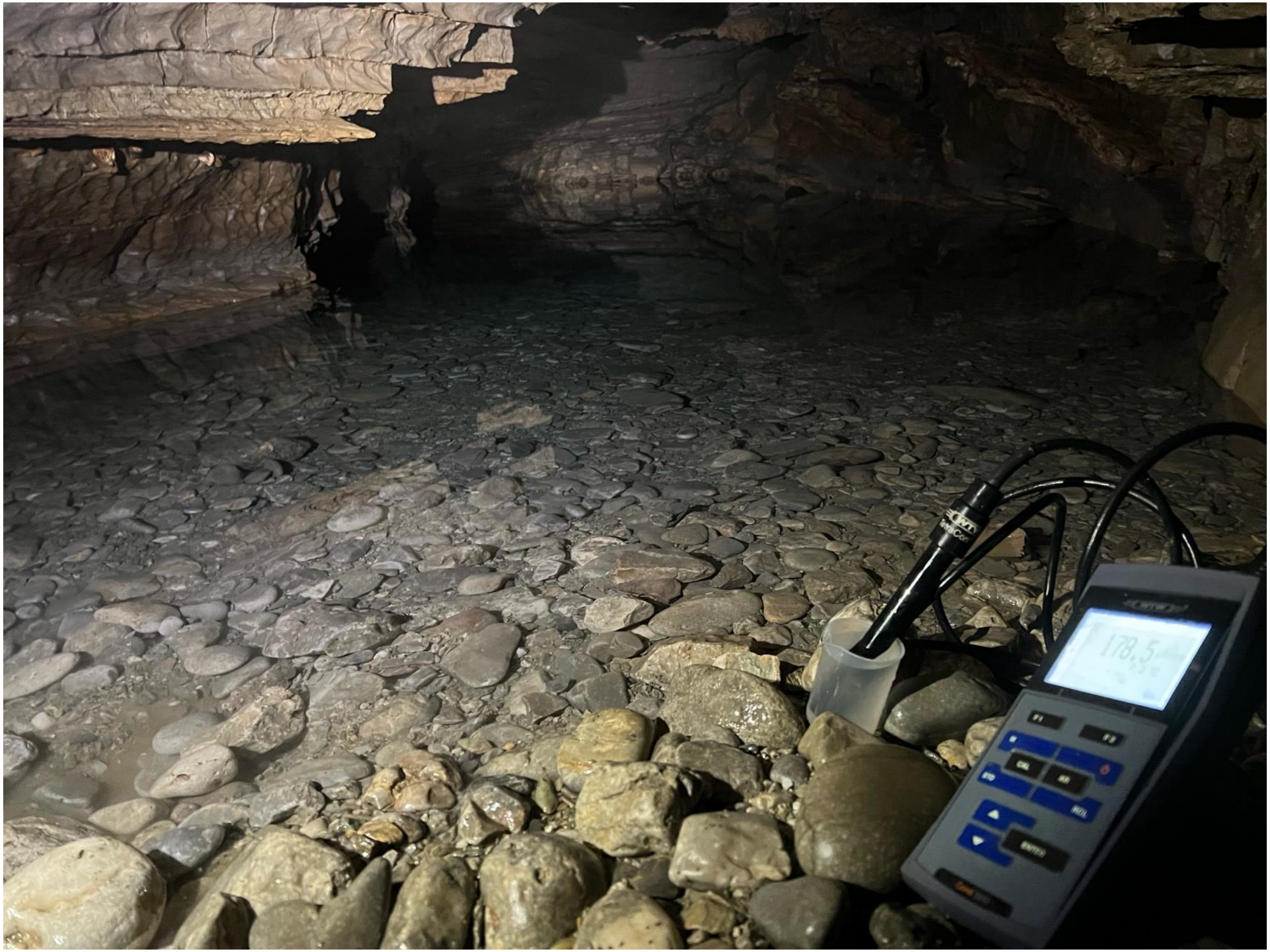

### gourgouthakas_-0678_siphon_6

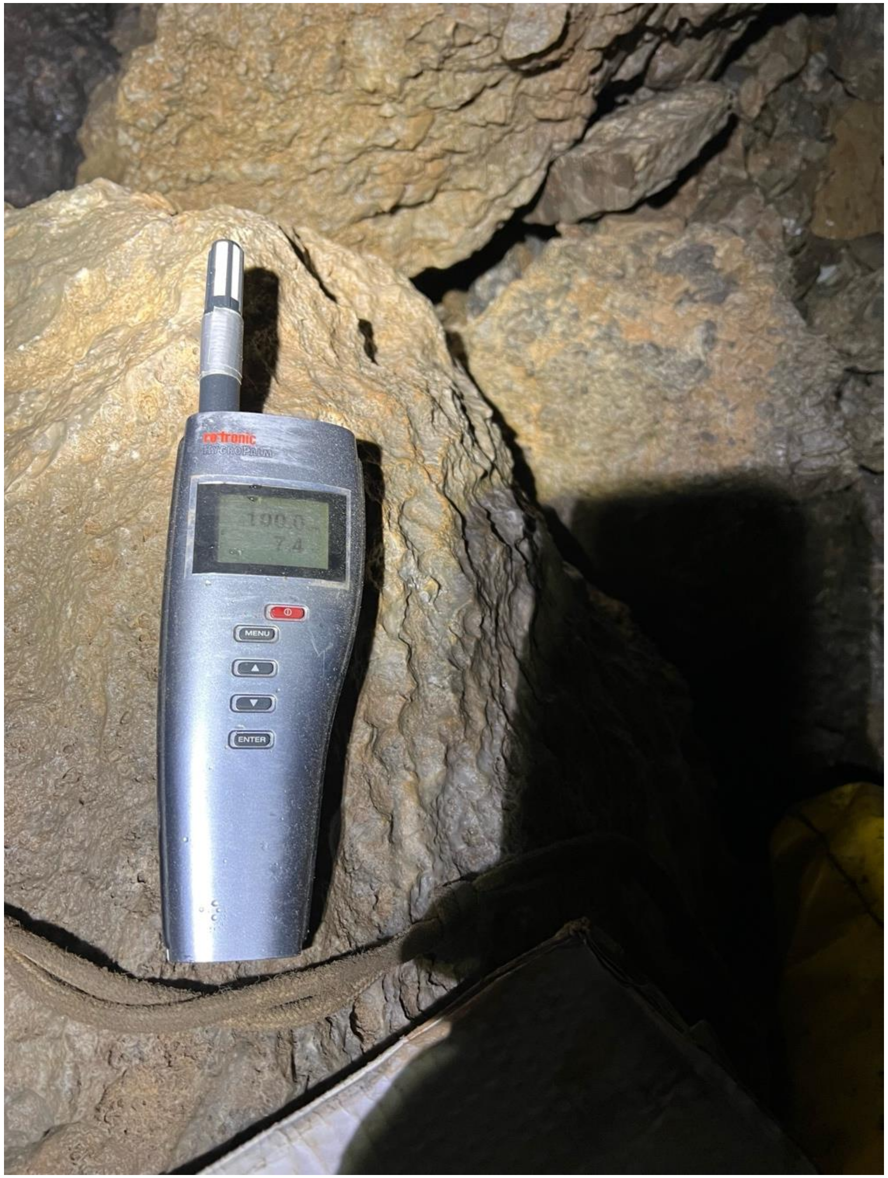

### gourgouthakas_-0700_camp_1

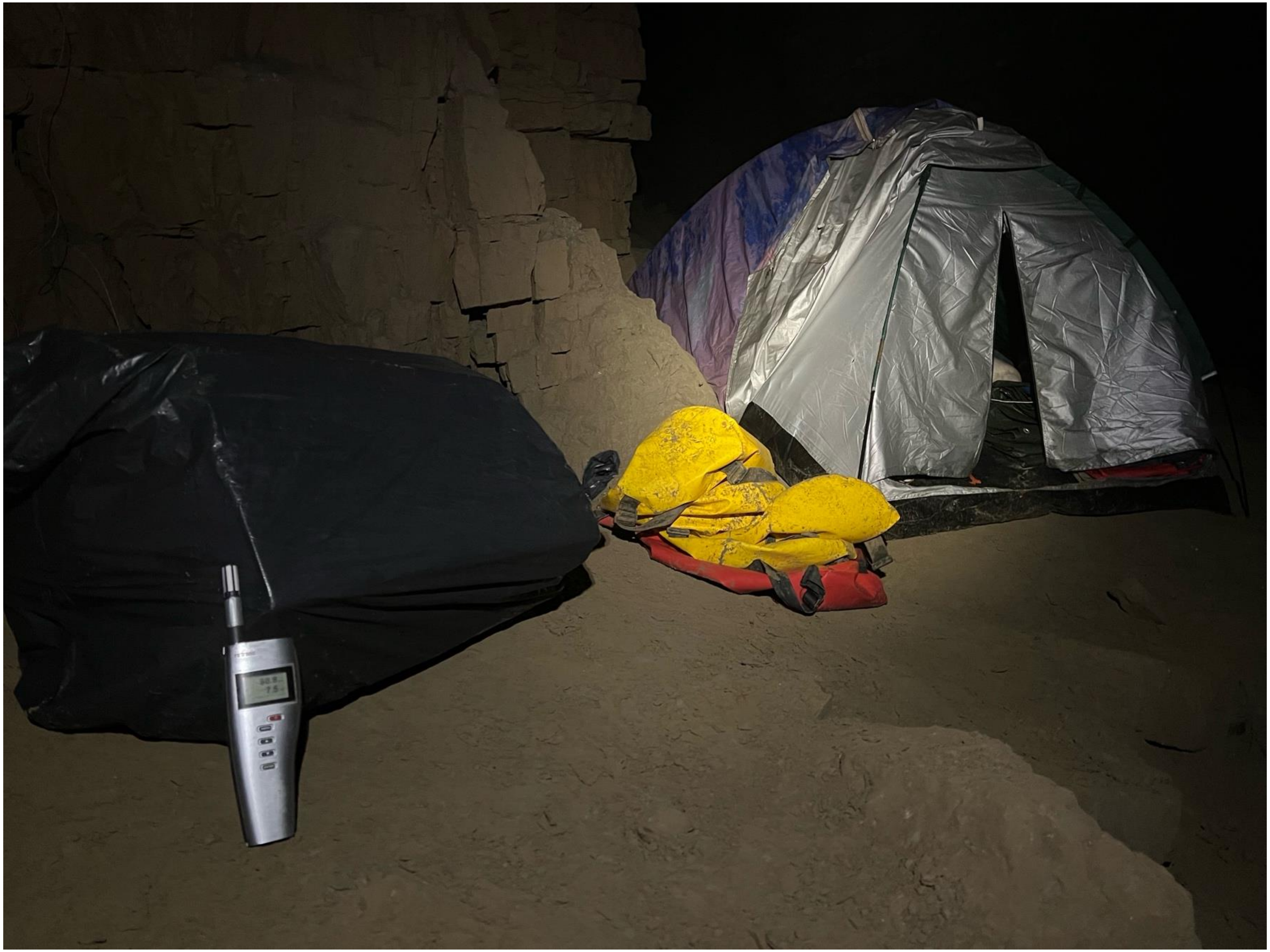

### gourgouthakas_-0700_camp_2

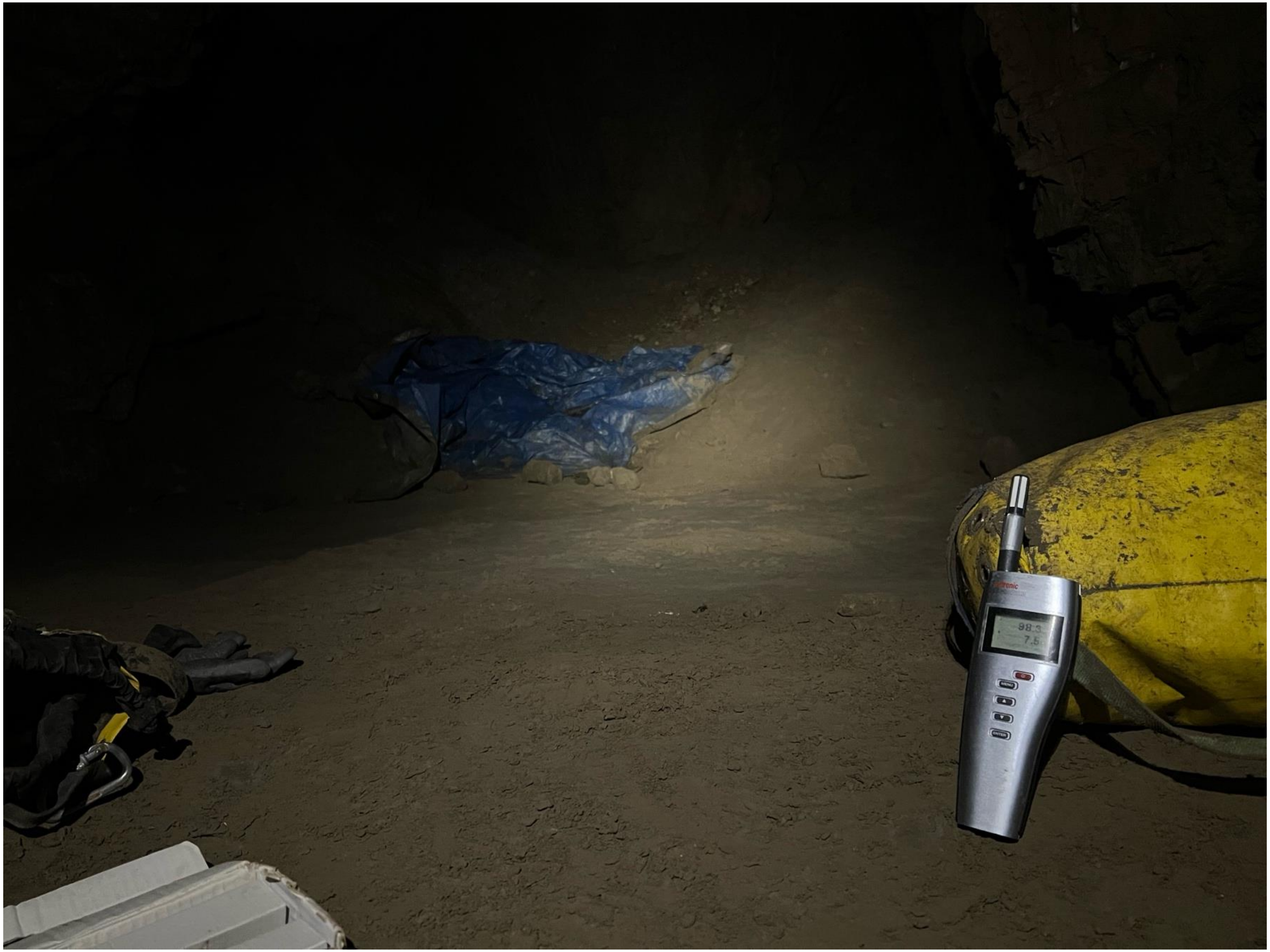

### gourgouthakas_-0713_marcel_waterfall_1

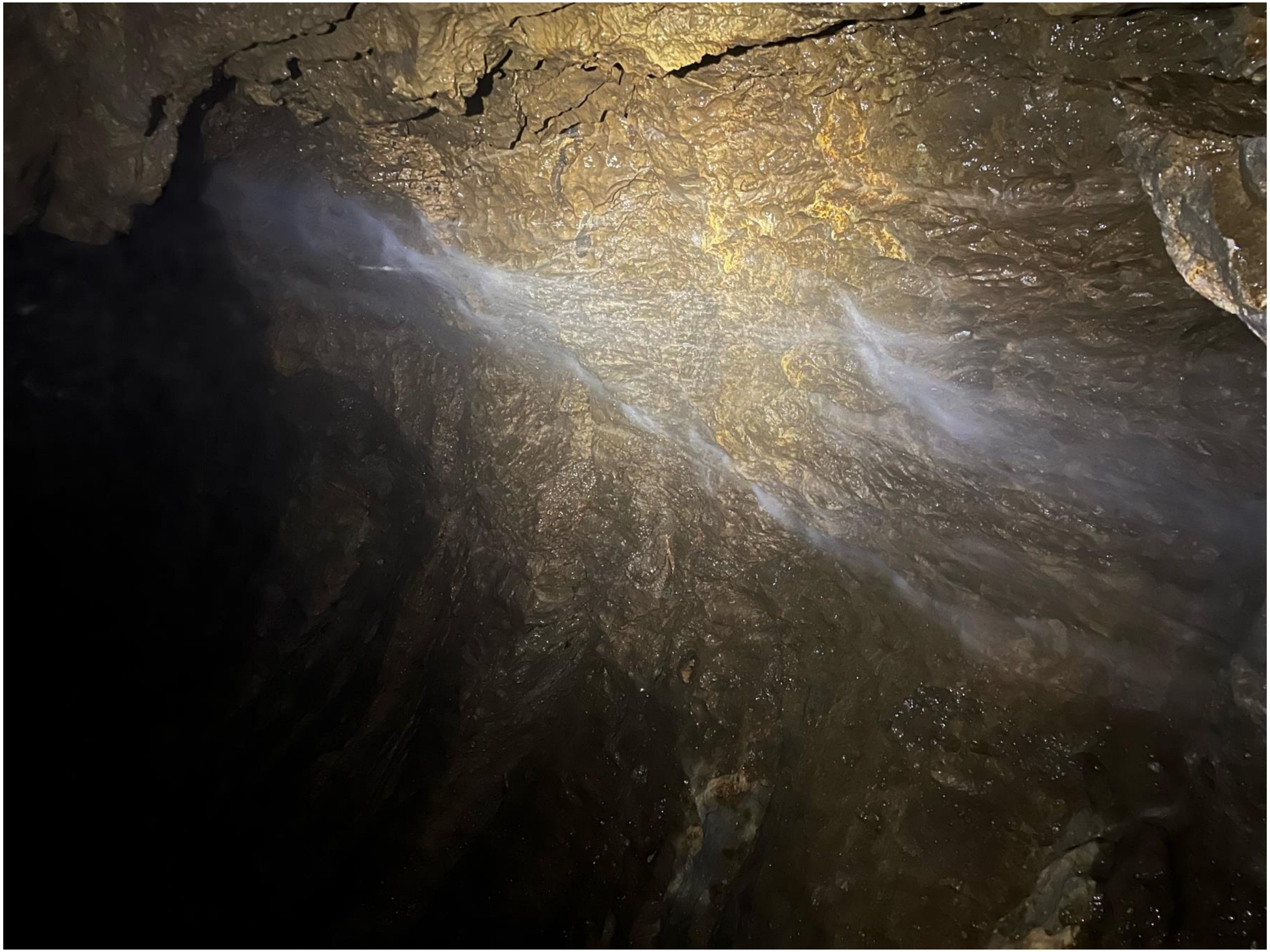

### gourgouthakas_-0713_marcel_waterfall_2

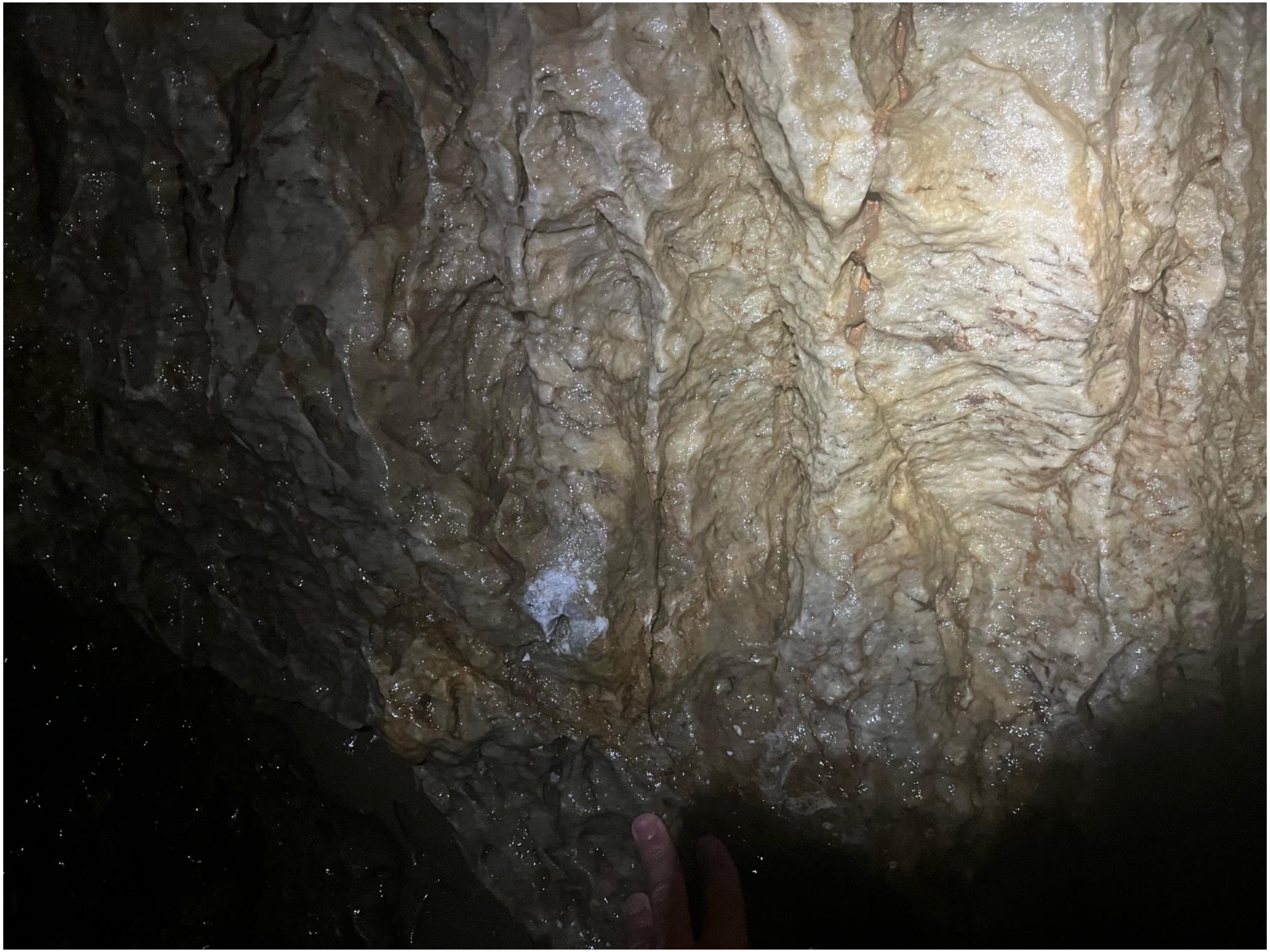

### gourgouthakas_-0713_marcel_waterfall_3

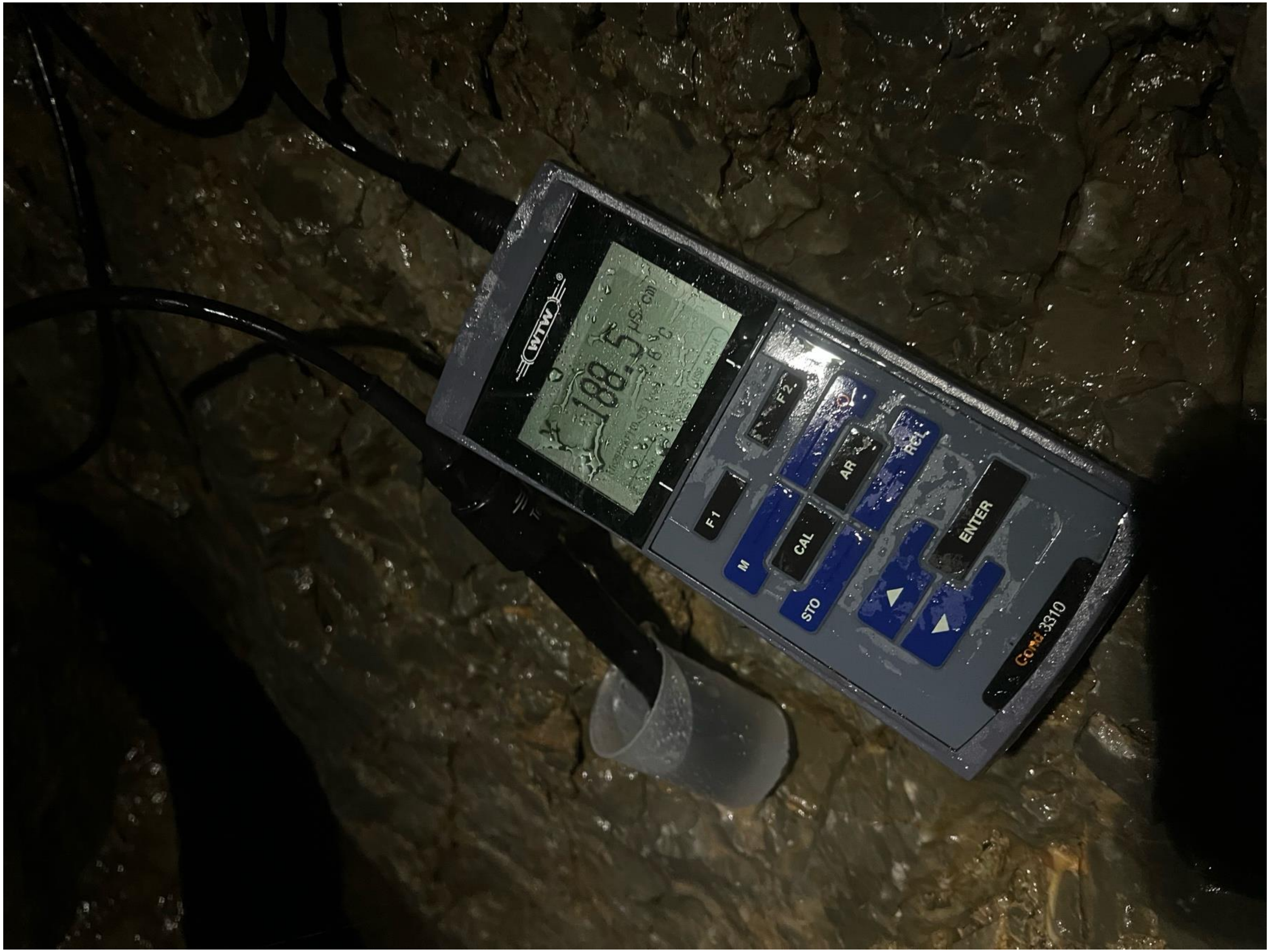

### gourgouthakas_-0713_marcel_waterfall_4

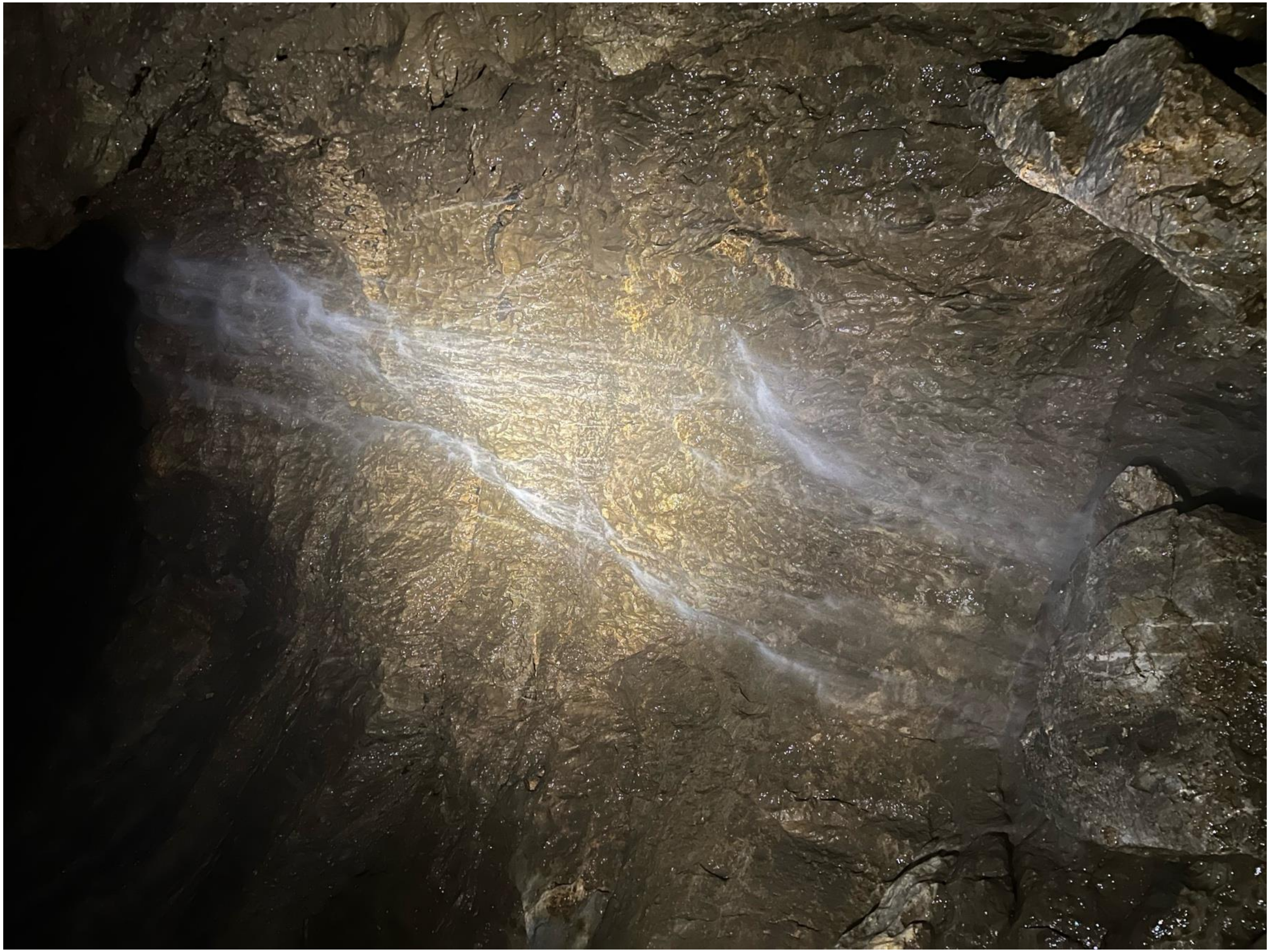

### gourgouthakas_-0713_marcel_waterfall_5

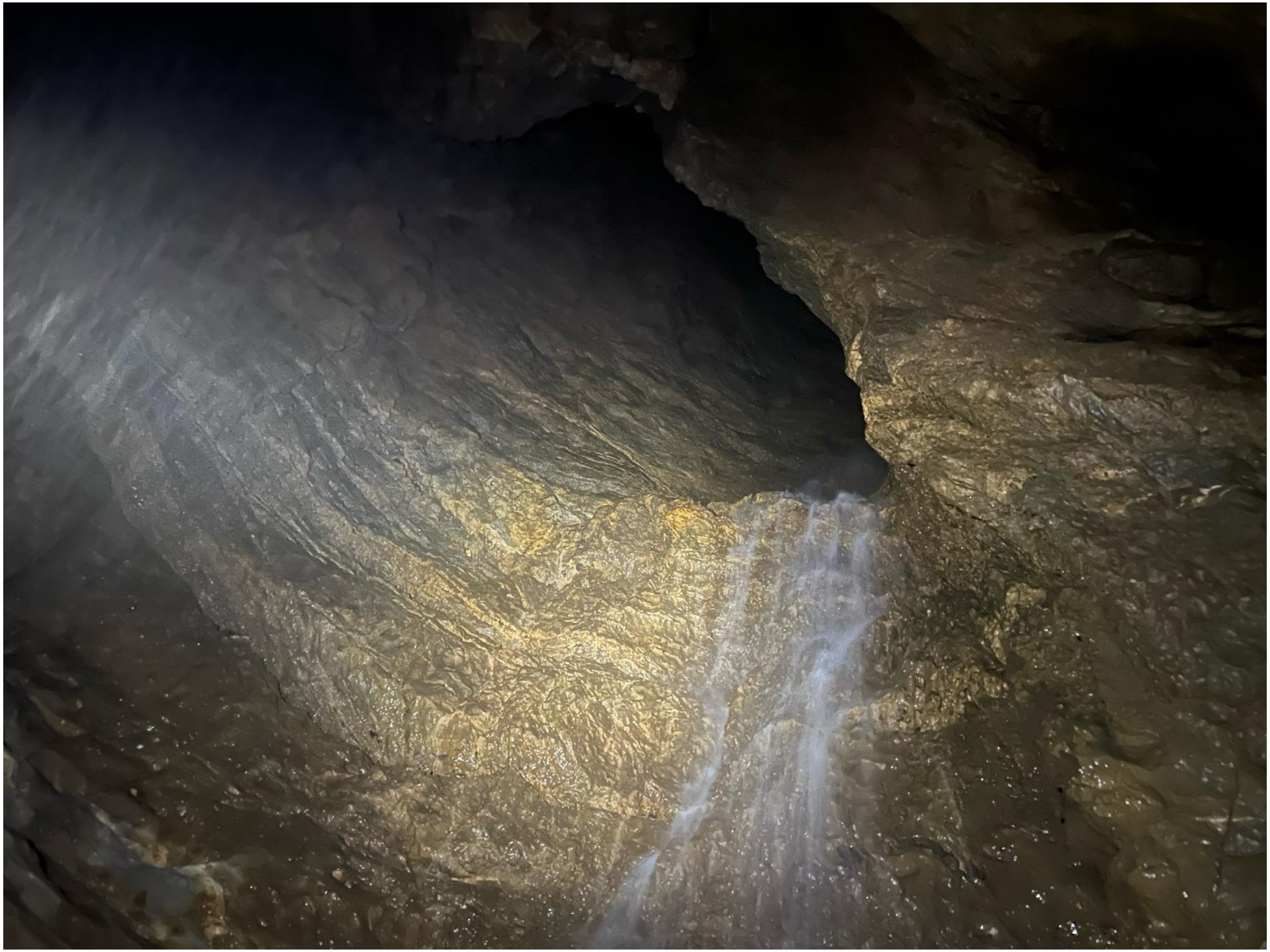

### gourgouthakas_-0900_water-curtain_1

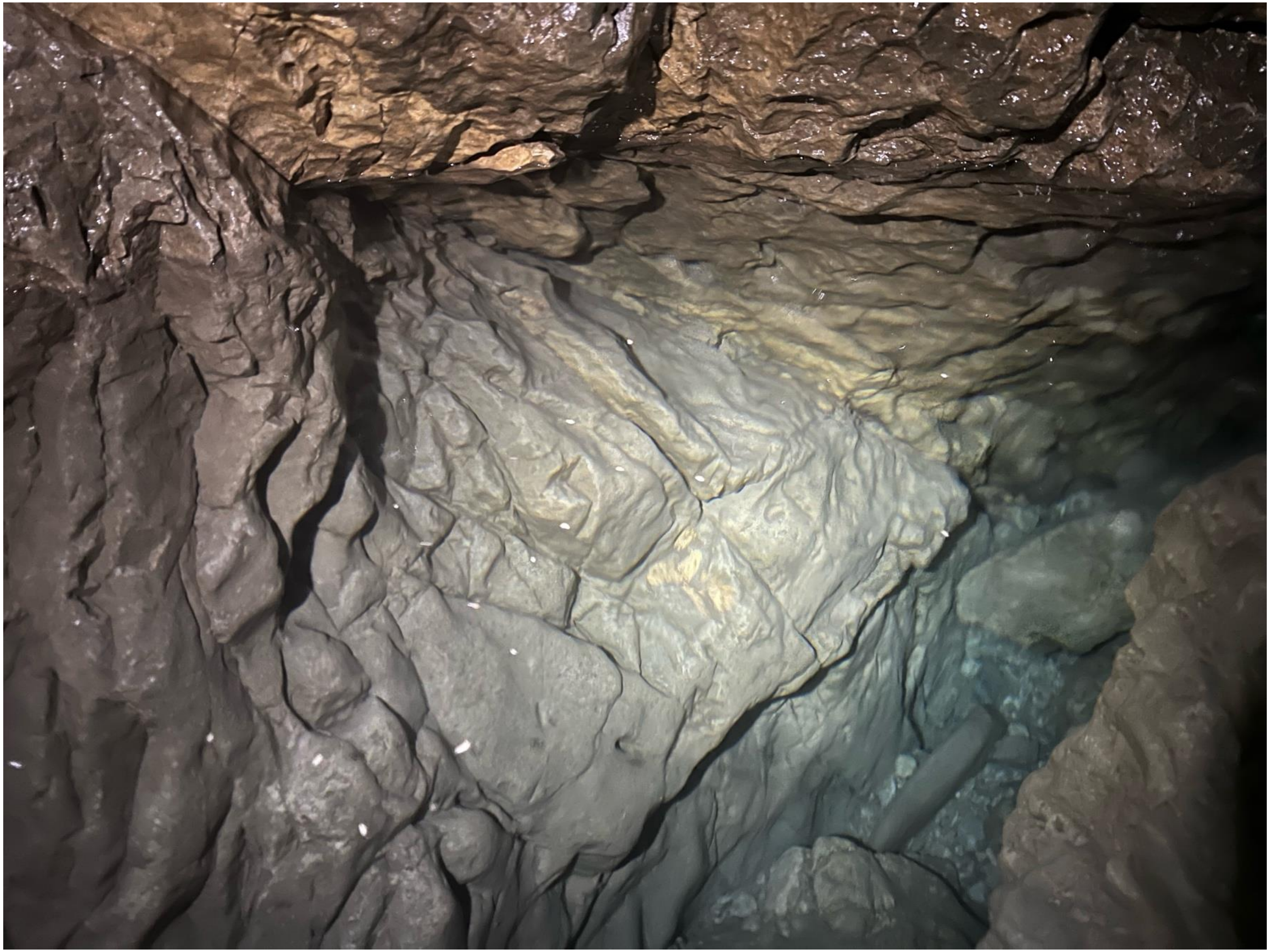

### gourgouthakas_-0900_water-curtain_3

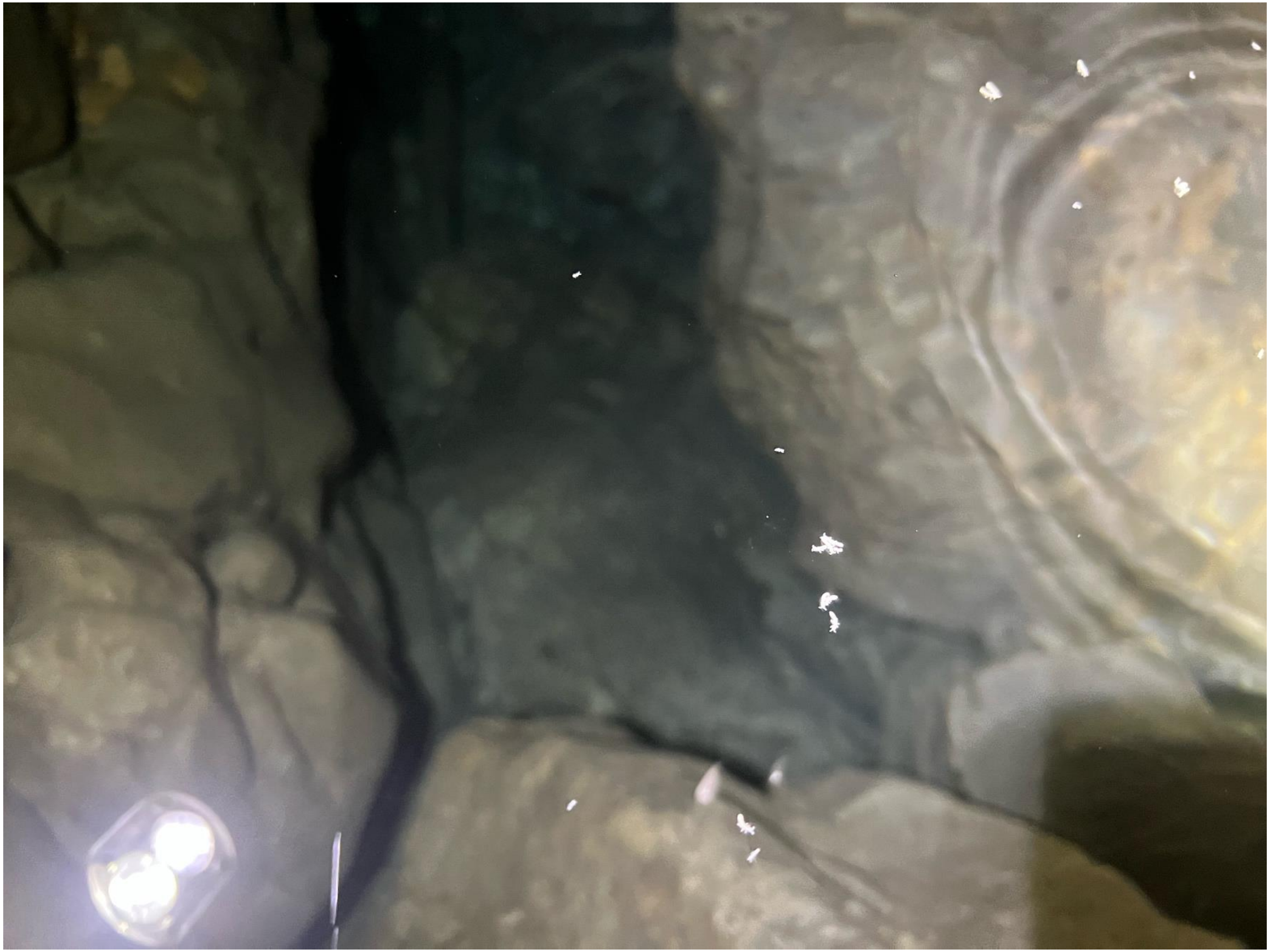

### gourgouthakas_-0900_water-curtain_4

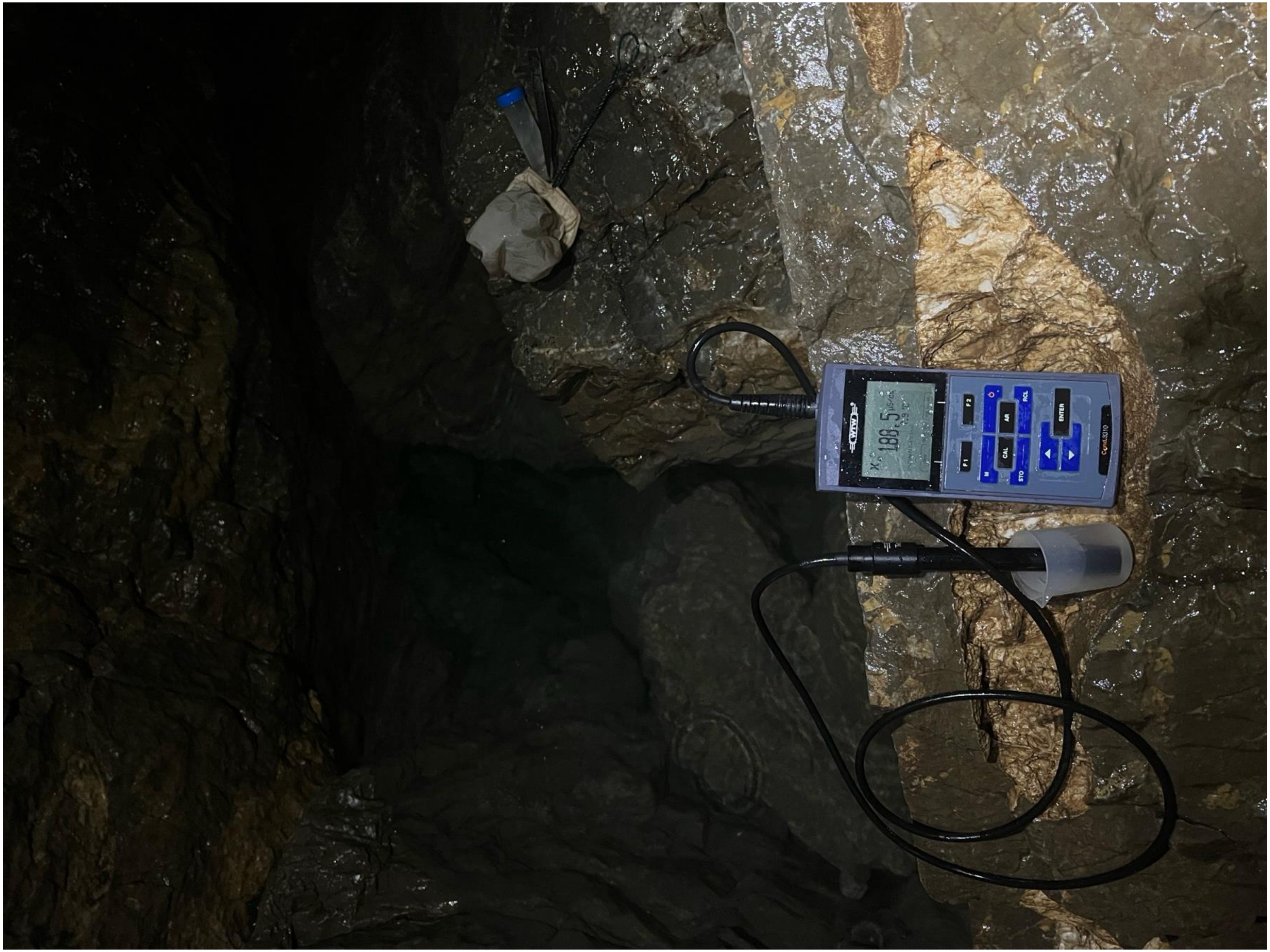

### gourgouthakas_-0900_water-curtain_6

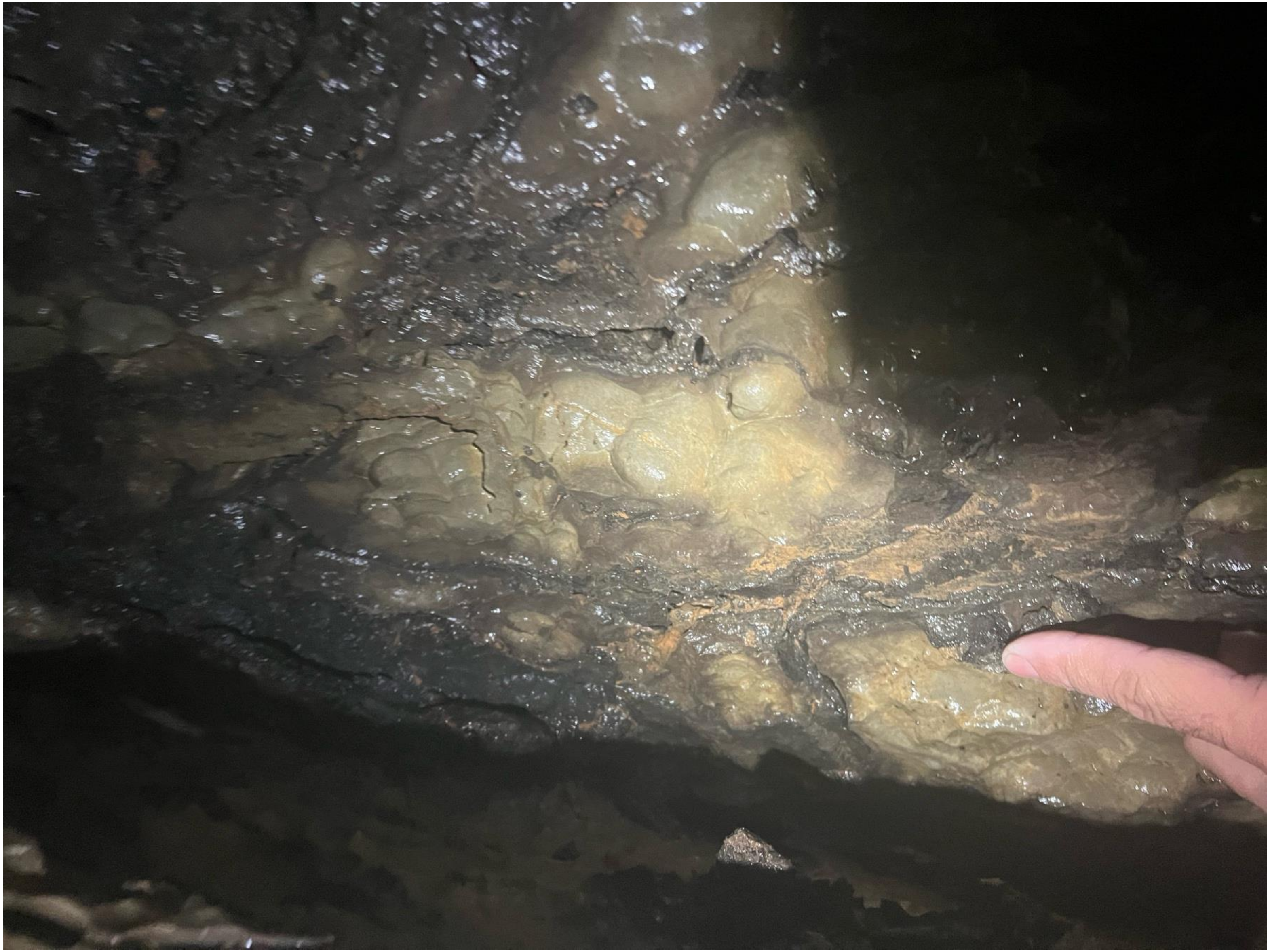

### gourgouthakas_-0900_water-curtain_7

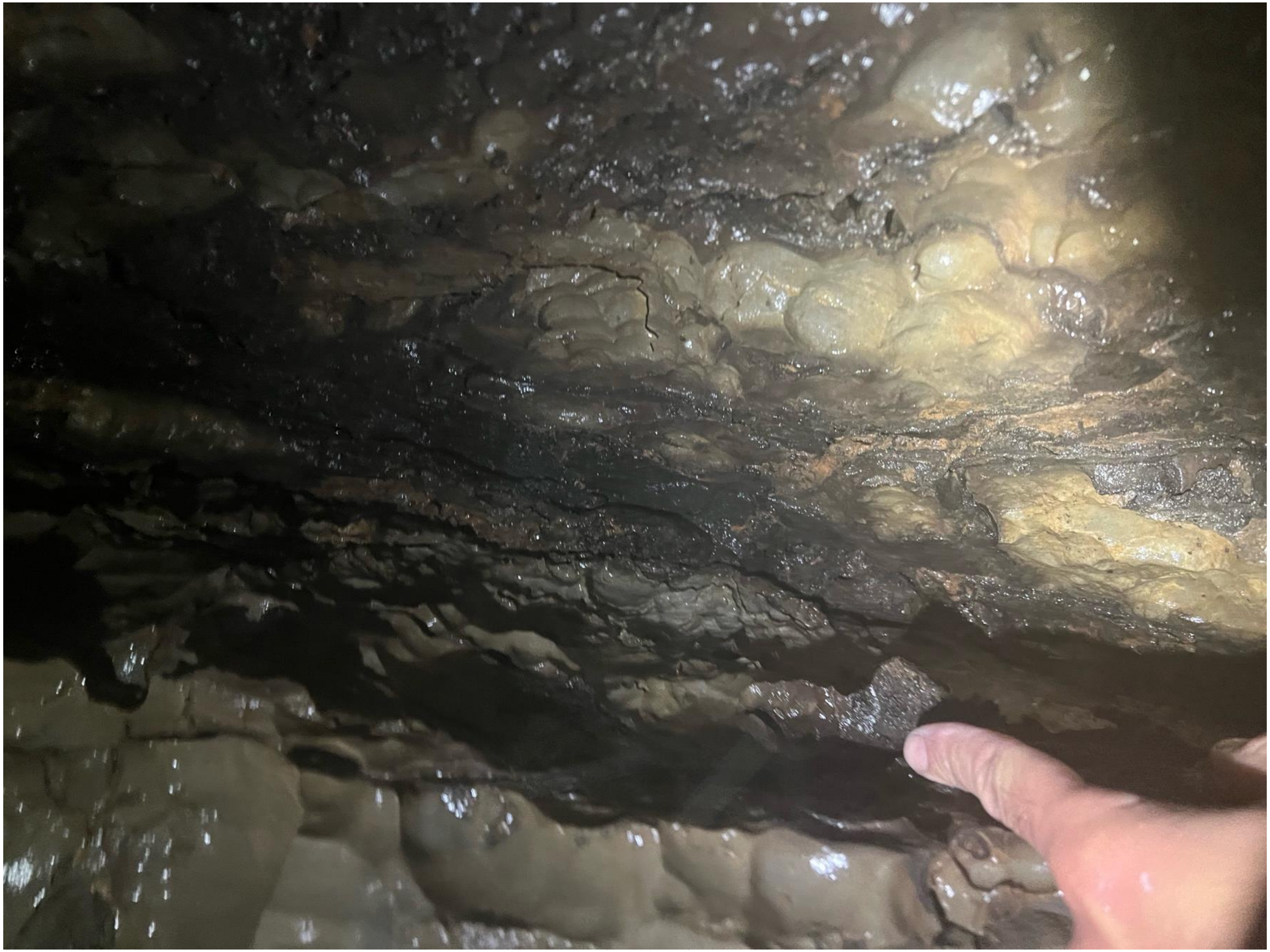

### gourgouthakas_-0900_water-curtain_8

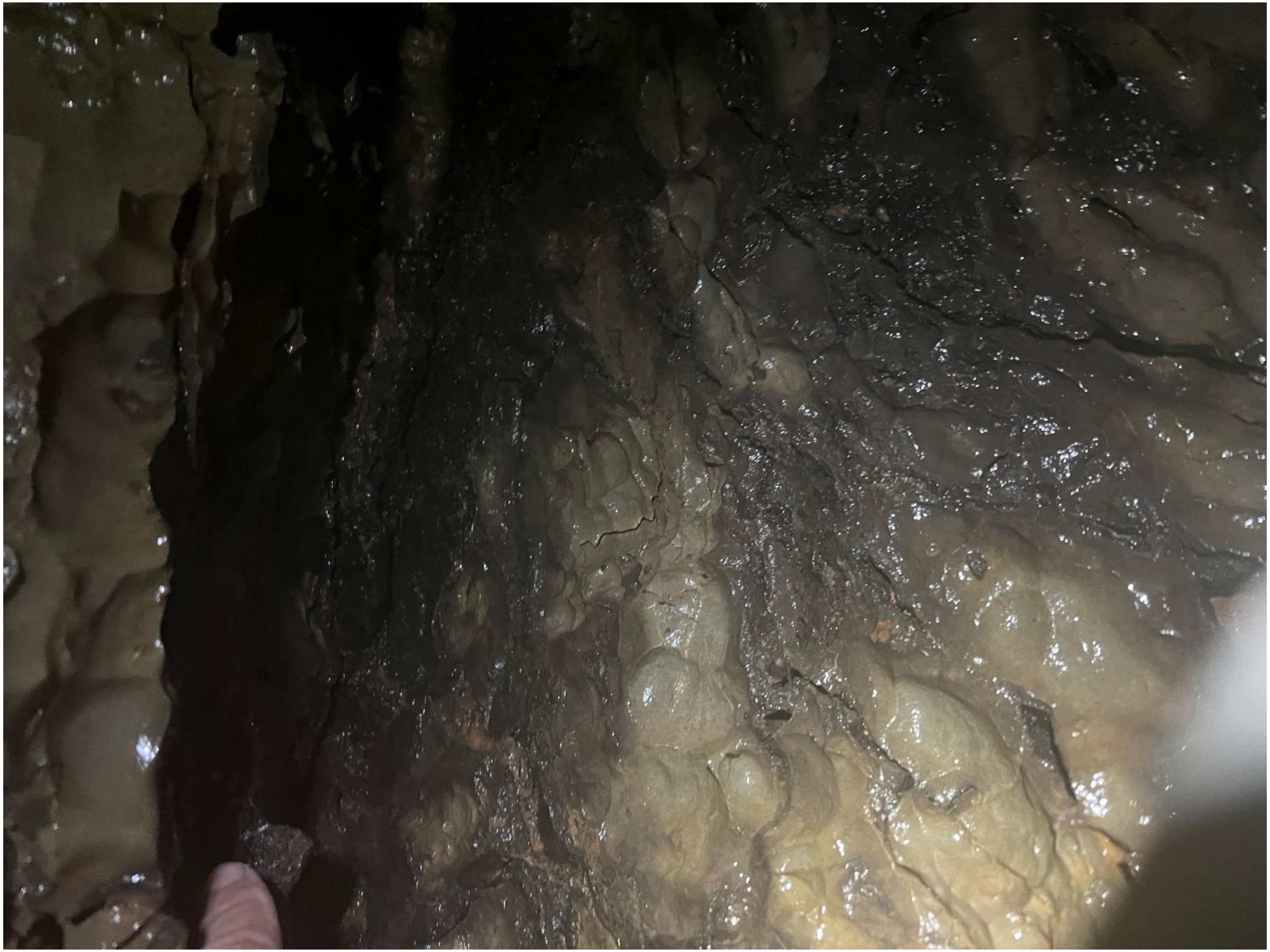

### gourgouthakas_-1050_traverse_1

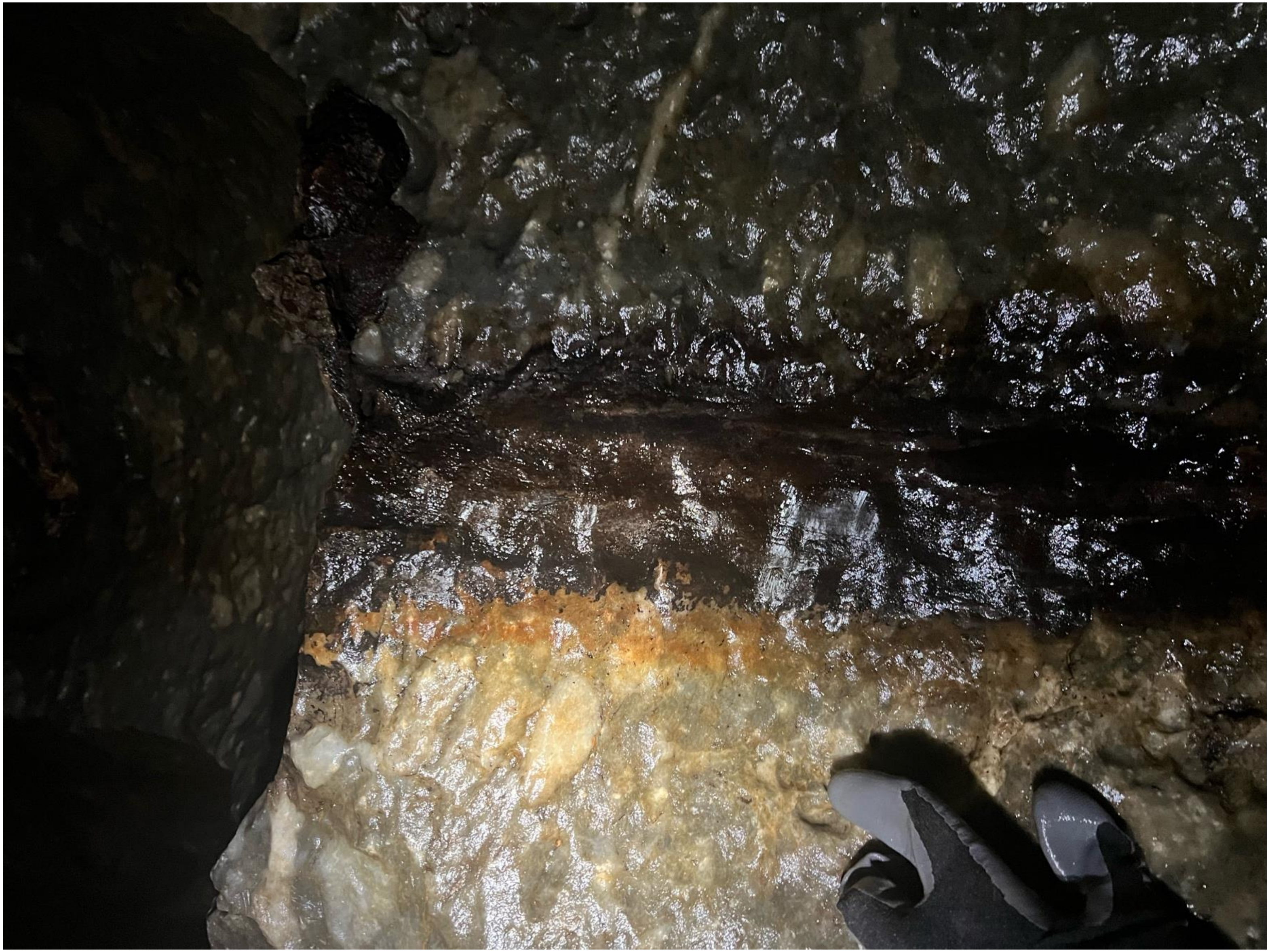

### gourgouthakas_-1050_traverse_2 2

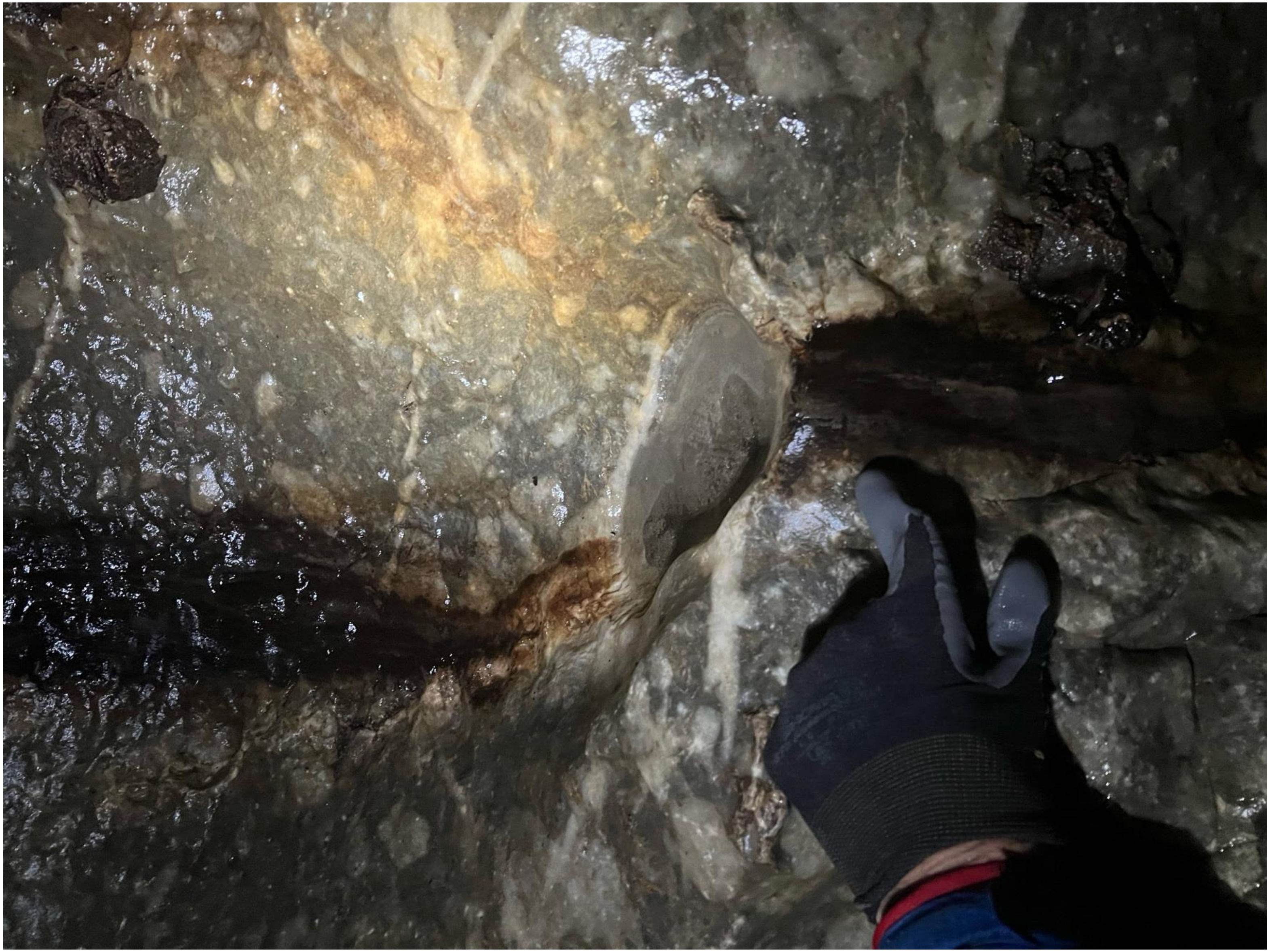

### gourgouthakas_-1050_traverse_2 3

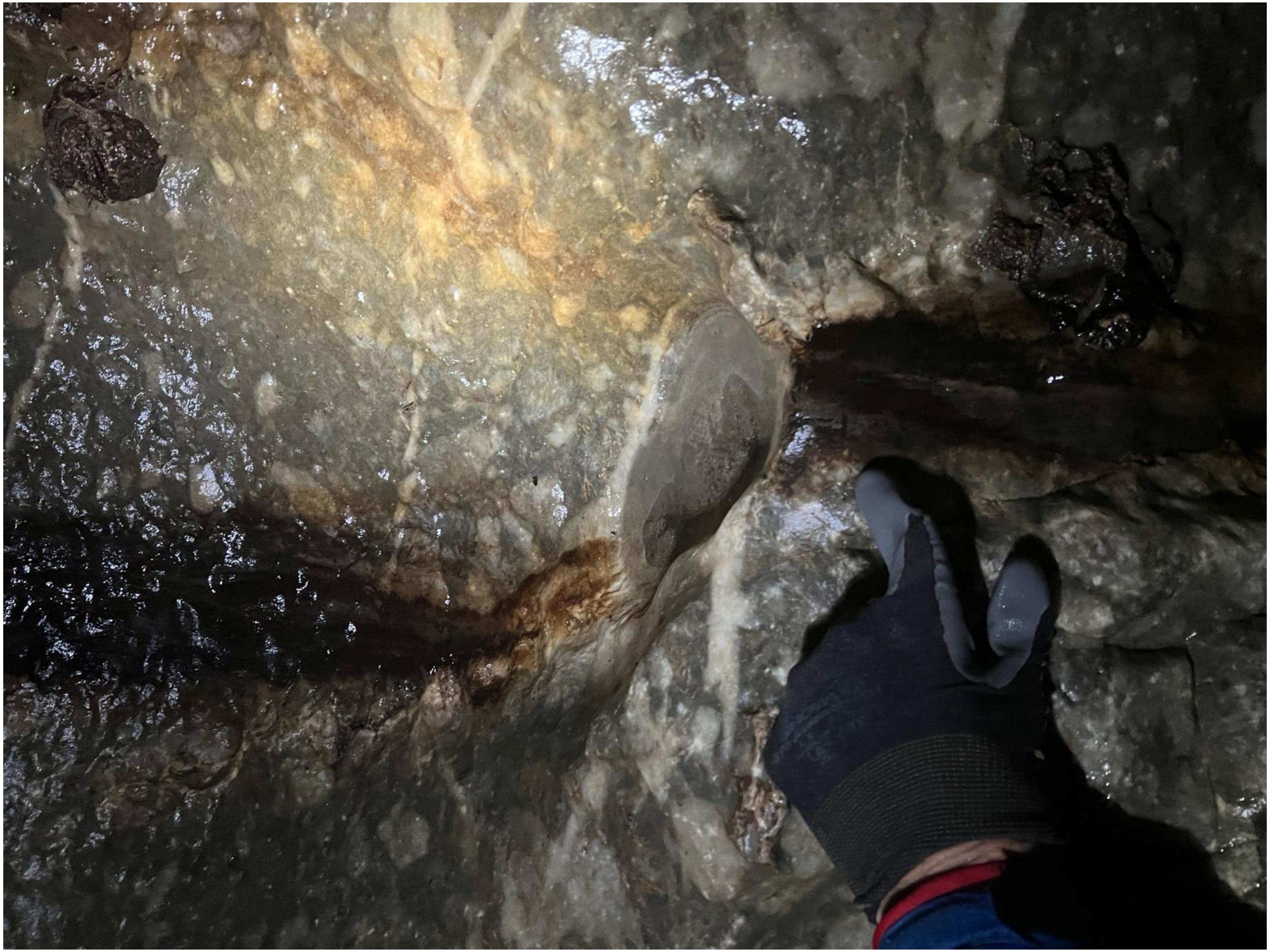

### gourgouthakas_-1050_traverse_2

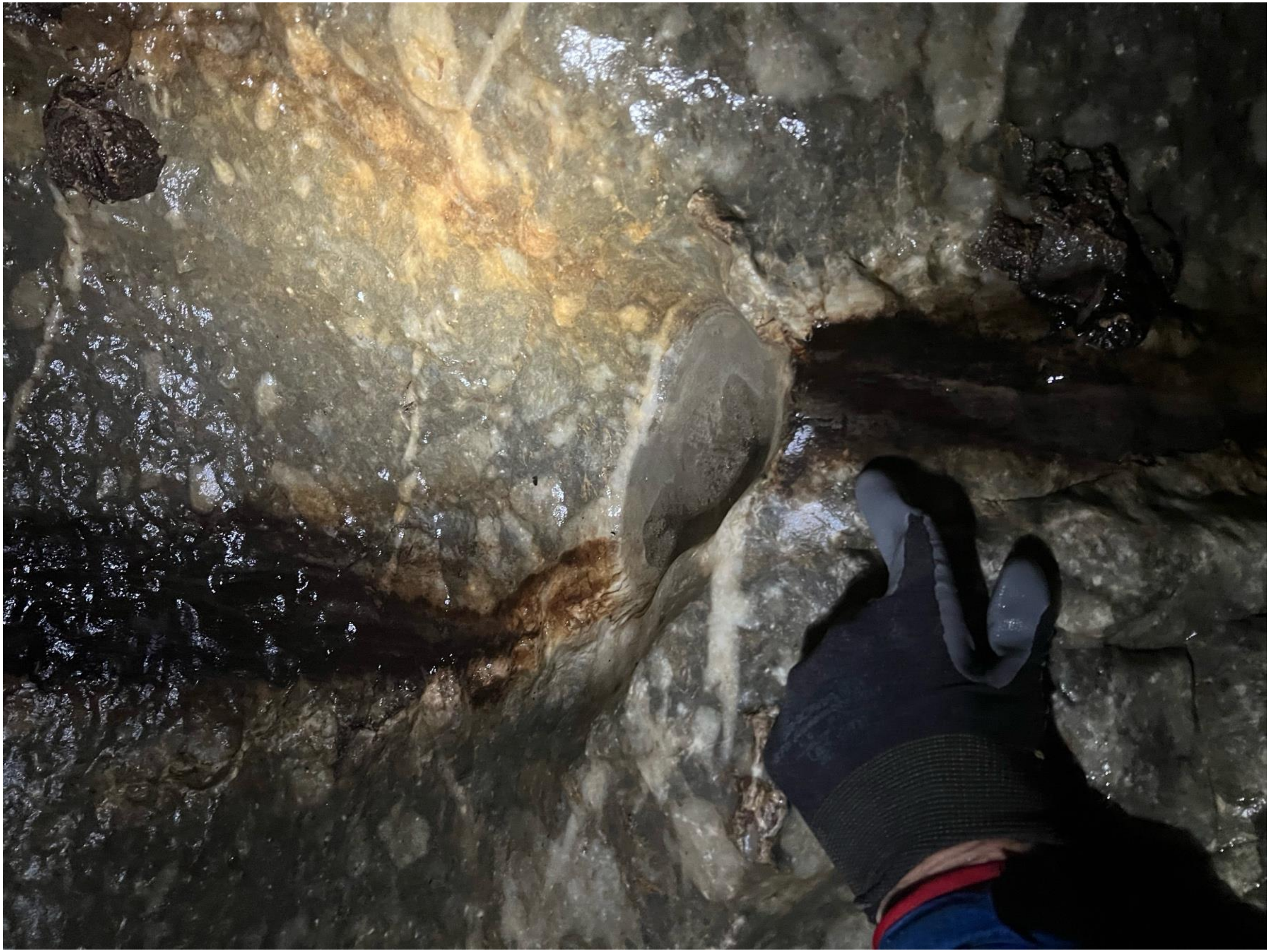

### gourgouthakas_-1050_traverse_3

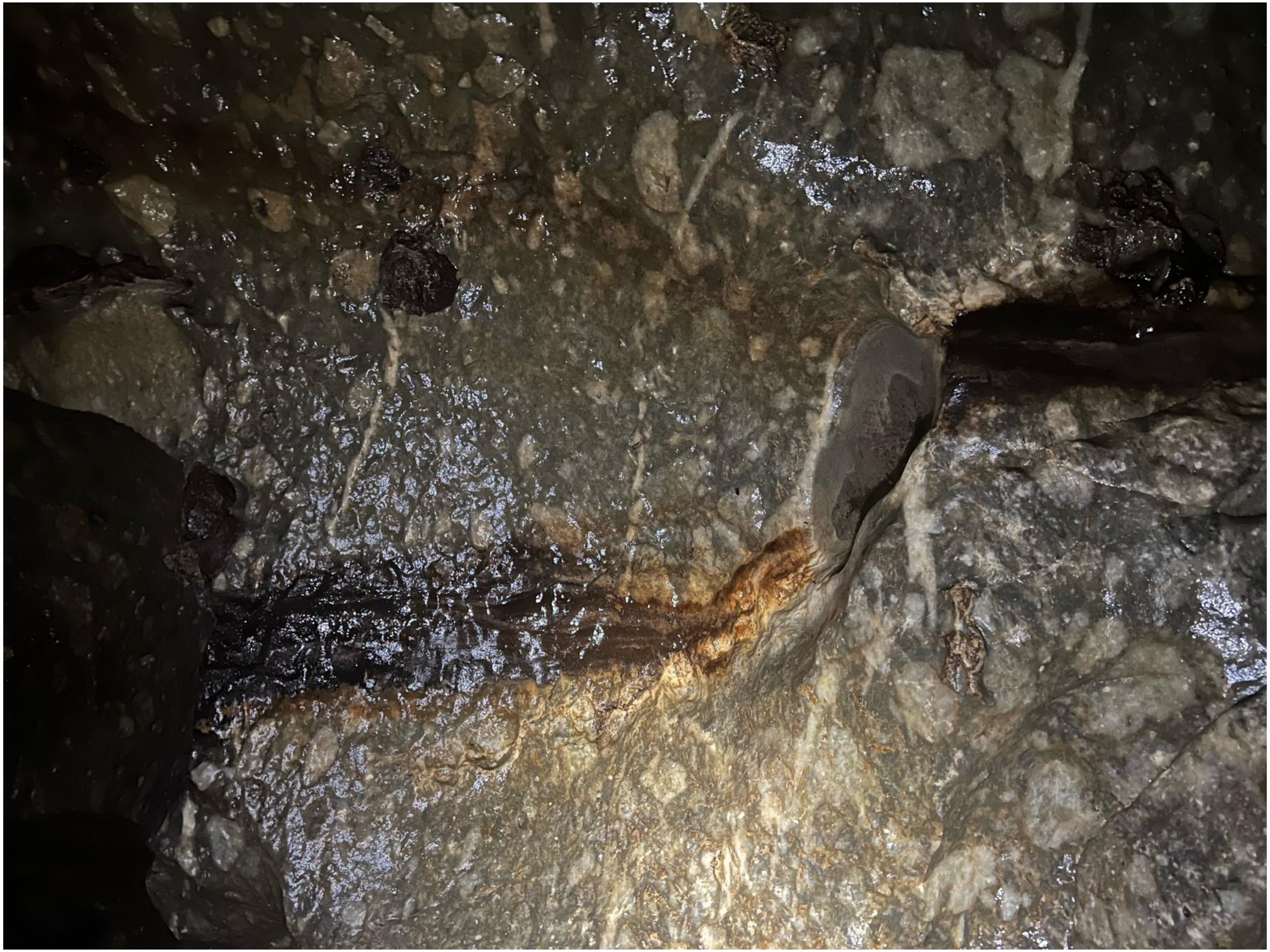

### gourgouthakas_-1100_siphon_1

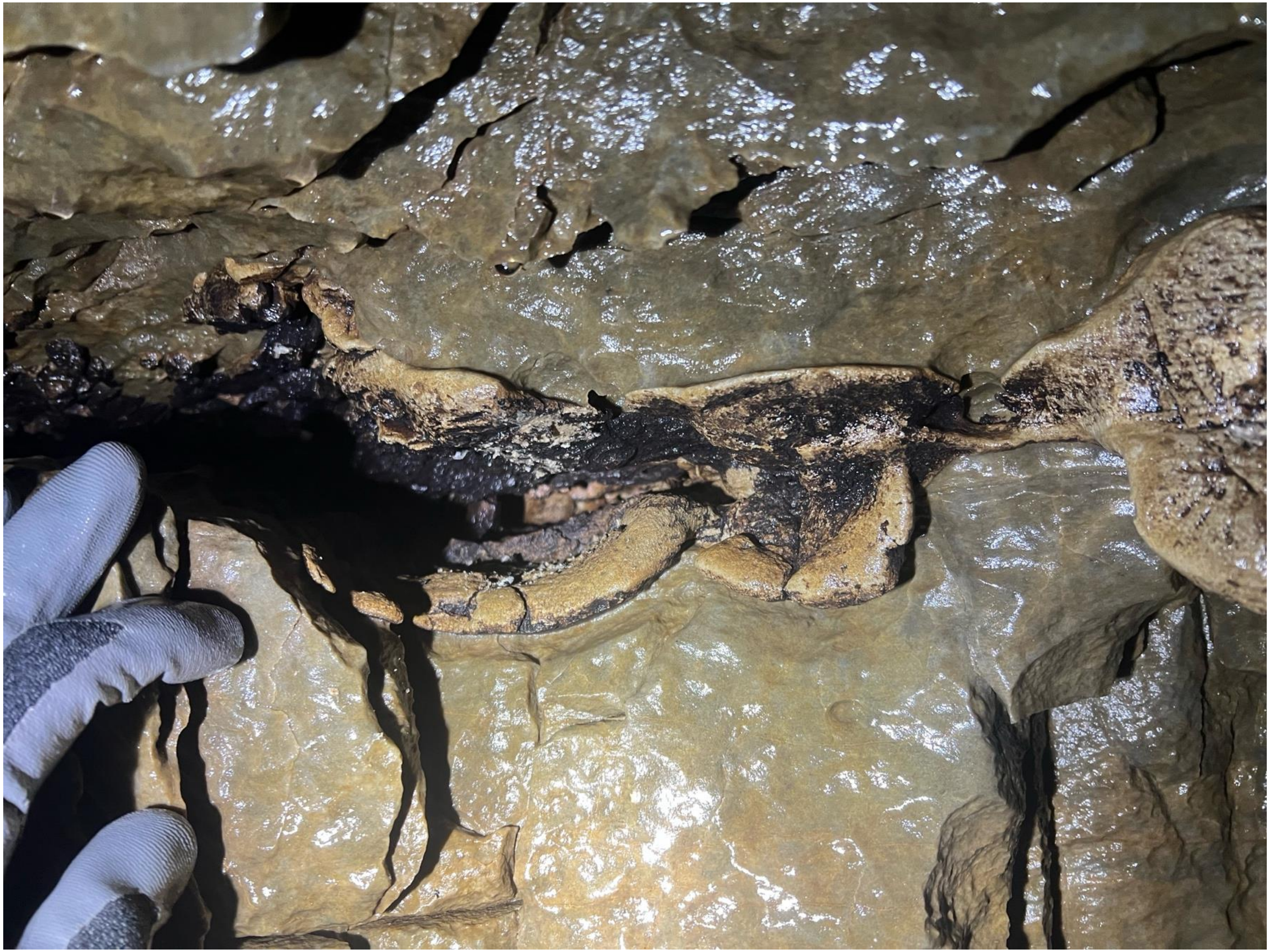

### gourgouthakas_-1100_siphon_2

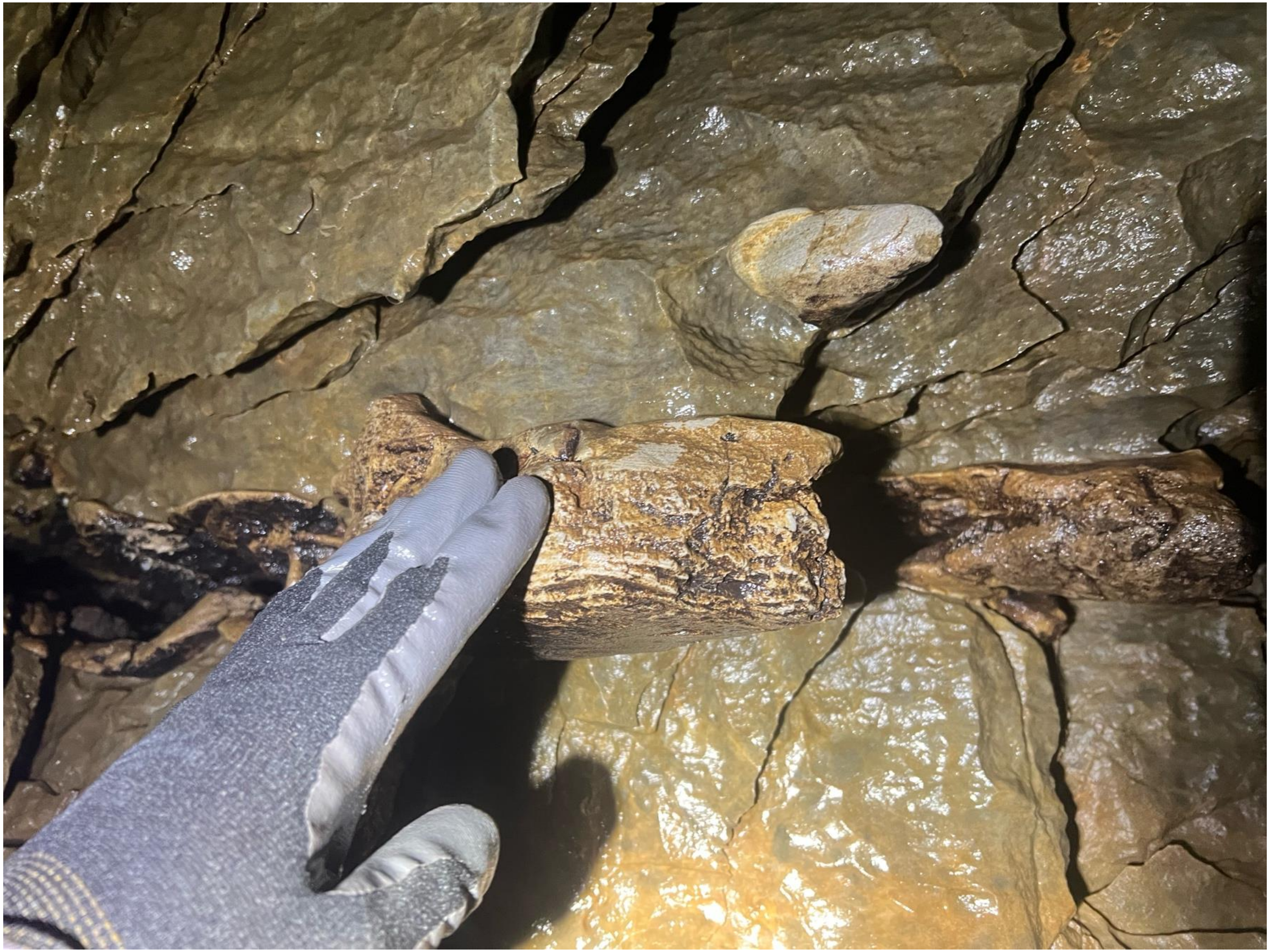

### gourgouthakas_-1100_siphon_3

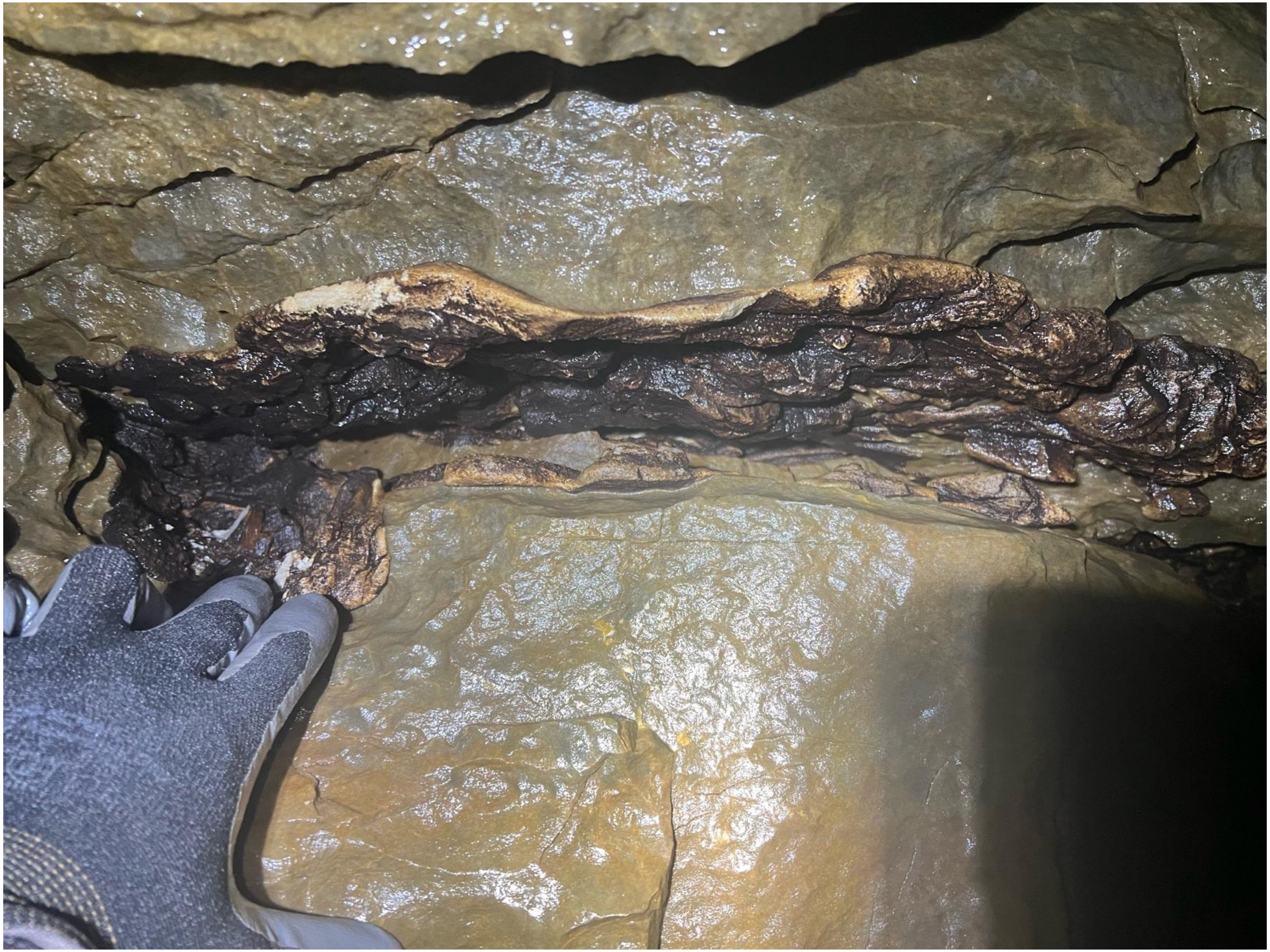

### gourgouthakas_-1100_siphon_4

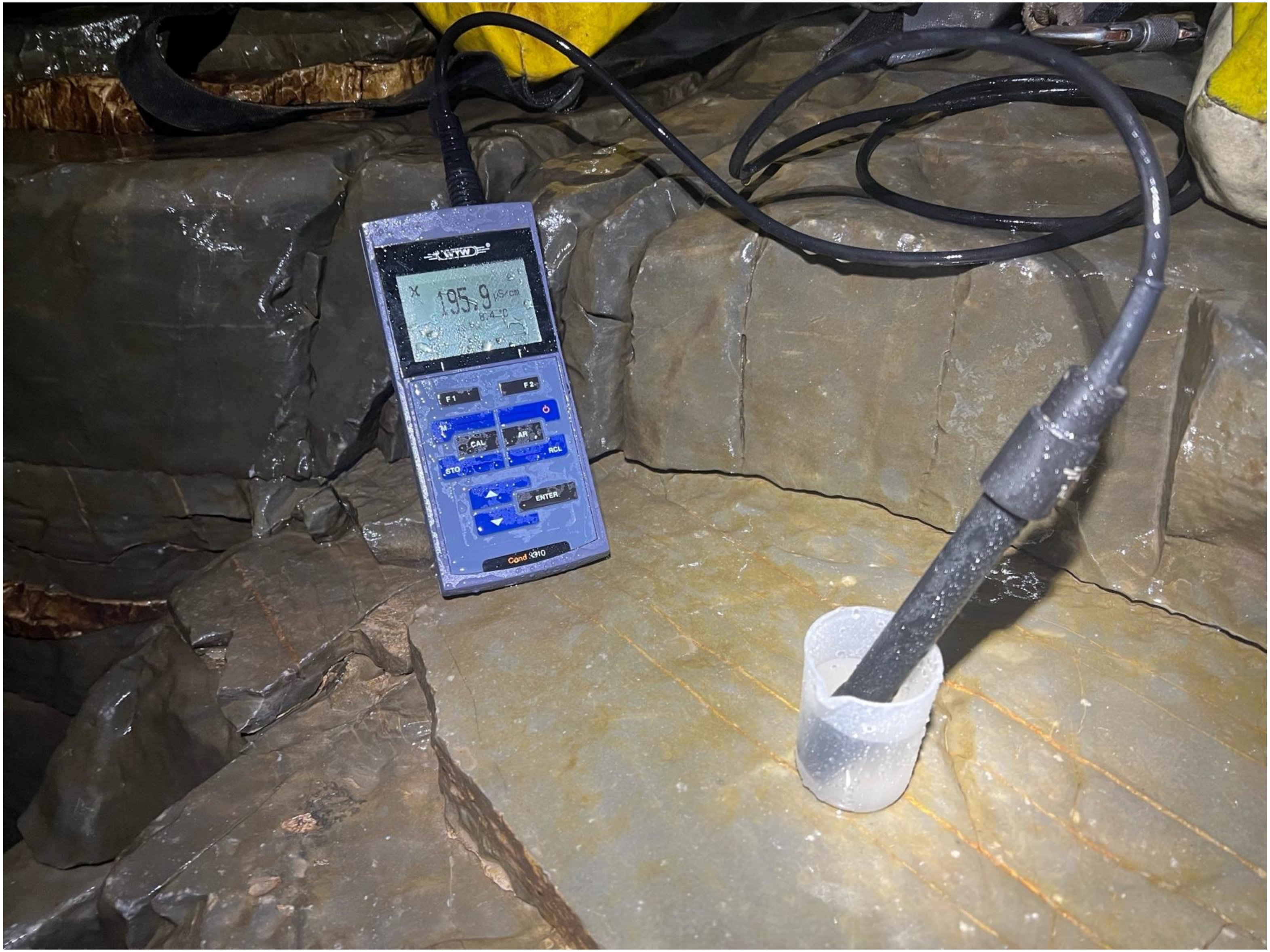

### gourgouthakas_-1100_siphon_5

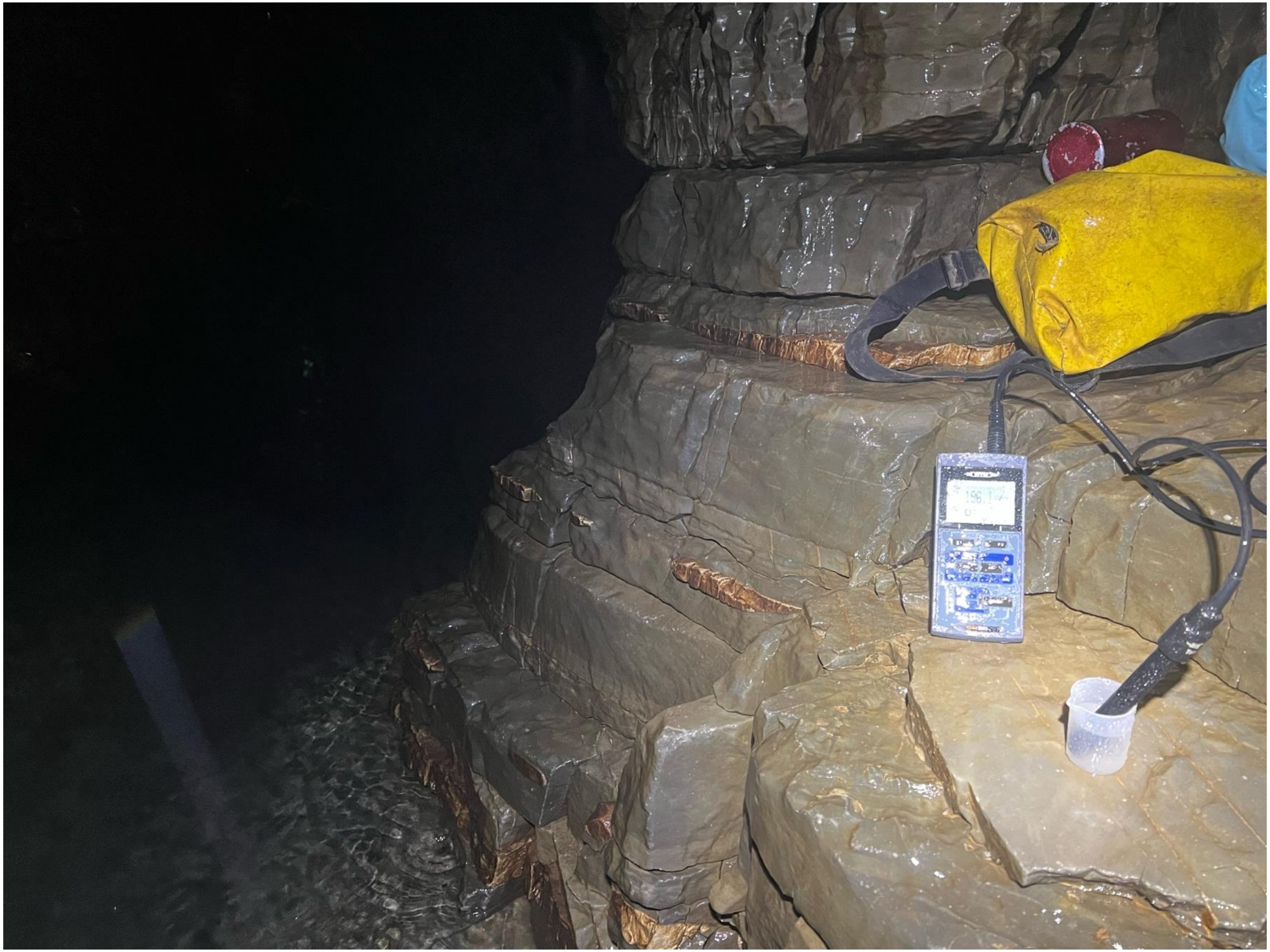

### gourgouthakas_-1100_siphon_6

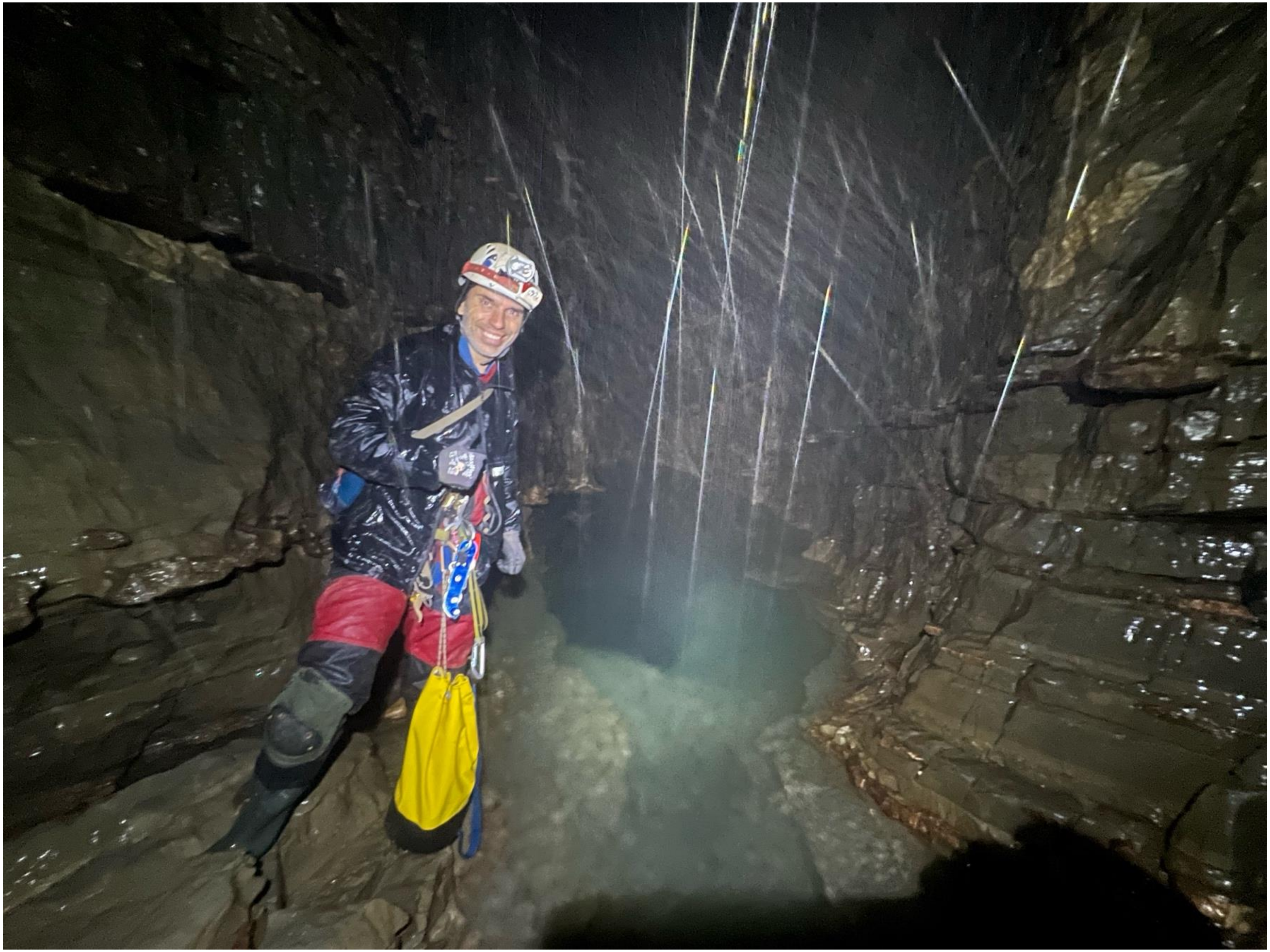

### gourgouthakas_-1100_siphon_7

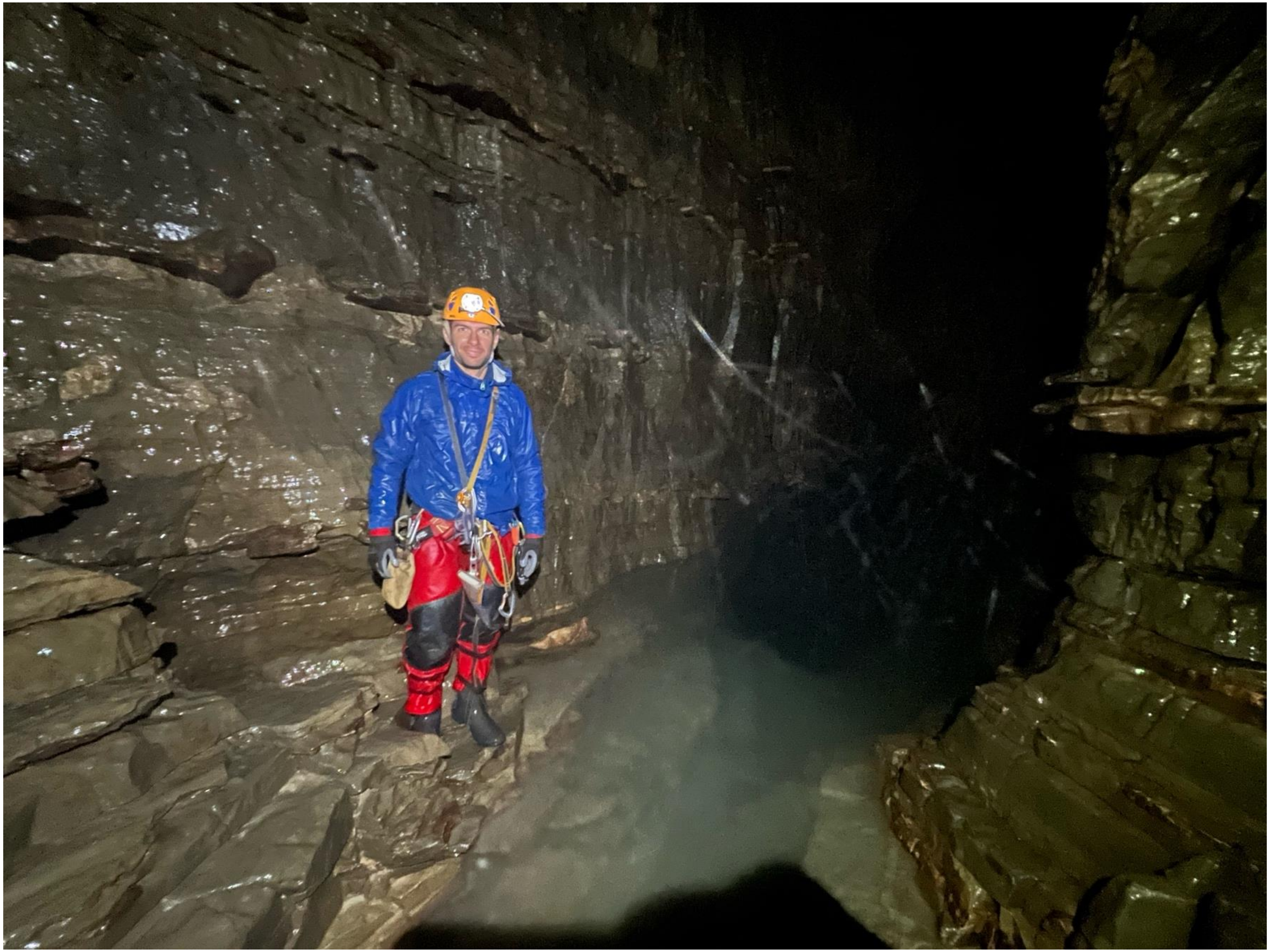

### gourgouthakas_samples_1

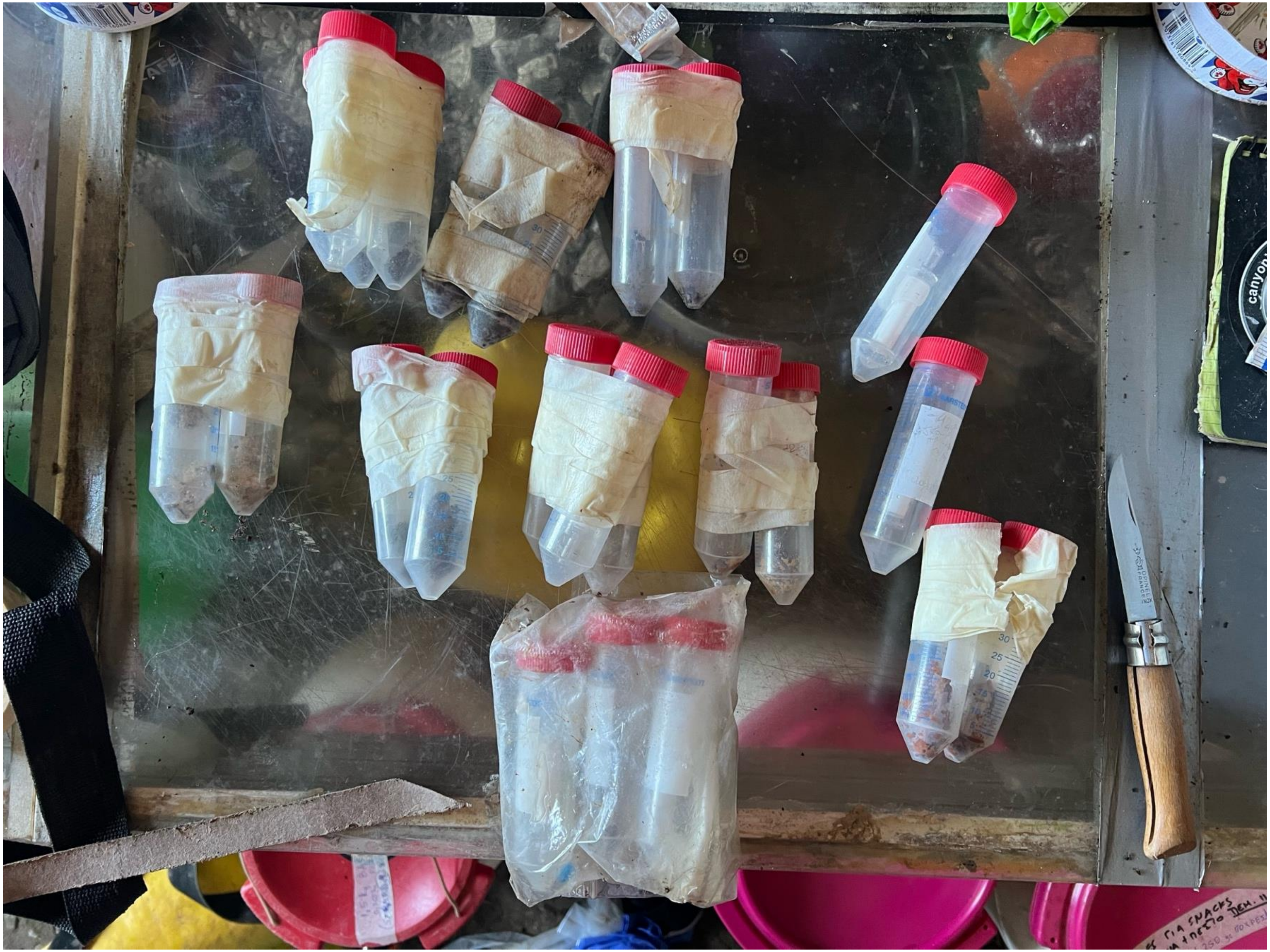

### gourgouthakas_samples_2

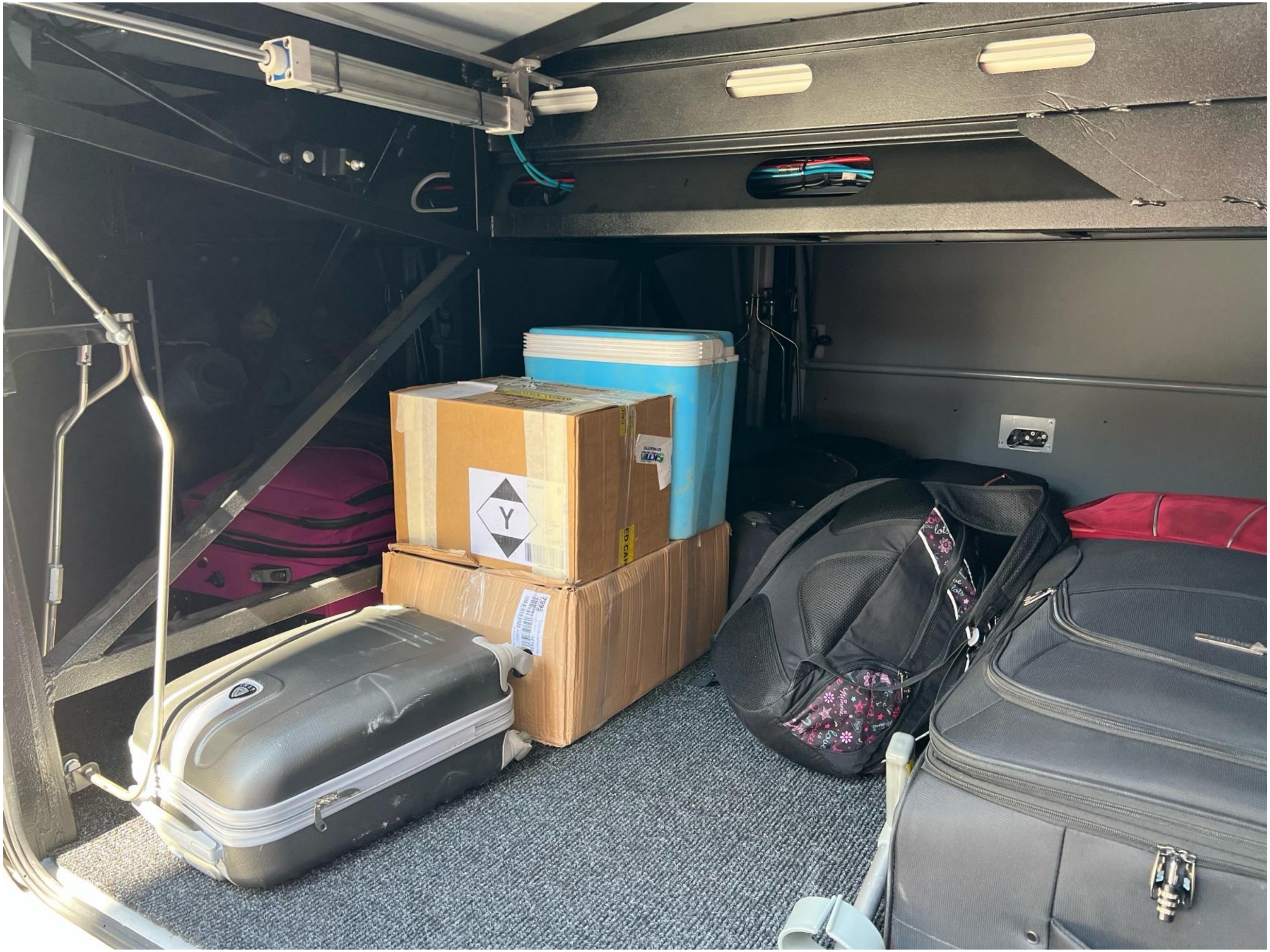

### gourgouthakas_scenery_1

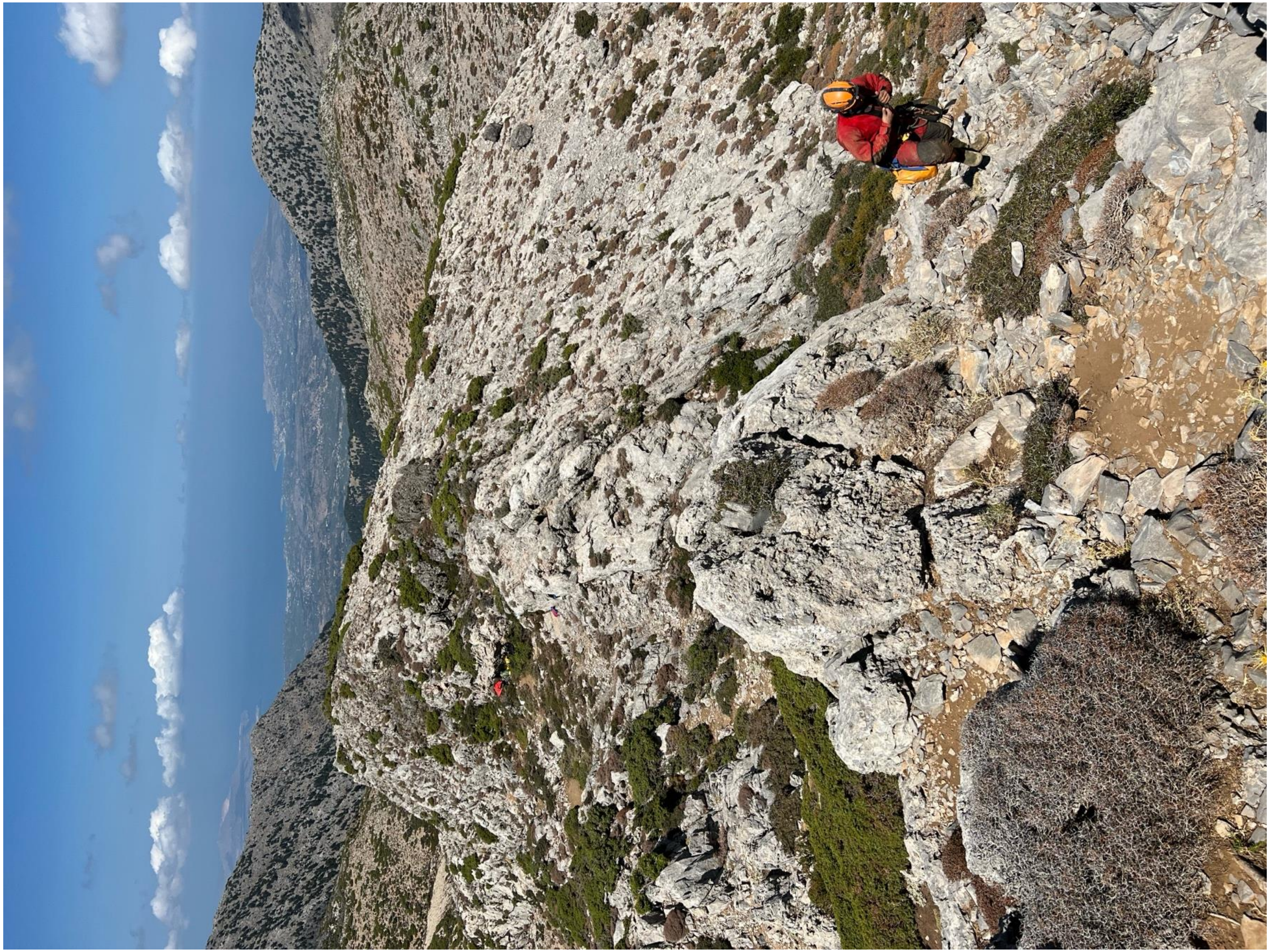

### gourgouthakas_scenery_2

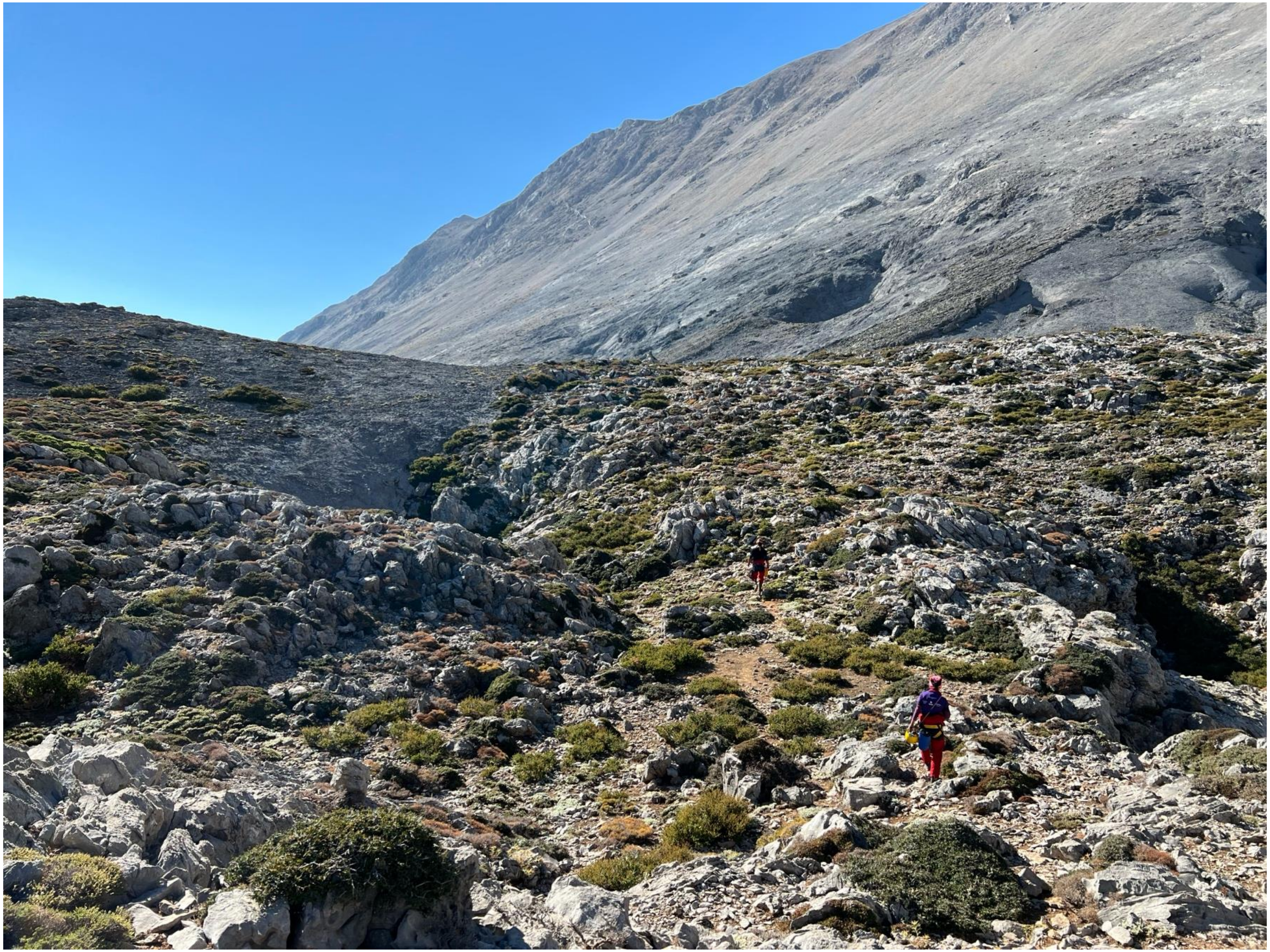

### gourgouthakas_scenery_3

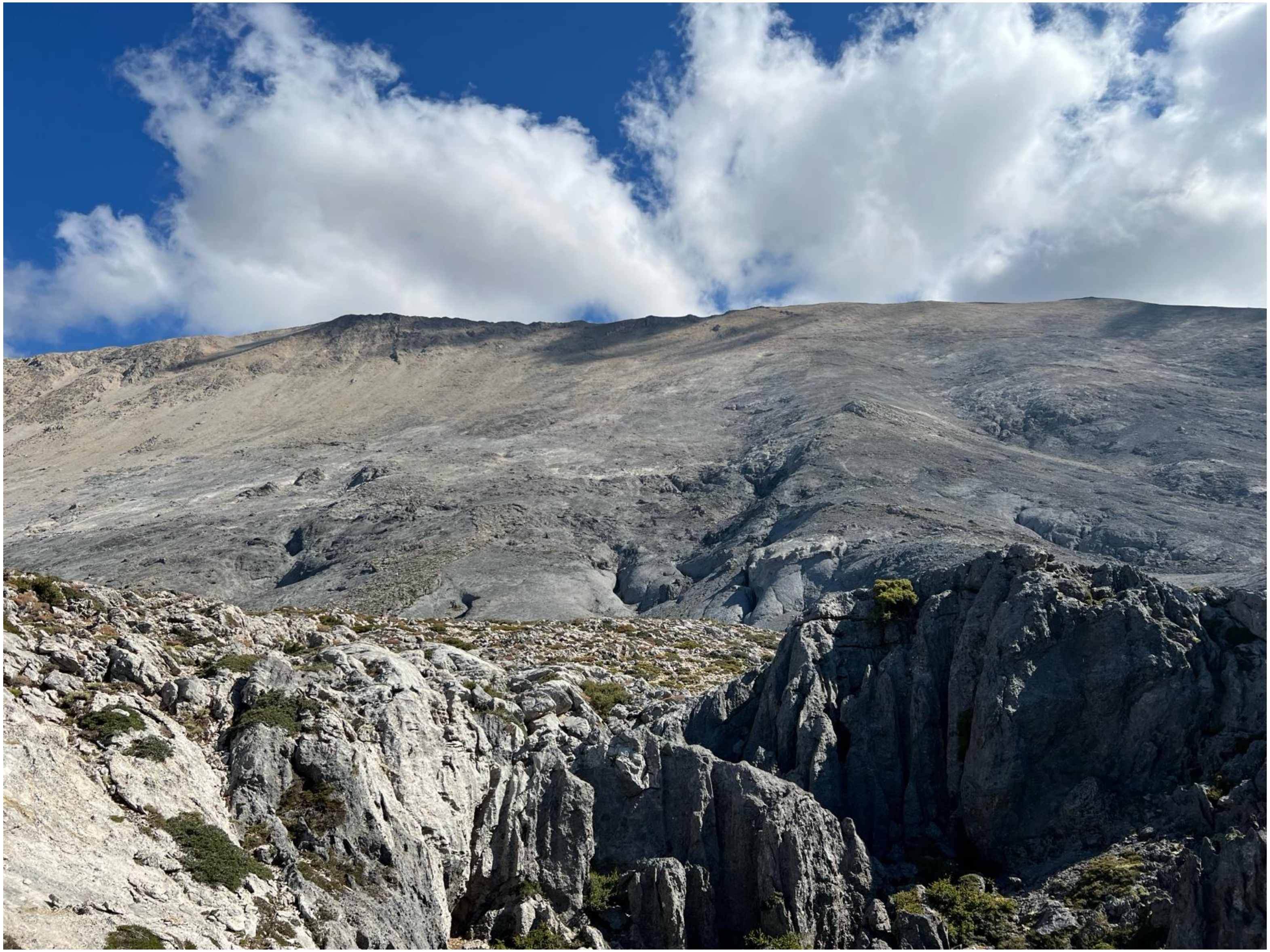

### gourgouthakas_scenery_4

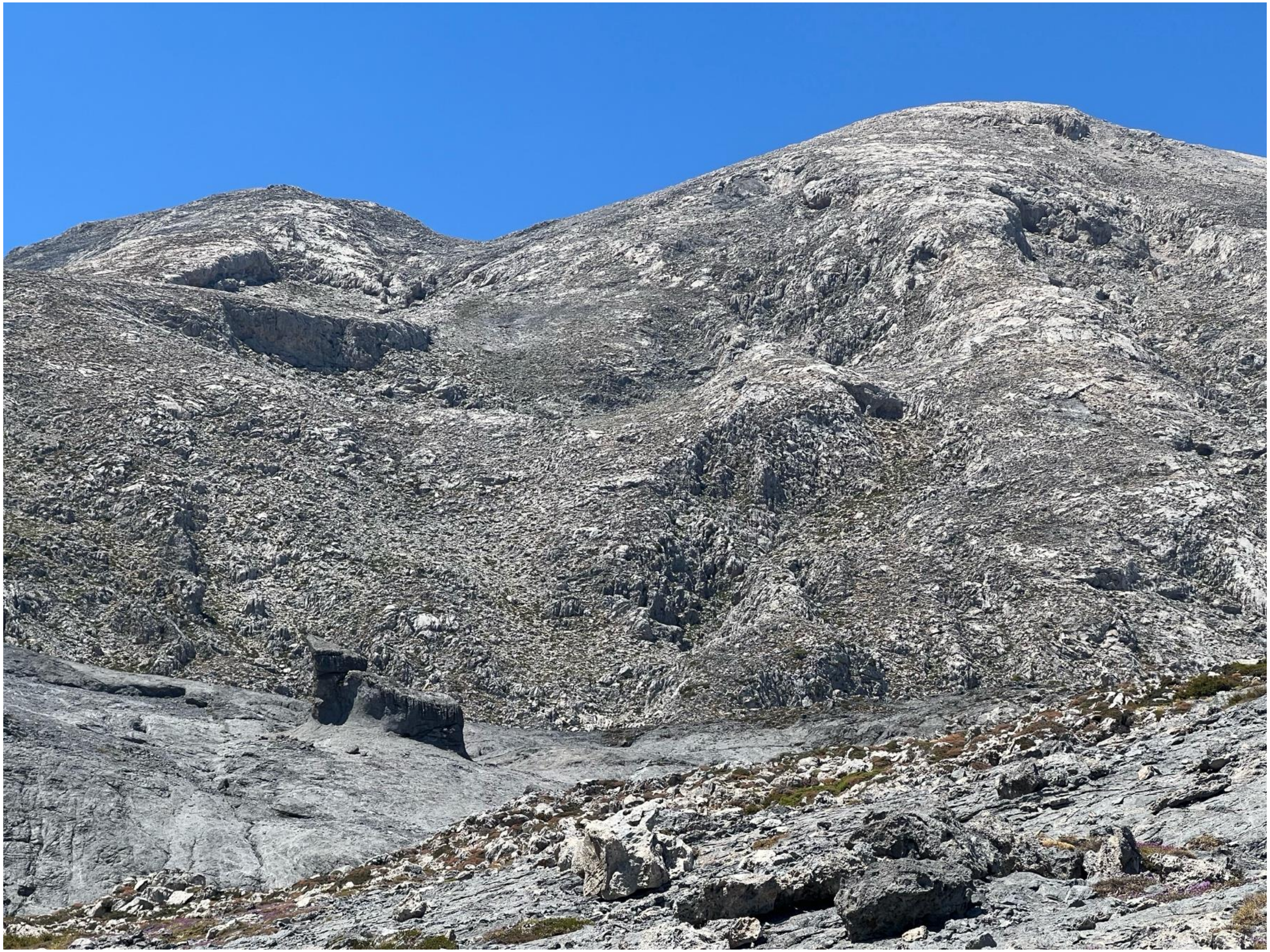
