## Supplementary Presentation 2 for "A Thousand Meters Deep: Vertical Profiling of the Subterranean Microbes of Gourgouthakas Cave"

frontiers

#### *Xanthomonas campestris* pv. *campestris* (Xcc)

**Xcc, NA**

- SRL1296
- SRL1288
- SRL927
- SRL926
- SRL1297
- **SRL1302**
- SRL938
- SRL929
- **SRL1304**
- SRL1303

24h

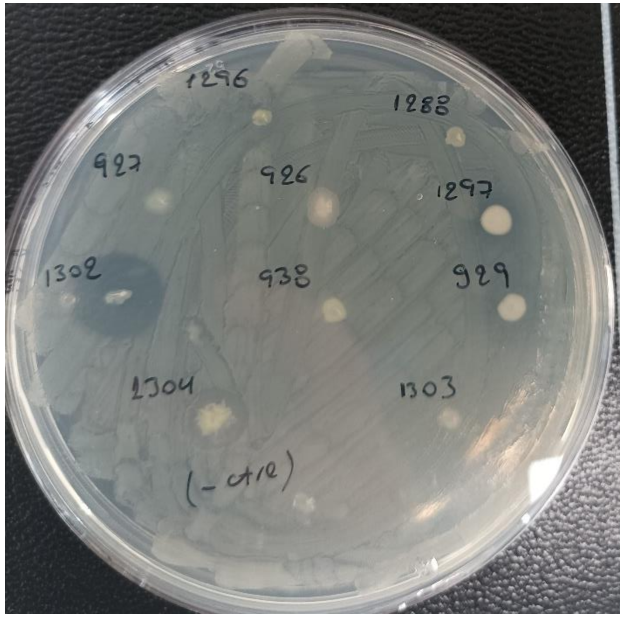


72h

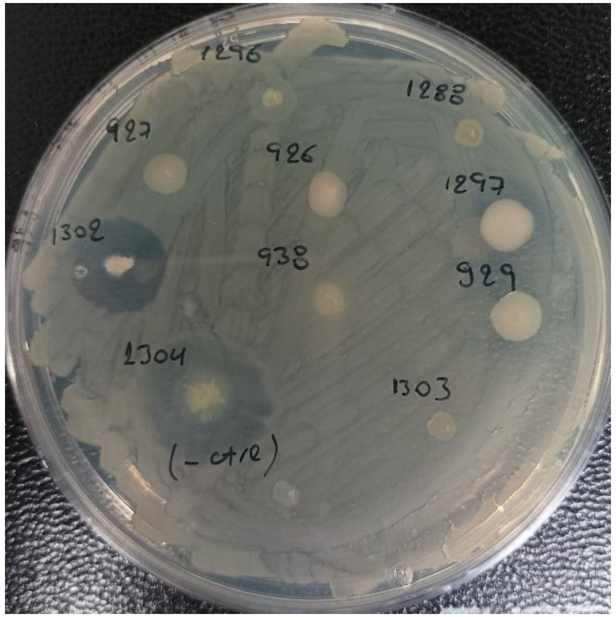


144h

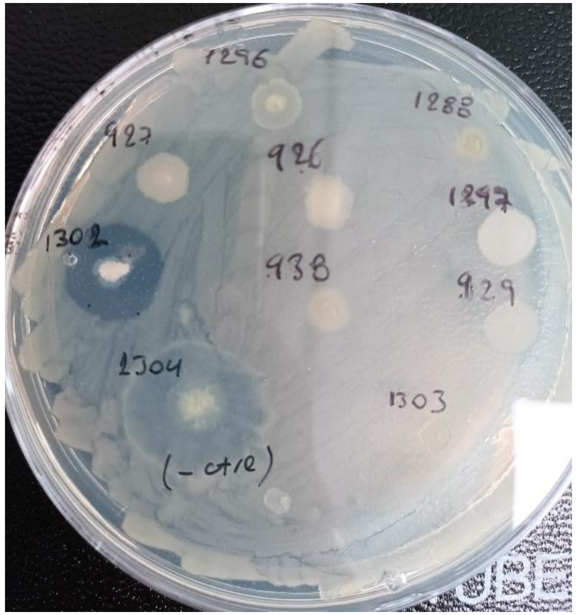


**Xcc, NA ½**

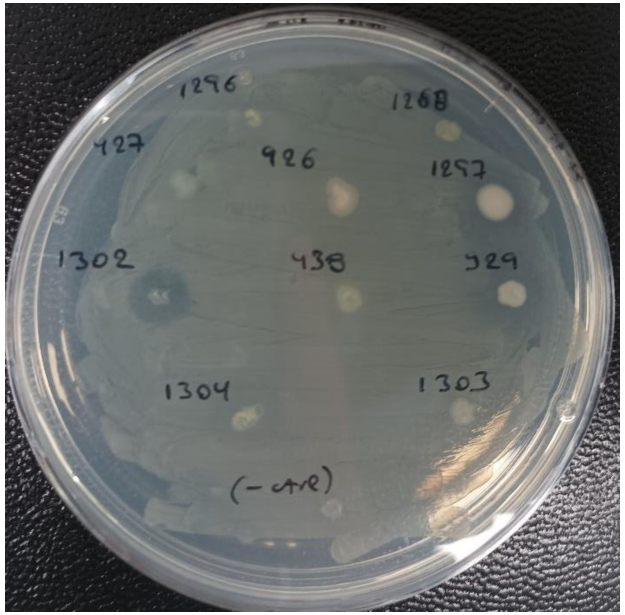


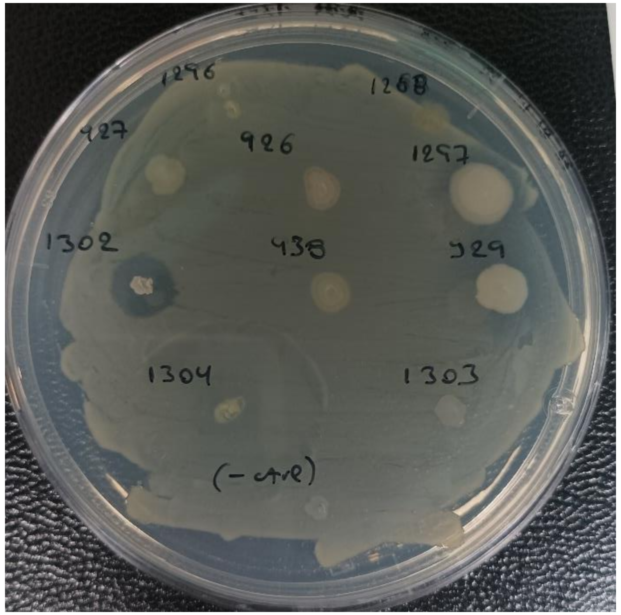

**Xcc, NA**

- SRL712
- SRL711
- SRL891
- SRL698
- SRL713
- SRL925
- SRL924
- SRL768
- SRL814

24h | 72h | 144h

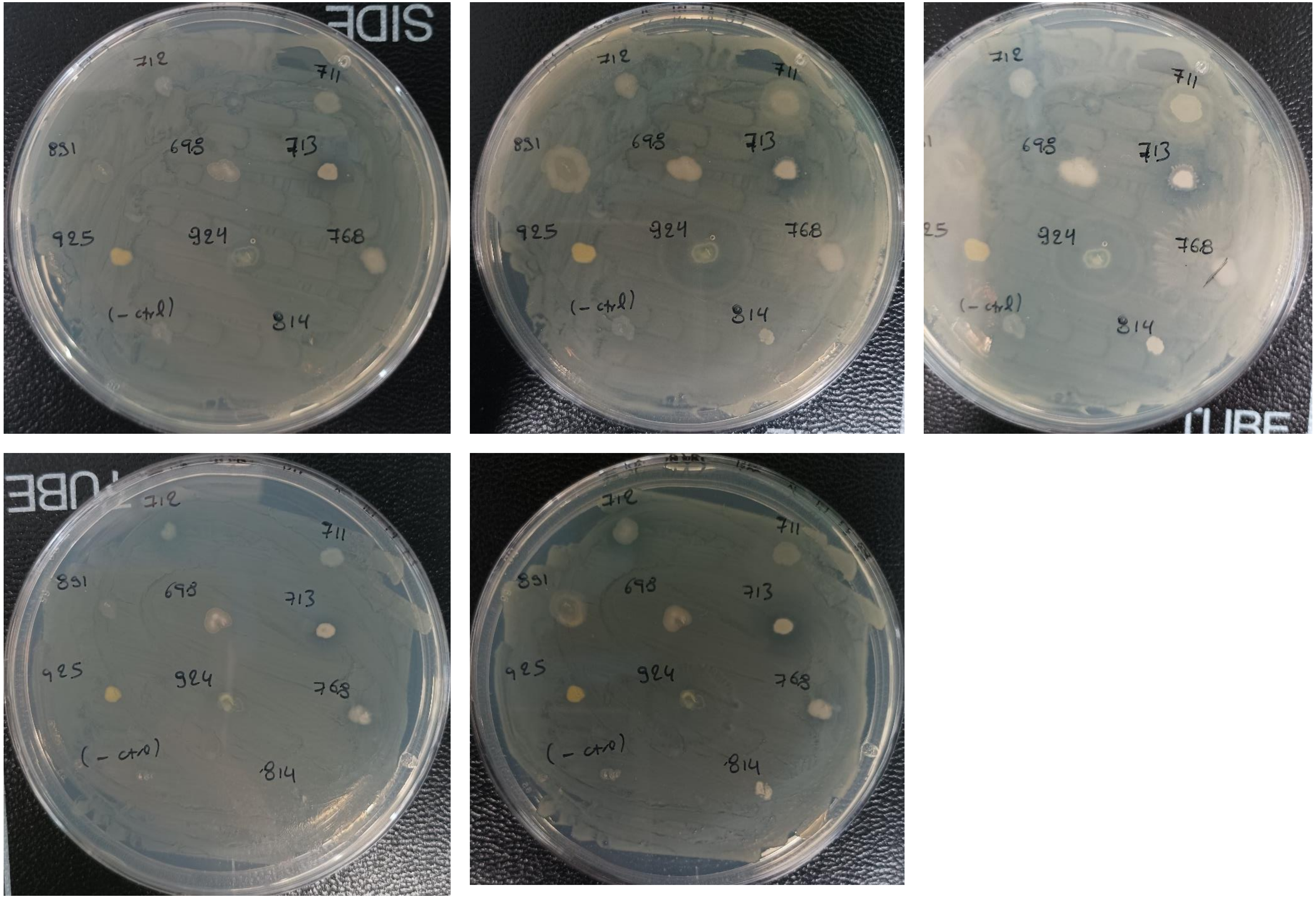


**Xcc, NA ½**

**Xcc, NA**

- SRL773
- **SRL769**
- **SRL771**
- SRL772
- SRL728
- SRL730
- SRL699
- **SRL726**
- SRL684

24h 72h

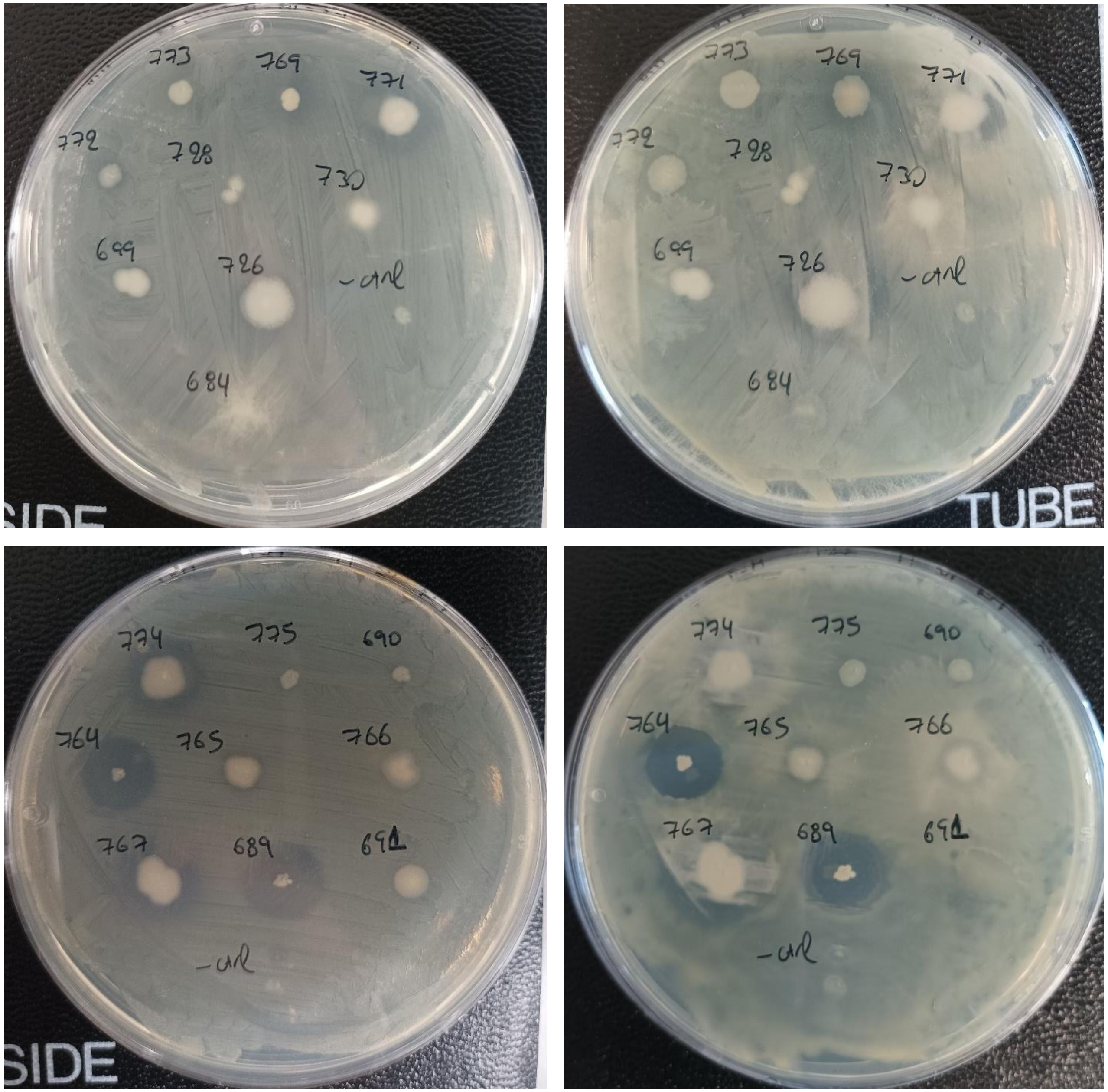


- **SRL774**
- SRL775
- SRL690
- **SRL764**
- **SRL765**
- SRL766
- SRL767
- **SRL689**
- SRL691

**Xcc, NA**

- **SRL820**
- **SRL821**
- SRL822
- **SRL823**
- SRL824
- SRL826
- SRL829
- SRL858
- SRL890
- SRL931

- SRL800
- SRL809
- **SRL810**
- **SRL811**
- SRL812
- SRL815
- **SRL816**
- SRL817
- **SRL818**

24h | 48h | 112h

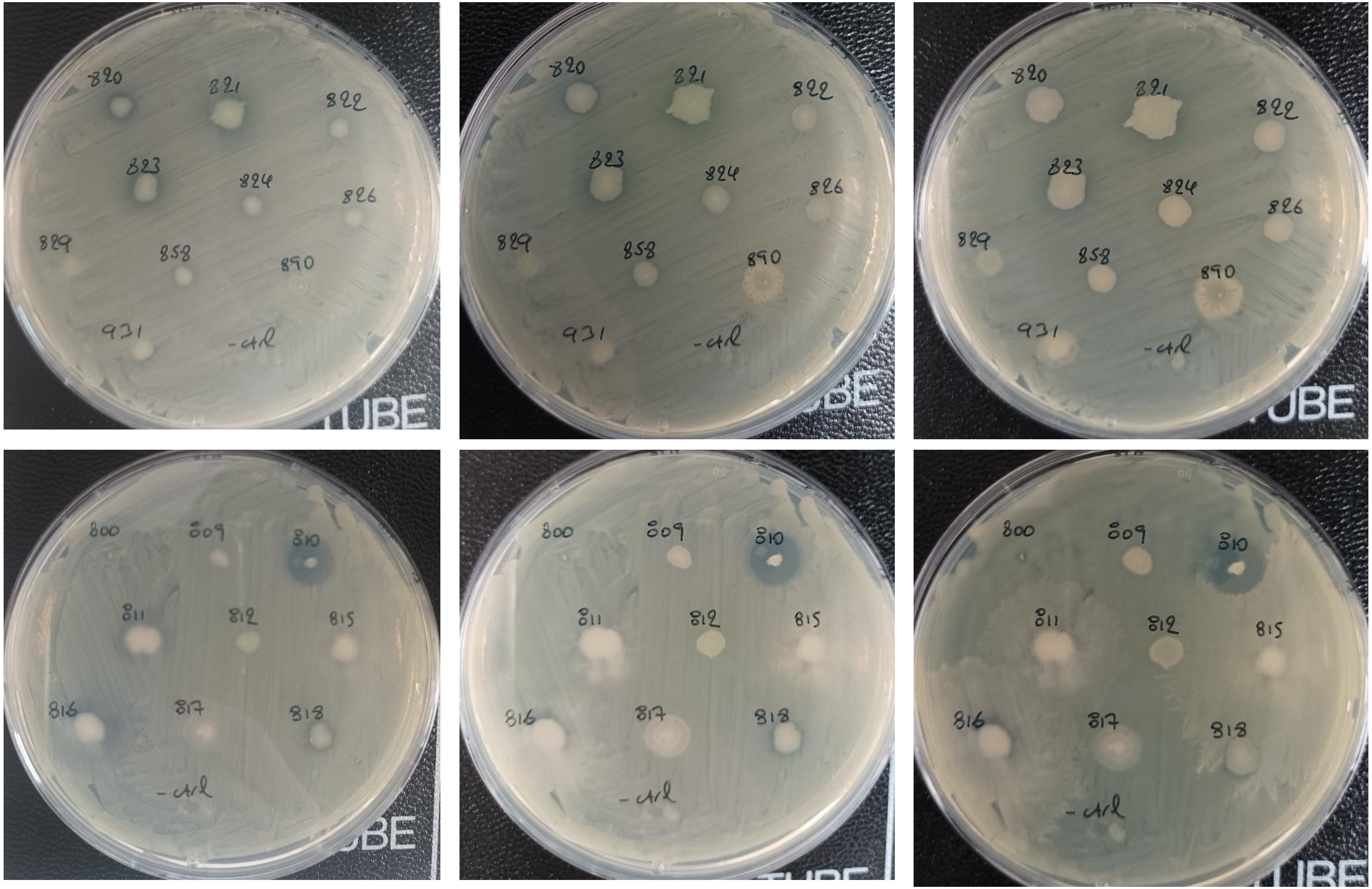

**Xcc, NA**

- SRL934
- SRL936
- **SRL937**
- SRL1281
- **SRL1282**
- SRL1283
- SRL1290
- SRL1291
- SRL1292

24h 48h 72h

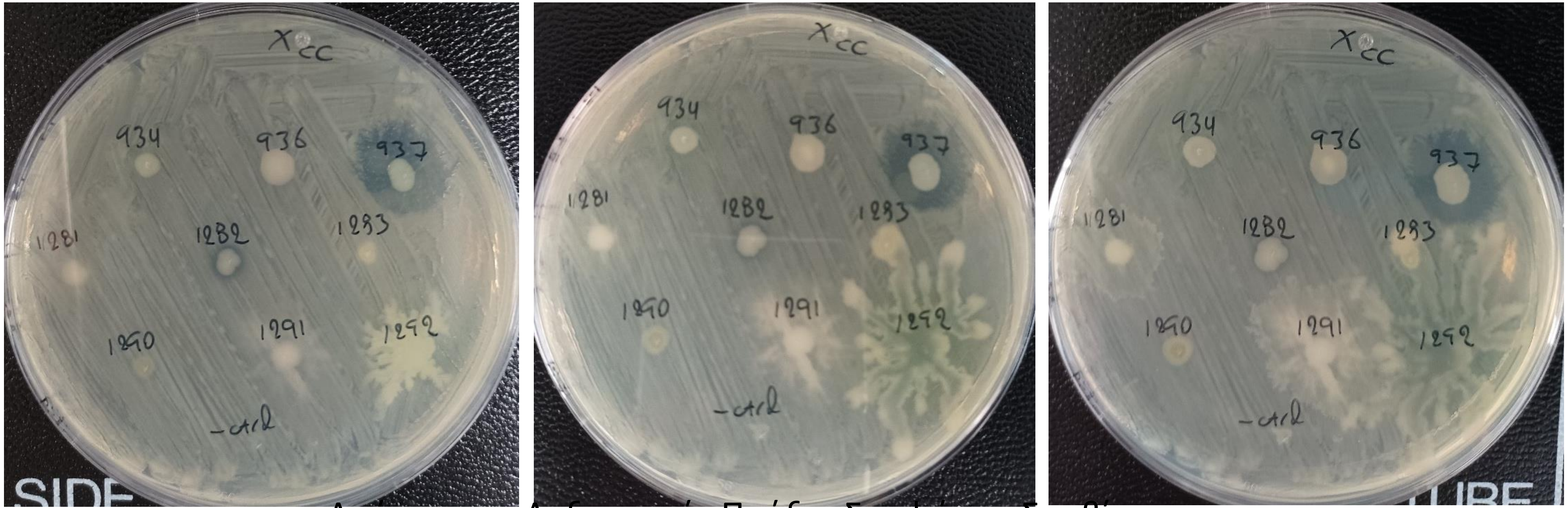


Απόσπασμα Ληξιαρχικής Πράξης Συμφώνου Συμβίωσης

- SRL1298
- SRL1299
- SRL1300
- **SRL1301**
- SRL1305
- SRL1307
- SRL1309
- SRL1310

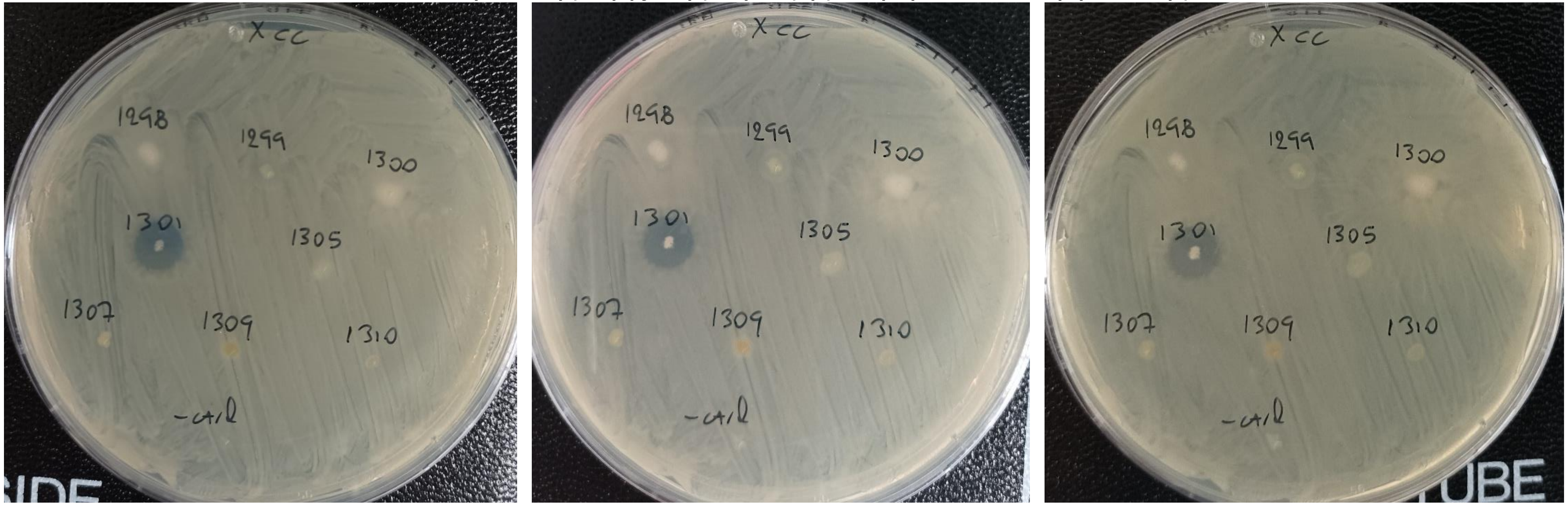

**Xcc, NA**

24h 48h 144h

- SRL780
- **SRL813**
- SRL930
- SRL939
- SRL1280
- SRL1289

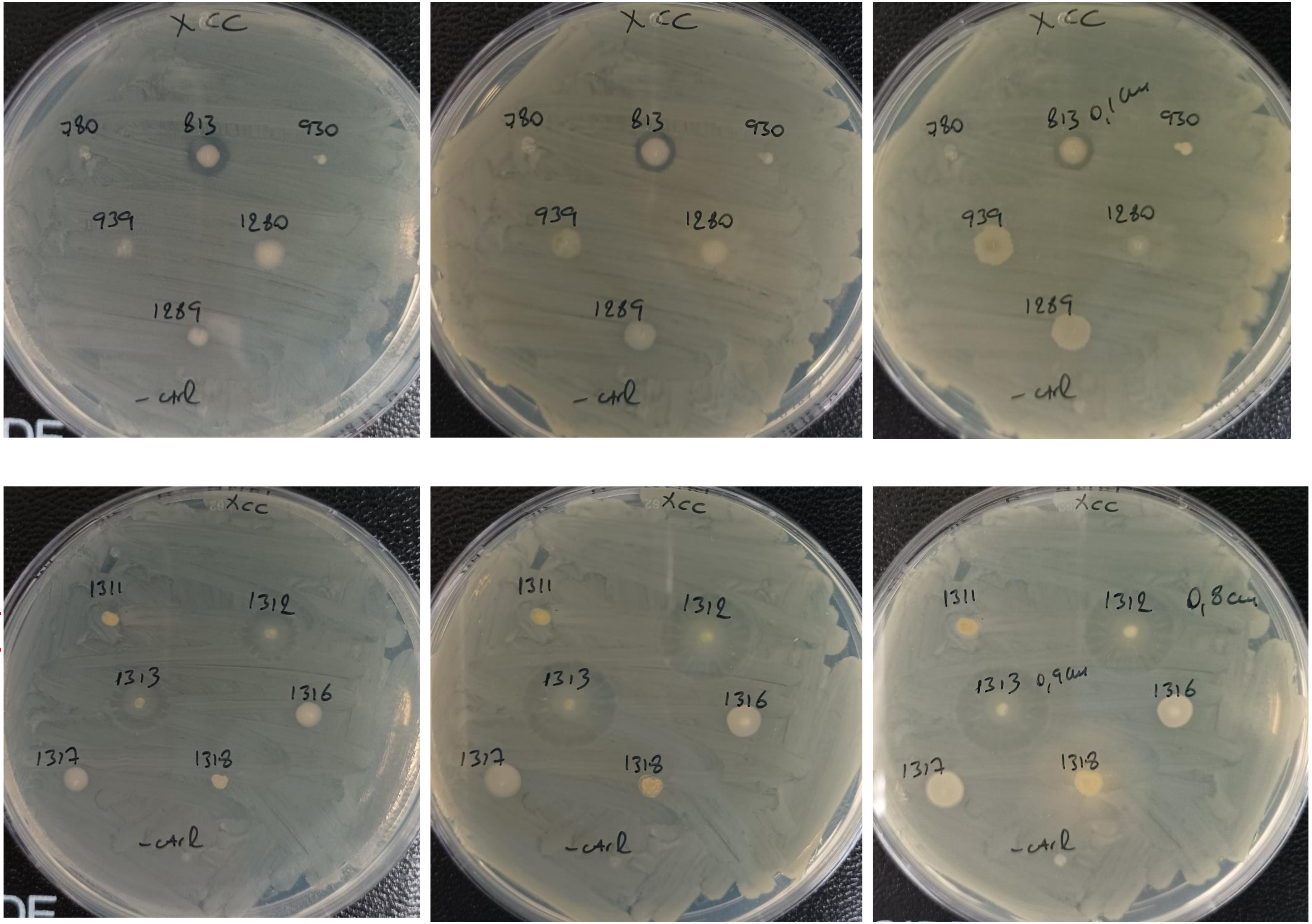


- SRL1311
- **SRL1312**
- **SRL1313**
- SRL1316
- SRL1317
- SRL1318

**Xcc, NA**

- SRL1346
- SRL1171
- SRL1120
- **SRL1336**
- SRL888
- SRL1325
- SRL1326
- SRL1058

24h 48h

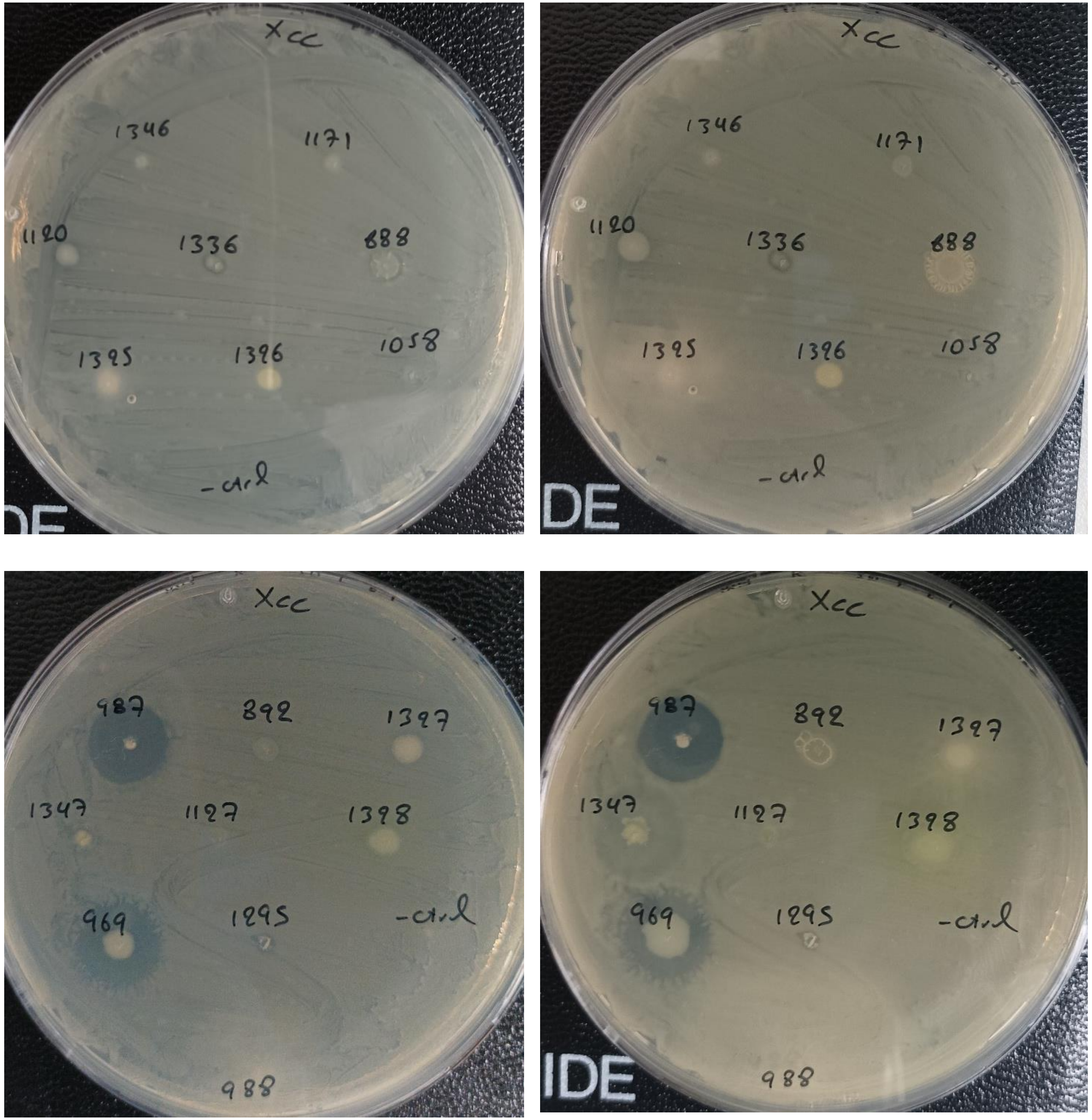


- **SRL987**
- SRL892
- SRL1327
- **SRL1347**
- SRL1127
- SRL1328
- **SRL969**
- SRL1295
- SRL988

**Xcc, NA**

24h | 48h | 96h

- SRL732
- SRL727
- SRL932
- **SRL979**
- SRL1024
- SRL1015
- SRL975
- SRL971
- SRL1323

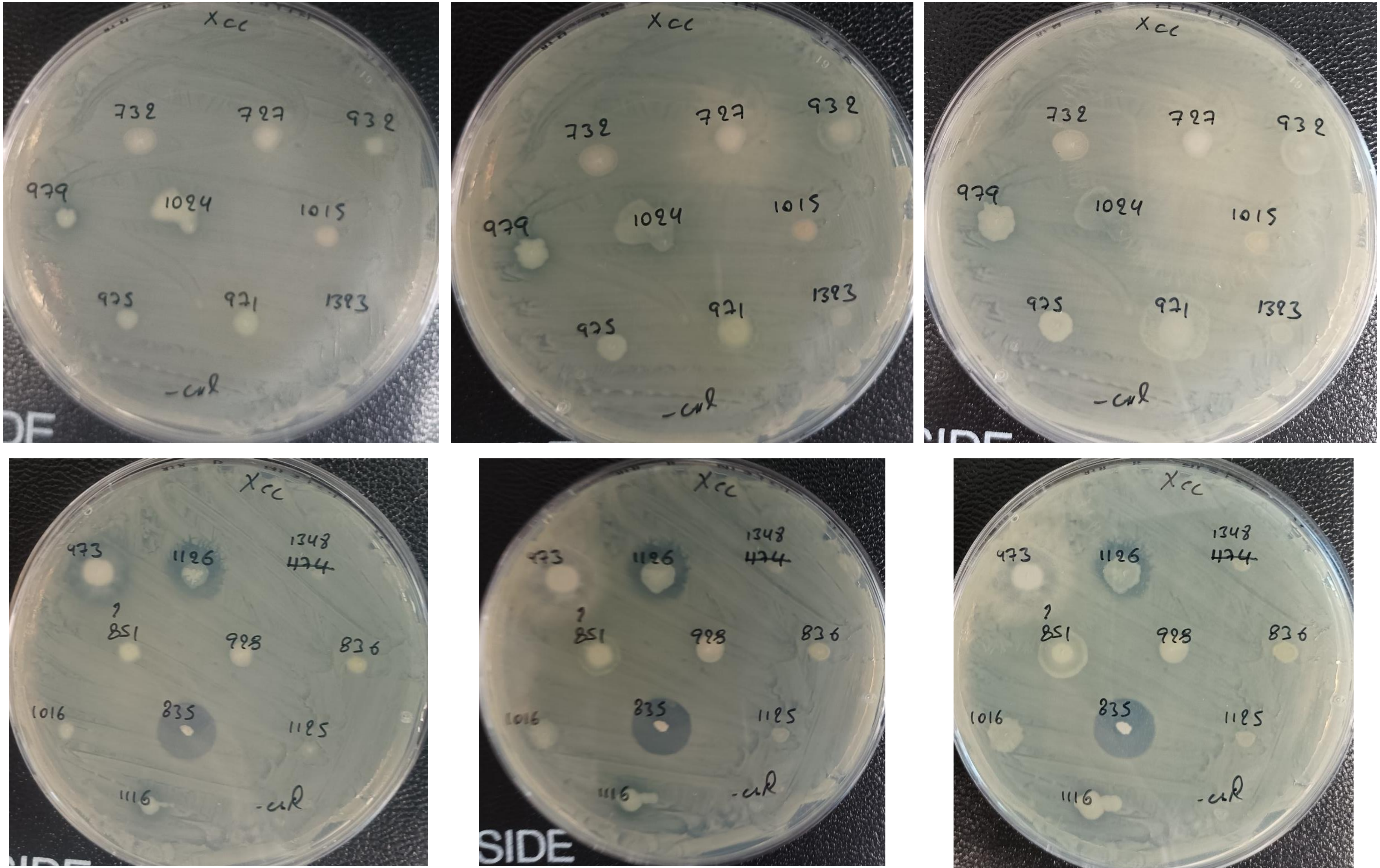


- **SRL973**
- **SRL1126**
- SRL1348
- SRL851
- SRL928
- SRL836
- SRL1016
- **SRL835**
- SRL1125
- SRL1116

**Xcc, NA**

- SRL1123
- SRL1114
- SRL1322
- SRL1333
- SRL1055
- **SRL778**
- SRL1343
- SRL1175
- SRL1113

- **SRL1342**
- SRL1124
- **SRL1115**
- **SRL982**
- **SRL983**
- SRL889
- SRL1049
- SRL1308
- SRL838
- **SRL837**

24h | 48h | 96h

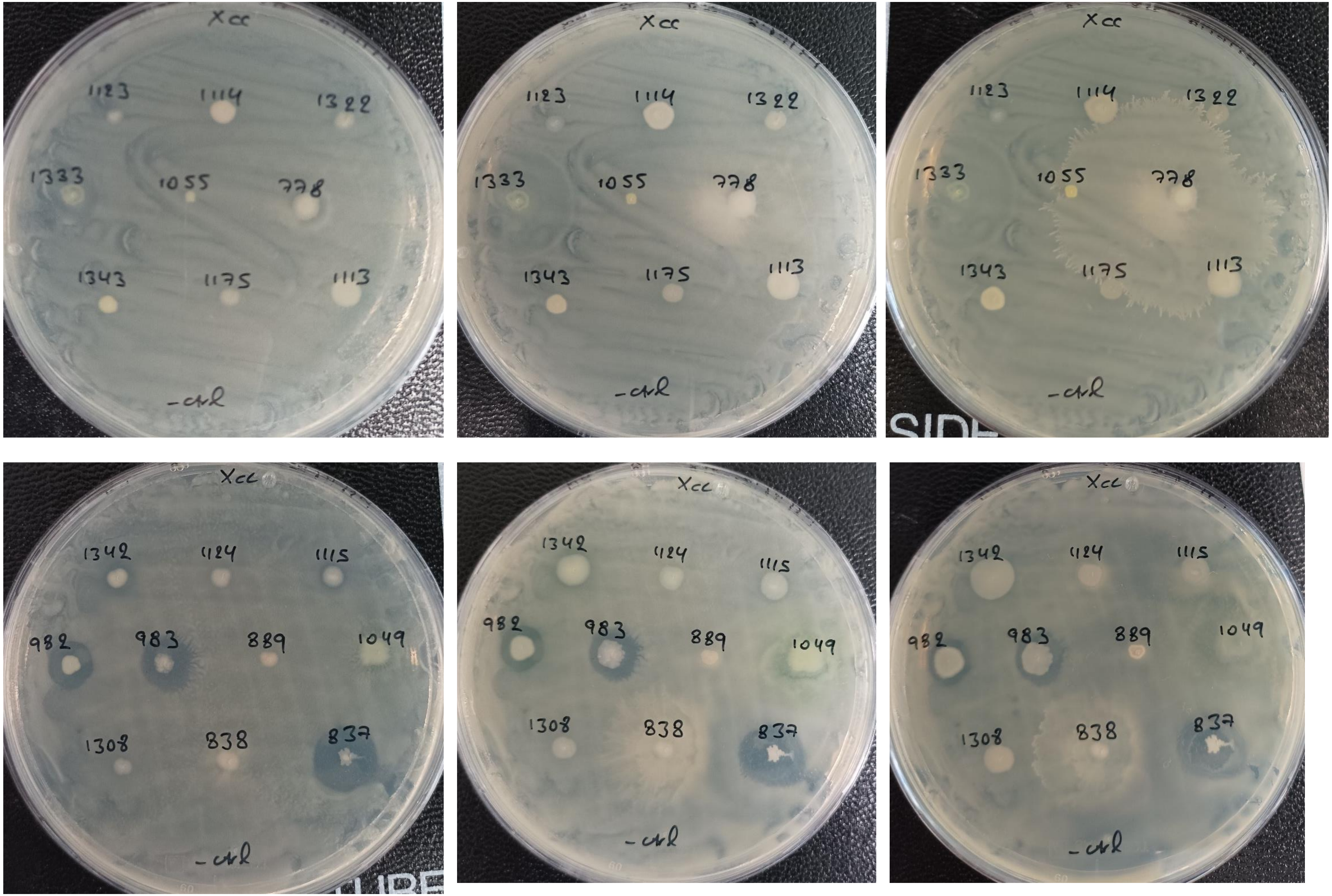

**Xcc, NA**

- **SRL1324**
- **SRL986**
- SRL1334
- **SRL871**
- SRL990
- SRL1344
- SRL1121
- SRL1345
- **SRL879**
- **SRL917**

- SRL874
- SRL981
- **SRL974**
- SRL976
- SRL1057
- SRL1048
- SRL729
- SRL834

24h | 48h | 96h

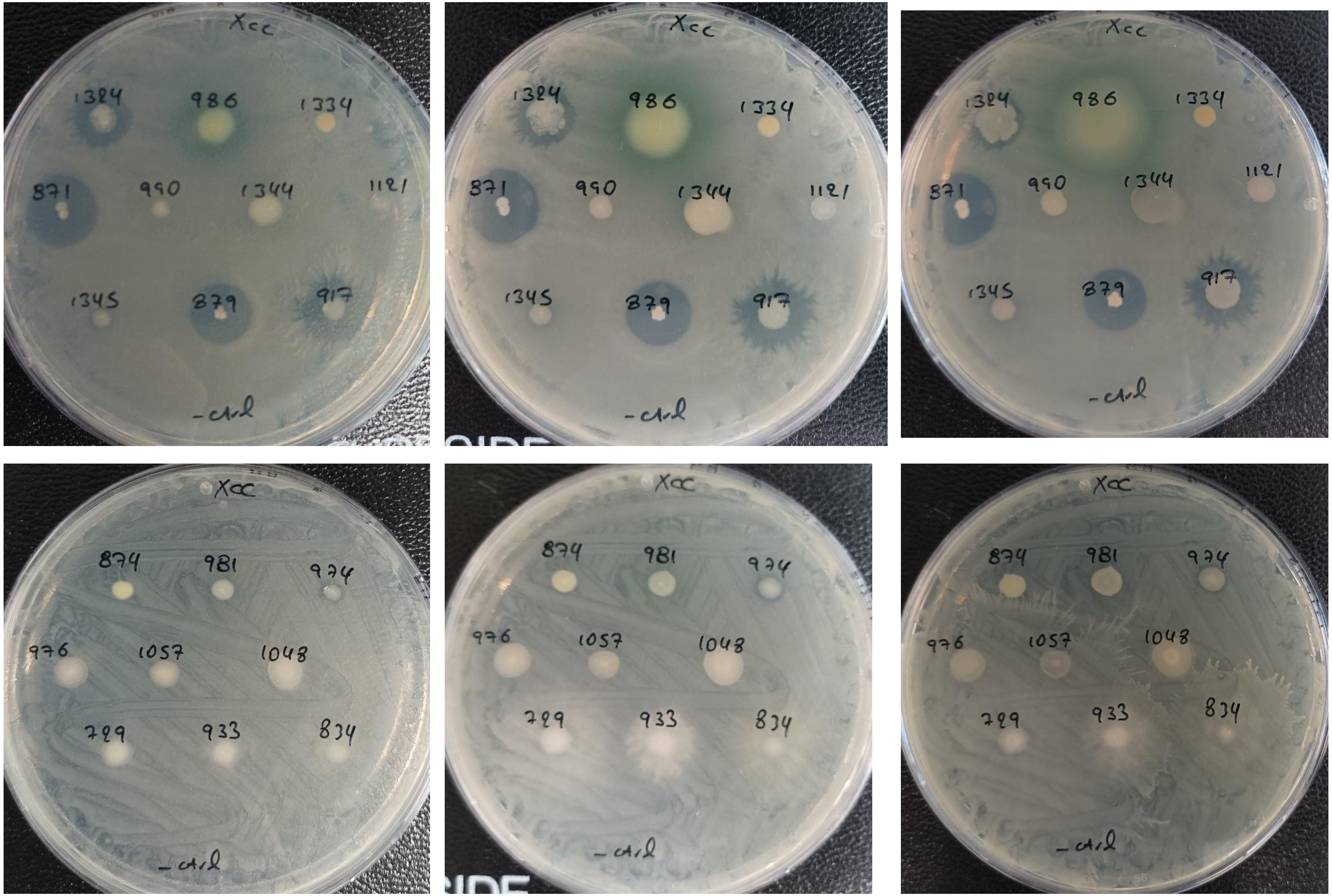

***Paracidovorax citrulli* (Pc)**

**Pc, NA**

- SRL1296
- SRL1288
- SRL927
- SRL926
- SRL1297
- **SRL1302**
- SRL938
- SRL929
- SRL1304
- SRL1303

72h

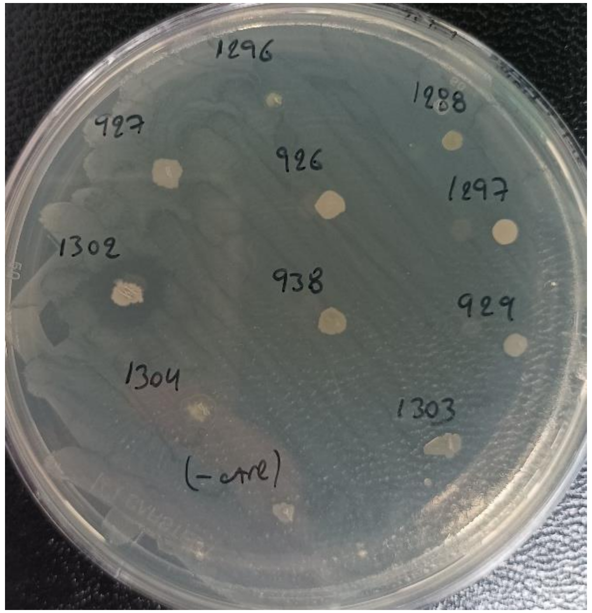


**Pc, NA ½**

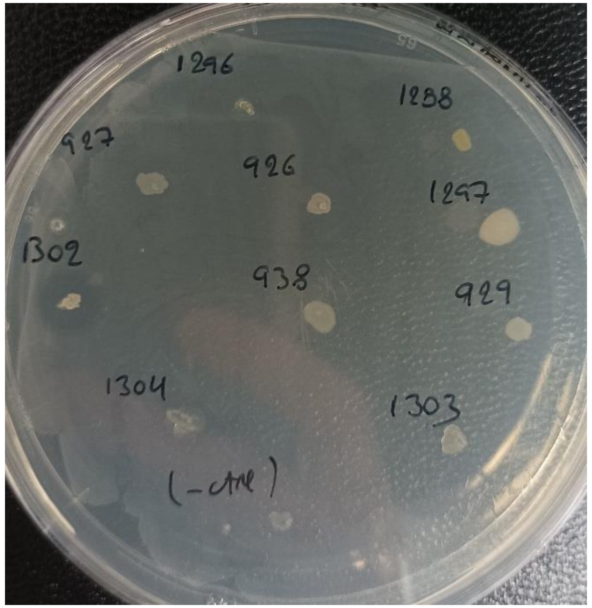


**Pc, NA**

- SRL712
- SRL711
- SRL891
- SRL698
- SRL713
- SRL925
- SRL924
- SRL768
- SRL814

72h

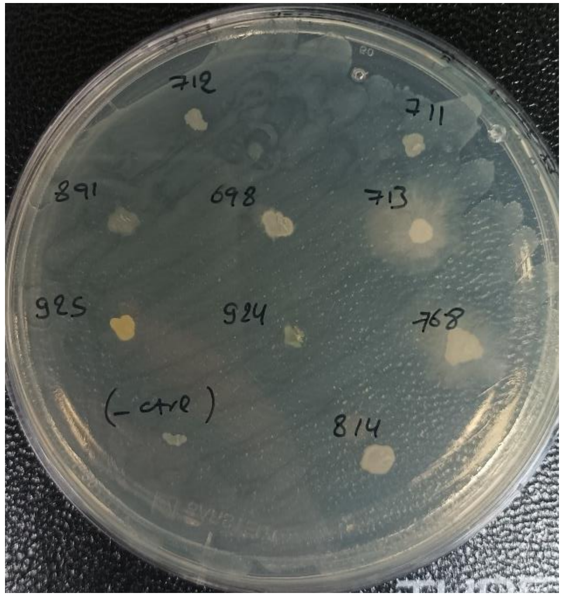


**Pc, NA ½**

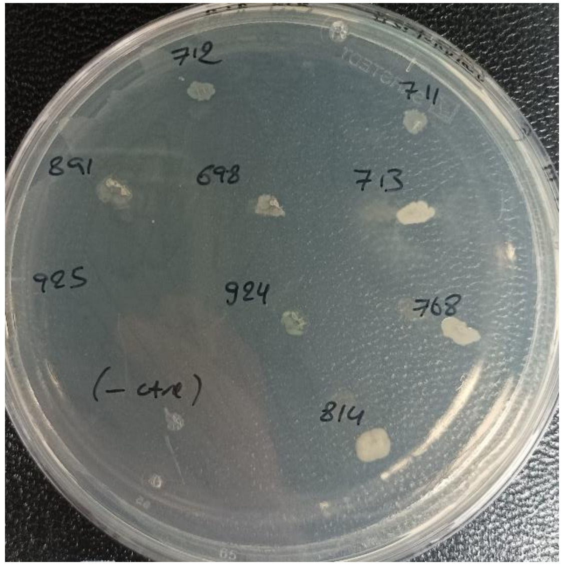

**Pc, NA**

- **SRL773**
- **SRL769**
- SRL771
- **SRL772**
- SRL728
- SRL730
- SRL699
- **SRL726**
- SRL684

- SRL774
- SRL775
- SRL690
- **SRL764**
- SRL765
- SRL766
- SRL767
- **SRL689**
- SRL691

24h 72h

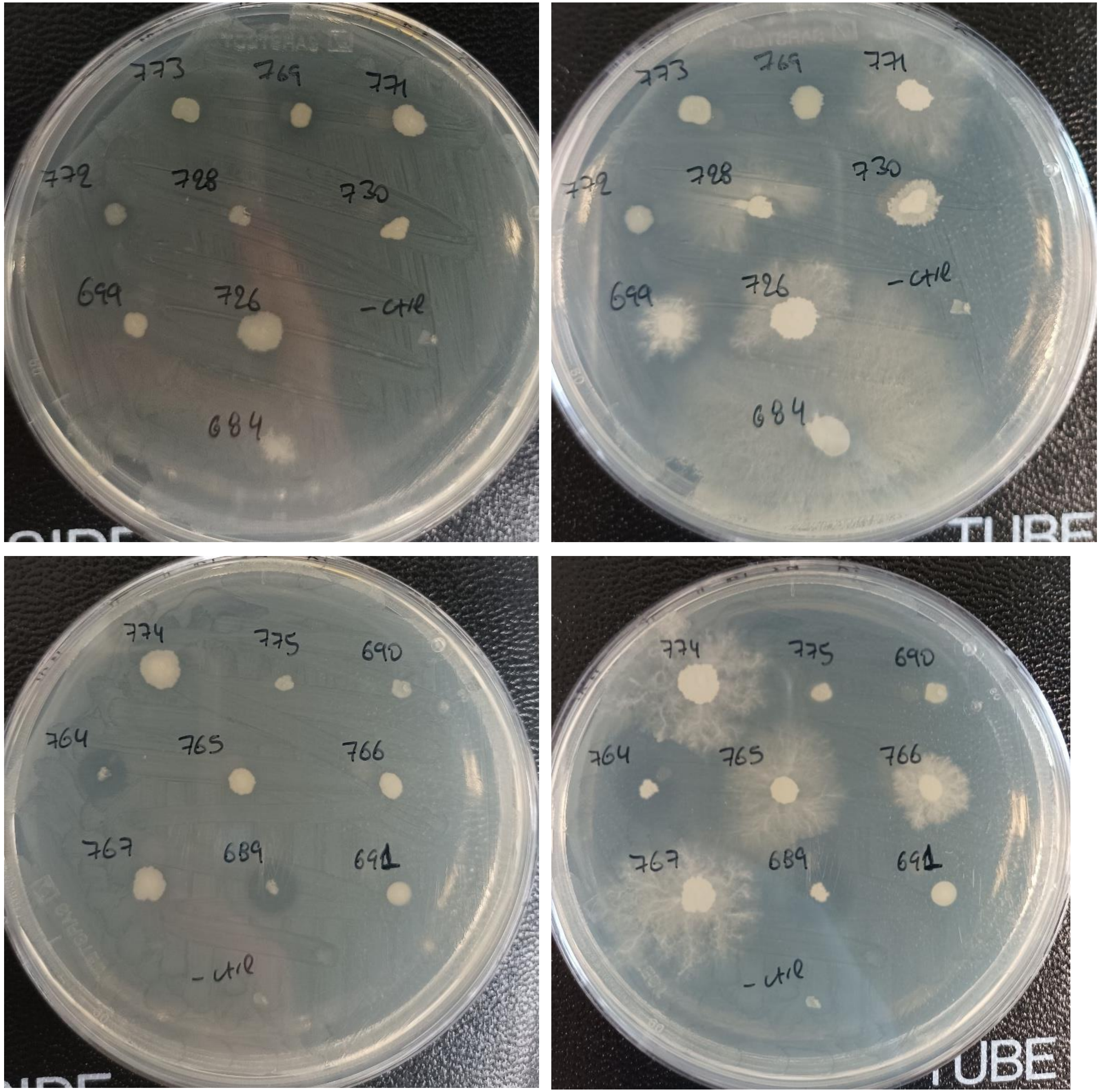

**Pc, NA**

24h 48h 112h

- SRL820
- SRL821
- SRL822
- SRL823
- SRL824
- SRL826
- SRL829
- SRL858
- SRL890
- SRL931

- SRL800
- SRL809
- SRL810
- SRL811
- SRL812
- SRL815
- SRL816
- SRL817
- SRL818

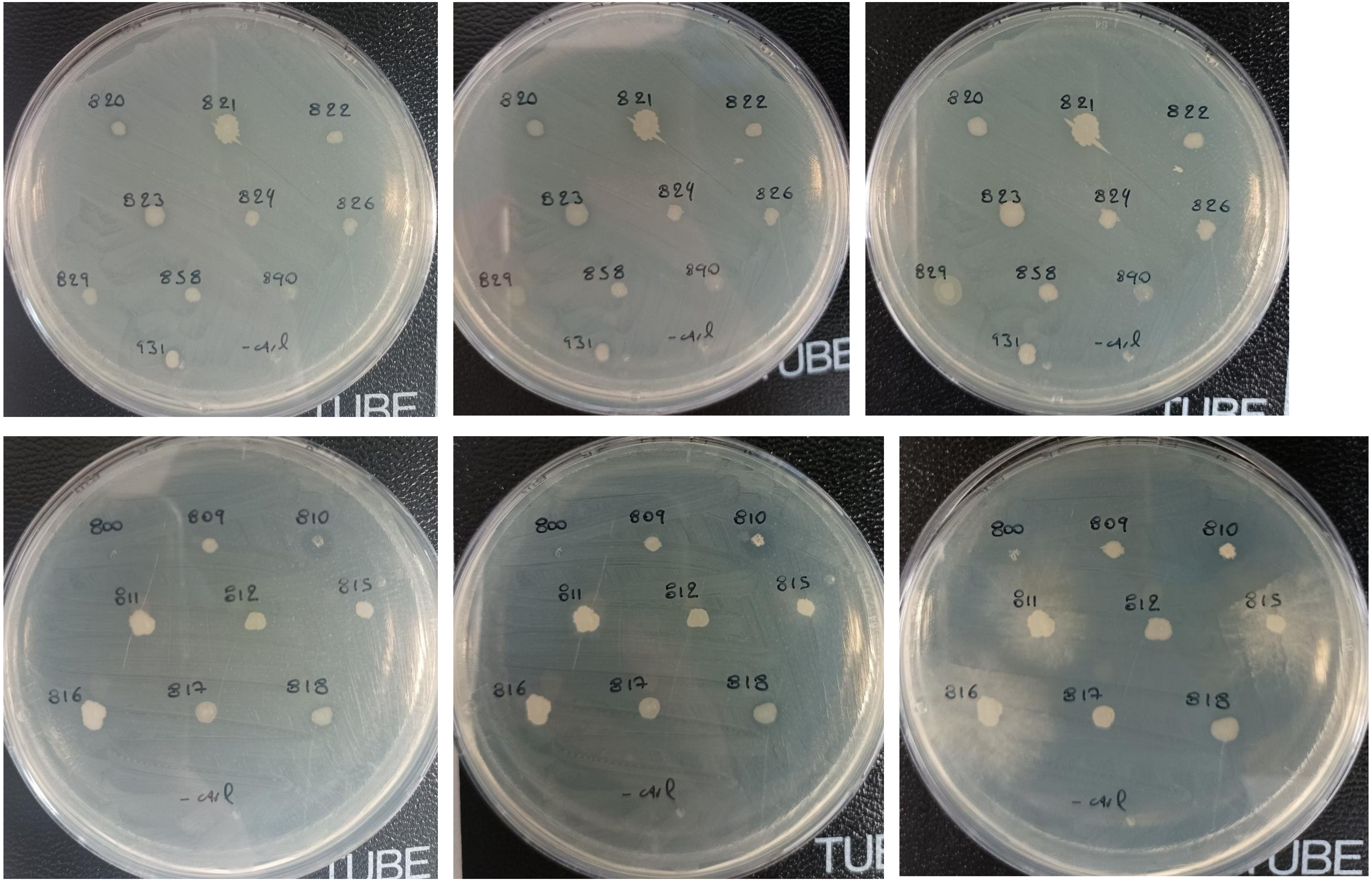

**Pc, NA**

- SRL934
- SRL936
- **SRL937**
- SRL1281
- **SRL1282**
- SRL1283
- SRL1290
- SRL1291
- SRL1292

- SRL1298
- SRL1299
- SRL1300
- **SRL1301**
- SRL1305
- SRL1307
- SRL1309
- SRL1310

24h | 48h | 72h

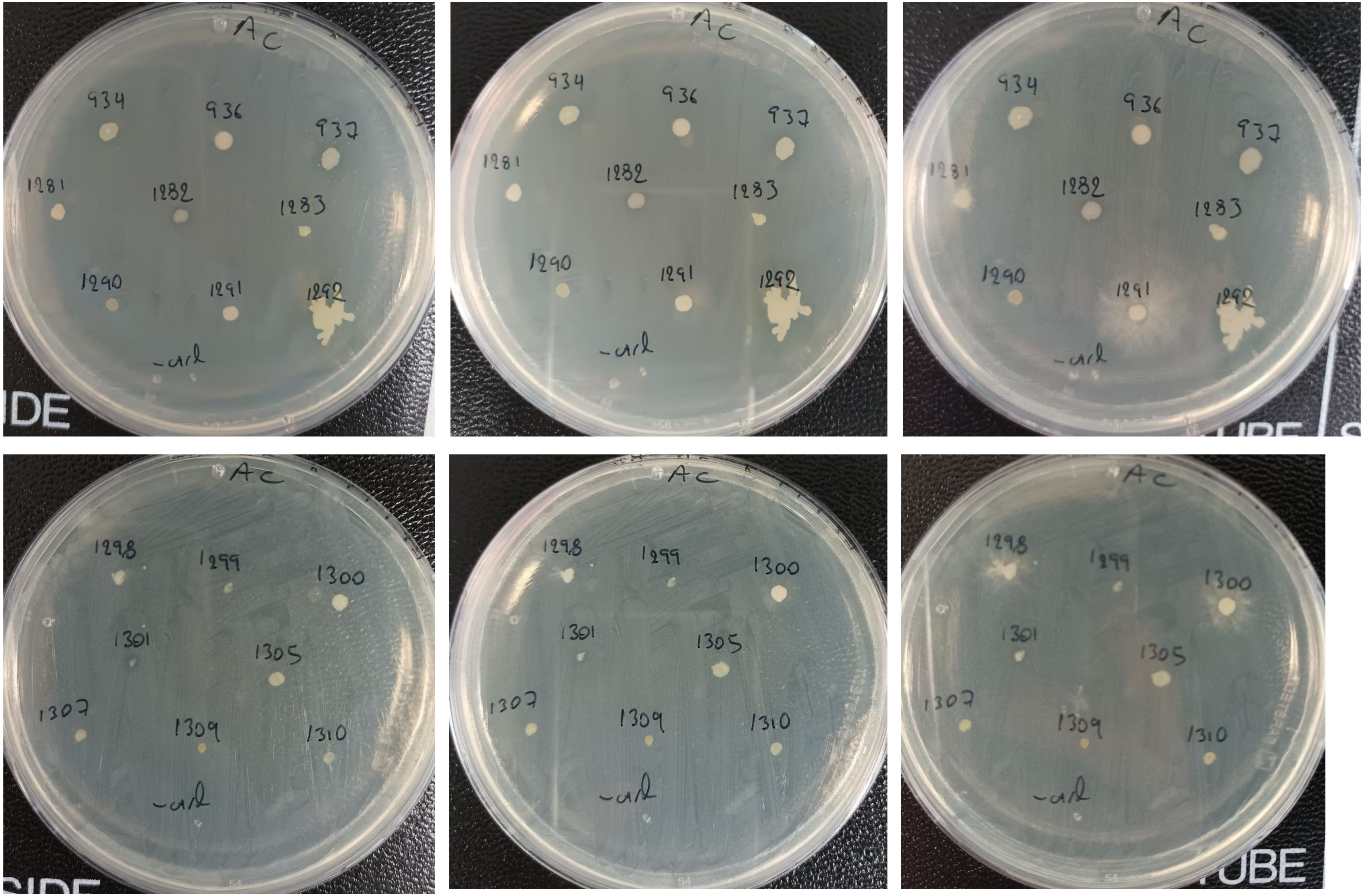

**Pc, NA**

- SRL780
- SRL813
- SRL930
- SRL939
- SRL1280
- SRL1289

- SRL1311
- SRL1312
- SRL1313
- SRL1316
- SRL1317
- SRL1318

24h 48h 144h

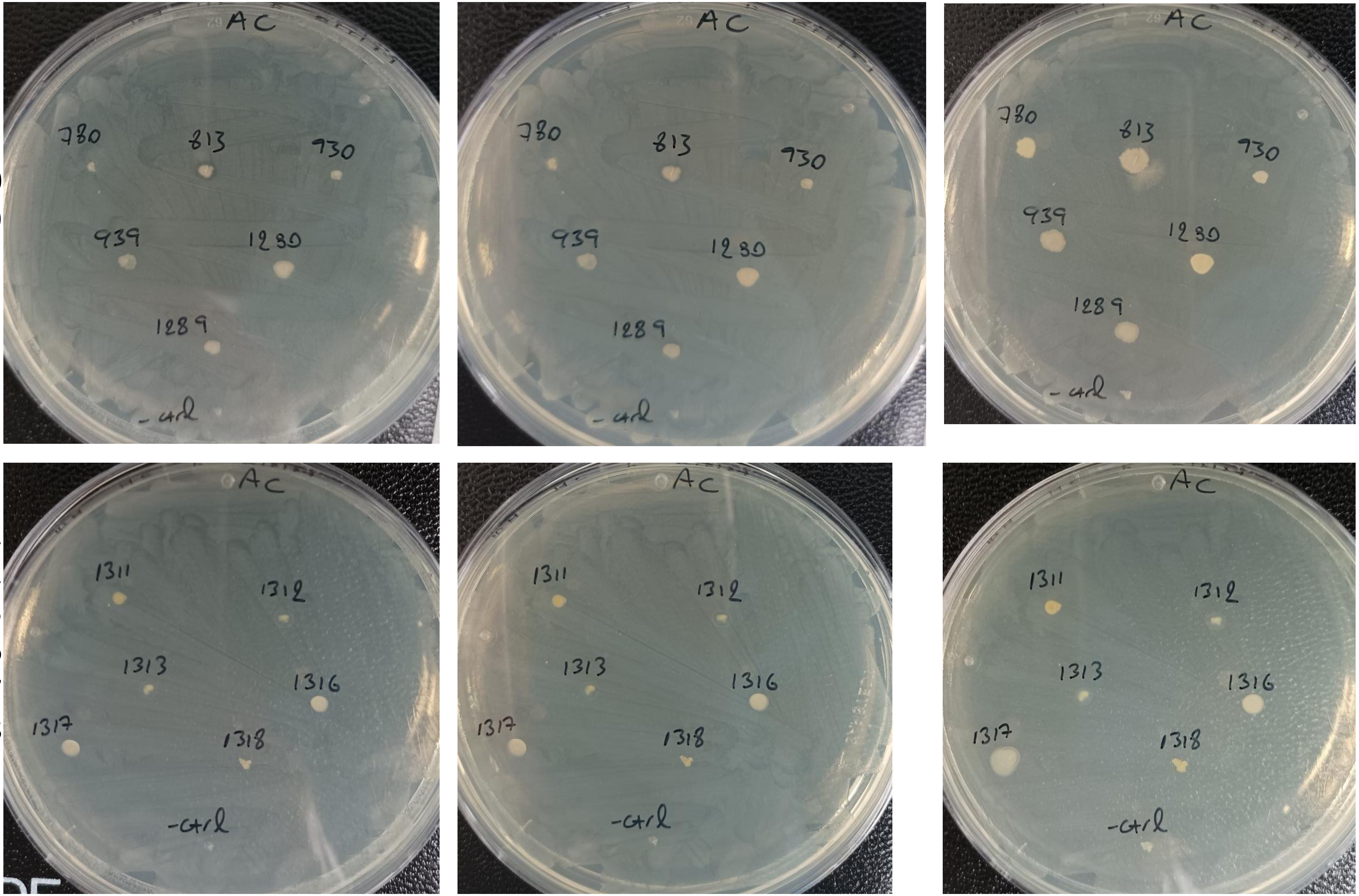

**Pc, NA**

24h | 48h

- SRL1123
- SRL1114
- SRL1322
- SRL1333
- SRL1055
- SRL778
- SRL1343
- SRL1175
- SRL1113

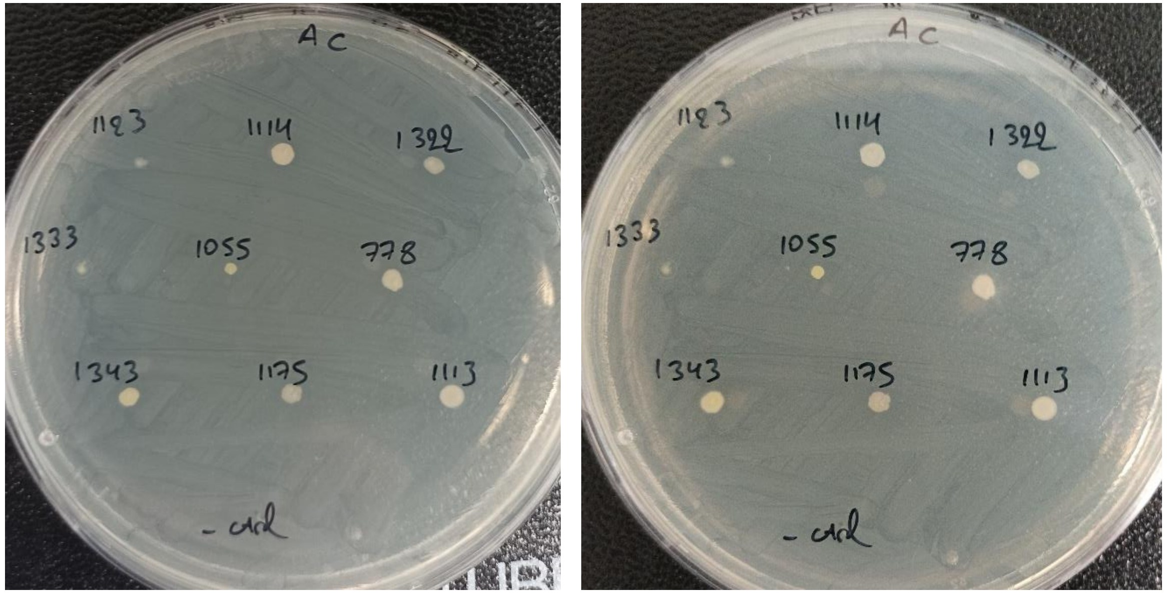


- SRL1342
- SRL1124
- SRL1115
- **SRL982**
- **SRL983**
- SRL889
- SRL1049
- **SRL1308**
- SRL838
- **SRL837**

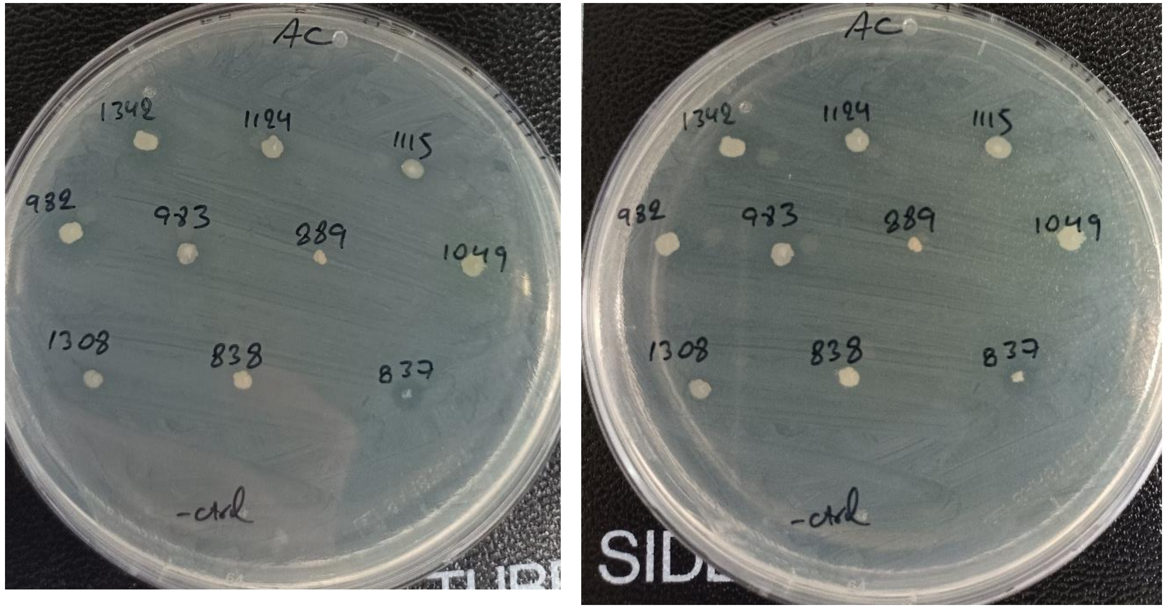

**Pc, NA**

- SRL1324
- **SRL986**
- SRL1334
- **SRL871**
- SRL990
- SRL1344
- SRL1121
- SRL1345
- **SRL879**
- SRL917

- SRL874
- SRL981
- SRL974
- SRL976
- SRL1057
- SRL1048
- SRL729
- SRL834

24h 48h

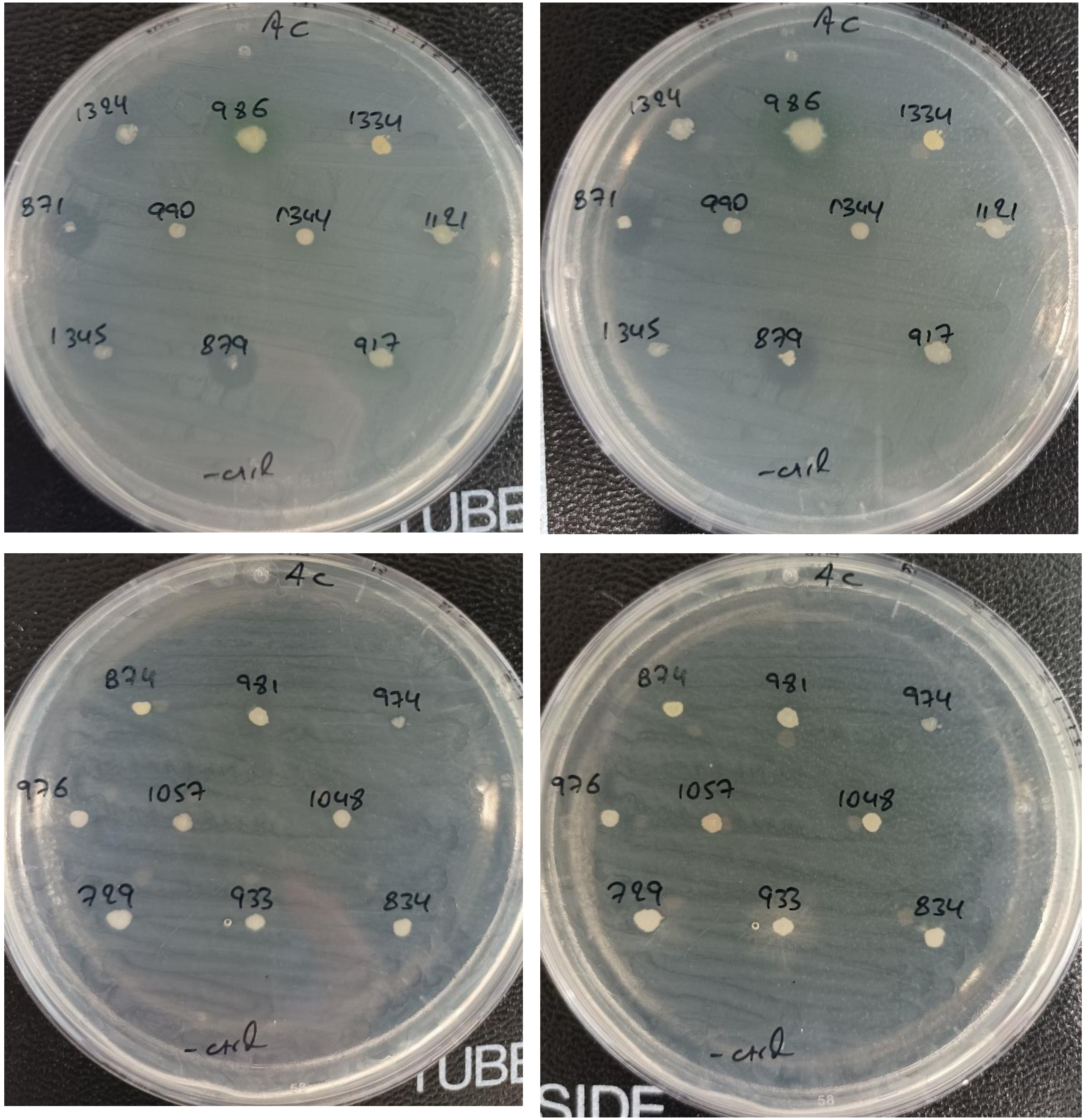

### *Clavibacter michiganensis* (Cm)

**Cm, NA**

72h

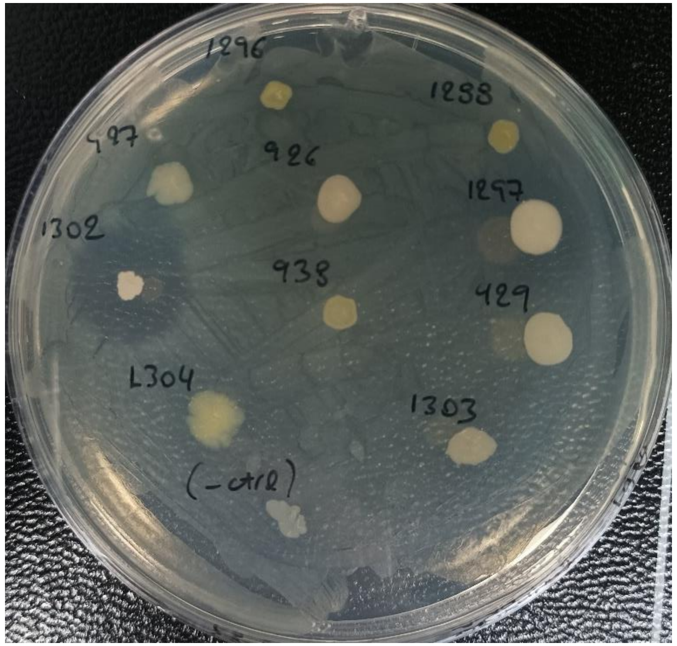


- SRL1296
- SRL1288
- SRL927
- SRL926
- SRL1297
- **SRL1302**
- SRL938
- SRL929
- SRL1304
- SRL1303

72h

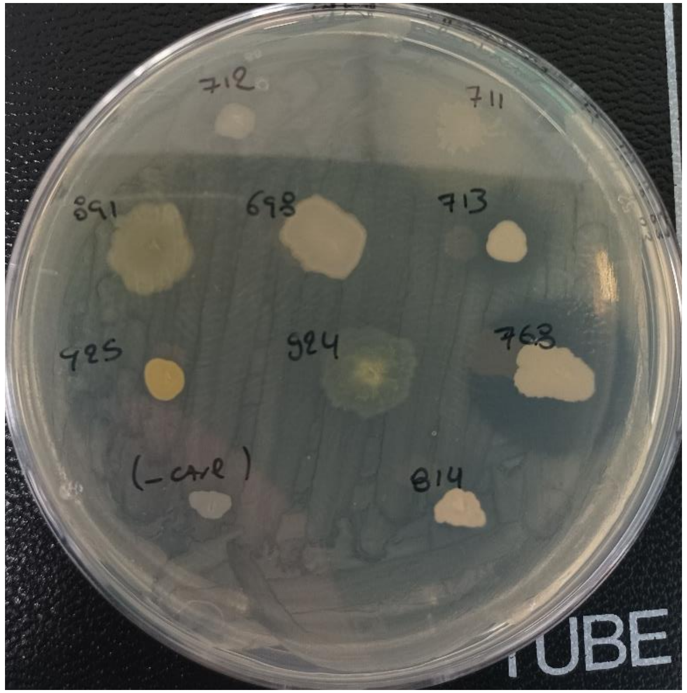


- SRL712
- SRL711
- SRL891
- SRL698
- **SRL713**
- SRL925
- SRL924
- **SRL768**
- SRL814

**Cm, NA ½**

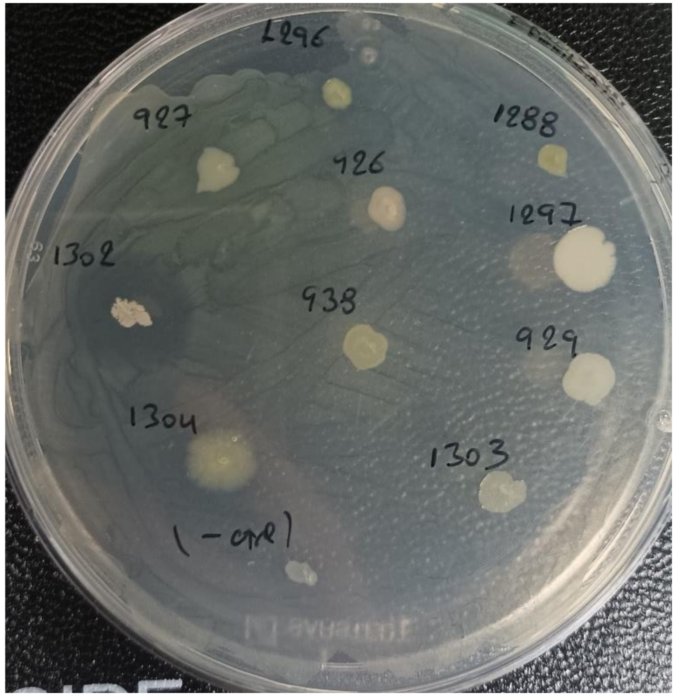


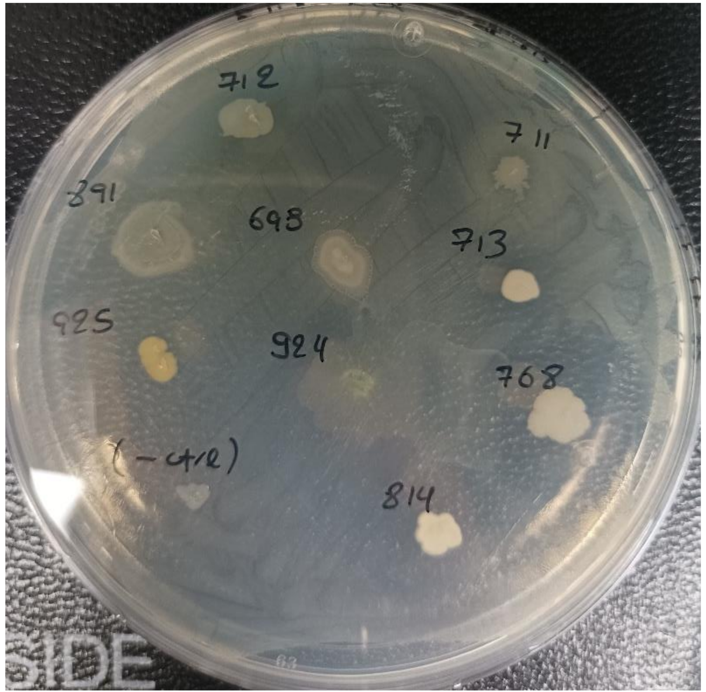

**Cm, NA**

- SRL730
- **SRL769**

48h

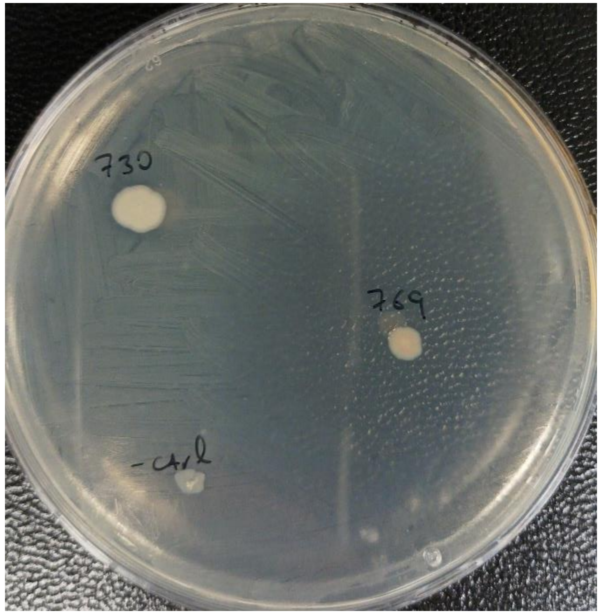


120h

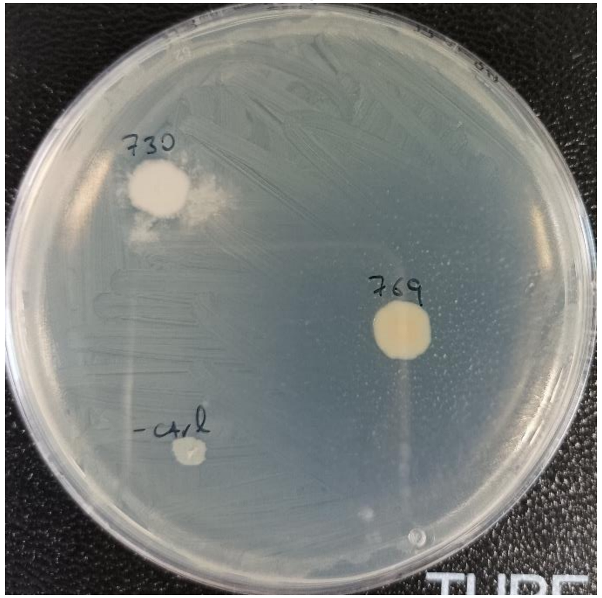


- SRL773
- SRL728

48h

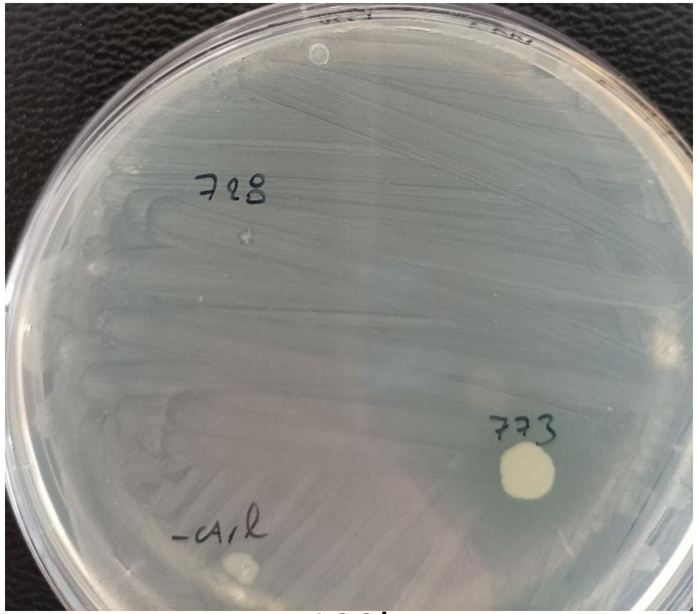


120h

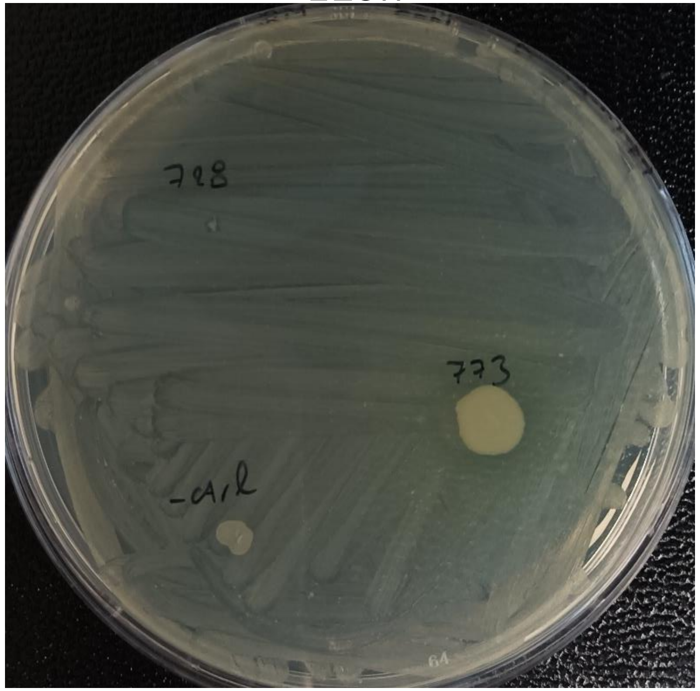


- SRL772
- SRL771

48h

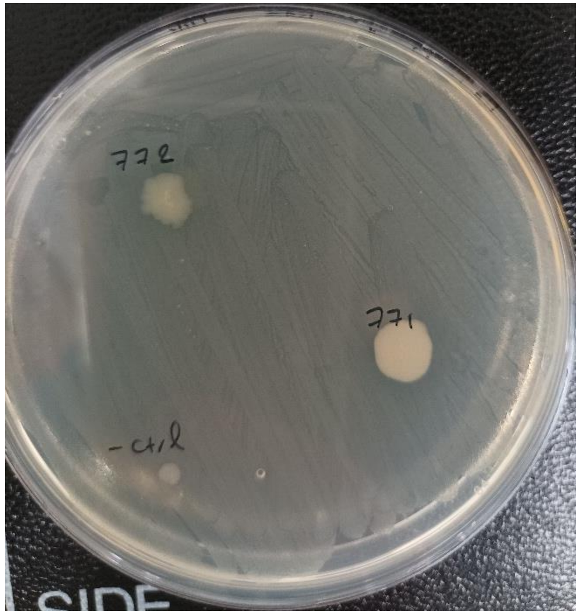

**Cm, NA**

- **SRL774**
- SRL775
- SRL 690
- **SRL764**
- **SRL765**
- **SRL766**
- **SRL767**
- **SRL689**
- SRL691

72h

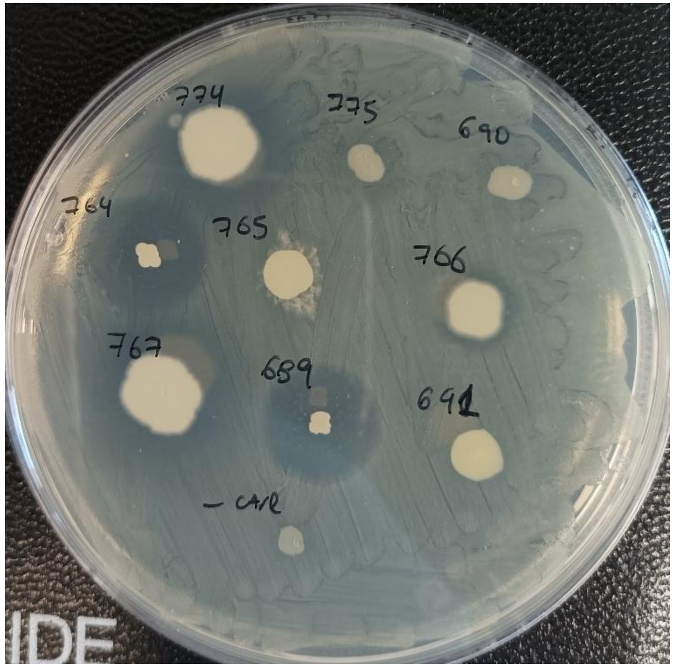

**Cm, NA**

48h | 112h

- SRL820
- SRL821
- SRL822
- SRL823
- SRL824
- SRL826
- SRL829
- SRL858
- SRL890
- **SRL931**

- SRL800
- SRL809
- **SRL810**
- SRL811
- SRL812
- **SRL815**
- SRL816
- SRL817
- SRL818

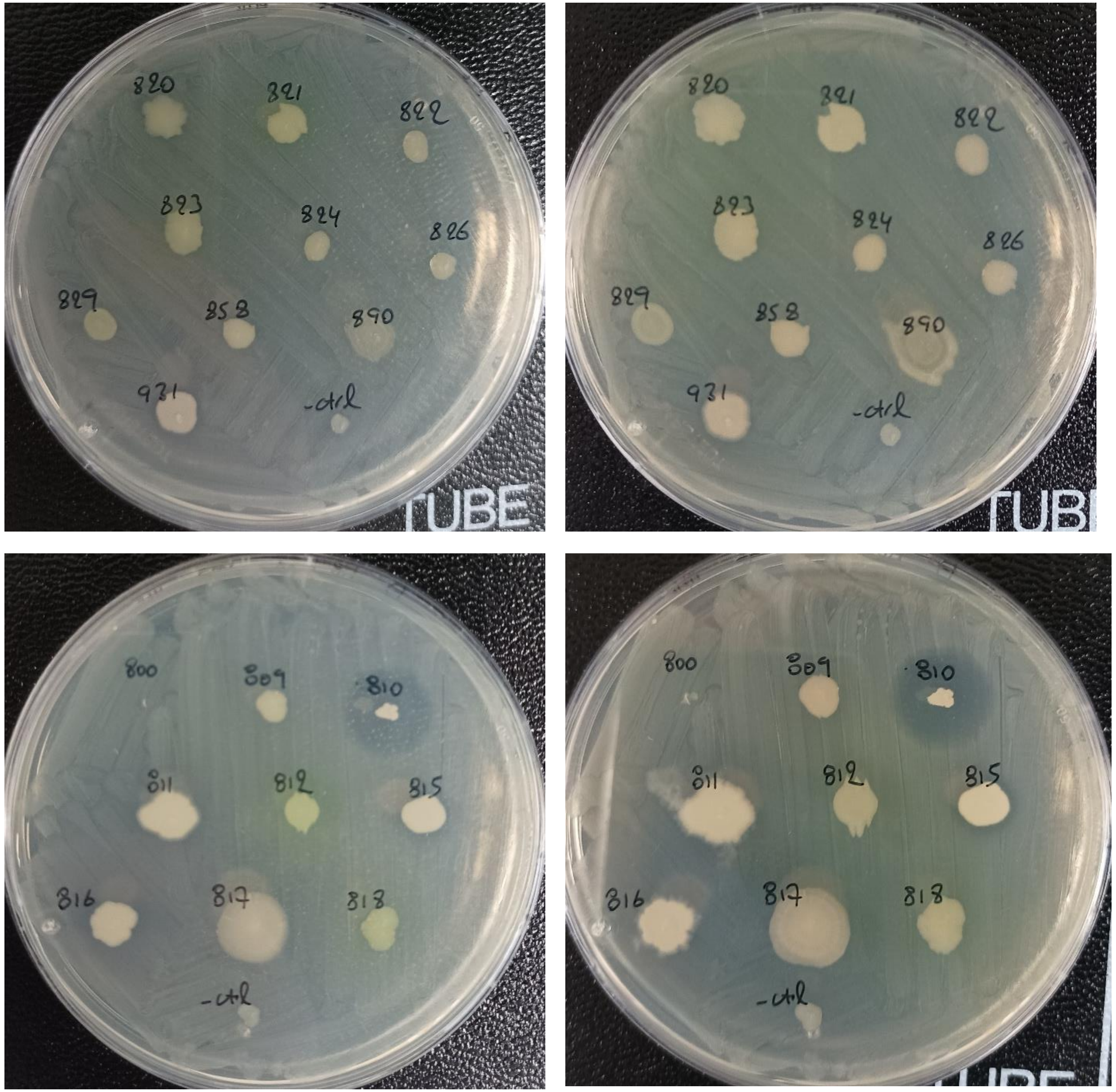

**Cm, NA**

48h | 144h

- SRL780
- SRL813
- SRL930
- SRL939
- SRL1280
- SRL1289

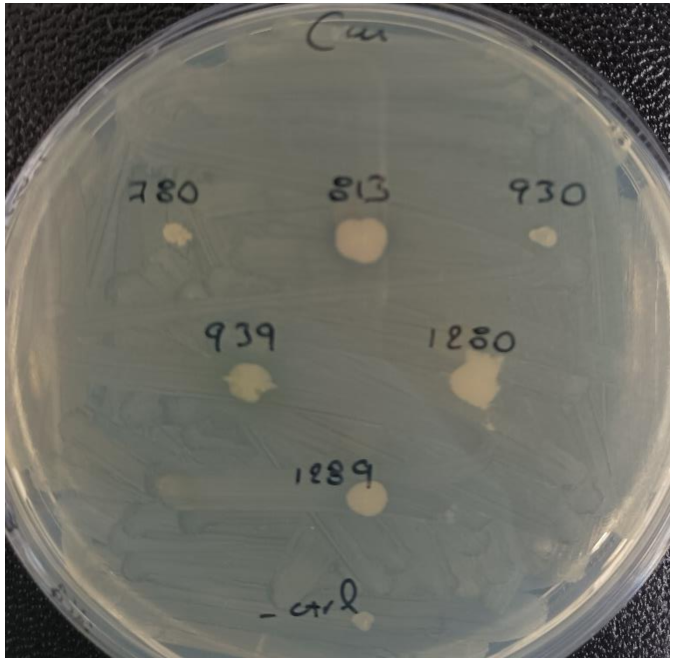


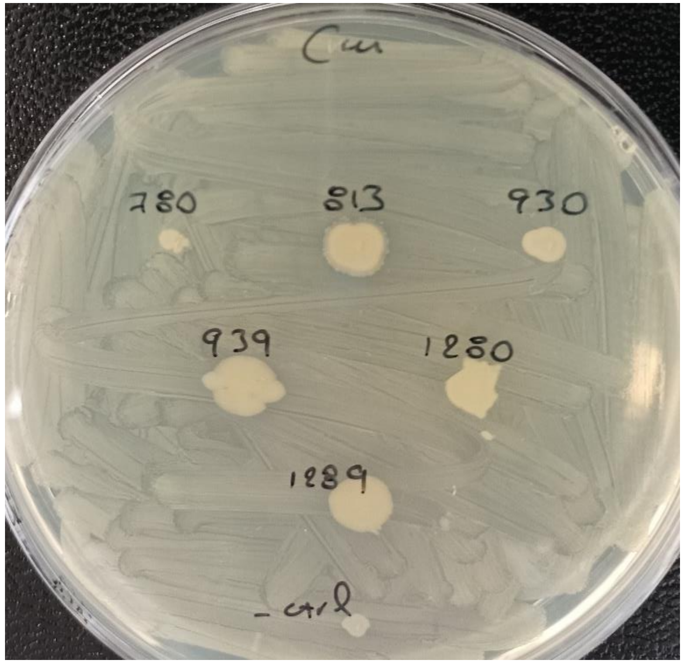


- SRL1311
- SRL1312
- SRL1313
- SRL1316
- SRL1317
- SRL1318

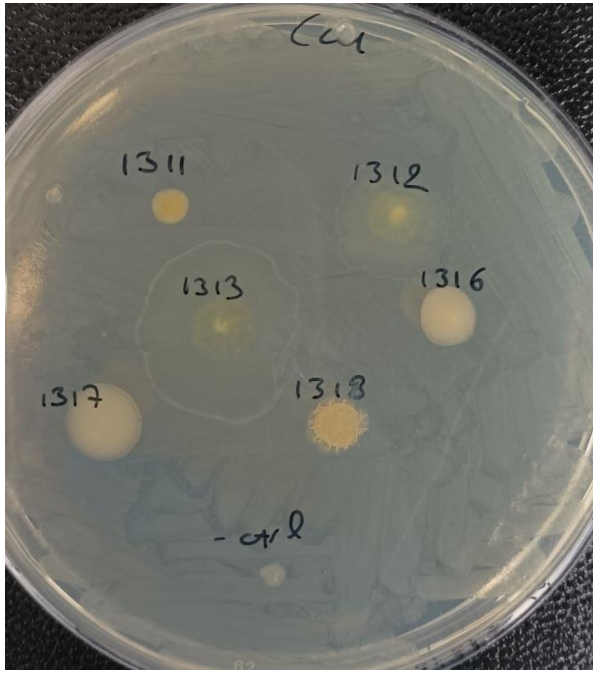


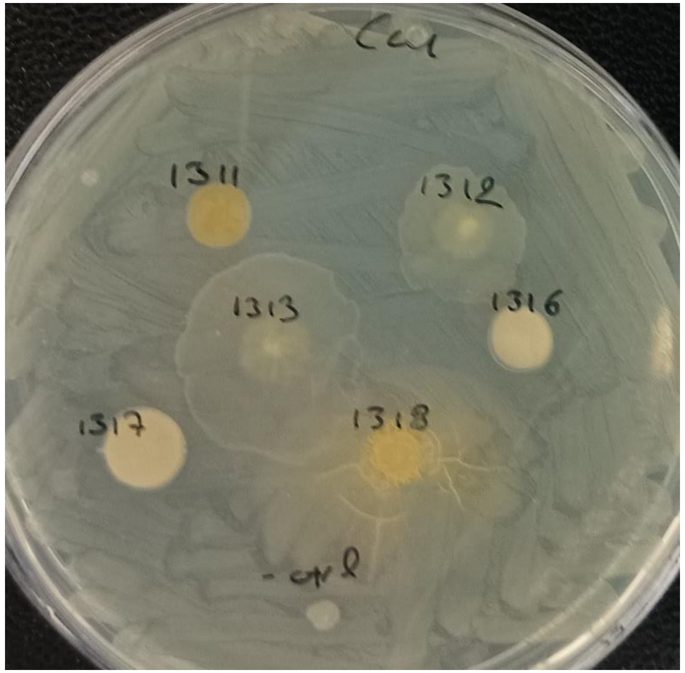

**Cm, NA**

- SRL1123
- SRL1114
- SRL1322
- SRL1333
- SRL1055
- **SRL778**
- SRL1343
- SRL1175
- SRL1113

48h

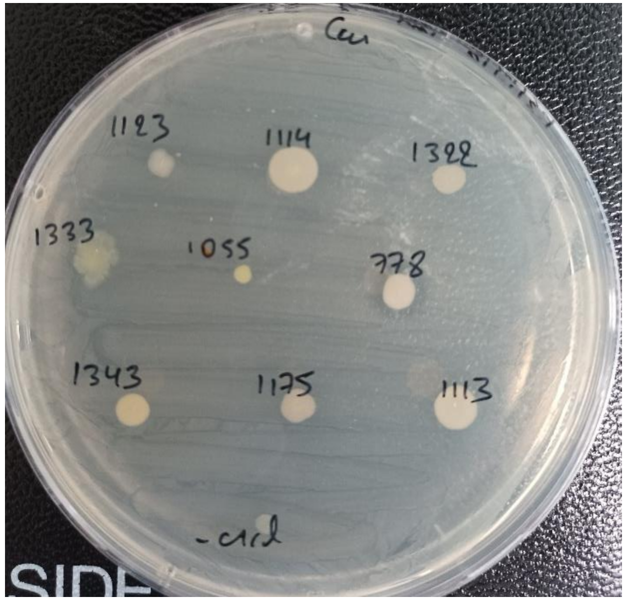


96h

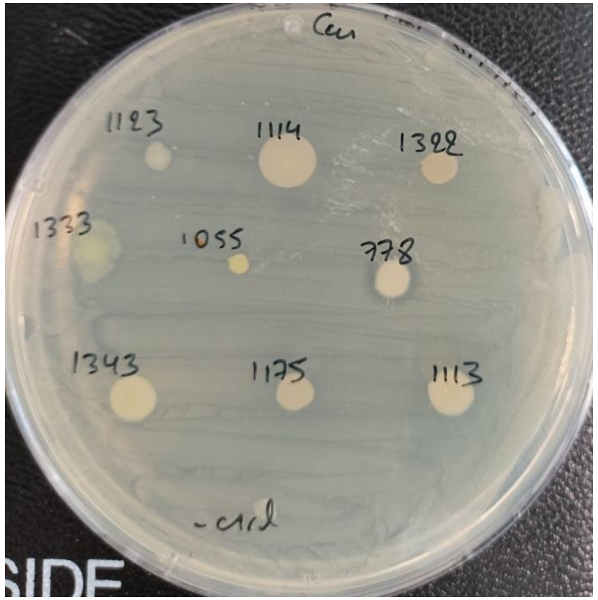


- SRL1342
- SRL1124
- SRL1115
- **SRL982**
- **SRL983**
- SRL889
- **SRL1049**
- SRL1308
- SRL838
- **SRL837**

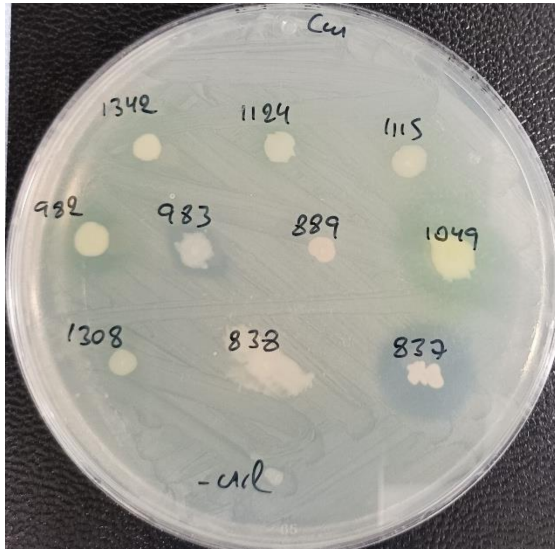


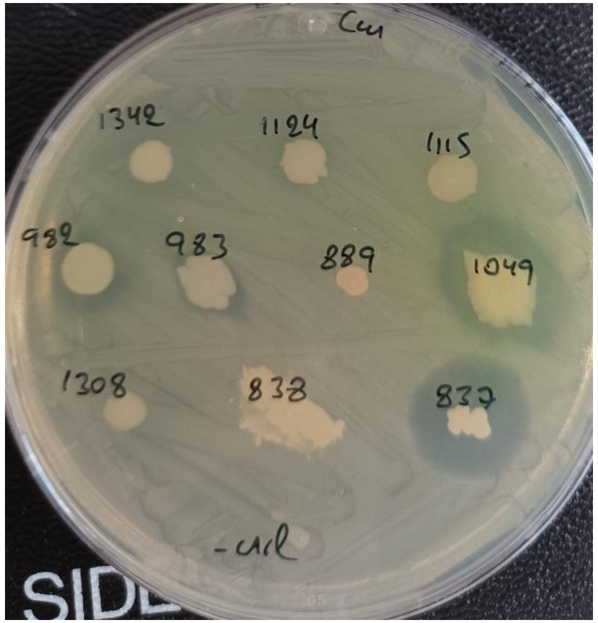

**Cm, NA**

- SRL1324
- **SRL986**
- SRL1334
- **SRL871**
- SRL990
- SRL1344
- SRL1121
- SRL1345
- **SRL879**
- **SRL917**

48h 96h

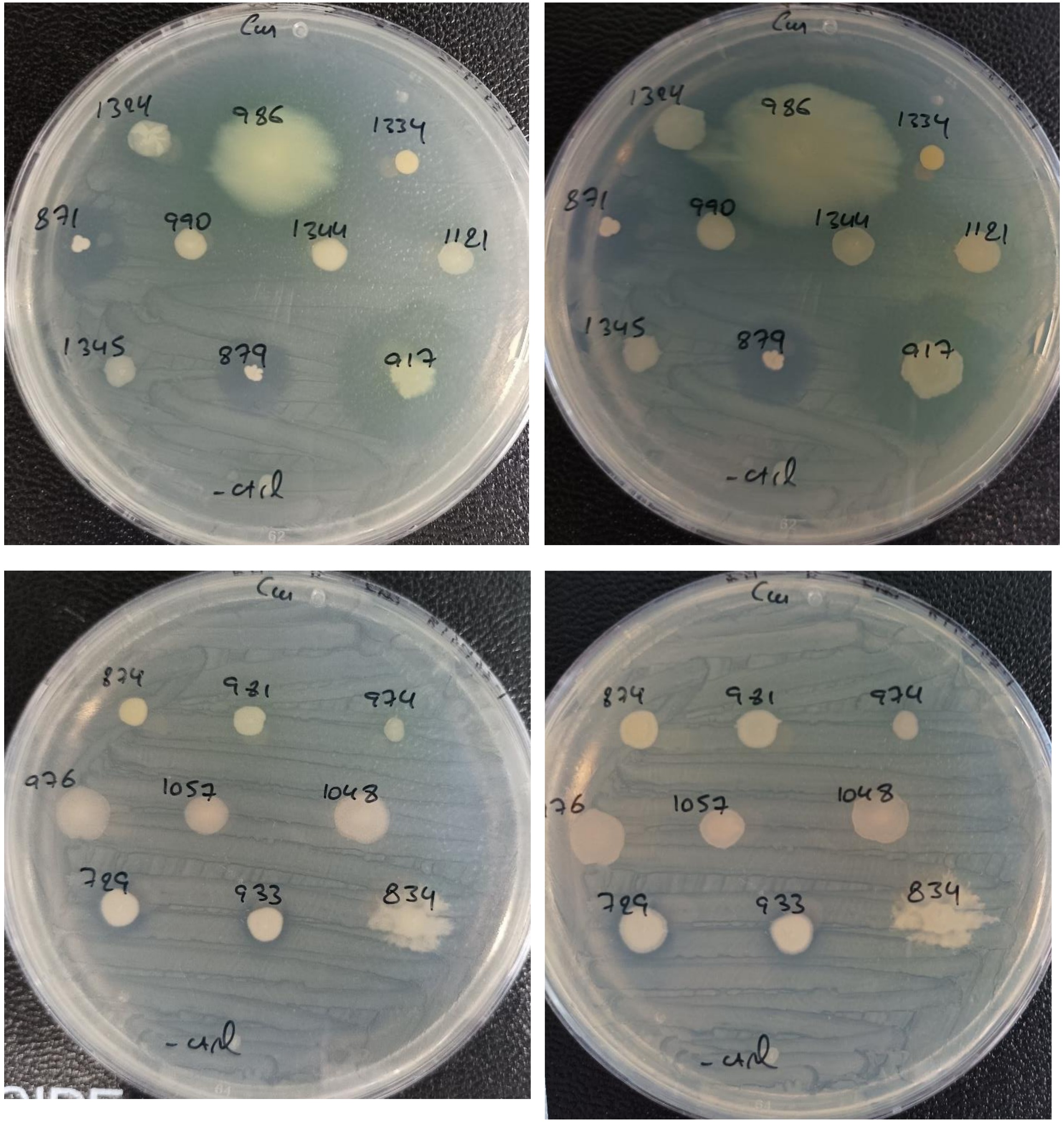


- SRL874
- SRL981
- SRL974
- SRL976
- SRL1057
- SRL1048
- **SRL729**
- SRL834
- **SRL933**

### ***Ralstonia solanacearum* (Rs)**

**Rs, NA**

- SRL1303
- **SRL1302**
- SRL814
- SRL1288
- SRL1296
- SRL1297
- SRL925
- SRL891
- SRL760

- **SRL689**
- SRL690
- SRL691
- SRL699
- SRL726
- SRL728
- SRL730
- SRL771
- SRL684

24h | 48h | 112h

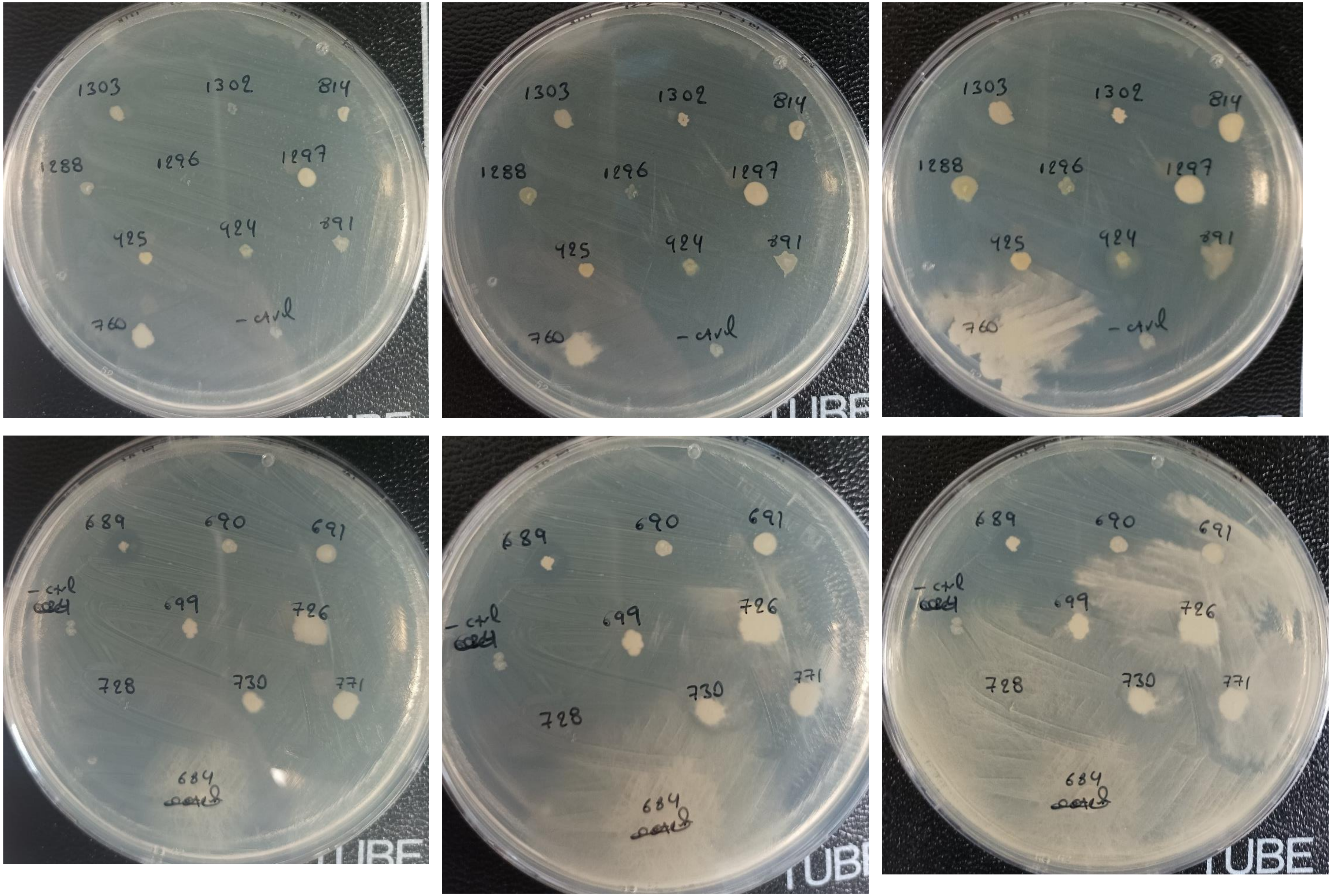

**Rs, NA**

- SRL772
- SRL773
- SRL769
- SRL774
- SRL775
- **SRL764**
- SRL765
- SRL766
- SRL767

- SRL698
- SRL711
- SRL712
- SRL713
- SRL926
- SRL927
- SRL929
- SRL938
- SRL1304

24h 48h 112h

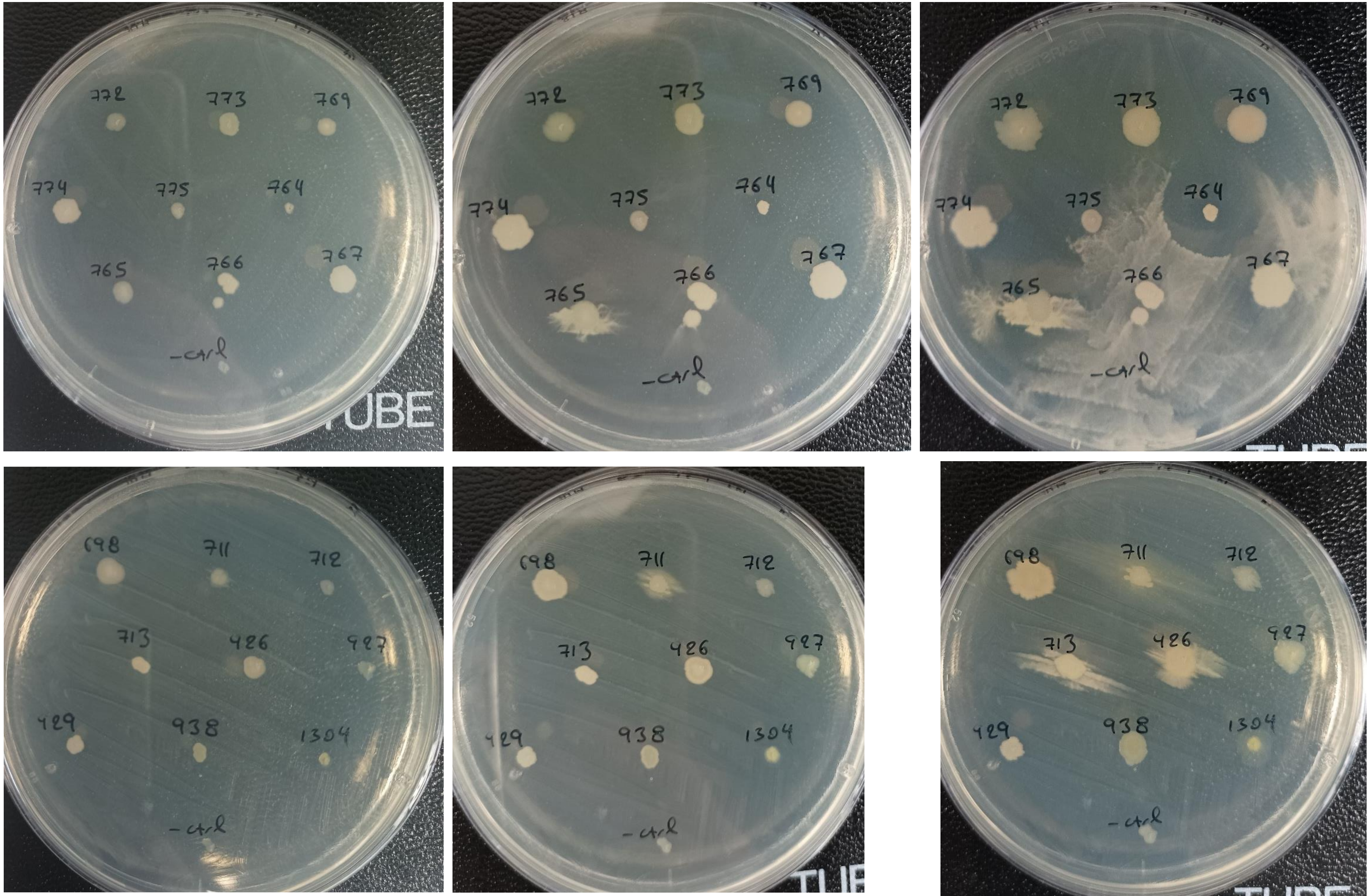

**Rs, NA**

- SRL934
- SRL936
- **SRL937**
- SRL1281
- **SRL1282**
- SRL1283
- SRL1290
- SRL1291
- SRL1292

24h

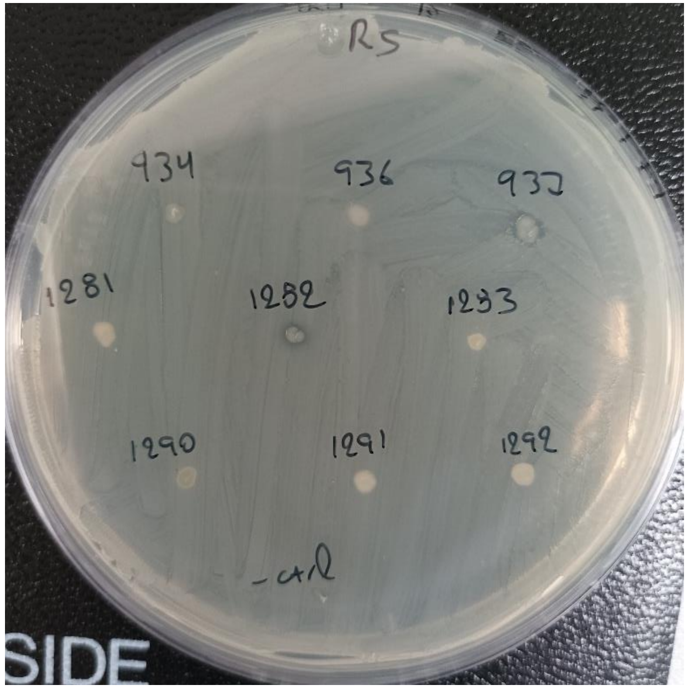


48h

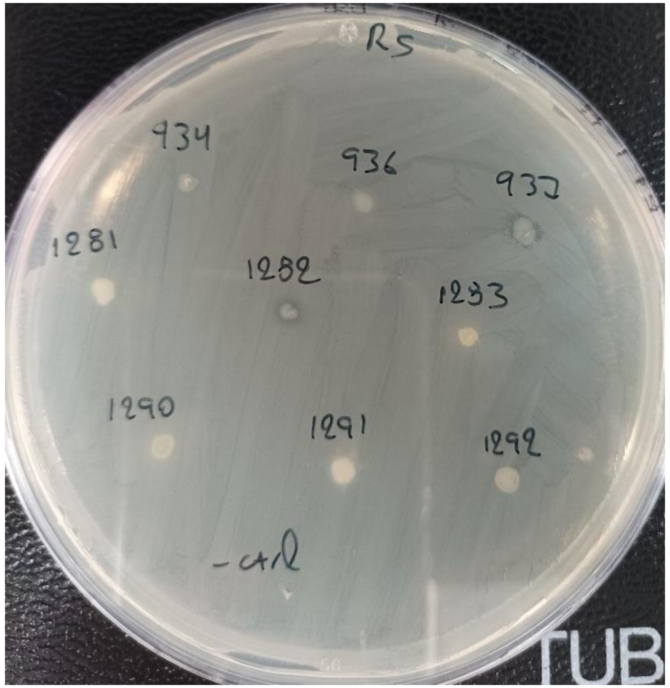


- SRL1298
- SRL1299
- SRL1300
- **SRL1301**
- SRL1305
- SRL1307
- SRL1309
- SRL1310

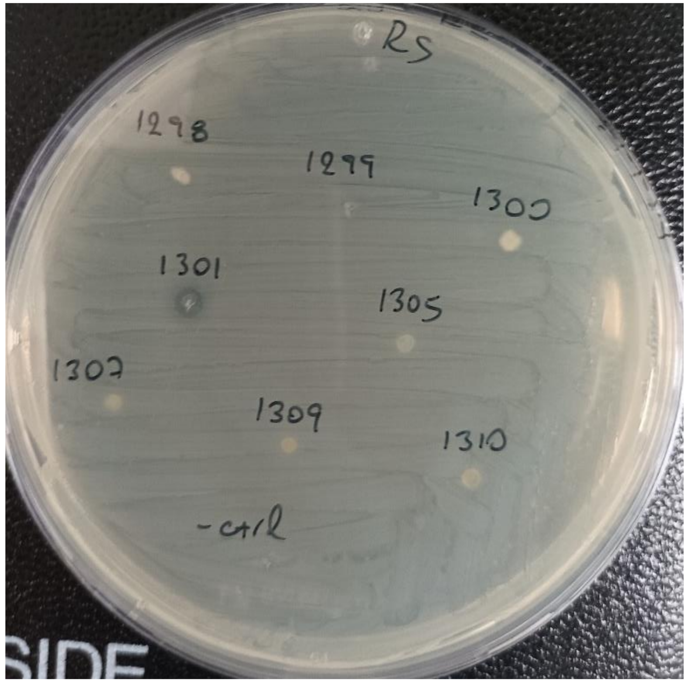


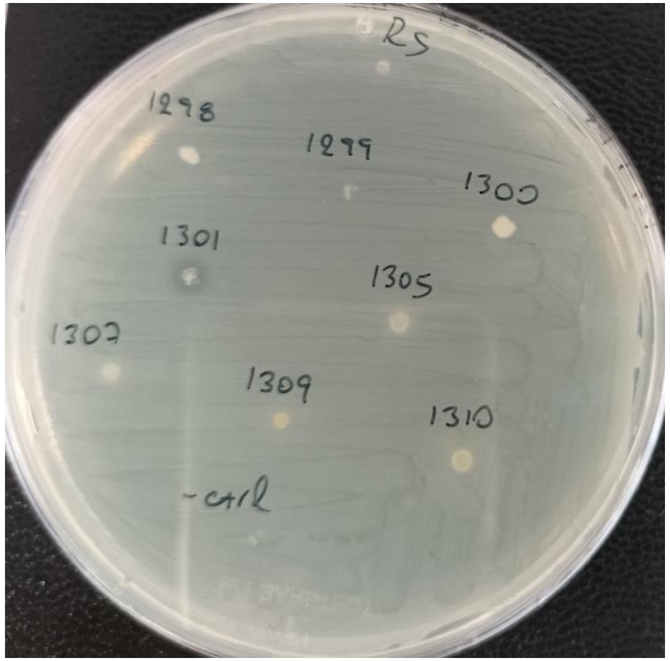

**Rs, NA**

| 24h | 48h | 112h |
|---|---|---|

- SRL820
- SRL821
- SRL822
- SRL823
- SRL824
- SRL826
- SRL829
- SRL858
- SRL890
- **SRL931**

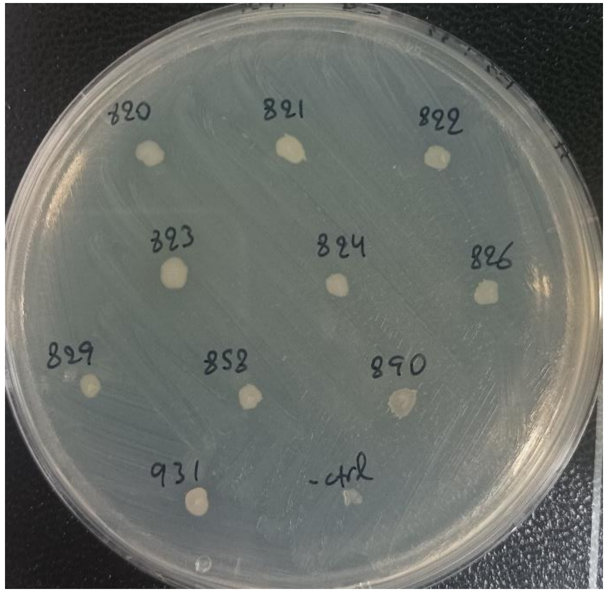

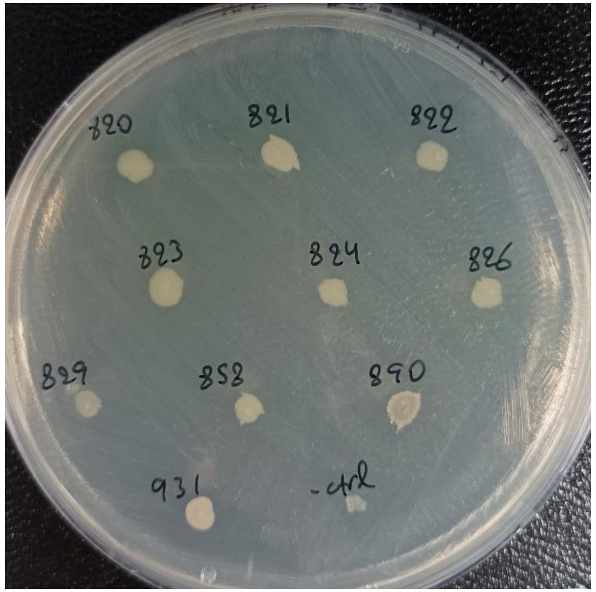

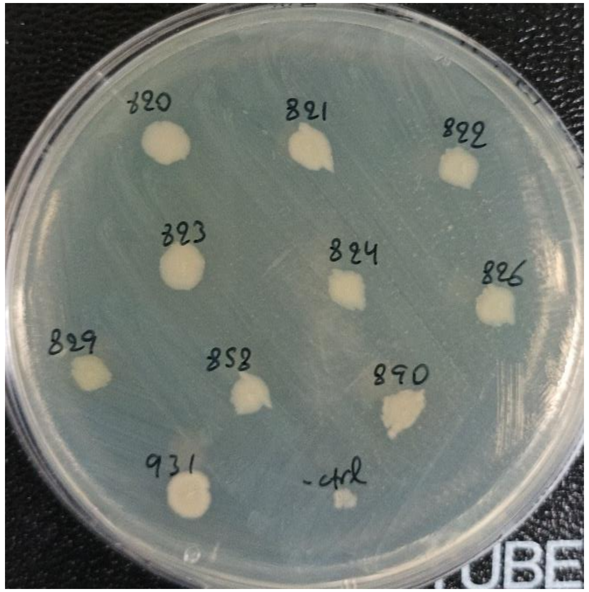


- SRL800
- SRL809
- **SRL810**
- SRL811
- SRL812
- SRL815
- SRL816
- SRL817
- SRL818

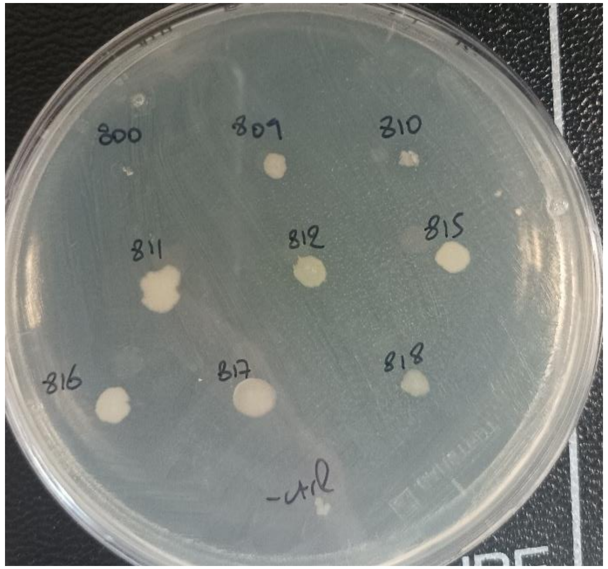

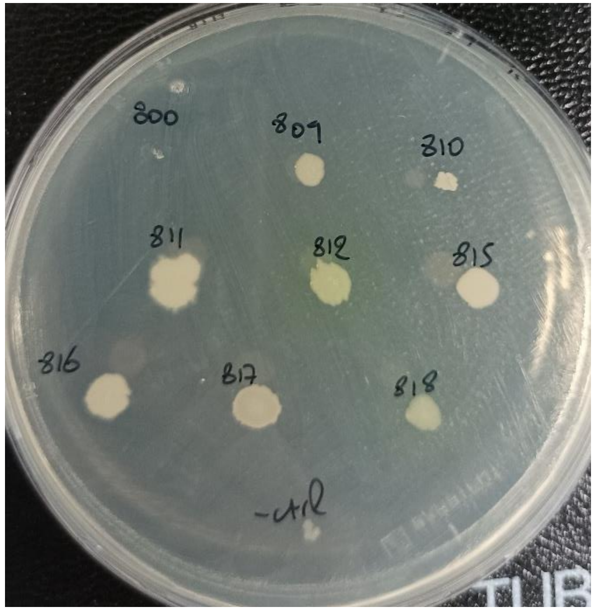

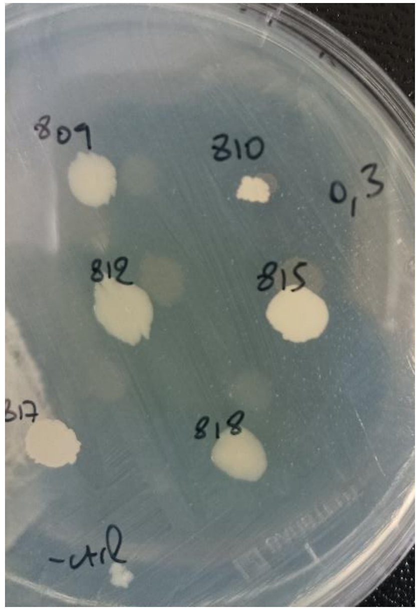

**Rs, NA**

42h 72h

- SRL780
- SRL813
- SRL930
- SRL939
- SRL1280
- SRL1289

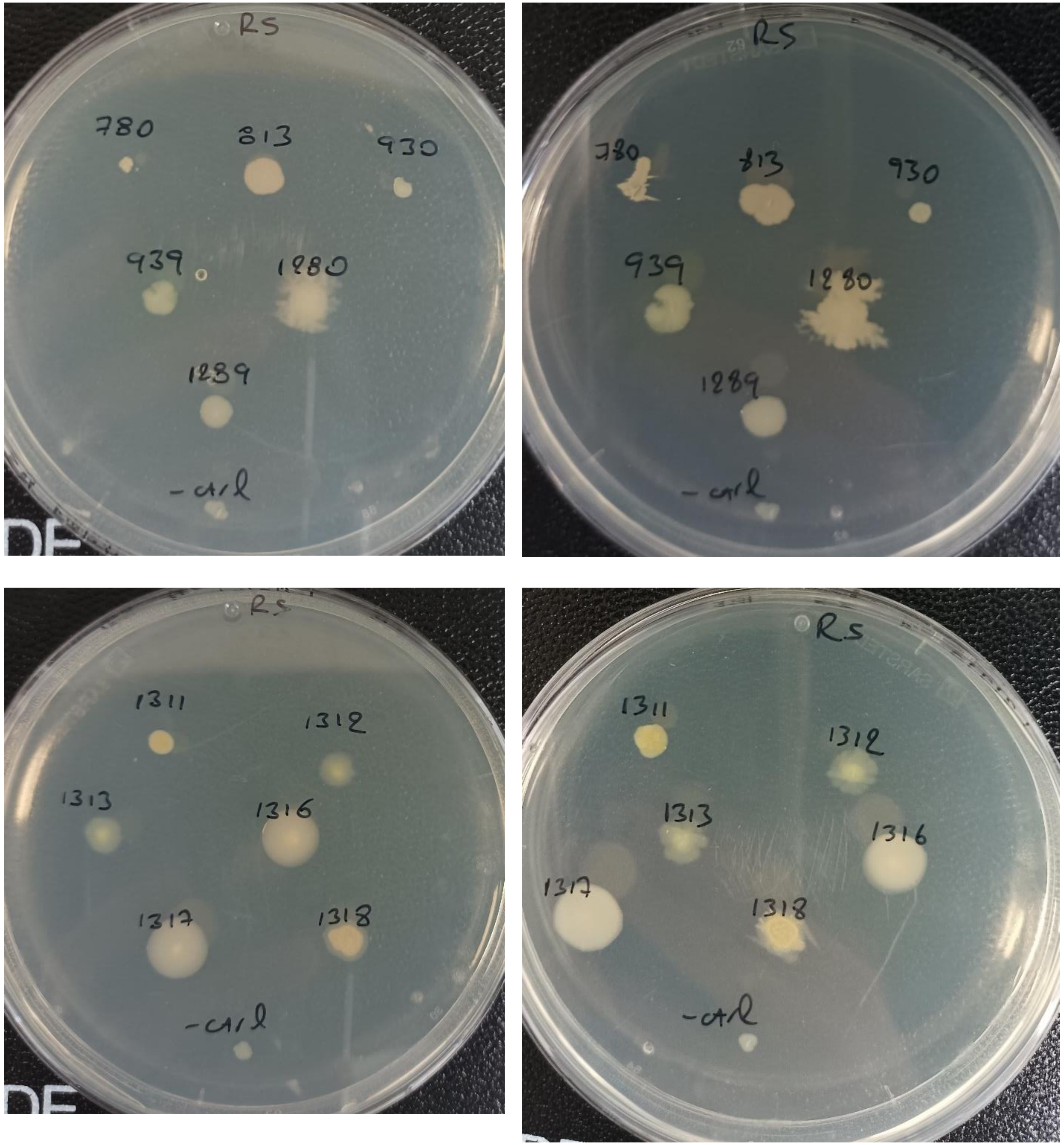


- SRL1311
- SRL1312
- SRL1313
- SRL1316
- SRL1317
- SRL1318

**Rs, NA**

48h | 72h

- SRL1116
- SRL874
- SRL981
- SRL974
- SRL976
- SRL1057
- SRL1048
- SRL729
- SRL933

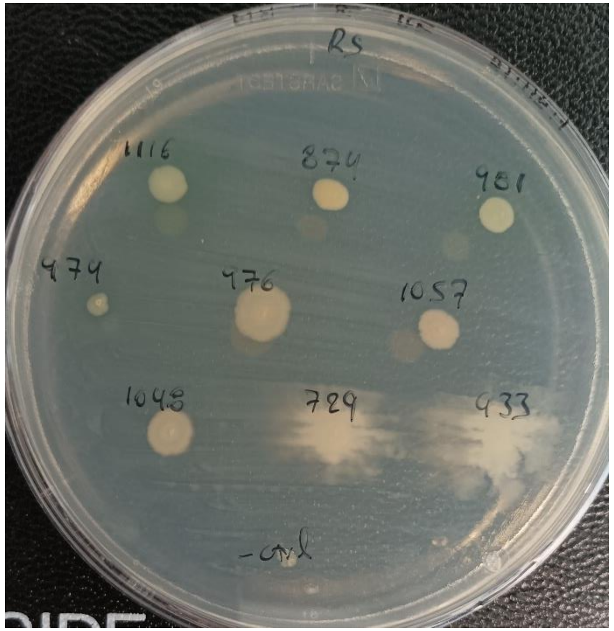


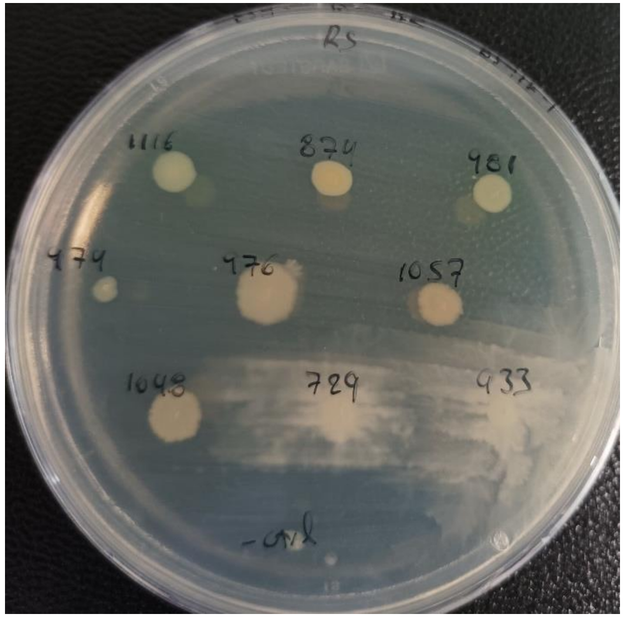


- SRL834
- SRL1342
- SRL1124
- SRL1115
- SRL982
- SRL983
- SRL889
- SRL1308

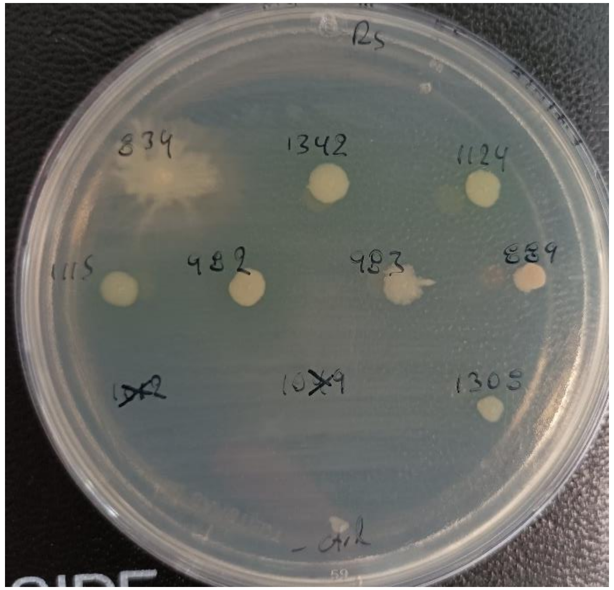


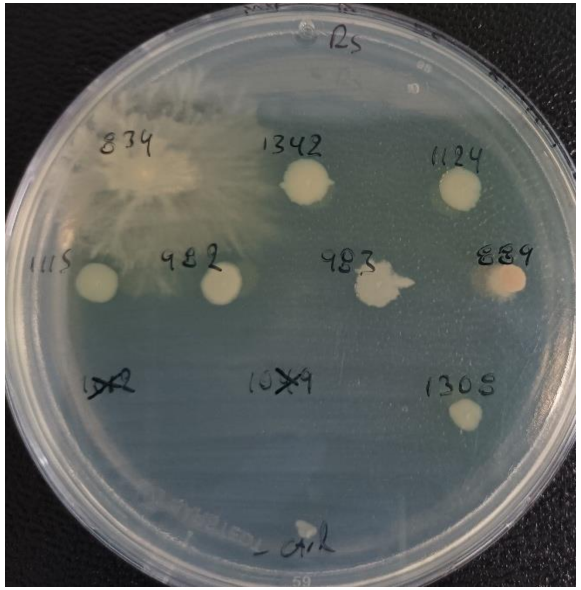

**Rs, NA**

- SRL838
- **SRL837**
- SRL1123
- SRL1114
- SRL1322
- SRL1333
- SRL1055
- SRL778
- SRL1343
- SRL1175

48h

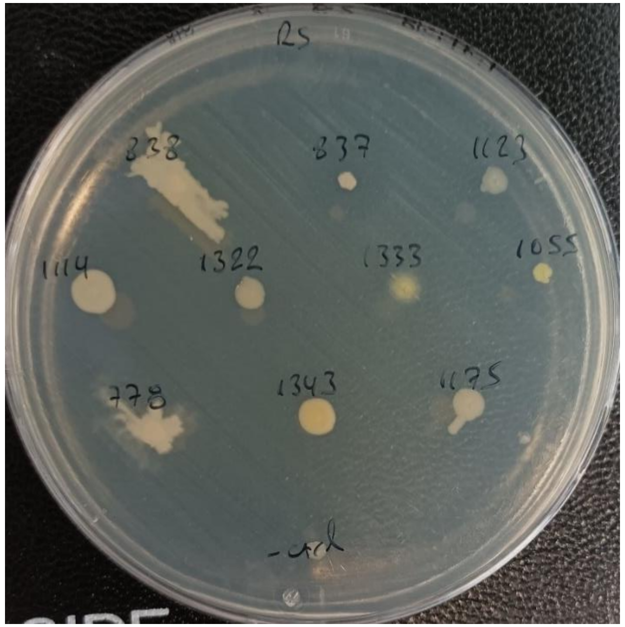

72h

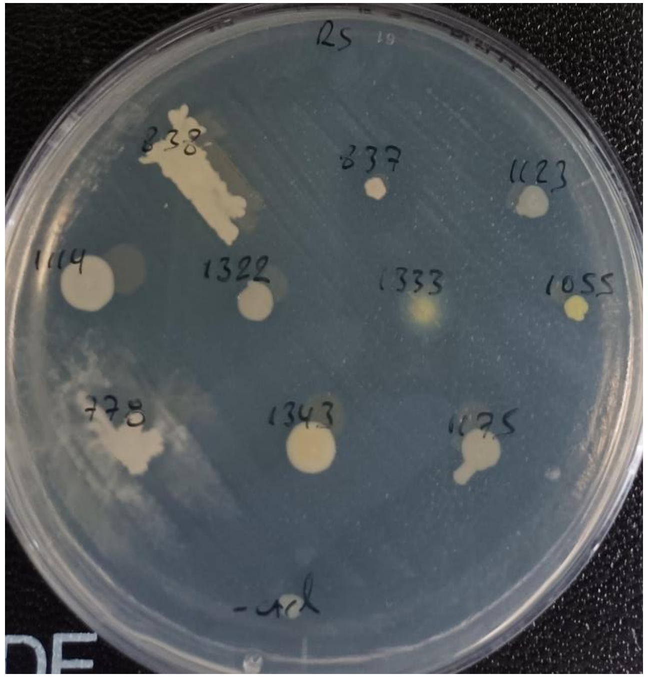

- SRL1113
- SRL1324
- SRL1334
- **SRL871**
- SRL990
- SRL1344
- SRL1121
- **SRL1345**
- **SRL879**
- SRL917

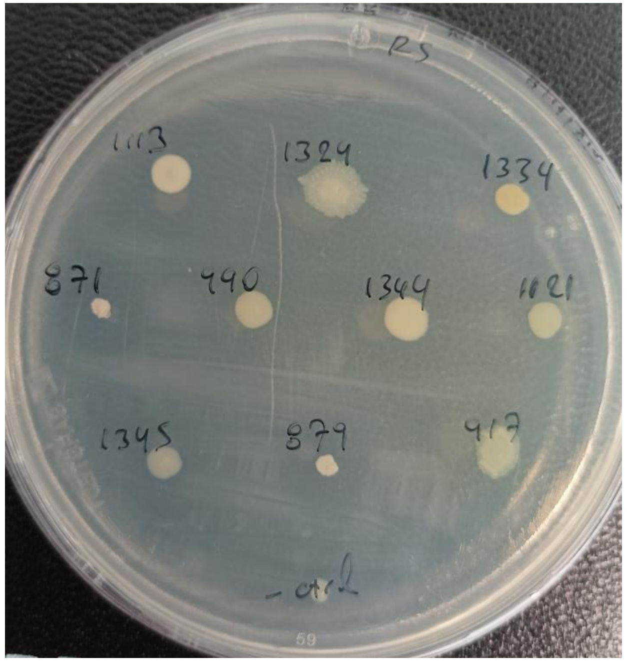

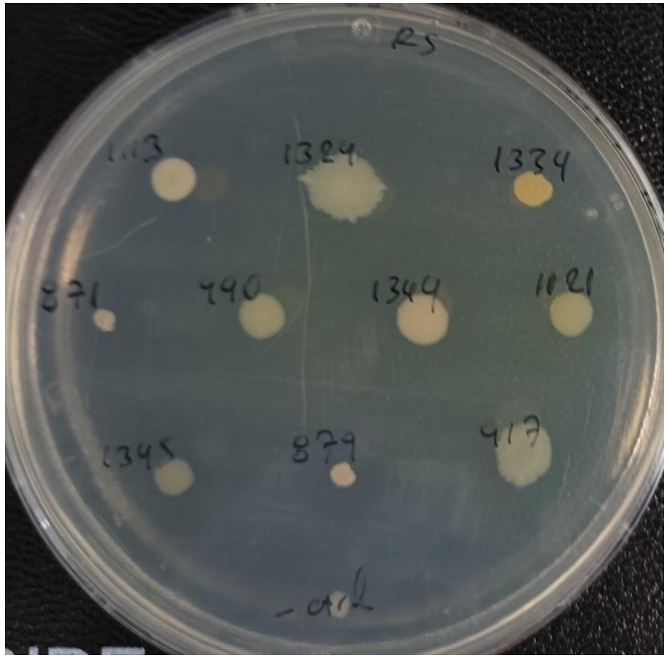

*Streptomyces* spp, SRL740, SRL742 and SRL1060
*Nocardiopsis* sp., SRL1020

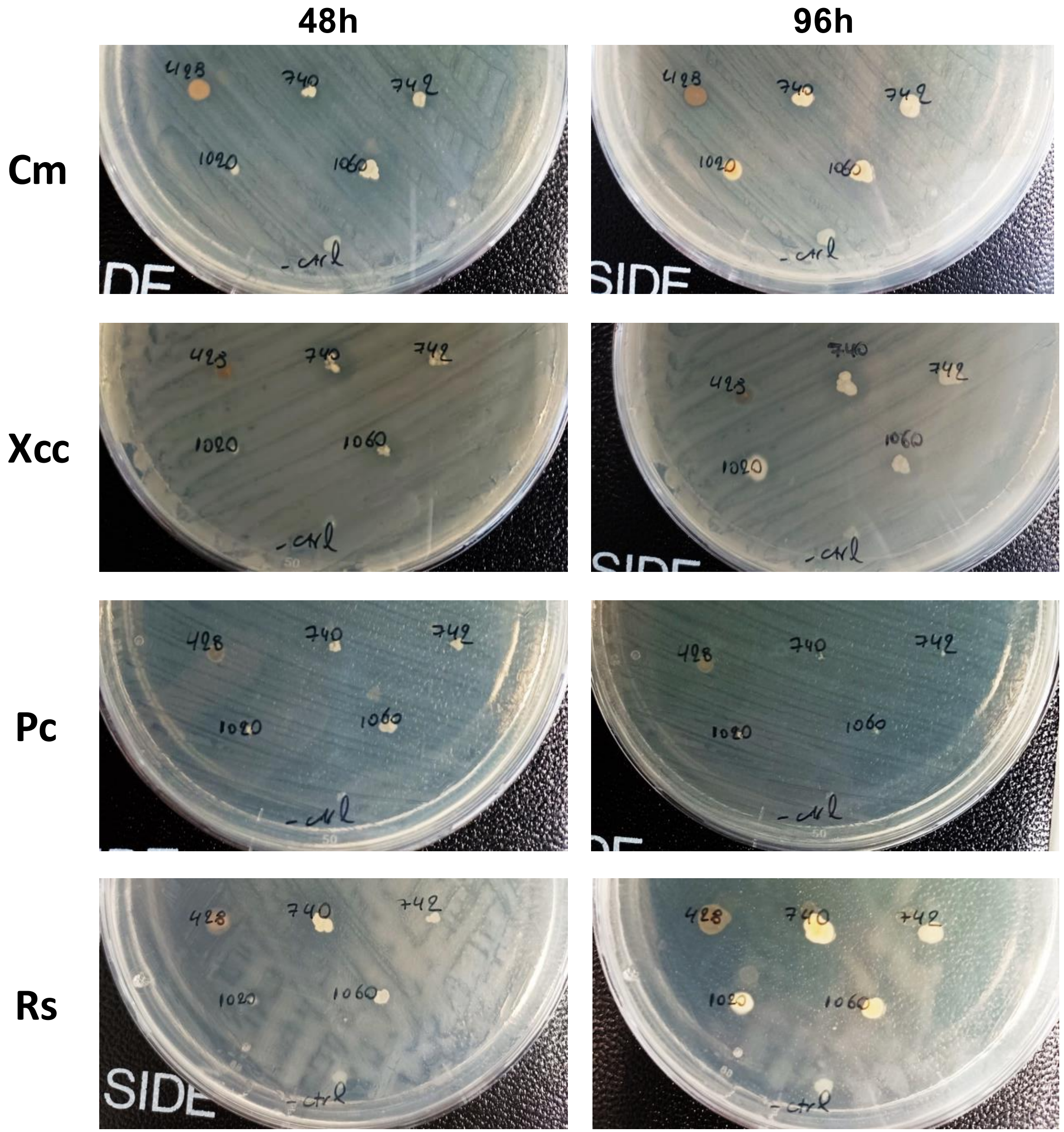
