## Supplementary Presentation 3 for "A Thousand Meters Deep: Vertical Profiling of the Subterranean Microbes of Gourgouthakas Cave"

### *Phytopthora nicotianae*

*Ph.nicotianae*

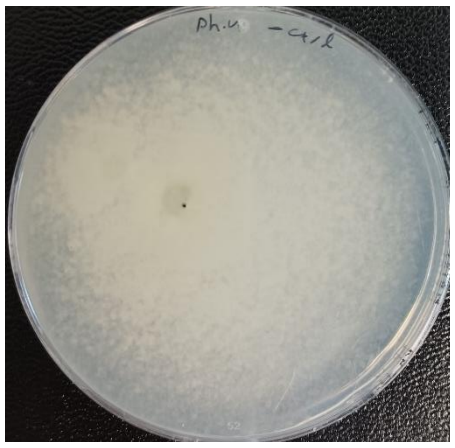

*Ph.nicotianae*/SRL684

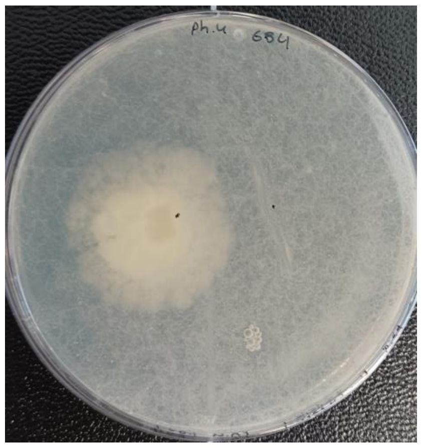

*Ph.nicotianae*/SRL689

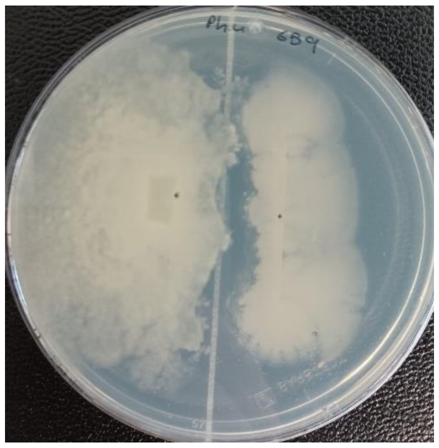

*Ph.nicotianae*/SRL690

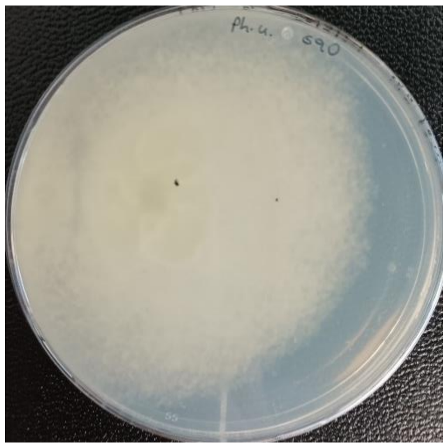

*Ph.nicotianae*/SRL691

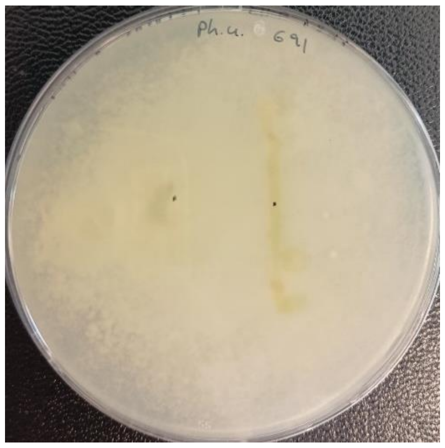

*Ph.nicotianae*/SRL698

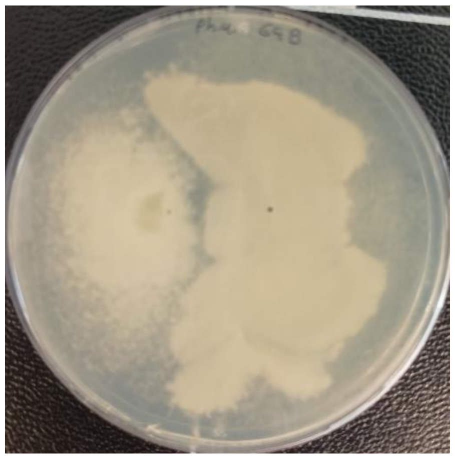

*Ph.nicotianae*/SRL699

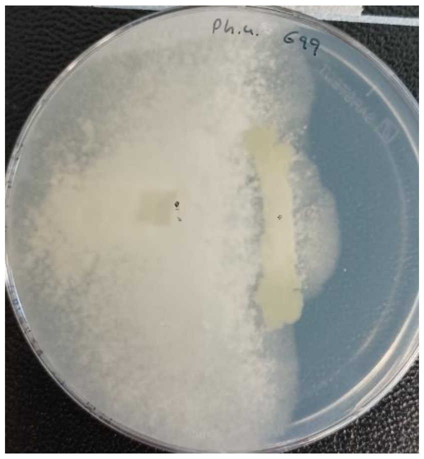

*Ph.nicotianae*/SRL711

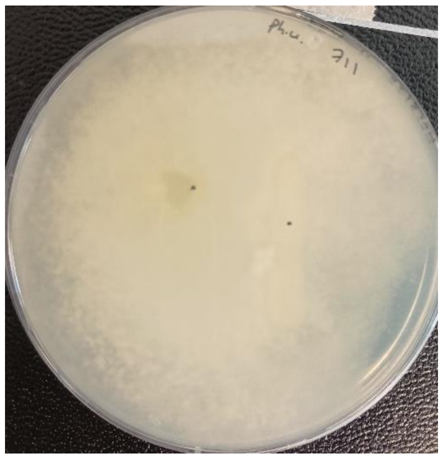

*Ph.nicotianae*/SRL712

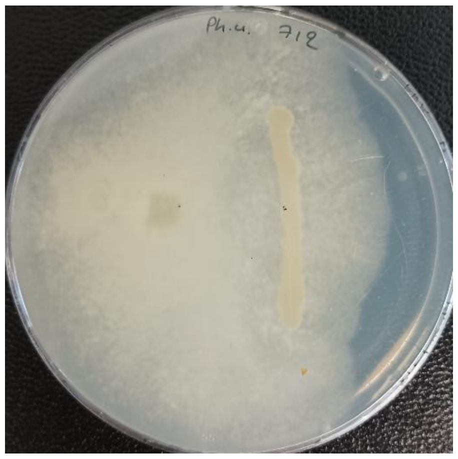

*Ph.nicotianae*/SRL713

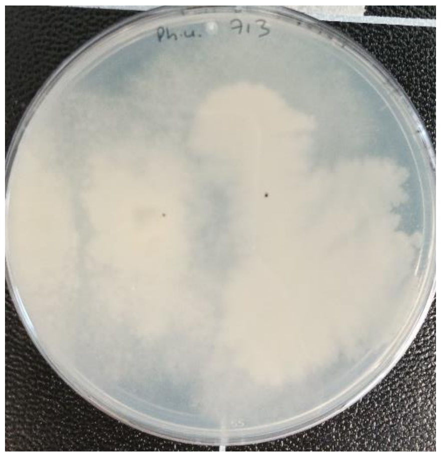

*Ph.nicotianae*/SRL726

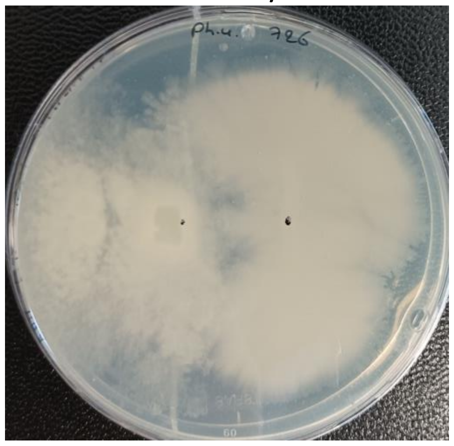

*Ph.nicotianae*/SRL727

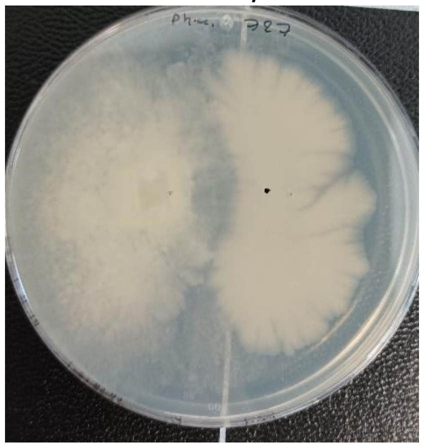

*Ph.nicotianae*/SRL728

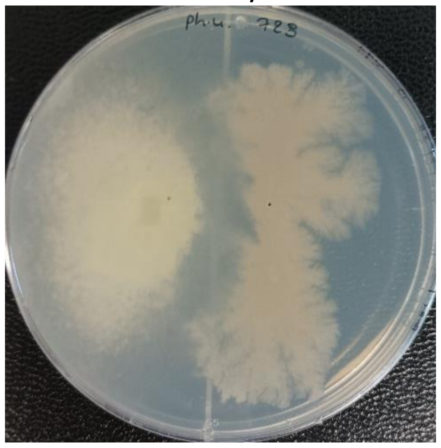

*Ph.nicotianae*/SRL729

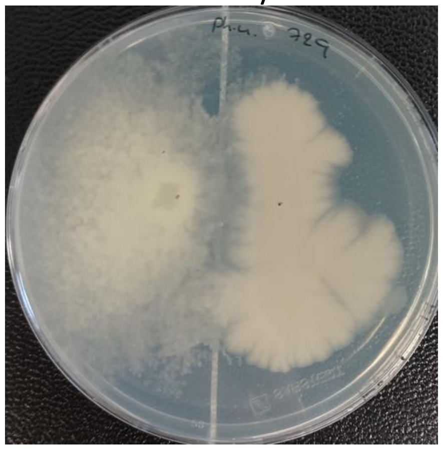

*Ph.nicotianae*/SRL730

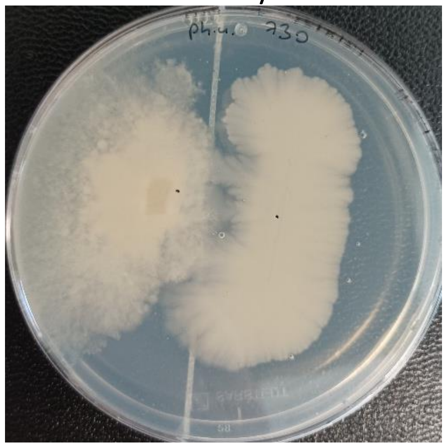

*Ph.nicotianae*/SRL731

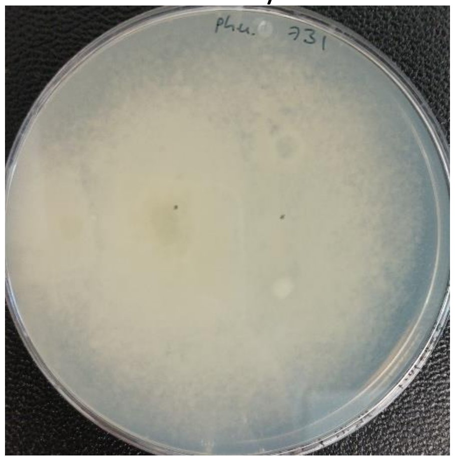

*Ph.nicotianae*/SRL732

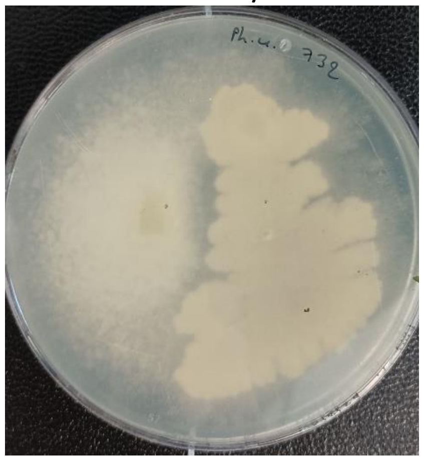

*Ph.nicotianae*/SRL764

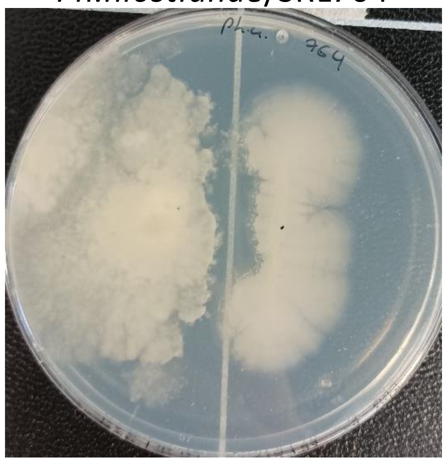

*Ph.nicotianae*/SRL765

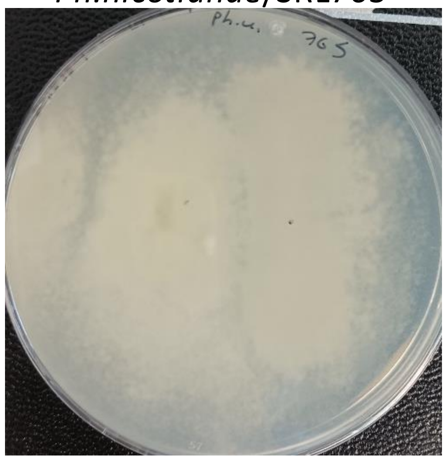

*Ph.nicotianae*/SRL766

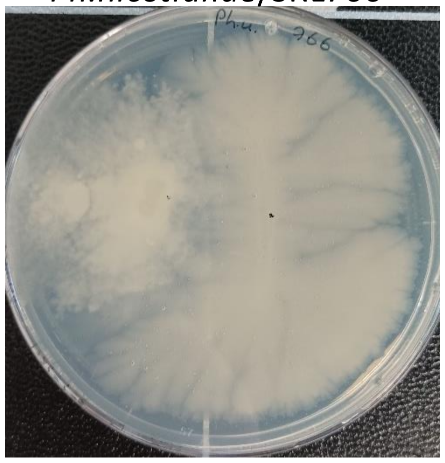

*Ph.nicotianae*/SRL767

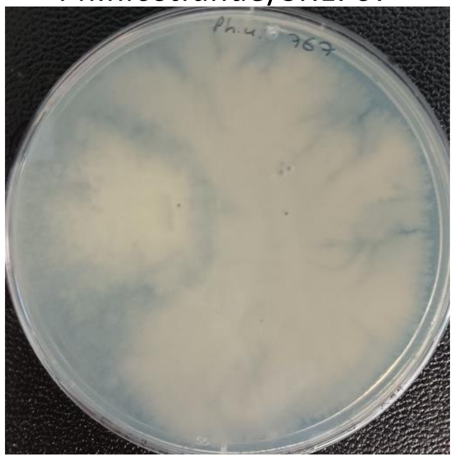

*Ph.nicotianae*/SRL768

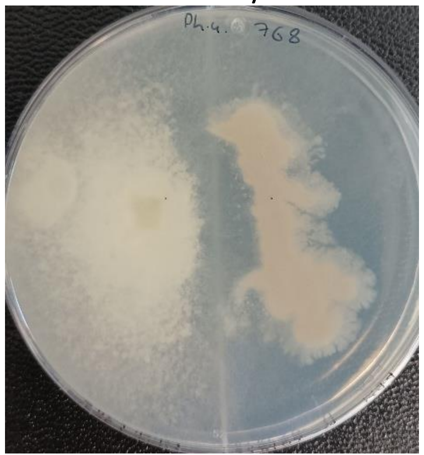

*Ph.nicotianae*/SRL769

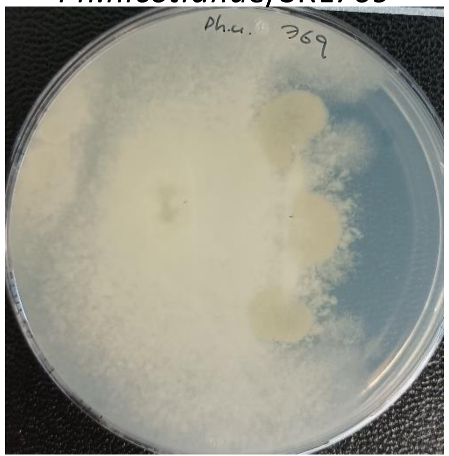

*Ph.nicotianae*/SRL771

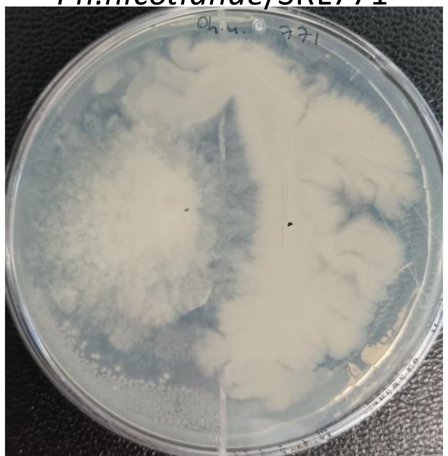

(A)

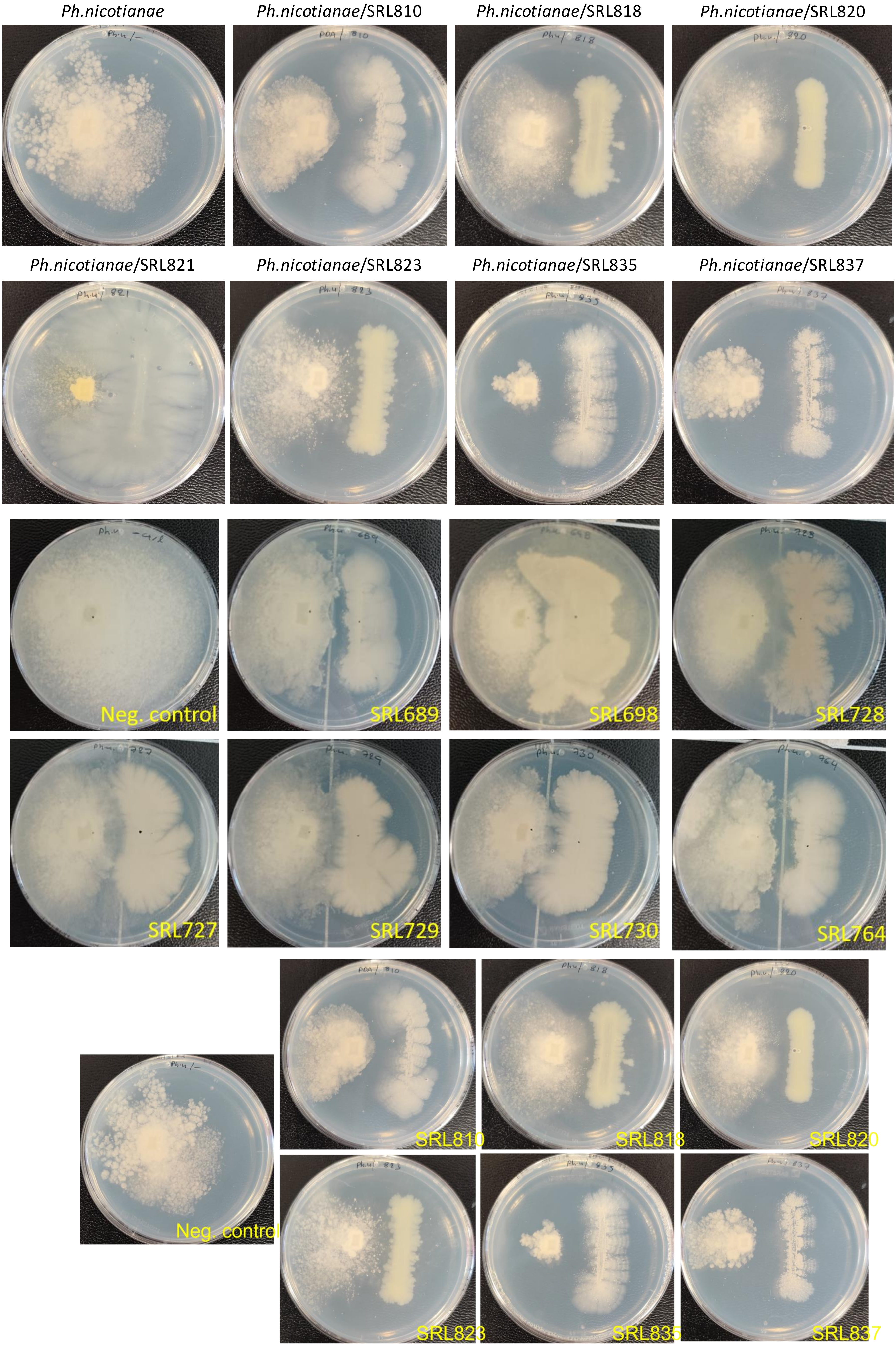

Ph.nicotianae
Ph.nicotianae/SRL810
Ph.nicotianae/SRL818
Ph.nicotianae/SRL820
Ph.nicotianae/SRL821
Ph.nicotianae/SRL823
Ph.nicotianae/SRL835
Ph.nicotianae/SRL837
Neg. control
SRL689
SRL698
SRL728
SRL727
SRL729
SRL730
SRL764
SRL810
SRL818
SRL820
Neg. control
SRL823
SRL835
SRL837

### *Verticillium dahliae*

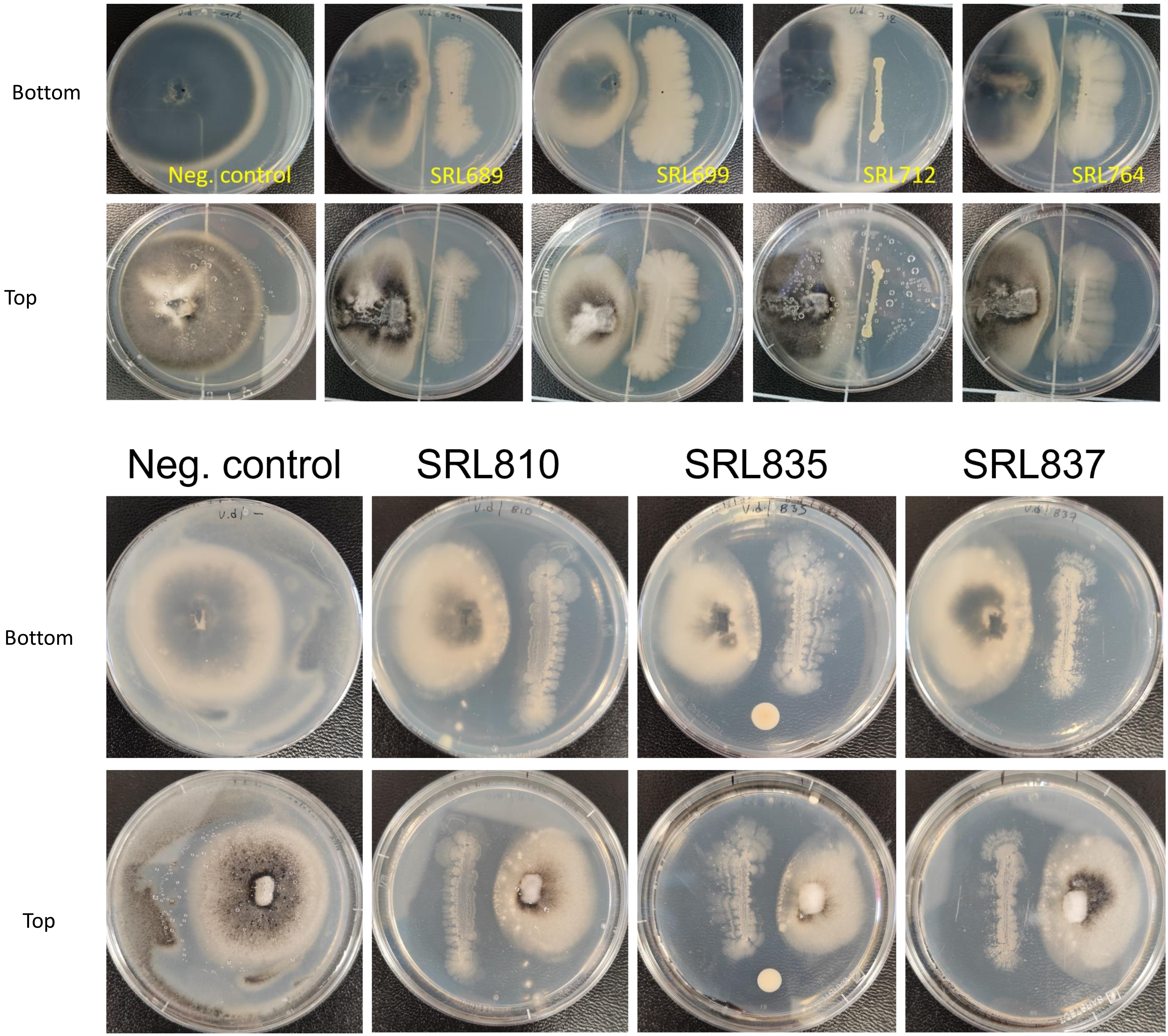
