## Supplementary figures and images for "A Thousand Meters Deep: Vertical Profiling of the Subterranean Microbes of Gourgouthakas Cave"

### Supplementary Figure 1

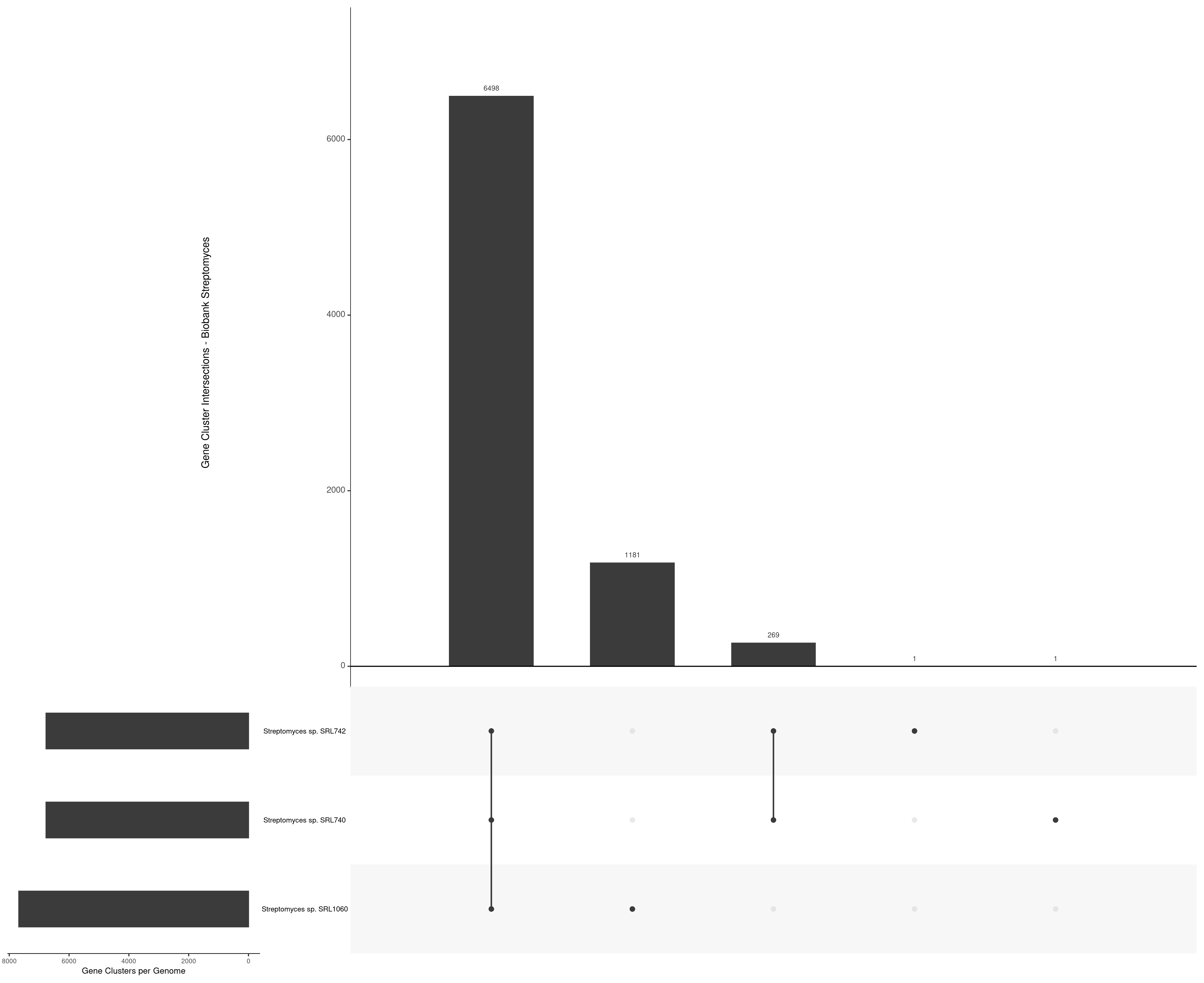
